## Supplementary Information for "Conformational restriction shapes inhibition of a multidrug efflux adaptor protein"

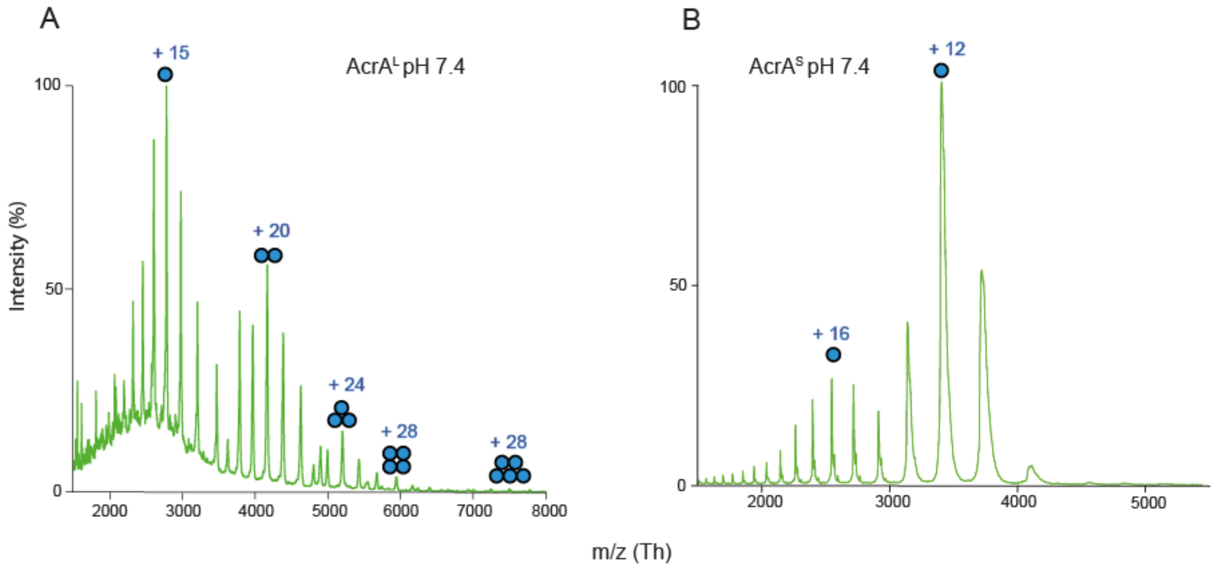


**Figure S1. Native-MS of AcrA constructs at pH 7.4. A.** Native-MS characterisation of AcrA$^{L}$ construct at pH 7.4. Protein buffer exchanged to 100 mM ammonium acetate prior to MS, in the presence of 2 x critical micelle concentration (CMC) of DDM at 0.03 %. AcrA$^{L}$ presents as a mix of oligomers up until pentamers. **B.** Native-MS characterisation of AcrA$^{S}$ construct at pH 7.4. Protein buffer exchanged to 100 mM ammonium acetate prior to MS. AcrA$^{S}$ presents as a monomer. See **Table S3** for masses.

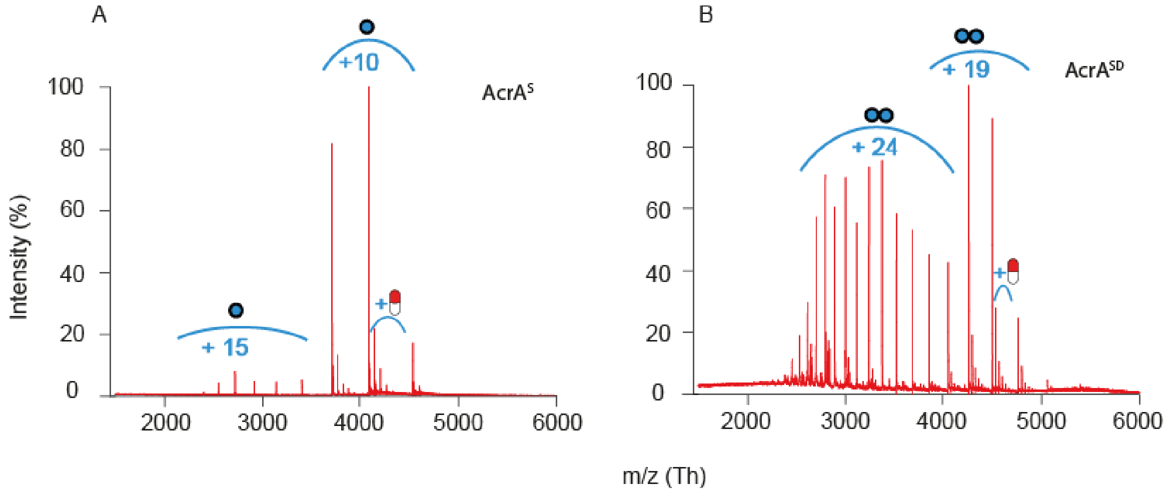


**Figure S2. Native-MS of AcrA$^{S/SD}$ constructs and novobiocin at pH 6.0. A.** Native-MS characterisation of AcrA$^{S}$ construct with novobiocin at pH 6.0. Protein buffer exchanged to 100 mM ammonium acetate prior to MS and protein diluted to 10 µM. Novobiocin added to a concentration of 30 µM, 5% DMSO final. Satellite peaks representing drug binding can be seen adjacent to peaks in the lower charge state distribution. **B.** Native-MS characterisation of AcrA$^{SD}$ construct with novobiocin at pH 6.0. Protein buffer exchanged to 100 mM ammonium acetate prior to MS and diluted to 10 µM. Novobiocin added to a concentration of 100 µM, 10% DMSO final. Satellite peaks representing drug binding can be seen adjacent to peaks in the lower charge state distribution. See **Table S4** for masses.

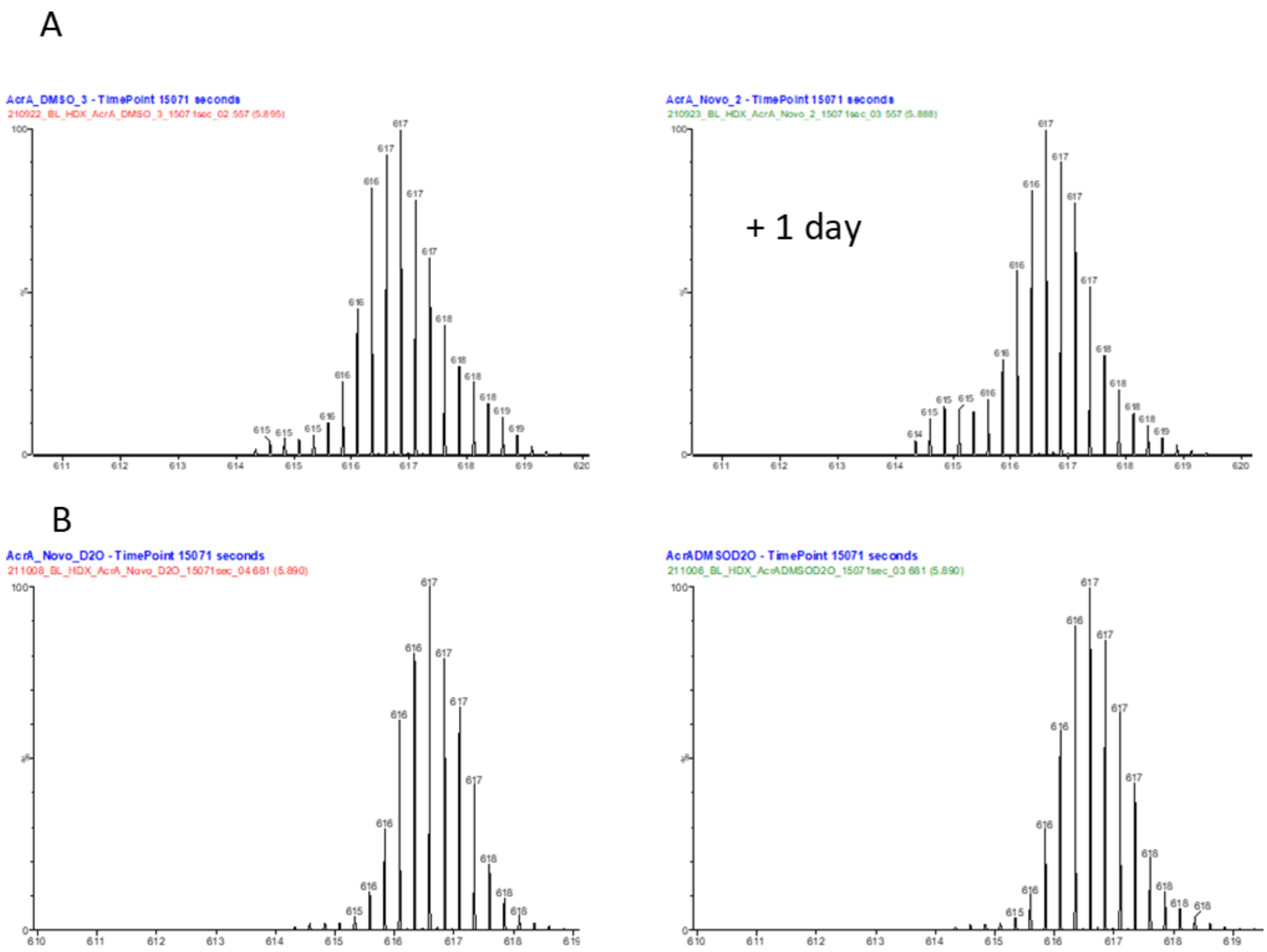


**Figure S3. Optimization of HDX-MS conditions. A.** One peptide selected throughout, after labelling in deuterated buffer for 4 hours. This represents two datasets for one protein taken one day after each other. There is an area in the low m/z (614-616 *m/z*) which is present, which represents protein aggregation/carryover. **B.** The same peptide and two subsequent datasets taken one day after each other after optimization steps. This was the addition of a SEC ‘clean-up’ stage of the protein sample before experiments and increasing pepsin washes from 2 to 3.

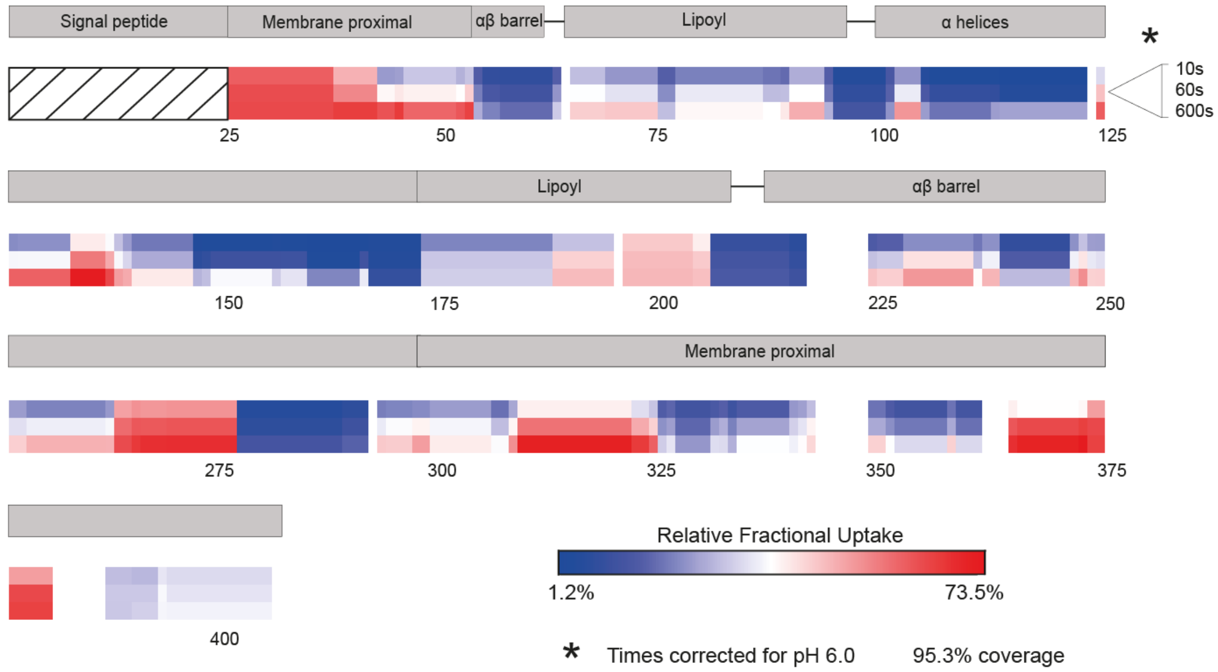


**Figure S4. Heatmap of AcrA$^{S}$ at pH 6.0.** Relative fraction uptake analysis of AcrA$^{S}$ at pH 6.0 for three time points (pH 7.4 corrected).

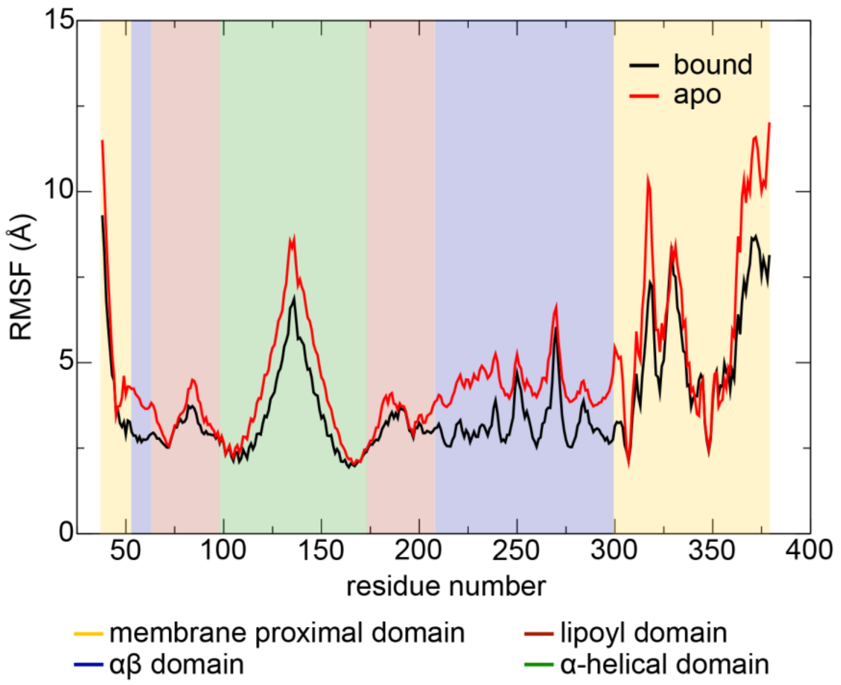


**Figure S5. Root-mean-square fluctuations (RMSF) from MD simulations of an AcrA monomer**. RMSF of AcrA for the bound (black) and apo (red) states. Curves represent averages from three 100-ns simulations. The shading indicates the domains of AcrA.

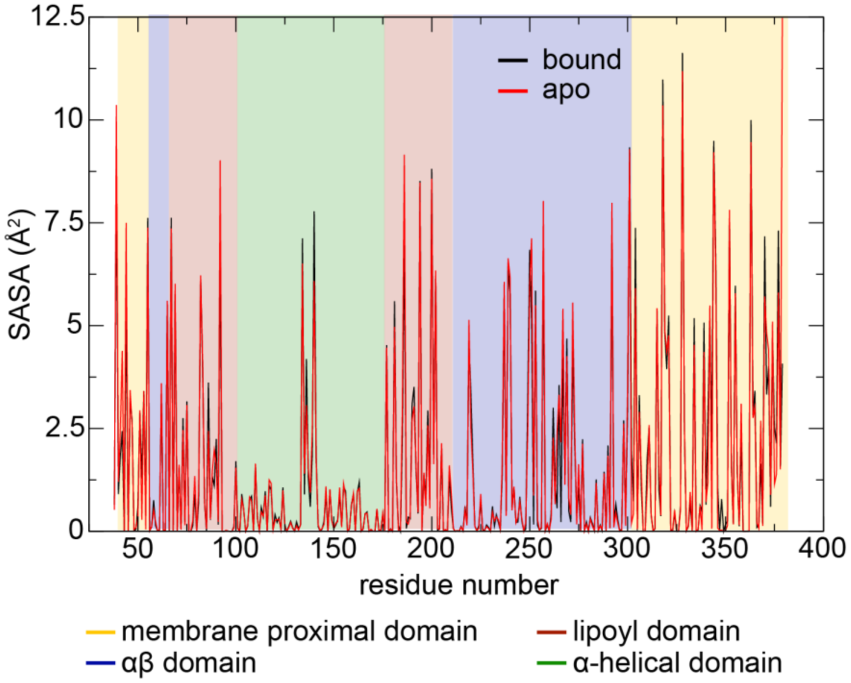


**Figure S6. Solvent accessible surface area (SASA) from MD simulations of an AcrA monomer**. SASA of AcrA for the bound (black) and apo (red) states. Curves represent averages from three 100-ns simulations. The shading indicates the domains of AcrA.

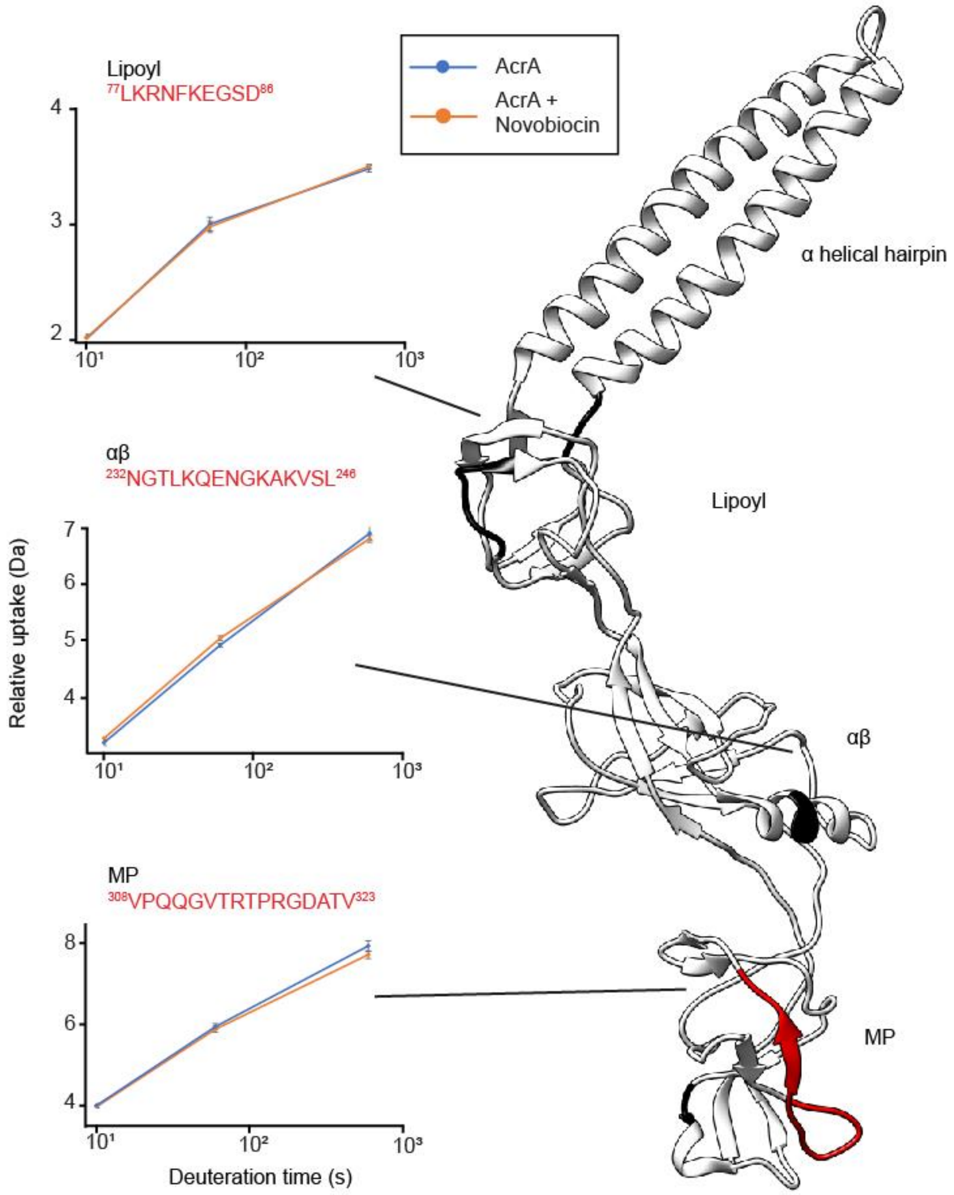


**Figure S7. The effect of novobiocin on AcrA$^S$ structural dynamics.** ΔHDX for ((AcrA$^S$ + novobiocin) – AcrA$^S$) for the latest time point is painted onto the AcrA structure (PDB:5O66) using HDeXplosion and Chimera.[1,2] We defined significance to be ≥ 0.34 Da change (see Methods) with a P-value ≤ 0.01 in a Welch's *t*-test (*n* = 4 technical replicates). White areas represent regions with insignificant ΔHDX, and black areas represent regions with no peptide coverage. Three peptide uptake plots can be shown, in areas that saw significant protection with NSC 60339. All supporting HDX-MS peptide data can be found in the Source Data file.

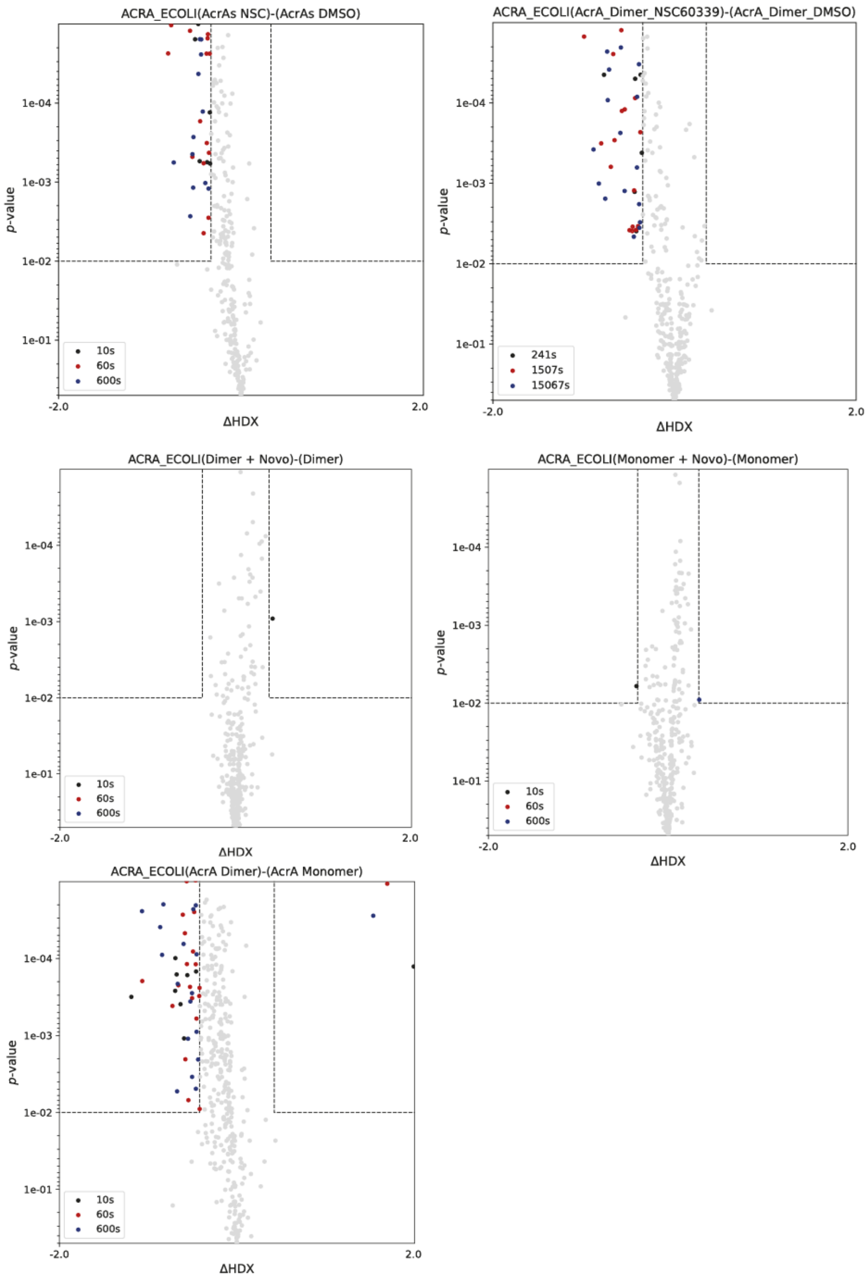


**Figure S8. Volcano plots for HDX-MS experiments**. ΔHDX cutoff values determined by individual statistical analysis (see Methods). Gray plots are insignificant, coloured spots represent significantly changed peptides at a certain time point.

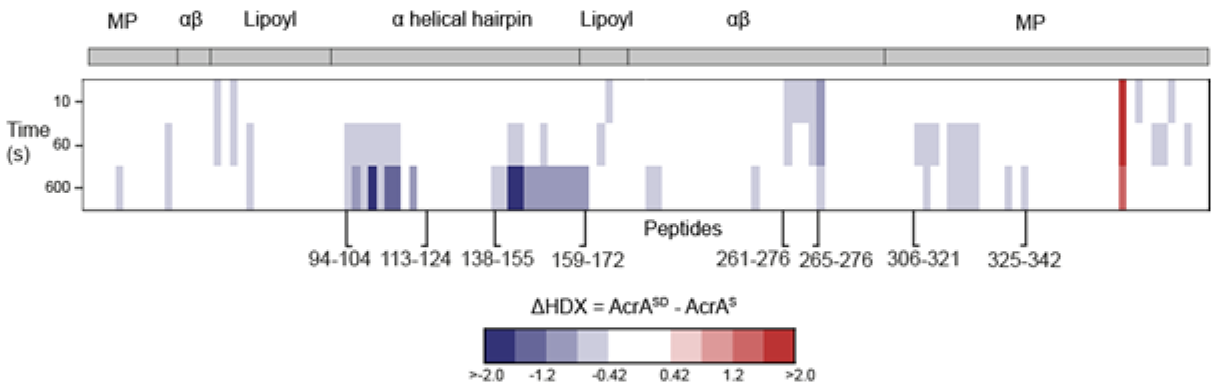


**Figure S9. Effects of dimerization on AcrA.** Chiclet plot displaying the differential HDX (ΔHDX) plots for $AcrA^{SD}$ - $AcrA^{S}$ for all time points collected. Blue signifies areas with decreased HDX between states. We defined significance to be ≥ 0.42 Da change (see Methods) with a P-value ≤ 0.01 in a Welch's *t*-test (*n* = 4 technical replicates). White areas represent regions with insignificant ΔHDX. All supporting HDX-MS peptide data can be found in the Source Data file

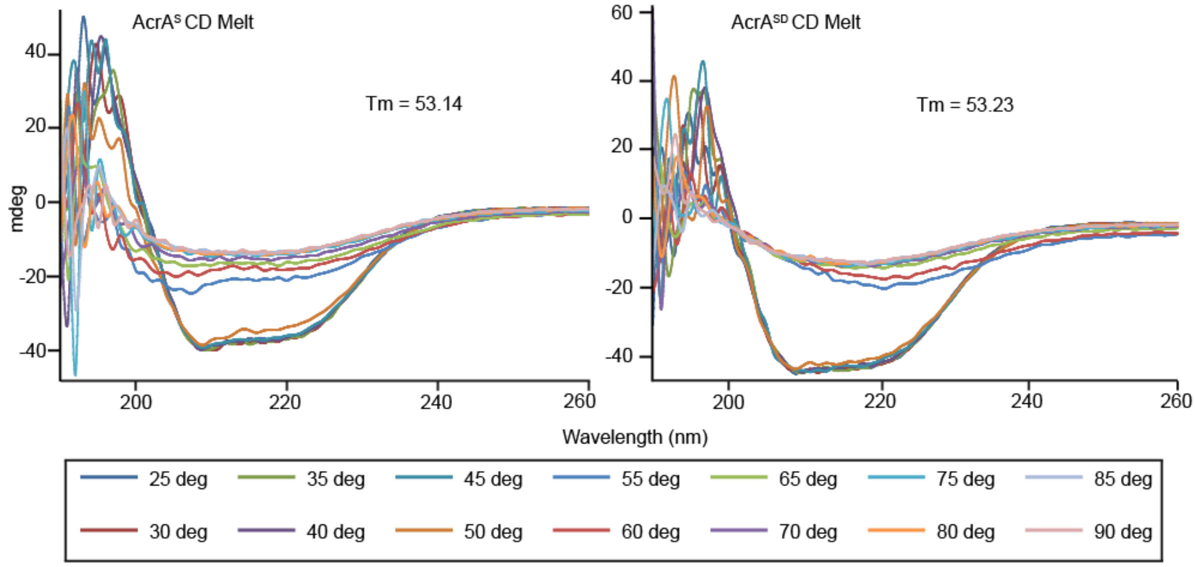


**Figure S10. Circular dichroism thermal melts of $AcrA^{S}$ and $AcrA^{SD}$.** Proteins diluted to 0.4 mg $ml^{-1}$ and CD measured from 190-260 nm in a 0.5 mm pathlength cell, from 25 ˚C to 90 ˚C, at 5 ˚C intervals. Tm calculated using the values at 222 nm.

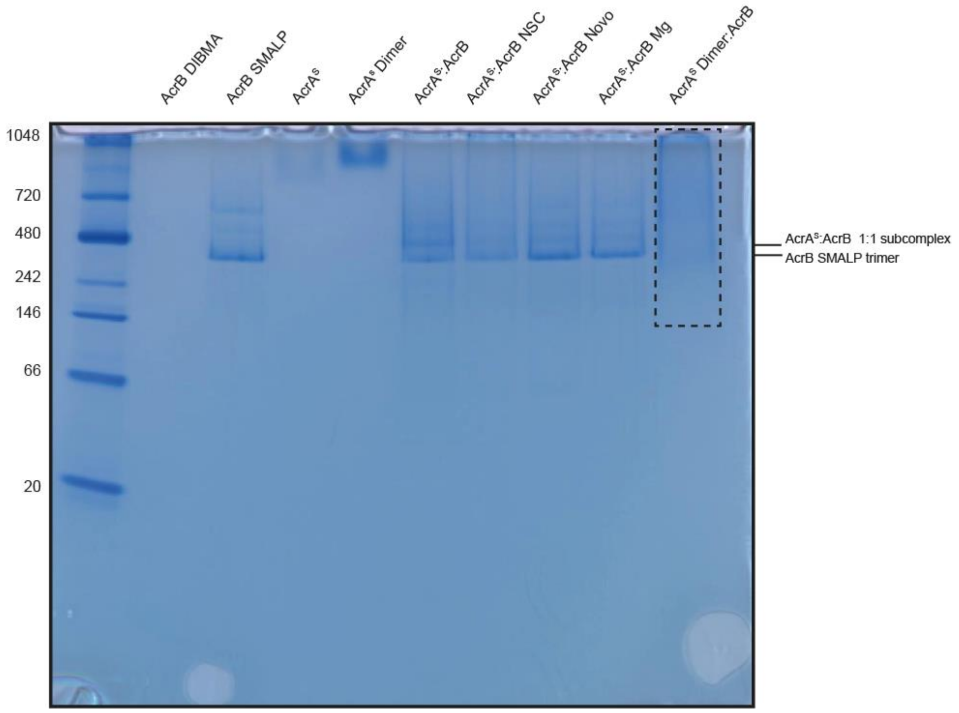


**Figure S11. SMA-PAGE of AcrA constructs and AcrB SMALPs.** Proteins loaded at 2 µM, NSC 60339 at 500 µM, novobiocin at 30 µM, $Mg^{2+}$ at 1 mM. AcrA$^S$:AcrB subcomplex can be seen at 1:1 ratio. AcrA$^{SD}$:AcrB shows various different stoichiometries, which is highlighted by the dotted box. Gel ran as described in the methods.

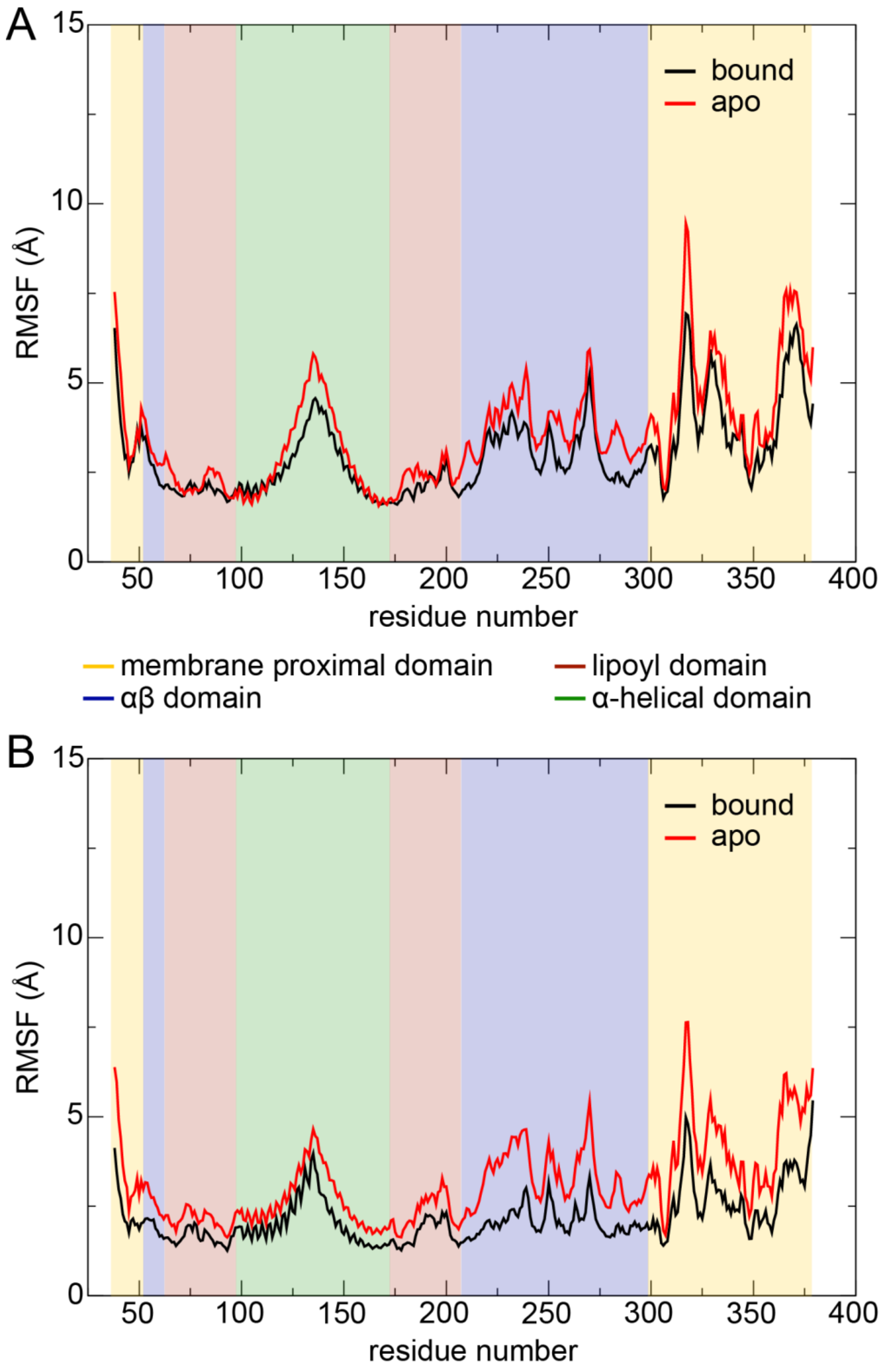


**Figure S12. RMSF from MD simulations of an AcrA dimer**. RMSF of AcrA for the bound (black) and apo (red) states. Curves represent averages from three 100-ns simulations. The shading indicates the domains of AcrA. (A) First protomer. (B) Second protomer.

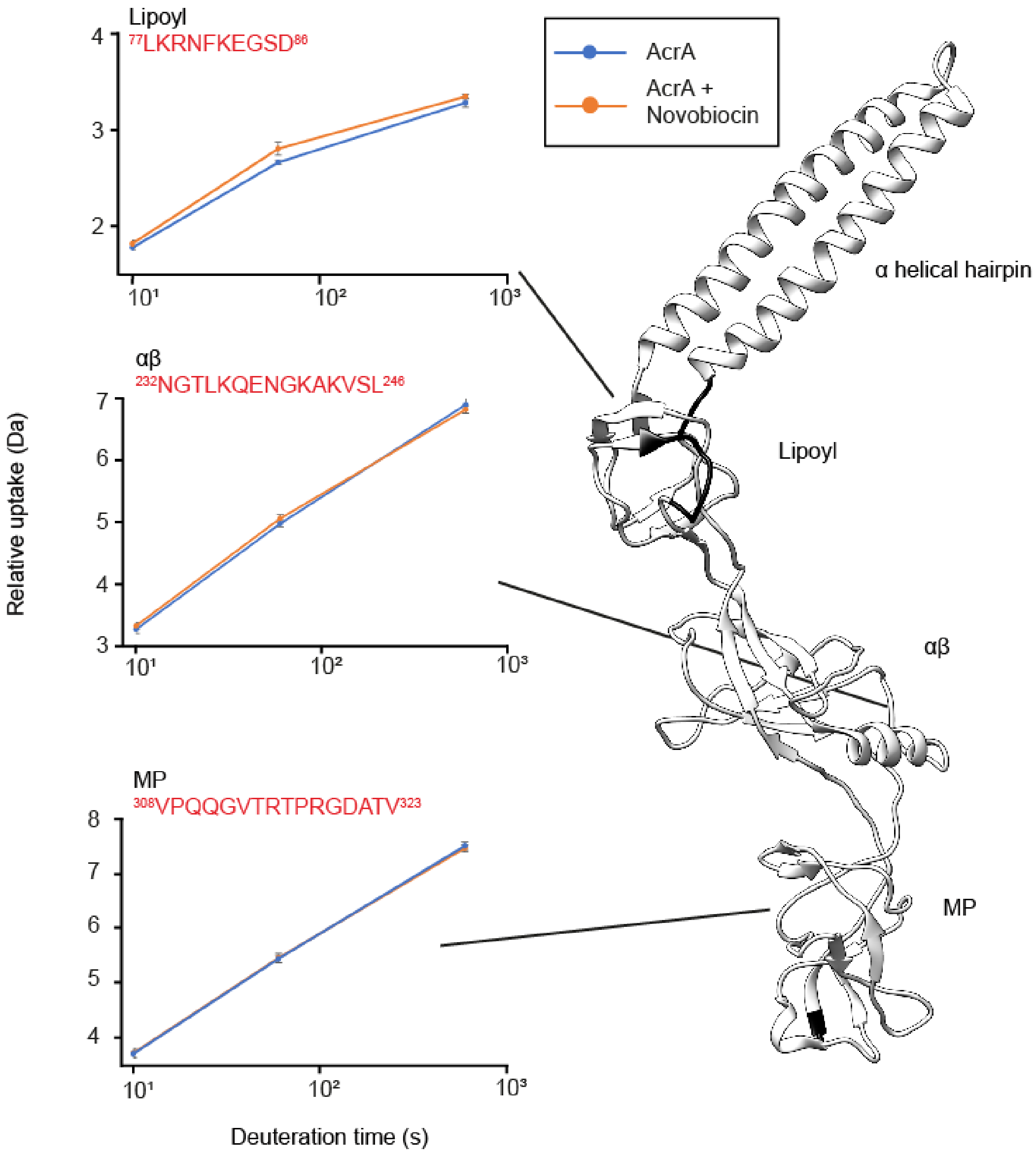


**Figure S13. The effect of novobiocin on AcrA$^{SD}$ structural dynamics. A.** Chiclet plot displaying the differential HDX (ΔHDX) plots for AcrA$^{S}$ +/- novobiocin for all time points collected. Blue signifies areas with decreased HDX between states. We defined significance to be ≥ 0.38 Da change (see Methods) using a P-value ≤ 0.01 in a Welch's *t*-test (*n* = 4 technical replicates). White areas represent regions with insignificant ΔHDX, and grey areas represent regions with no peptide coverage. All supporting HDX-MS peptide data can be found in the Source Data file. **B.** ΔHDX for ((AcrA$^{SD}$ + novobiocin) – AcrA$^{SD}$) for the latest time point is painted onto the AcrA structure (PDB:5O66) using HDeXplosion and Chimera.[1,2]

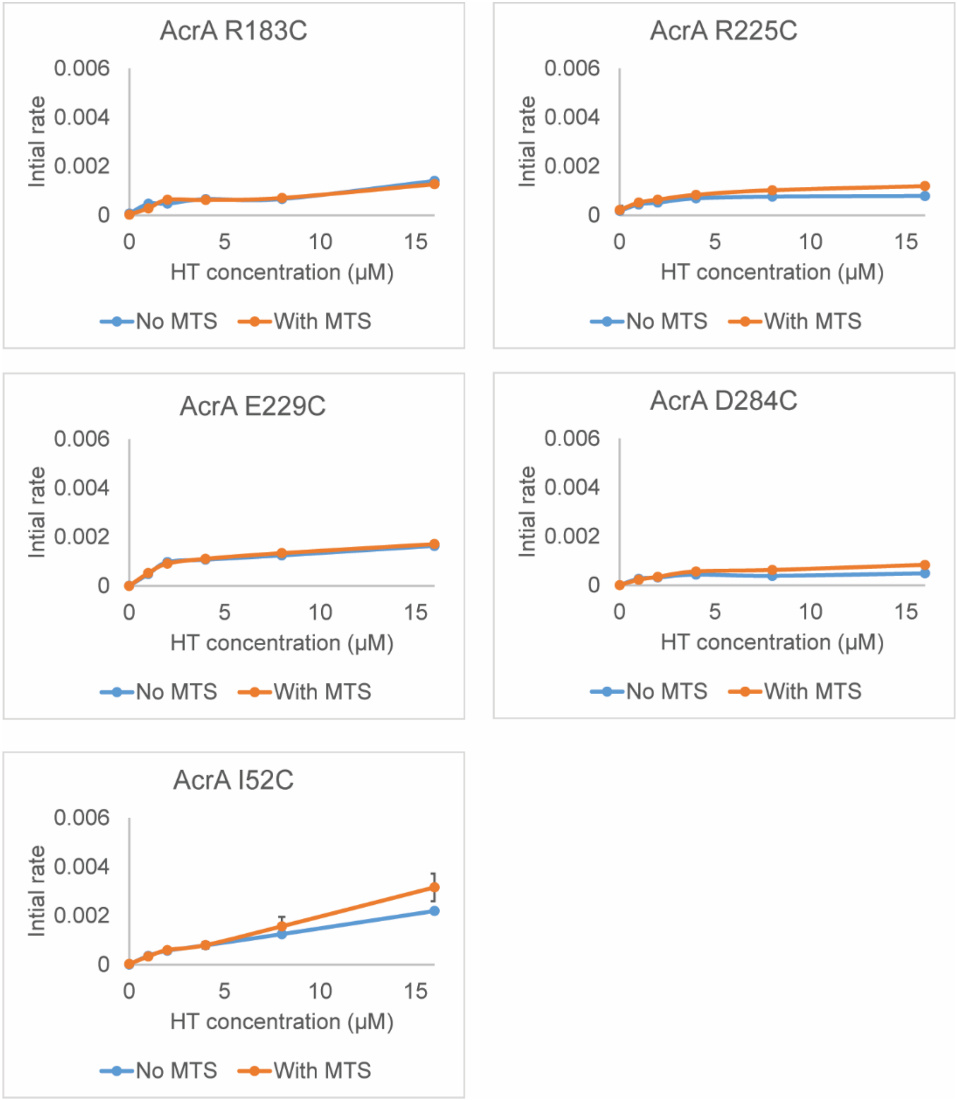


**Figure S14. The effect of Cys-reactive MTS probe on the efficiency of AcrAB-TolC.** *E. coli* Δ9-Pore cells producing AcrAB-TolC complex carrying the indicated AcrA variants were split into two aliquots and one of the aliquots was treated with a Cys-reactive probe MTS. After incubation for 15 min at 37C, cells were washed and the intracellular accumulation of Hoechst was analysed as described previously.[3] Kinetic data were fitted into a simple two-exponential model and the calculated initial rates of Hoechst accumulation (µM/s) were plotted as a function of the externally added concentration of Hoechst.

**Table 1. Primers**

| Protein | | Primer | Sequence 5'-3' |
|---|---|---|---|
| $AcrA^{SD}$ | | | |
| | AcrA (1) | F1 | GTCCGCCC ATG GCA GACGACAAACAGGCCCAACAAGGTGGC |
| | | R1 | GTCCGCGGATCC**ACCAGAAGAATTACC**AGACTTGGACTGTTC AGGCTGAGC |
| | AcrA (2) | F1 | GCGCGC GGATCC GAC GAC AAA CAG GCC CAA CAA GGT |
| | | R1 | GCGGGG CTCGAG AGACTTGGACTGTTCAGGCTGAGCA |

**Table S2. Minimal inhibitory concentrations of D9-Pore cells**

| Strains | SDS (µg/ml) | NOV (µg/ml) | ERY (µg/ml) | Van (µg/ml) | MTS (µM) |
|---|---|---|---|---|---|
| Δ9-Pore pUC18 | 8-16 | <1 | <1 | 8-16 | >200 |
| Δ9-Pore p151 | 500-1000 | 64-128 | 32-64 | 8 | >200 |
| Δ9-Pore p151 $AcrA_{L50C}B$ | 1000->1000 | 128 | 32-64 | 8 | >200 |
| Δ9-Pore p151 $AcrA_{I52C}B$ | 1000 | 128 | 32-64 | 8 | >200 |
| Δ9-Pore p151 $AcrA_{T205C}B$ | 1000->1000 | 128 | 32-64 | 16 | >200 |
| Δ9-Pore p151 $AcrA_{R225C}B$ | 1000 | 64 | 32 | 16 | >200 |
| Δ9-Pore p151 $AcrA_{E229C}B$ | 1000 | 64 | 32 | 16 | >200 |
| Δ9-Pore p151 $AcrA_{N232C}B$ | 500 | 64 | 32 | 8 | >200 |
| Δ9-Pore p151 $AcrA_{D284C}B$ | >1000 | 64 | 32 | 8 | >200 |
| Δ9-Pore p151 $AcrA_{R183C}B$ | >1000 | 64 | 32 | 16 | >200 |

**Table S3. Native-MS masses table 1.** Reported is the standard error of the mean within a single spectrum. Positive mass differences can be attributed to salt and/or detergent adducts.

| | **Measured mass (Da)** | **Standard Error (± Da)** | **Theoretical mass†,* (Da)** | **Mass difference (Da)** |
|---|---|---|---|---|
| **AcrA$^{l}$ pH 6.0 Monomer** | 41,627 | 8 | 41,624 | 3 |
| **AcrA$^{l}$ pH 6.0 Dimer** | 83,221 | 2 | 83,248 | -27 |
| **AcrA$^{l}$ pH 7.4 Monomer** | 41,632 | 7 | 41,624 | 8 |
| **AcrA$^{l}$ pH 7.4 Dimer** | 83,274 | 8 | 83,248 | 26 |
| **AcrA$^{l}$ pH 7.4 Trimer** | 124,879 | 10 | 124,872 | 7 |
| **AcrA$^{l}$ pH 7.4 Tetramer** | 166,485 | 25 | 166,496 | -11 |
| **AcrA$^{l}$ pH 7.4 Pentamer** | 210,023 | 33 | 208,120 | 1903 |
| **AcrA$^{s}$ pH 6.0 Monomer** | 40,849 | 2 | 40,817 | 32 |
| **AcrA$^{s}$ pH 7.4 Monomer** | 40,846 | 2 | 40,817 | 29 |
| ***AcrA$^{SD}$ pH 6.0** | 81,005 | 4 | 80969 | 36 |

†*The theoretical masses for AcrA$^{l}$ were calculated for AcrA modified with N-acyl-S-diacylglycerol containing two palmitoyl residues and one oleoyl residue.*

**Theoretical masses for AcrA$^{S}$ construct were amended for fMet processing.*

**Table S4. Native-MS masses table 2.** Reported is the standard error of the mean within a single spectrum. Positive mass differences can be attributed to salt and/or detergent adducts. Novobiocin mass = 613 Da.

| | **Measured mass (Da)** | **Standard Error (± Da)** | **Mass difference (Da)** |
|---|---|---|---|
| **AcrA$^{S}$ pH 6.0** | 40,841 | 2 | - |
| **+ Novobiocin pH 6.0** | 41,458 | 3 | 617 |
| **+ Novobiocin pH 6.0 (x2)** | 42,067 | 4 | 1226 |
| ***AcrA$^{SD}$ pH 6.0** | 81,000 | 3 | - |
| **+ Novobiocin pH 6.0** | 81,613 | 10 | 613 |
| **+ Novobiocin pH 6.0 (x2)** | 82,228 | 4 | 1228 |
| **+ Novobiocin pH 6.0 (x3)** | 82,954 | 2 | 1954 |
