## Supplementary material for "Conformational restriction shapes inhibition of a multidrug efflux adaptor protein": UptakePlots1

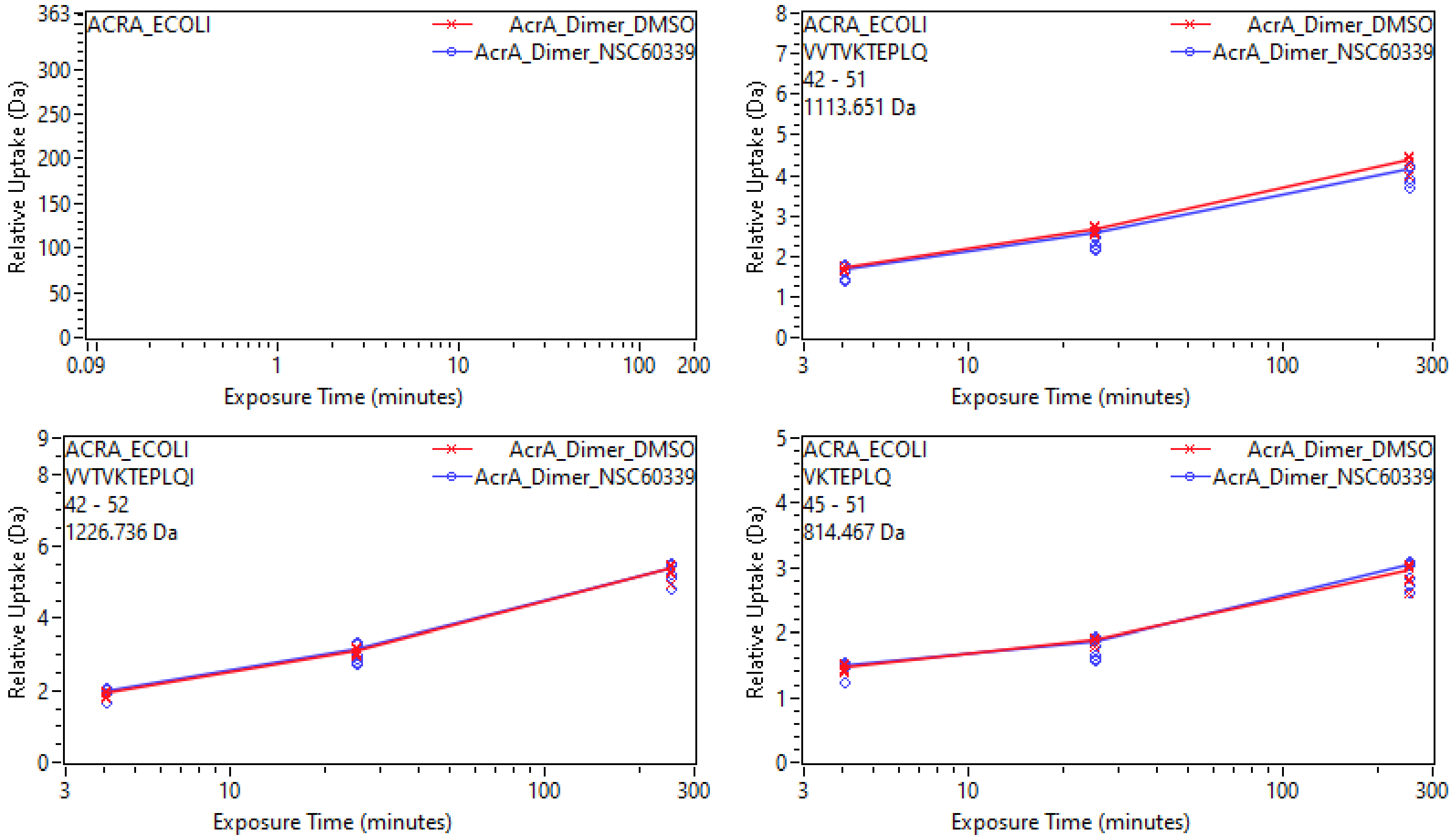
ACRA_ECOLI
AcrA_Dimer_DMSO
AcrA_Dimer_NSC60339
Relative Uptake (Da)
Exposure Time (minutes)
ACRA_ECOLI
VVTVKTEPLQ
42 - 51
1113.651 Da
ACRA_ECOLI
VVTVKTEPLQI
42 - 52
1226.736 Da
ACRA_ECOLI
VKTEPLQ
45 - 51
814.467 Da

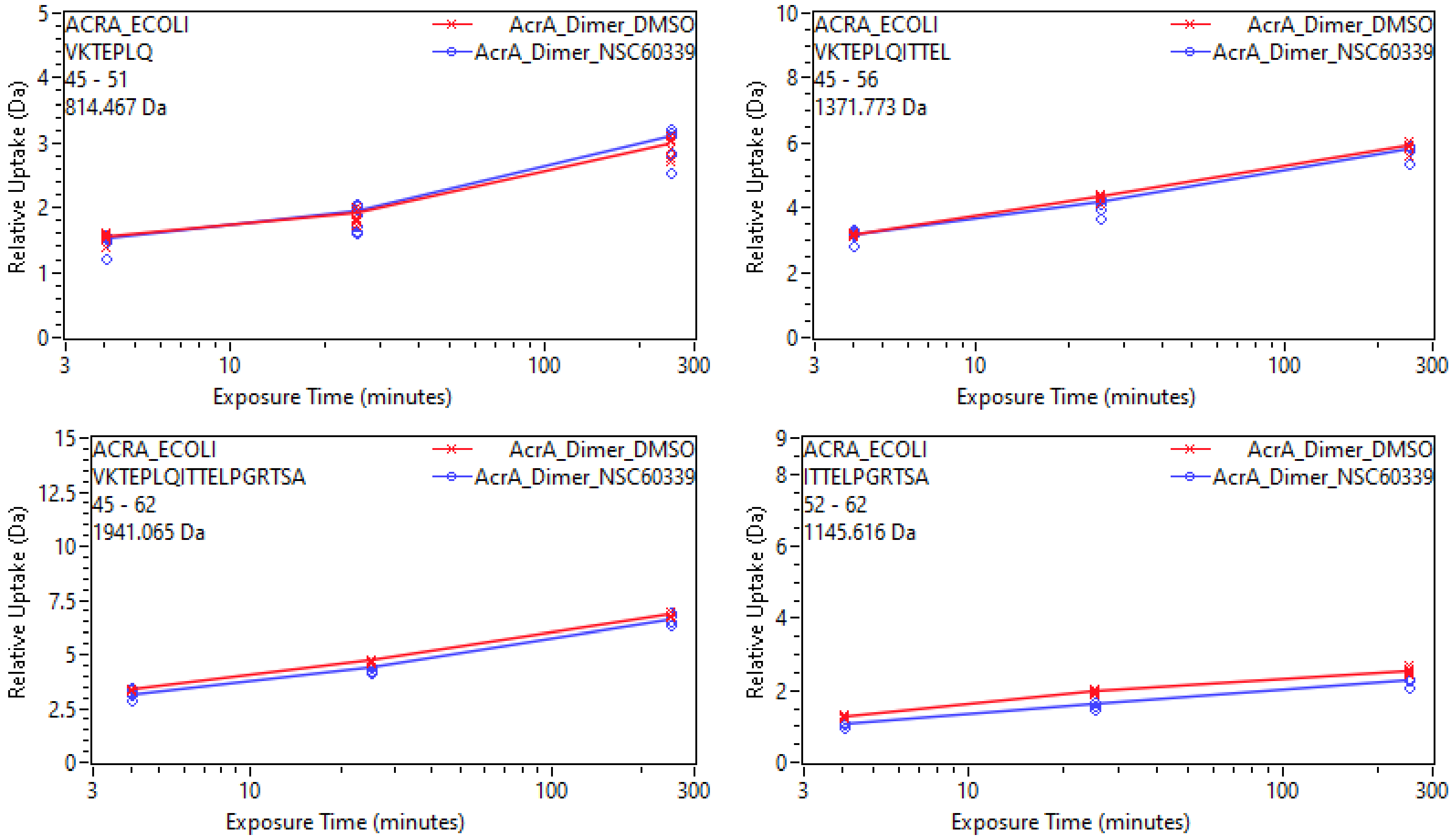
ACRA_ECOLI
VKTEPLQ
45 - 51
814.467 Da
AcrA_Dimer_DMSO
AcrA_Dimer_NSC60339
Relative Uptake (Da)
Exposure Time (minutes)
ACRA_ECOLI
VKTEPLQITTEL
45 - 56
1371.773 Da
ACRA_ECOLI
VKTEPLQITTELPGRTSA
45 - 62
1941.065 Da
ACRA_ECOLI
ITTELPGRTSA
52 - 62
1145.616 Da

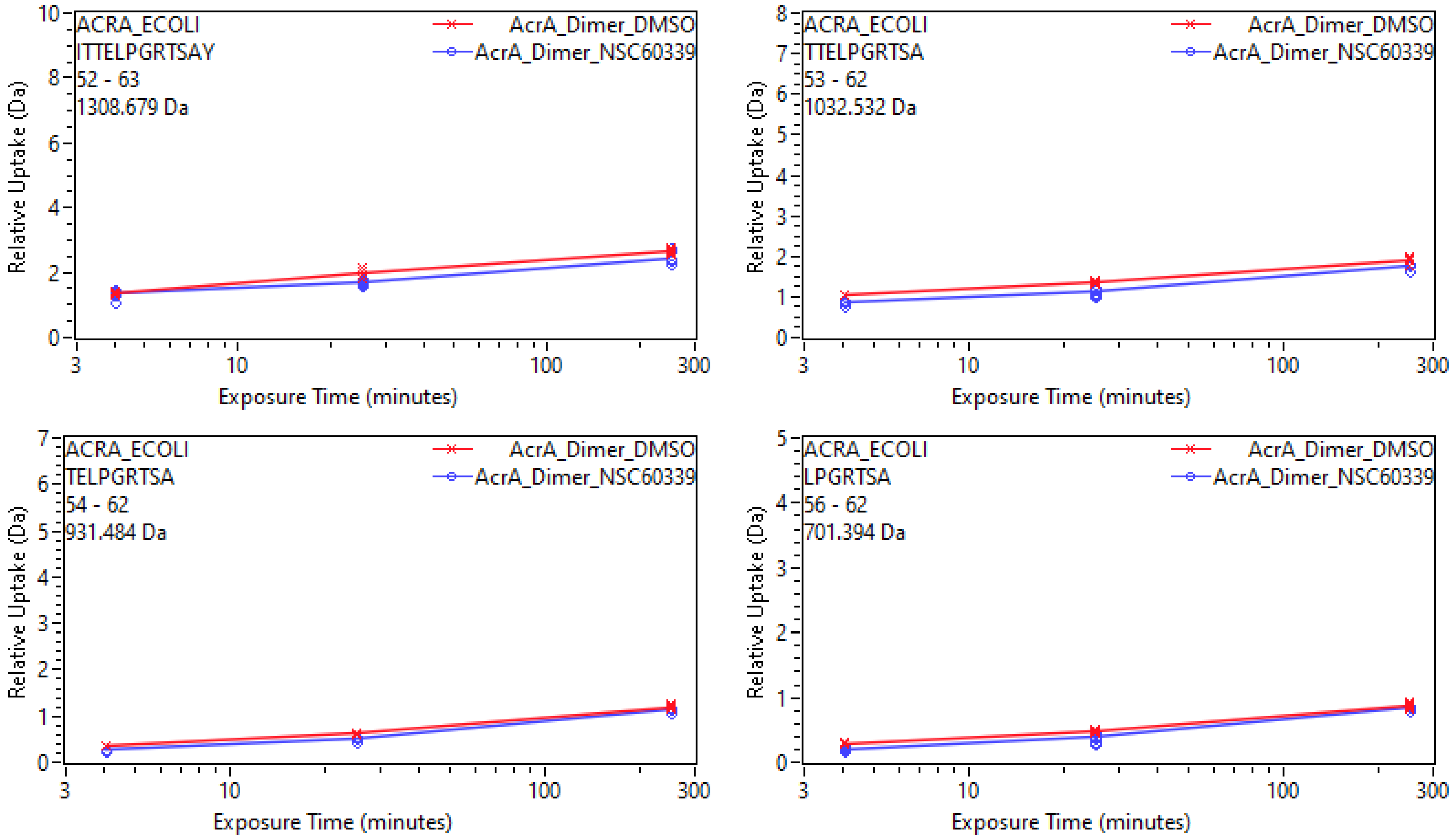
ACRA_ECOLI
ITTELPGRTSAY
52 - 63
1308.679 Da
AcrA_Dimer_DMSO
AcrA_Dimer_NSC60339
Relative Uptake (Da)
Exposure Time (minutes)
ACRA_ECOLI
TTELPGRTSA
53 - 62
1032.532 Da
ACRA_ECOLI
TELPGRTSA
54 - 62
931.484 Da
ACRA_ECOLI
LPGRTSA
56 - 62
701.394 Da

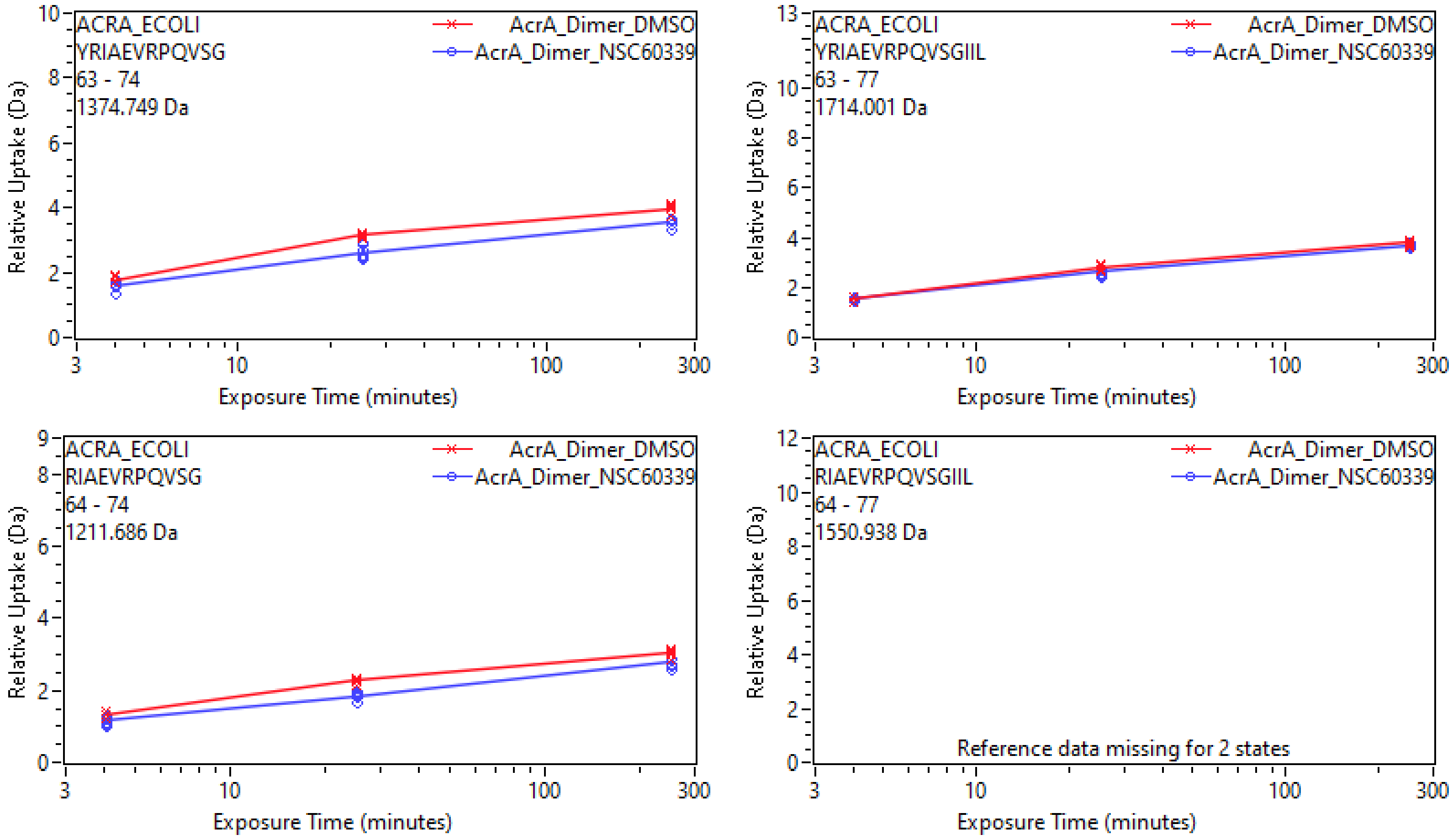
ACRA_ECOLI
YRIAEVRPQVSG
63 - 74
1374.749 Da
AcrA_Dimer_DMSO
AcrA_Dimer_NSC60339
Relative Uptake (Da)
Exposure Time (minutes)
ACRA_ECOLI
YRIAEVRPQVSGIIL
63 - 77
1714.001 Da
AcrA_Dimer_DMSO
AcrA_Dimer_NSC60339
Relative Uptake (Da)
Exposure Time (minutes)
ACRA_ECOLI
RIAEVRPQVSG
64 - 74
1211.686 Da
AcrA_Dimer_DMSO
AcrA_Dimer_NSC60339
Relative Uptake (Da)
Exposure Time (minutes)
ACRA_ECOLI
RIAEVRPQVSGIIL
64 - 77
1550.938 Da
AcrA_Dimer_DMSO
AcrA_Dimer_NSC60339
Reference data missing for 2 states
Relative Uptake (Da)
Exposure Time (minutes)

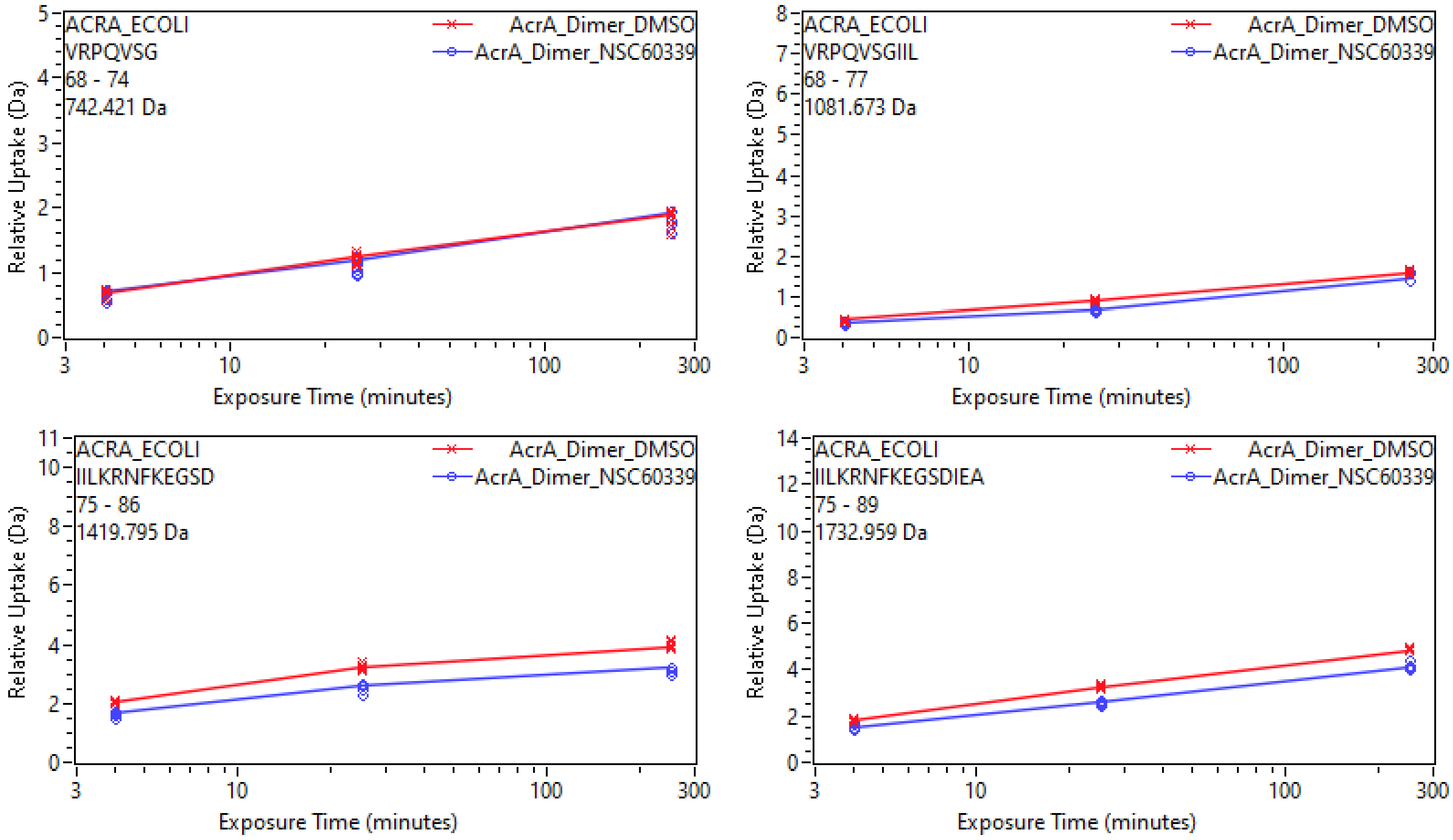

ACRA_ECOLI
VRPQVSG
68 - 74
742.421 Da
AcrA_Dimer_DMSO
AcrA_Dimer_NSC60339
Relative Uptake (Da)
Exposure Time (minutes)
ACRA_ECOLI
VRPQVSGIIL
68 - 77
1081.673 Da
ACRA_ECOLI
IILKRNFKEGSD
75 - 86
1419.795 Da
ACRA_ECOLI
IILKRNFKEGSDIEA
75 - 89
1732.959 Da

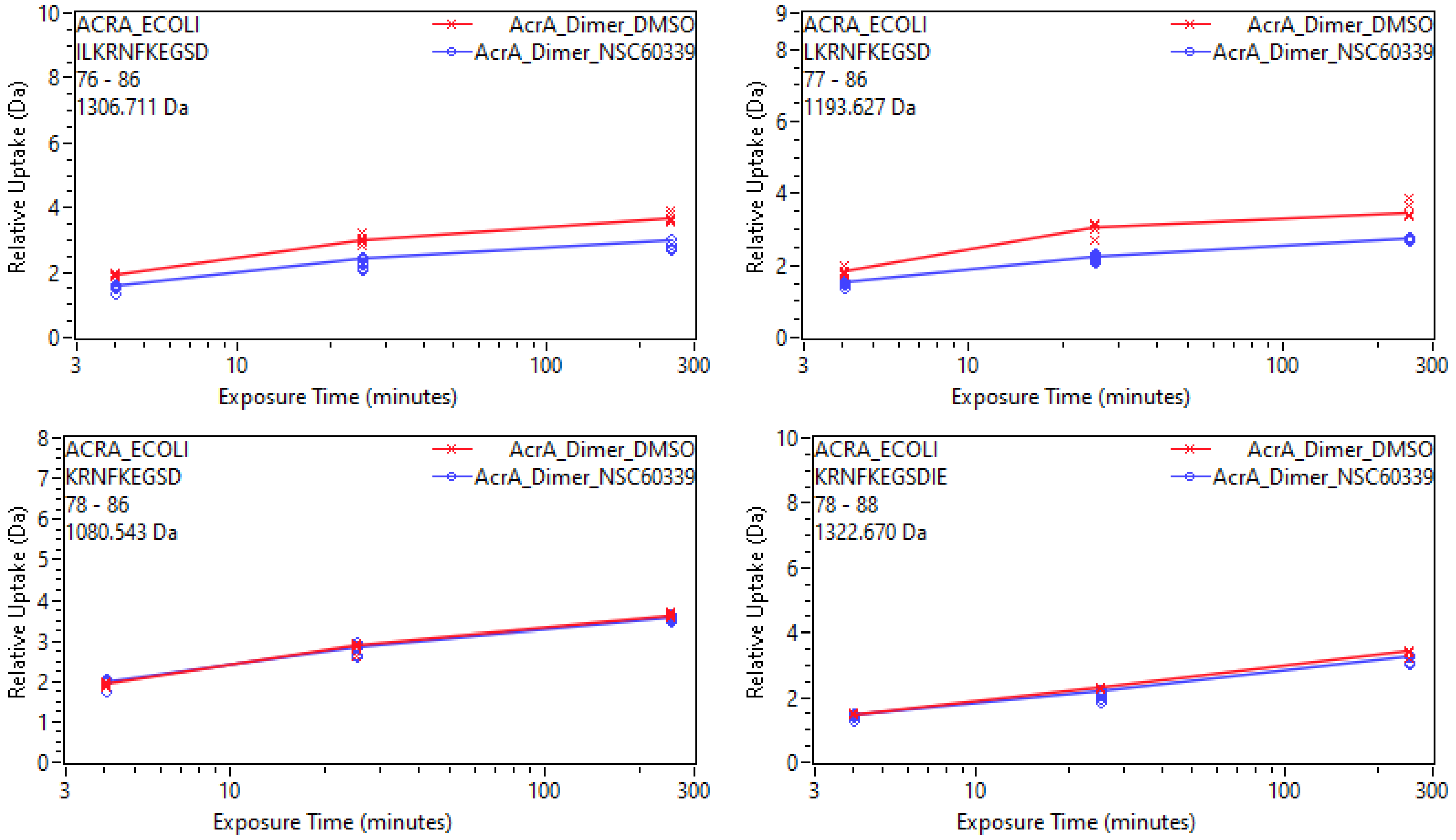
ACRA_ECOLI
ILKRNFKEGSD
76 - 86
1306.711 Da
AcrA_Dimer_DMSO
AcrA_Dimer_NSC60339
Relative Uptake (Da)
Exposure Time (minutes)
ACRA_ECOLI
LKRNFKEGSD
77 - 86
1193.627 Da
AcrA_Dimer_DMSO
AcrA_Dimer_NSC60339
Relative Uptake (Da)
Exposure Time (minutes)
ACRA_ECOLI
KRNFKEGSD
78 - 86
1080.543 Da
AcrA_Dimer_DMSO
AcrA_Dimer_NSC60339
Relative Uptake (Da)
Exposure Time (minutes)
ACRA_ECOLI
KRNFKEGSDIE
78 - 88
1322.670 Da
AcrA_Dimer_DMSO
AcrA_Dimer_NSC60339
Relative Uptake (Da)
Exposure Time (minutes)

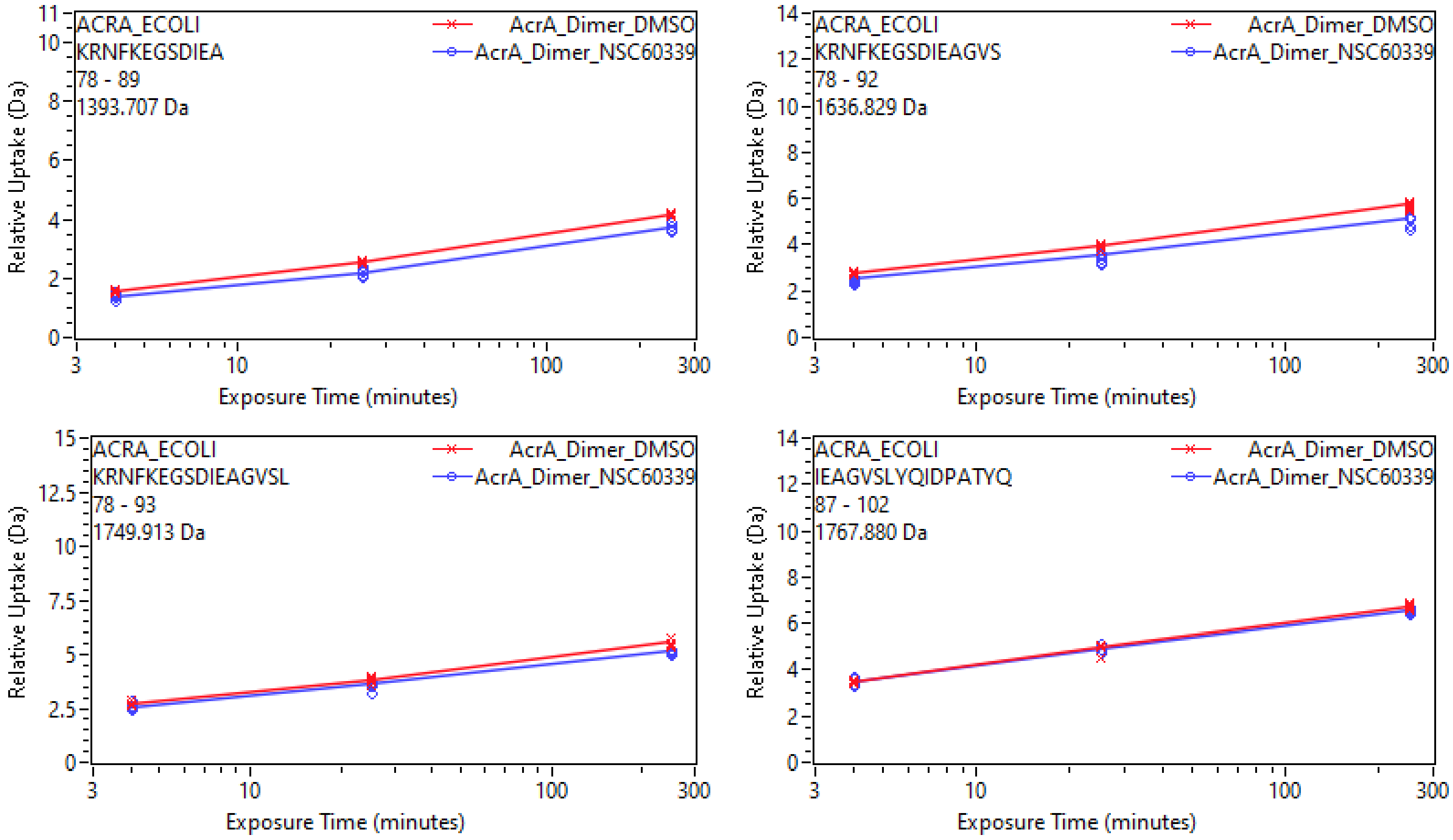

ACRA_ECOLI
KRNFKEGSDIEA
78 - 89
1393.707 Da
AcrA_Dimer_DMSO
AcrA_Dimer_NSC60339
Relative Uptake (Da)
Exposure Time (minutes)
ACRA_ECOLI
KRNFKEGSDIEAGVS
78 - 92
1636.829 Da
AcrA_Dimer_DMSO
AcrA_Dimer_NSC60339
Relative Uptake (Da)
Exposure Time (minutes)
ACRA_ECOLI
KRNFKEGSDIEAGVSL
78 - 93
1749.913 Da
AcrA_Dimer_DMSO
AcrA_Dimer_NSC60339
Relative Uptake (Da)
Exposure Time (minutes)
ACRA_ECOLI
IEAGVSLYQIDPATYQ
87 - 102
1767.880 Da
AcrA_Dimer_DMSO
AcrA_Dimer_NSC60339
Relative Uptake (Da)
Exposure Time (minutes)

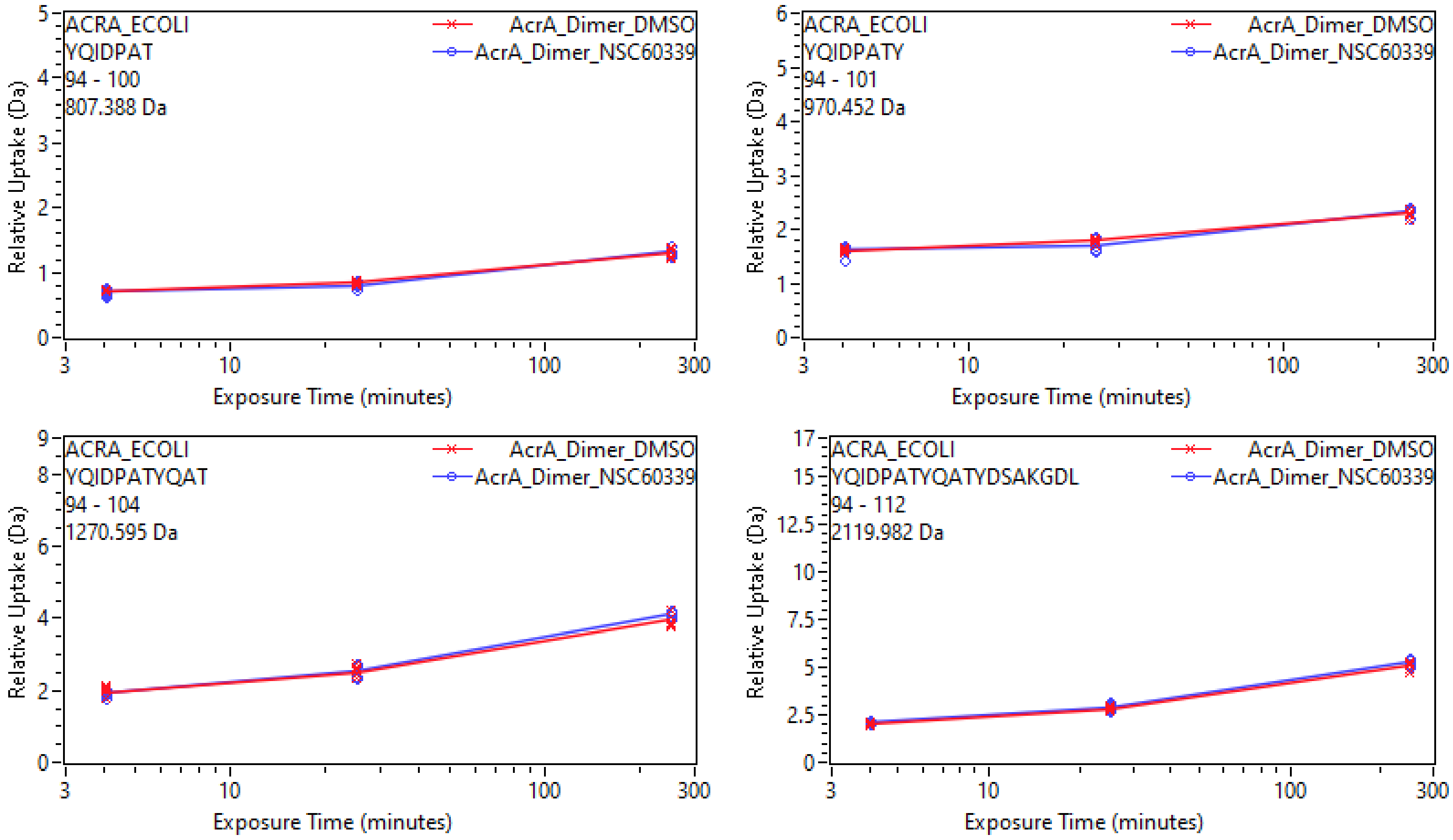
ACRA_ECOLI
YQIDPAT
94 - 100
807.388 Da
AcrA_Dimer_DMSO
AcrA_Dimer_NSC60339
Relative Uptake (Da)
Exposure Time (minutes)
ACRA_ECOLI
YQIDPATY
94 - 101
970.452 Da
ACRA_ECOLI
YQIDPATYQAT
94 - 104
1270.595 Da
ACRA_ECOLI
YQIDPATYQATYDSAKGDL
94 - 112
2119.982 Da

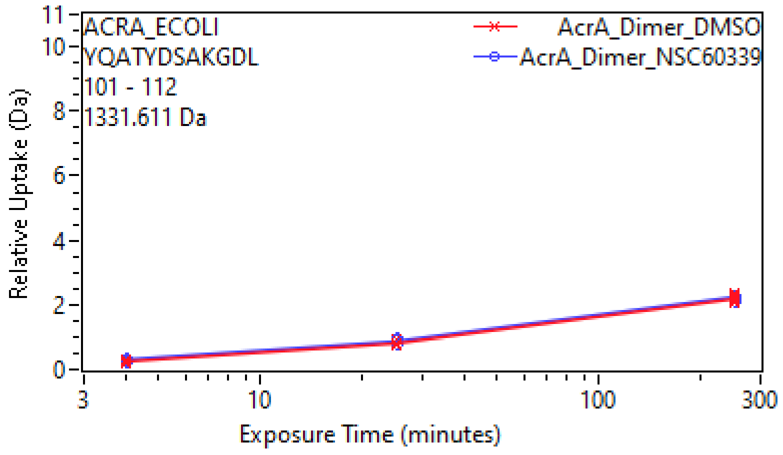
ACRA_ECOLI
YQATYDSAKGDL
101 - 112
1331.611 Da
AcrA_Dimer_DMSO
AcrA_Dimer_NSC60339
Relative Uptake (Da)
Exposure Time (minutes)

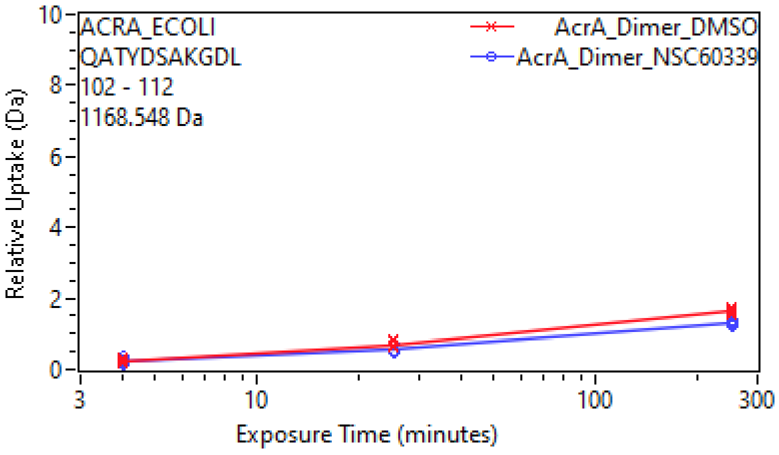
ACRA_ECOLI
QATYDSAKGDL
102 - 112
1168.548 Da
AcrA_Dimer_DMSO
AcrA_Dimer_NSC60339
Relative Uptake (Da)
Exposure Time (minutes)

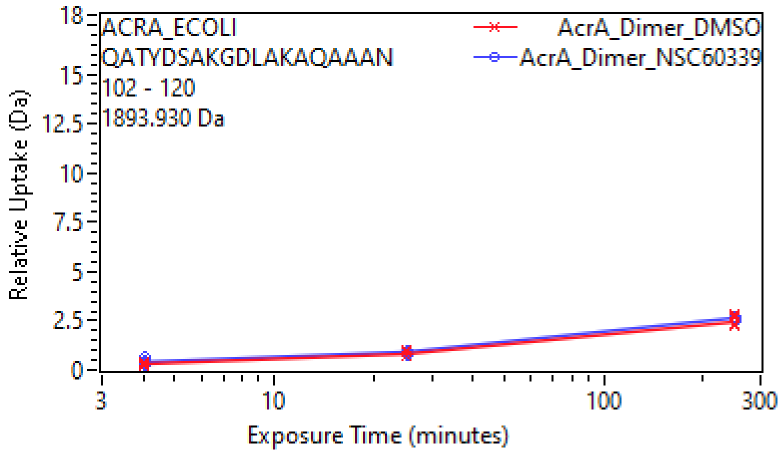
ACRA_ECOLI
QATYDSAKGDLAKAQAAAN
102 - 120
1893.930 Da
AcrA_Dimer_DMSO
AcrA_Dimer_NSC60339
Relative Uptake (Da)
Exposure Time (minutes)

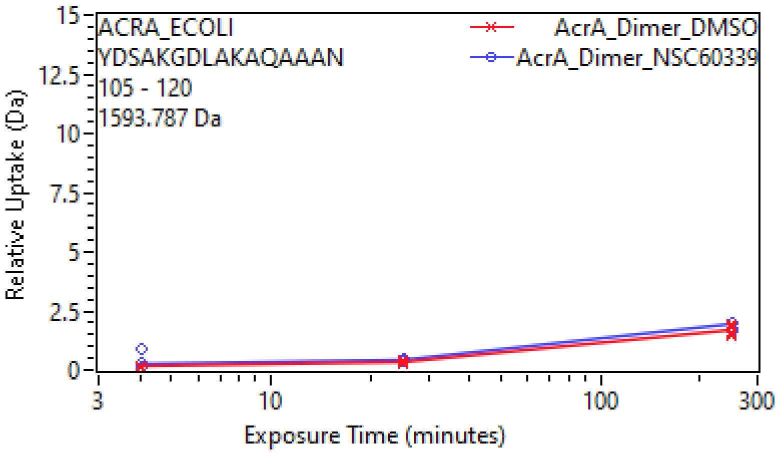
ACRA_ECOLI
YDSAKGDLAKAQAAAN
105 - 120
1593.787 Da
AcrA_Dimer_DMSO
AcrA_Dimer_NSC60339
Relative Uptake (Da)
Exposure Time (minutes)

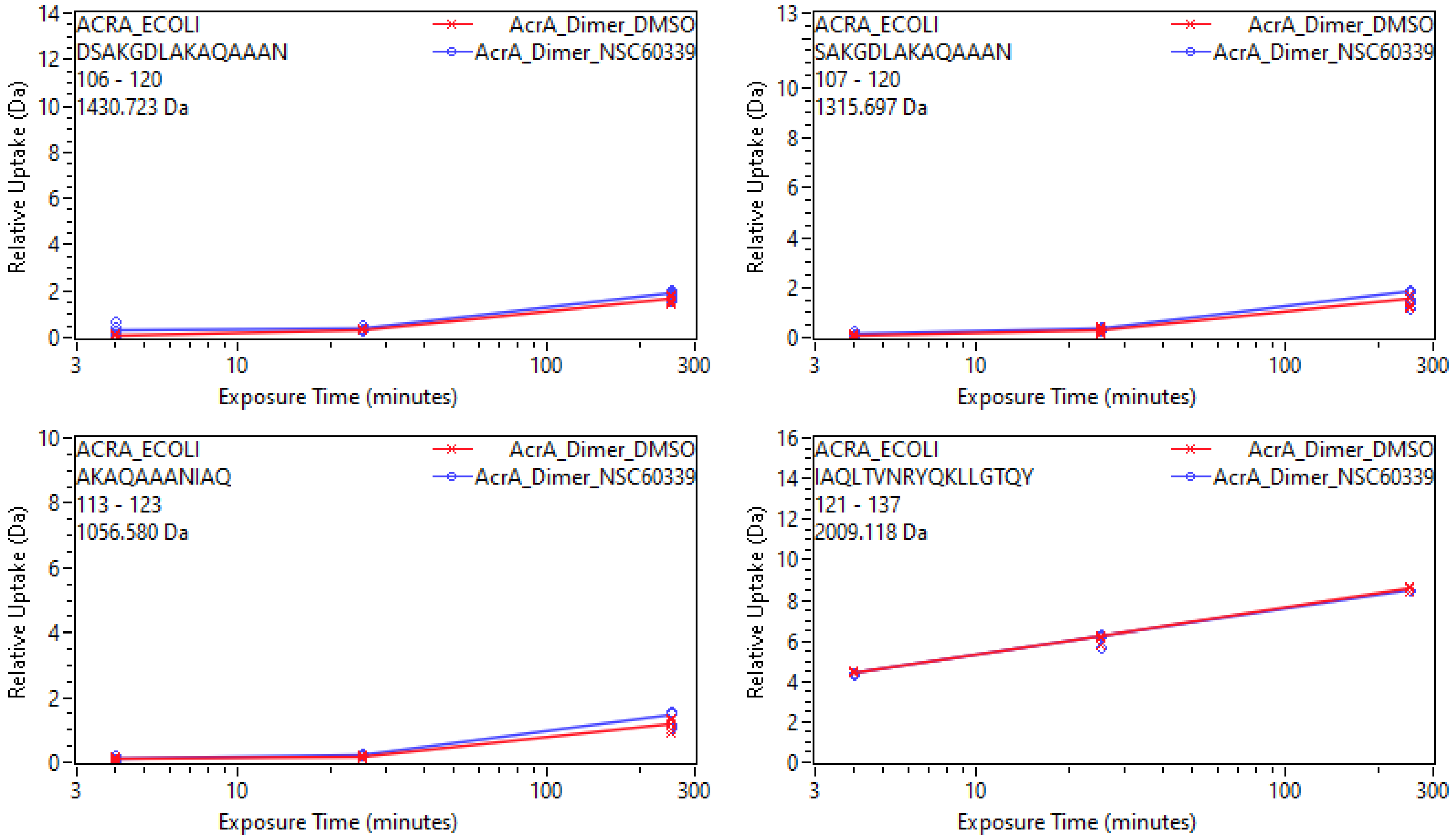
ACRA_ECOLI
DSAKGDLAKAQAAAN
106 - 120
1430.723 Da
AcrA_Dimer_DMSO
AcrA_Dimer_NSC60339
Relative Uptake (Da)
Exposure Time (minutes)
ACRA_ECOLI
SAKGDLAKAQAAAN
107 - 120
1315.697 Da
AcrA_Dimer_DMSO
AcrA_Dimer_NSC60339
Relative Uptake (Da)
Exposure Time (minutes)
ACRA_ECOLI
AKAQAAANIAQ
113 - 123
1056.580 Da
AcrA_Dimer_DMSO
AcrA_Dimer_NSC60339
Relative Uptake (Da)
Exposure Time (minutes)
ACRA_ECOLI
IAQLTVNRYQKLLGTQY
121 - 137
2009.118 Da
AcrA_Dimer_DMSO
AcrA_Dimer_NSC60339
Relative Uptake (Da)
Exposure Time (minutes)

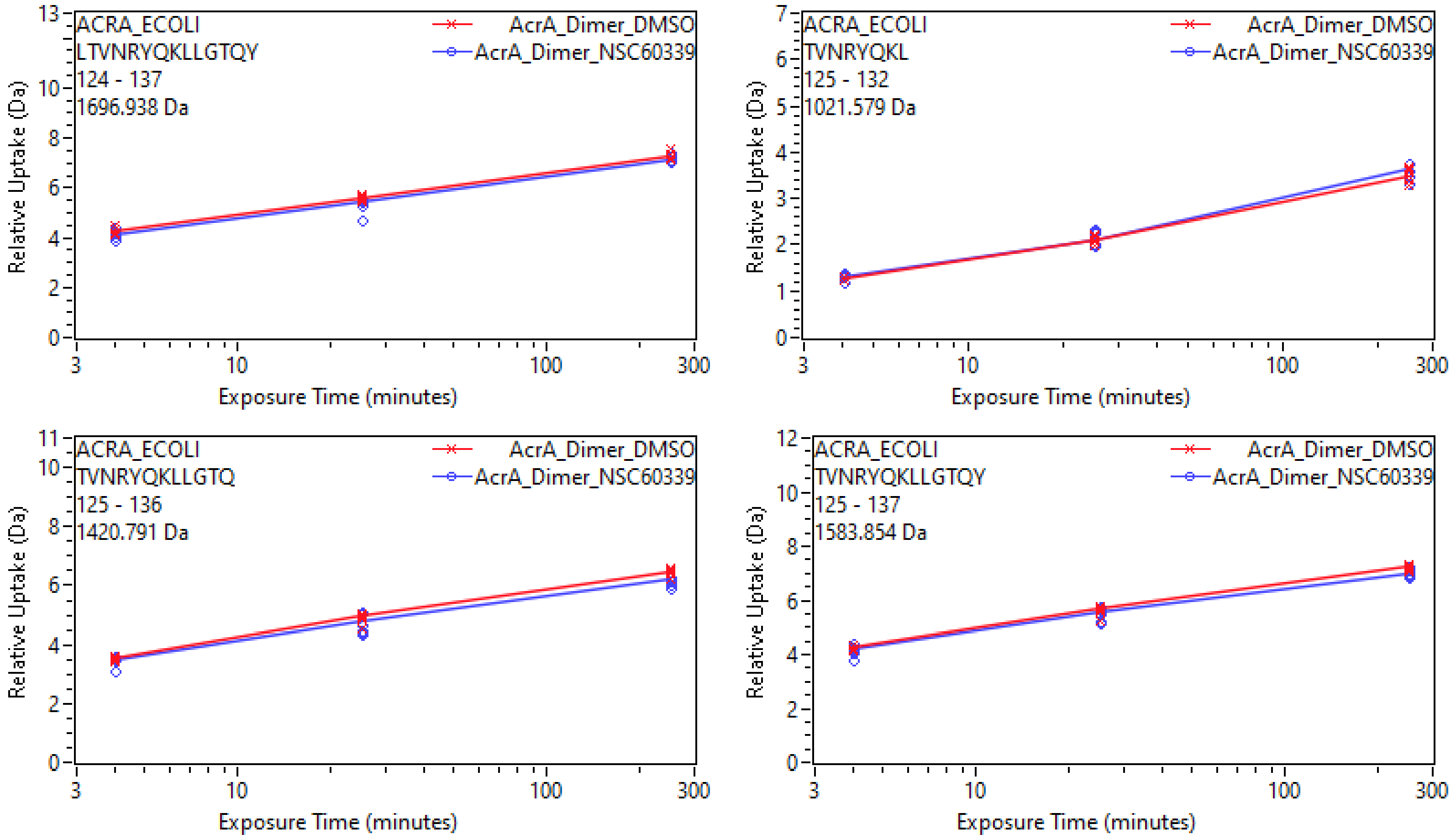
ACRA_ECOLI
LTVNRYQKLLGTQY
124 - 137
1696.938 Da
AcrA_Dimer_DMSO
AcrA_Dimer_NSC60339
Relative Uptake (Da)
Exposure Time (minutes)
ACRA_ECOLI
TVNRYQKL
125 - 132
1021.579 Da
AcrA_Dimer_DMSO
AcrA_Dimer_NSC60339
Relative Uptake (Da)
Exposure Time (minutes)
ACRA_ECOLI
TVNRYQKLLGTQ
125 - 136
1420.791 Da
AcrA_Dimer_DMSO
AcrA_Dimer_NSC60339
Relative Uptake (Da)
Exposure Time (minutes)
ACRA_ECOLI
TVNRYQKLLGTQY
125 - 137
1583.854 Da
AcrA_Dimer_DMSO
AcrA_Dimer_NSC60339
Relative Uptake (Da)
Exposure Time (minutes)

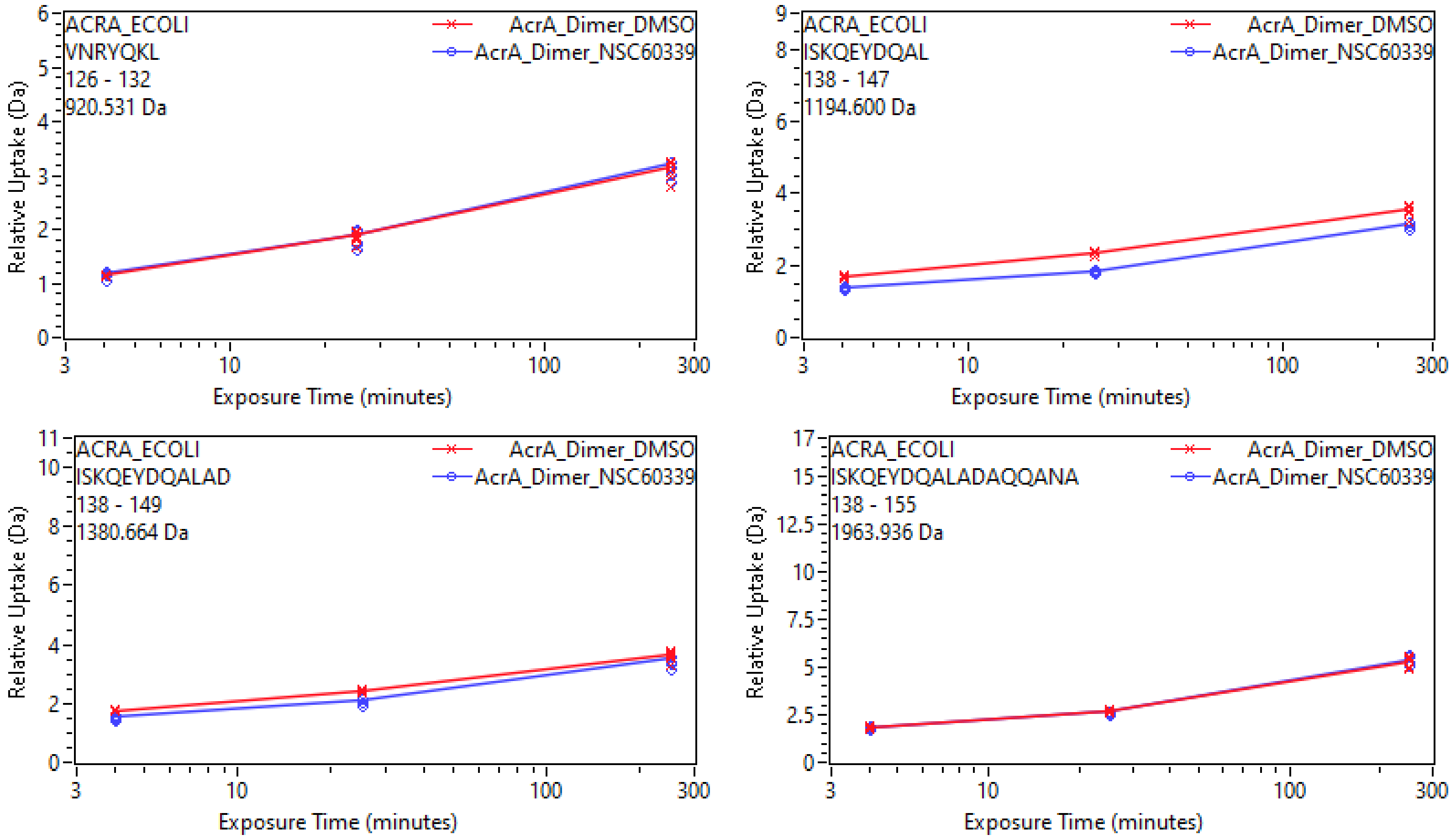
ACRA_ECOLI
VNRYQKL
126 - 132
920.531 Da
AcrA_Dimer_DMSO
AcrA_Dimer_NSC60339
Relative Uptake (Da)
Exposure Time (minutes)
ACRA_ECOLI
ISKQEYDQAL
138 - 147
1194.600 Da
AcrA_Dimer_DMSO
AcrA_Dimer_NSC60339
Relative Uptake (Da)
Exposure Time (minutes)
ACRA_ECOLI
ISKQEYDQALAD
138 - 149
1380.664 Da
AcrA_Dimer_DMSO
AcrA_Dimer_NSC60339
Relative Uptake (Da)
Exposure Time (minutes)
ACRA_ECOLI
ISKQEYDQALADAQQANA
138 - 155
1963.936 Da
AcrA_Dimer_DMSO
AcrA_Dimer_NSC60339
Relative Uptake (Da)
Exposure Time (minutes)

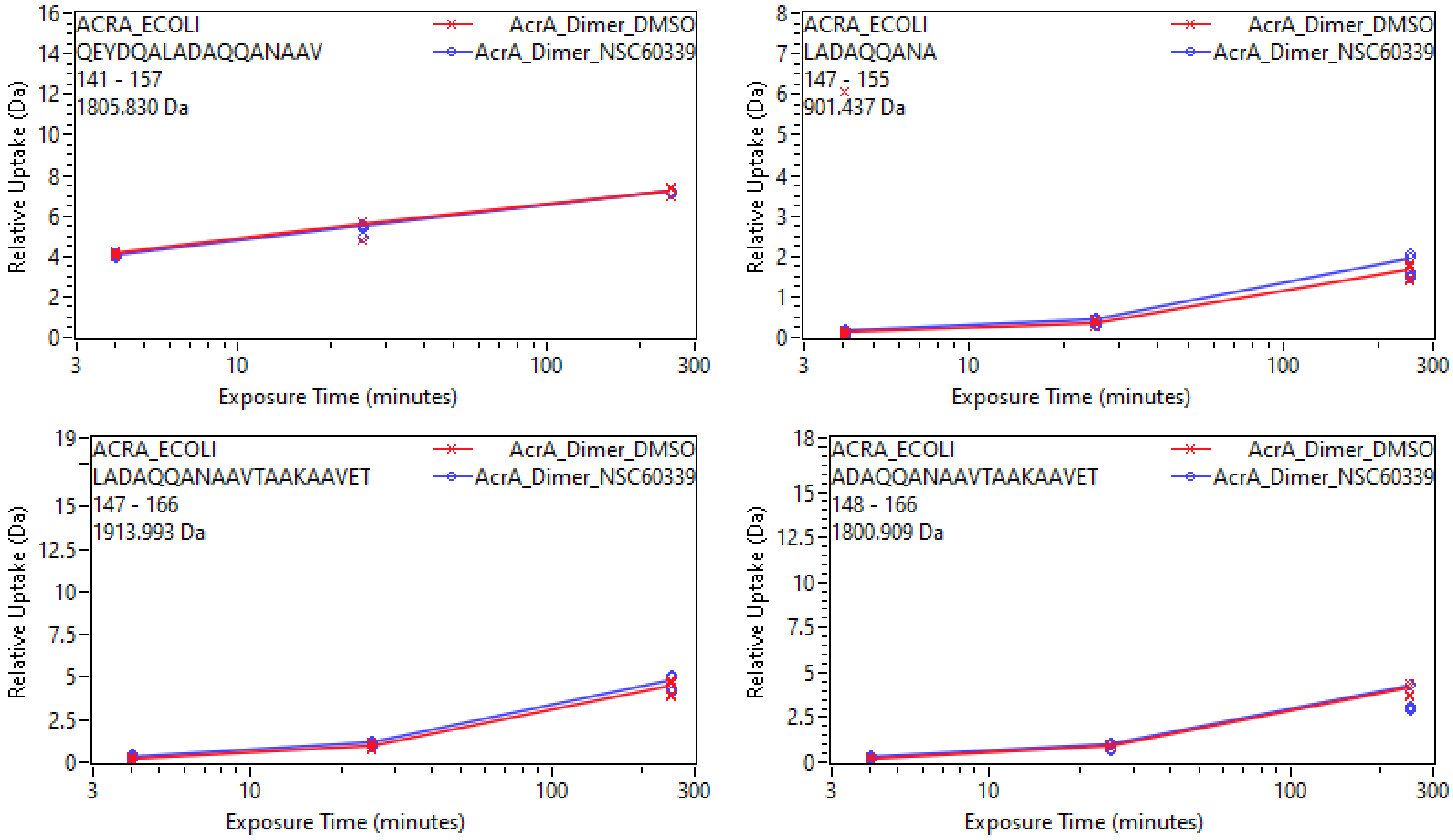

ACRA_ECOLI
QEYDQALADAQQANAAV
141 - 157
1805.830 Da
AcrA_Dimer_DMSO
AcrA_Dimer_NSC60339
Relative Uptake (Da)
Exposure Time (minutes)
ACRA_ECOLI
LADAQQANA
147 - 155
901.437 Da
AcrA_Dimer_DMSO
AcrA_Dimer_NSC60339
Relative Uptake (Da)
Exposure Time (minutes)
ACRA_ECOLI
LADAQQANAAVTAAKAAVET
147 - 166
1913.993 Da
AcrA_Dimer_DMSO
AcrA_Dimer_NSC60339
Relative Uptake (Da)
Exposure Time (minutes)
ACRA_ECOLI
ADAQQANAAVTAAKAAVET
148 - 166
1800.909 Da
AcrA_Dimer_DMSO
AcrA_Dimer_NSC60339
Relative Uptake (Da)
Exposure Time (minutes)

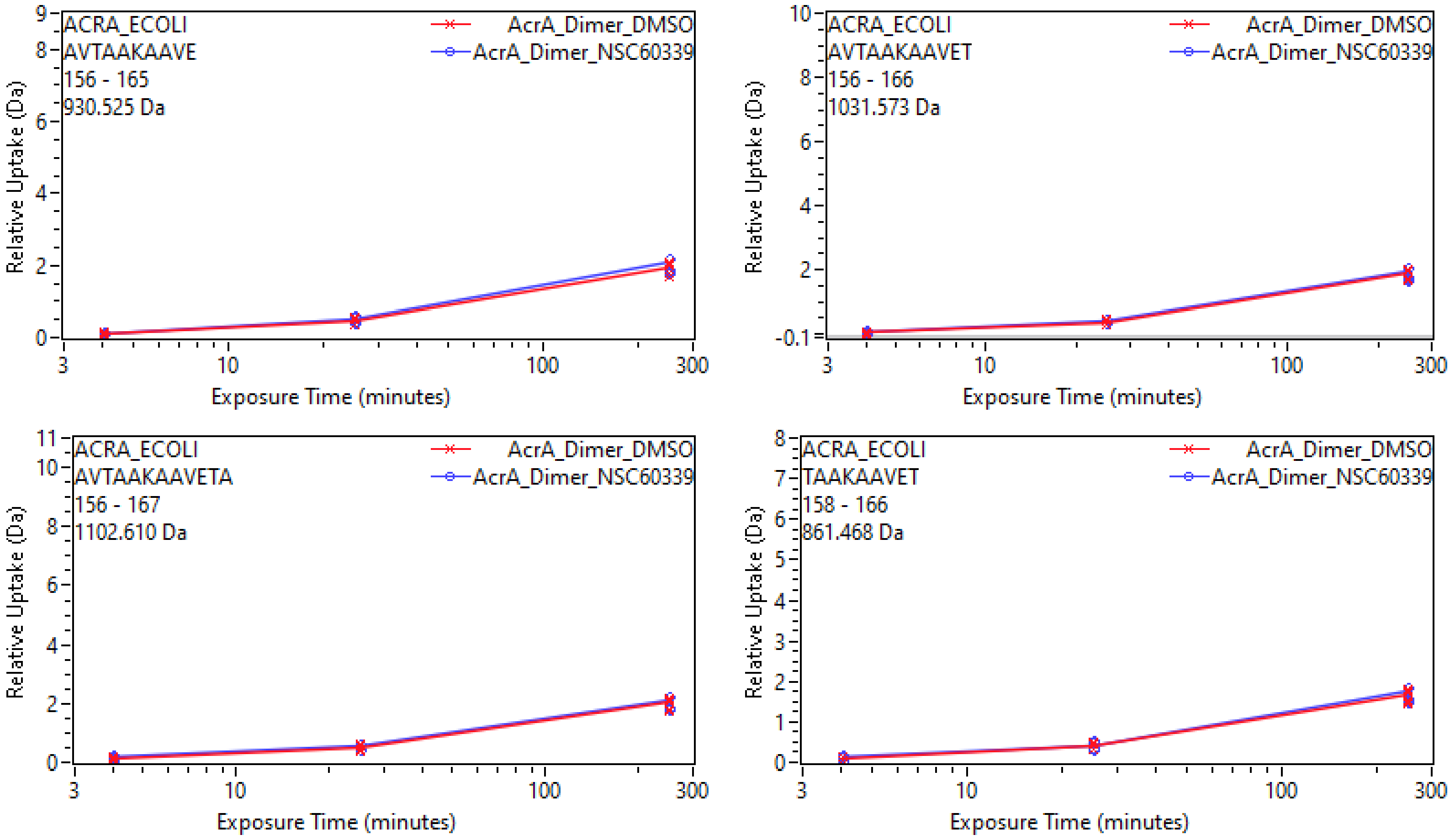
ACRA_ECOLI
AVTAAKAAVE
156 - 165
930.525 Da
AcrA_Dimer_DMSO
AcrA_Dimer_NSC60339
Relative Uptake (Da)
Exposure Time (minutes)
ACRA_ECOLI
AVTAAKAAVET
156 - 166
1031.573 Da
AcrA_Dimer_DMSO
AcrA_Dimer_NSC60339
Relative Uptake (Da)
Exposure Time (minutes)
ACRA_ECOLI
AVTAAKAAVETA
156 - 167
1102.610 Da
AcrA_Dimer_DMSO
AcrA_Dimer_NSC60339
Relative Uptake (Da)
Exposure Time (minutes)
ACRA_ECOLI
TAAKAAVET
158 - 166
861.468 Da
AcrA_Dimer_DMSO
AcrA_Dimer_NSC60339
Relative Uptake (Da)
Exposure Time (minutes)

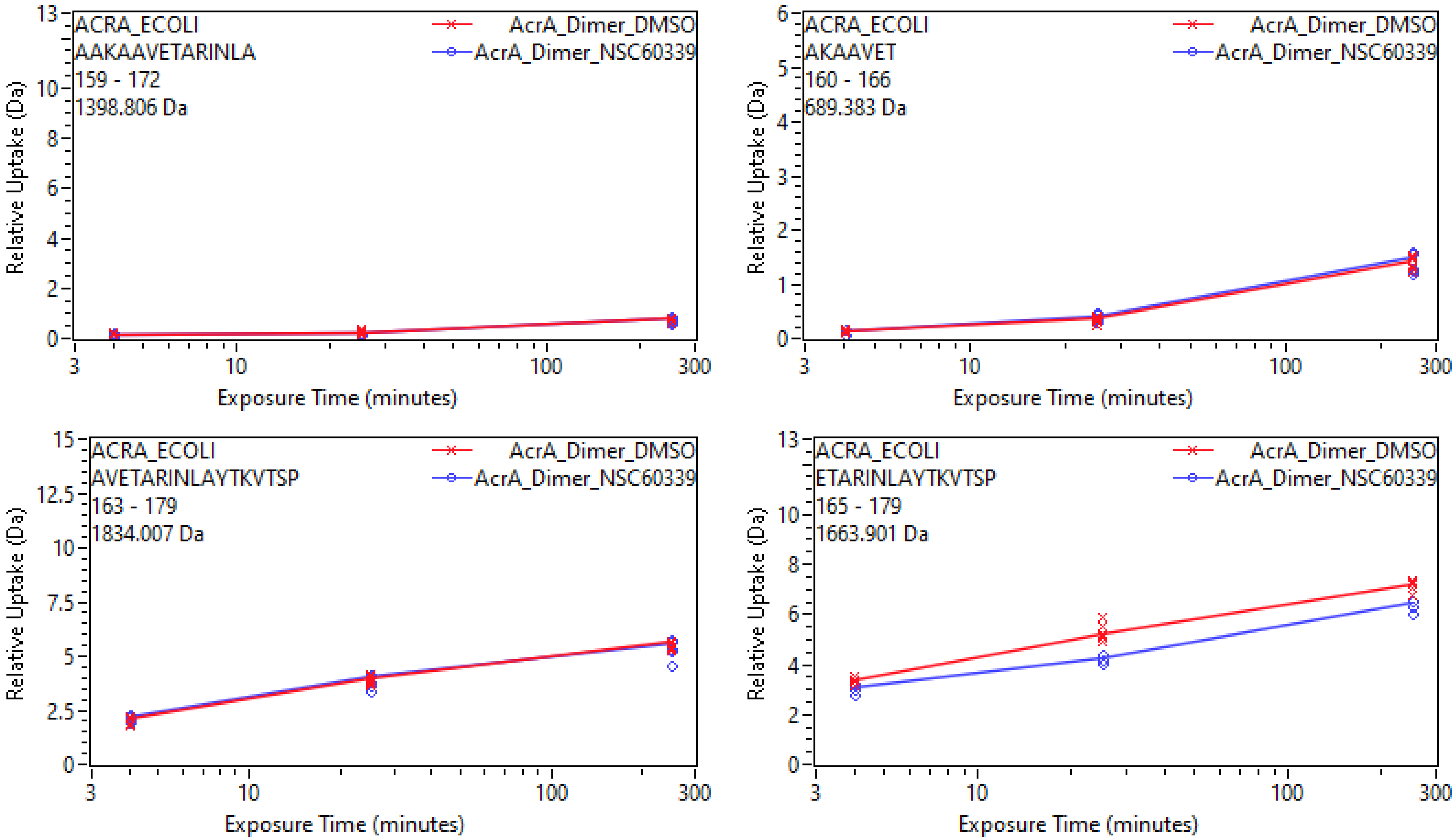
ACRA_ECOLI
AAKAAVETARINLA
159 - 172
1398.806 Da
AcrA_Dimer_DMSO
AcrA_Dimer_NSC60339
Relative Uptake (Da)
Exposure Time (minutes)
ACRA_ECOLI
AKAAVET
160 - 166
689.383 Da
AcrA_Dimer_DMSO
AcrA_Dimer_NSC60339
Relative Uptake (Da)
Exposure Time (minutes)
ACRA_ECOLI
AVETARINLAYTKVTSP
163 - 179
1834.007 Da
AcrA_Dimer_DMSO
AcrA_Dimer_NSC60339
Relative Uptake (Da)
Exposure Time (minutes)
ACRA_ECOLI
ETARINLAYTKVTSP
165 - 179
1663.901 Da
AcrA_Dimer_DMSO
AcrA_Dimer_NSC60339
Relative Uptake (Da)
Exposure Time (minutes)

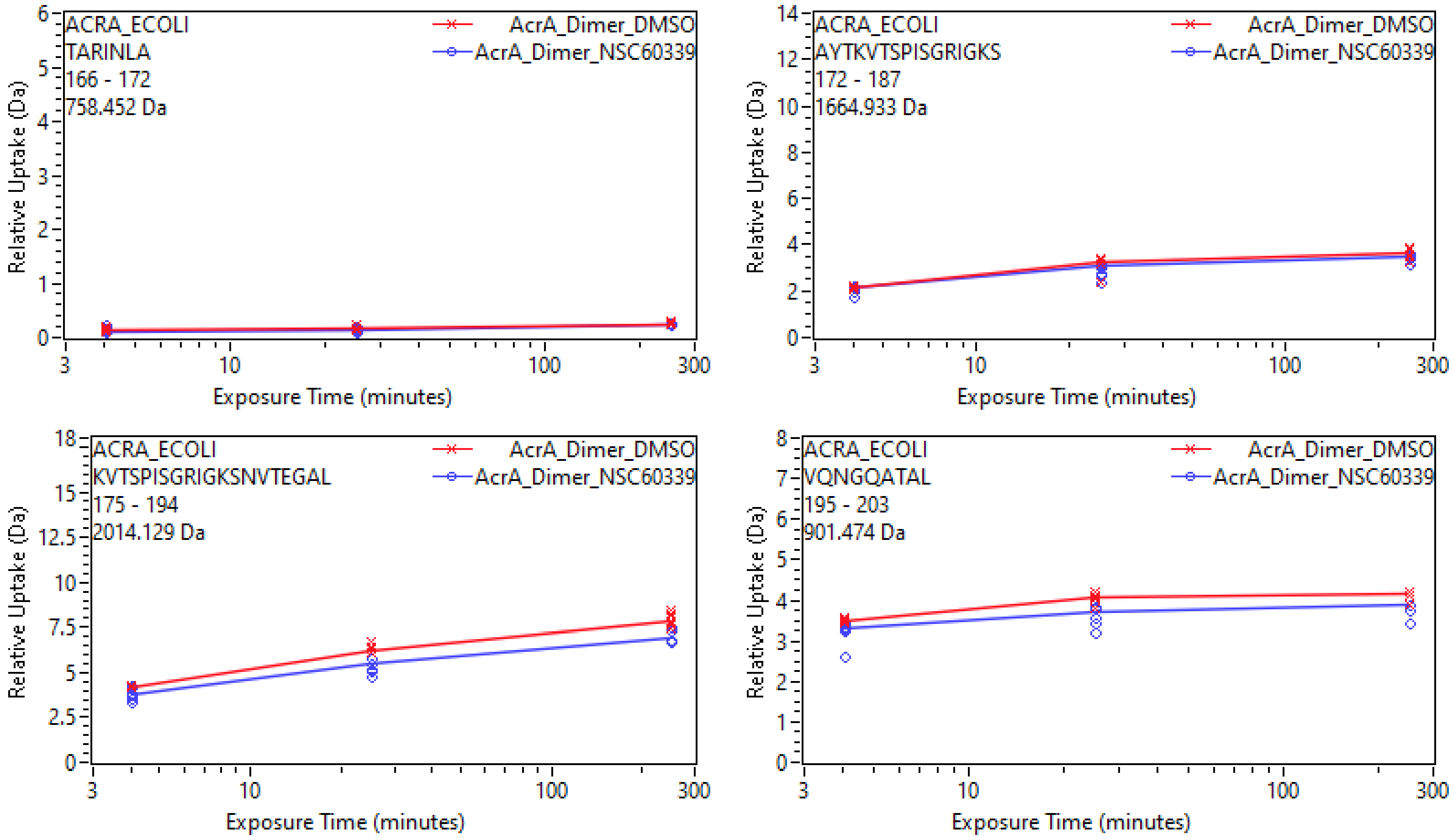

ACRA_ECOLI
TARINLA
166 - 172
758.452 Da
AcrA_Dimer_DMSO
AcrA_Dimer_NSC60339
Relative Uptake (Da)
Exposure Time (minutes)
ACRA_ECOLI
AYTKVTSPISGRIGKS
172 - 187
1664.933 Da
ACRA_ECOLI
KVTSPISGRIGKSNVTEGAL
175 - 194
2014.129 Da
ACRA_ECOLI
VQNGQATAL
195 - 203
901.474 Da

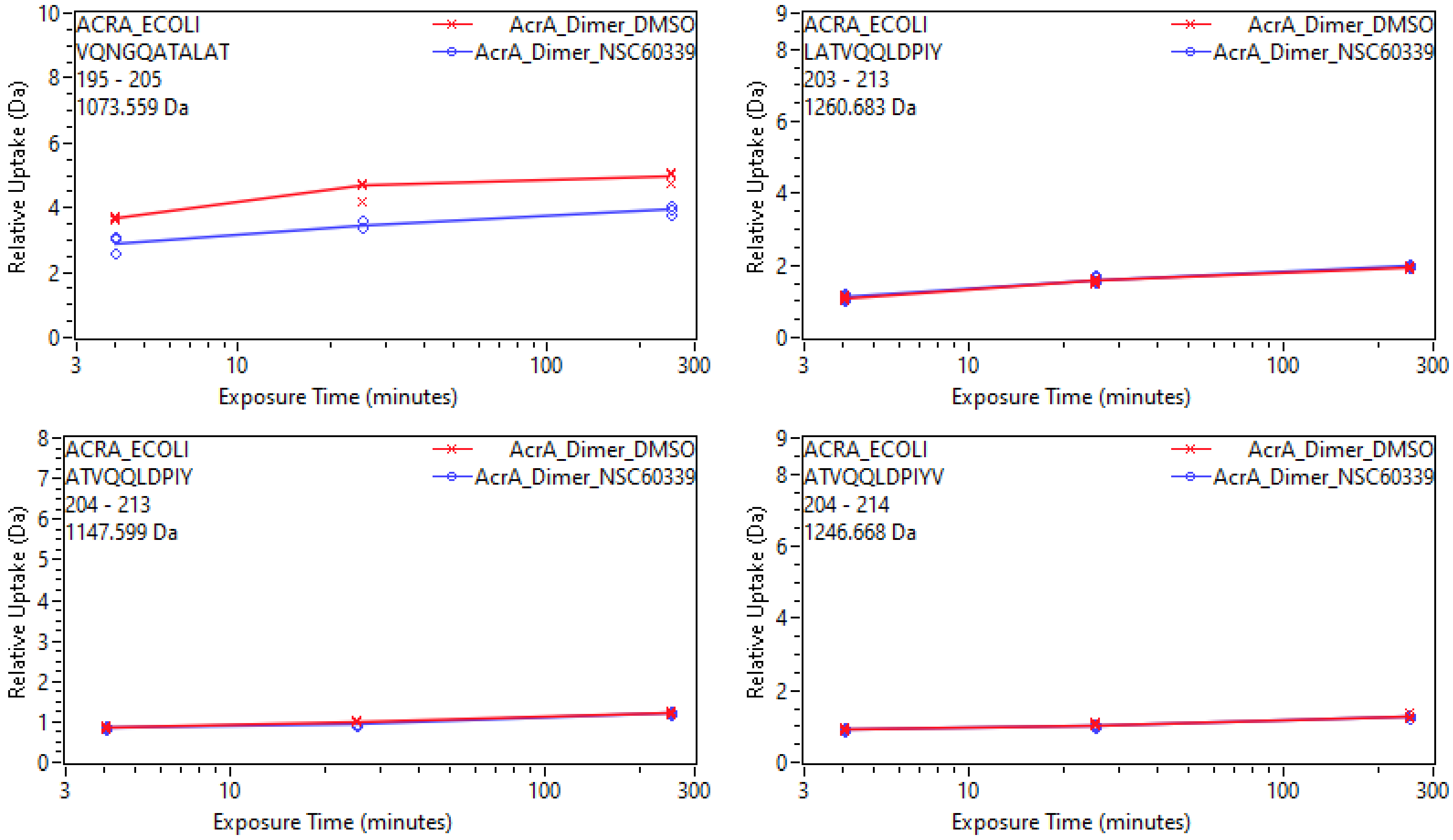

ACRA_ECOLI
VQNGQATALAT
195 - 205
1073.559 Da
AcrA_Dimer_DMSO
AcrA_Dimer_NSC60339
Relative Uptake (Da)
Exposure Time (minutes)
ACRA_ECOLI
LATVQQLDPIY
203 - 213
1260.683 Da
AcrA_Dimer_DMSO
AcrA_Dimer_NSC60339
Relative Uptake (Da)
Exposure Time (minutes)
ACRA_ECOLI
ATVQQLDPIY
204 - 213
1147.599 Da
AcrA_Dimer_DMSO
AcrA_Dimer_NSC60339
Relative Uptake (Da)
Exposure Time (minutes)
ACRA_ECOLI
ATVQQLDPIYV
204 - 214
1246.668 Da
AcrA_Dimer_DMSO
AcrA_Dimer_NSC60339
Relative Uptake (Da)
Exposure Time (minutes)

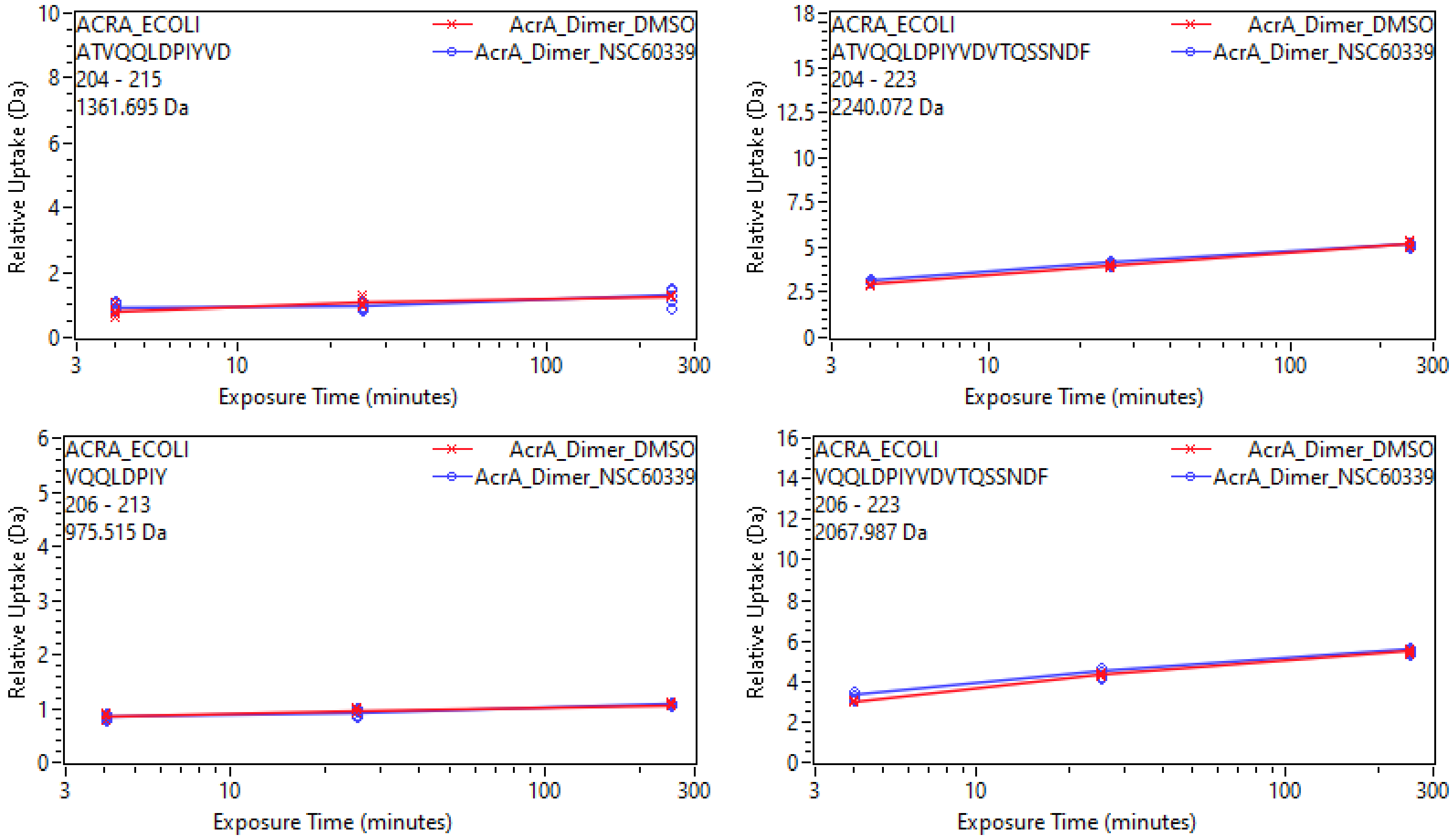

ACRA_ECOLI
ATVQQLDPIYVD
204 - 215
1361.695 Da
AcrA_Dimer_DMSO
AcrA_Dimer_NSC60339
Relative Uptake (Da)
Exposure Time (minutes)
ACRA_ECOLI
ATVQQLDPIYVDVTQSSNDF
204 - 223
2240.072 Da
AcrA_Dimer_DMSO
AcrA_Dimer_NSC60339
Relative Uptake (Da)
Exposure Time (minutes)
ACRA_ECOLI
VQQLDPIY
206 - 213
975.515 Da
AcrA_Dimer_DMSO
AcrA_Dimer_NSC60339
Relative Uptake (Da)
Exposure Time (minutes)
ACRA_ECOLI
VQQLDPIYVDVTQSSNDF
206 - 223
2067.987 Da
AcrA_Dimer_DMSO
AcrA_Dimer_NSC60339
Relative Uptake (Da)
Exposure Time (minutes)

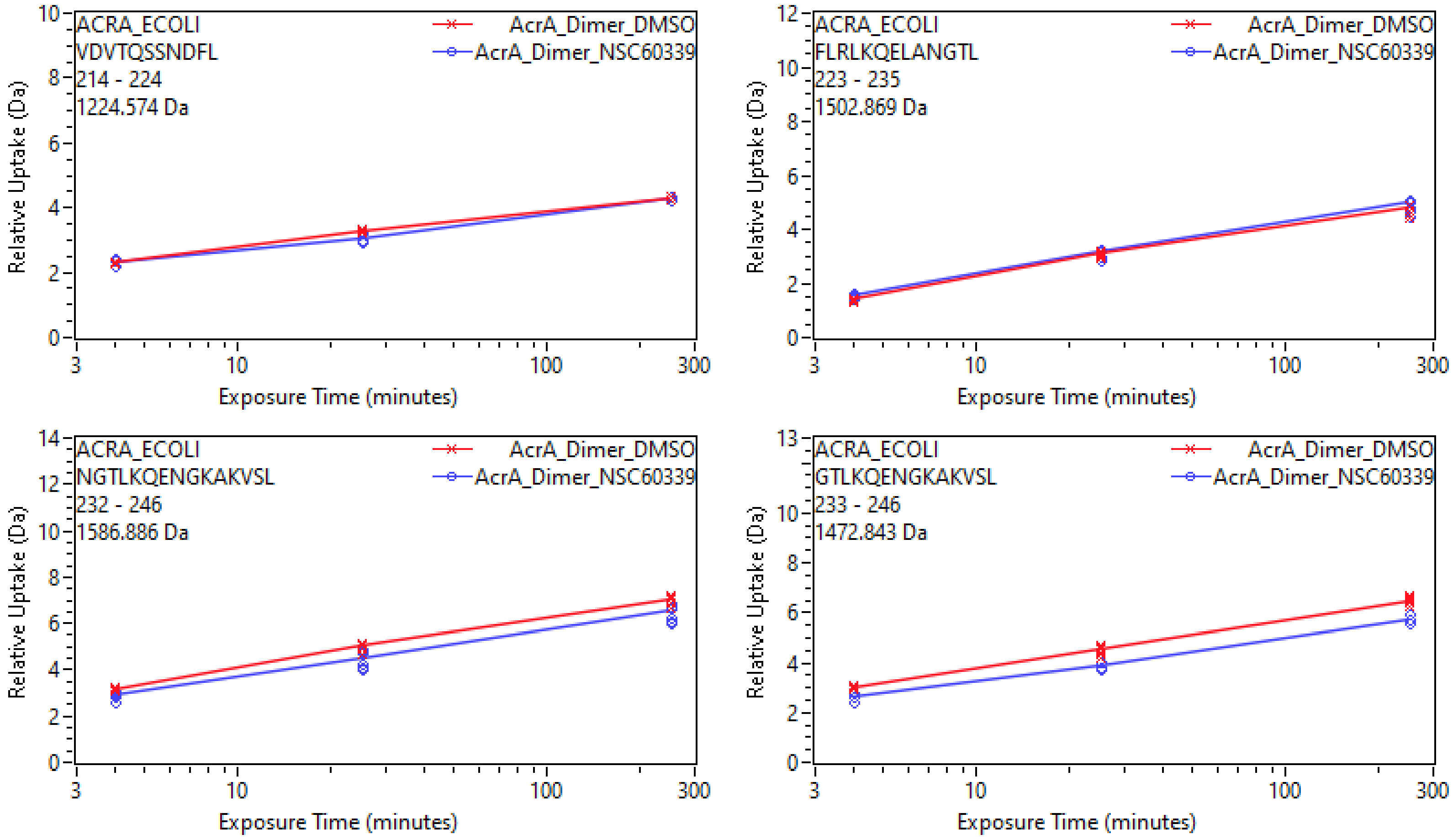
ACRA_ECOLI
VDVTQSSNDFL
214 - 224
1224.574 Da
AcrA_Dimer_DMSO
AcrA_Dimer_NSC60339
Relative Uptake (Da)
Exposure Time (minutes)
ACRA_ECOLI
FLRLKQELANGTL
223 - 235
1502.869 Da
AcrA_Dimer_DMSO
AcrA_Dimer_NSC60339
Relative Uptake (Da)
Exposure Time (minutes)
ACRA_ECOLI
NGTLKQENGKAKVSL
232 - 246
1586.886 Da
AcrA_Dimer_DMSO
AcrA_Dimer_NSC60339
Relative Uptake (Da)
Exposure Time (minutes)
ACRA_ECOLI
GTLKQENGKAKVSL
233 - 246
1472.843 Da
AcrA_Dimer_DMSO
AcrA_Dimer_NSC60339
Relative Uptake (Da)
Exposure Time (minutes)

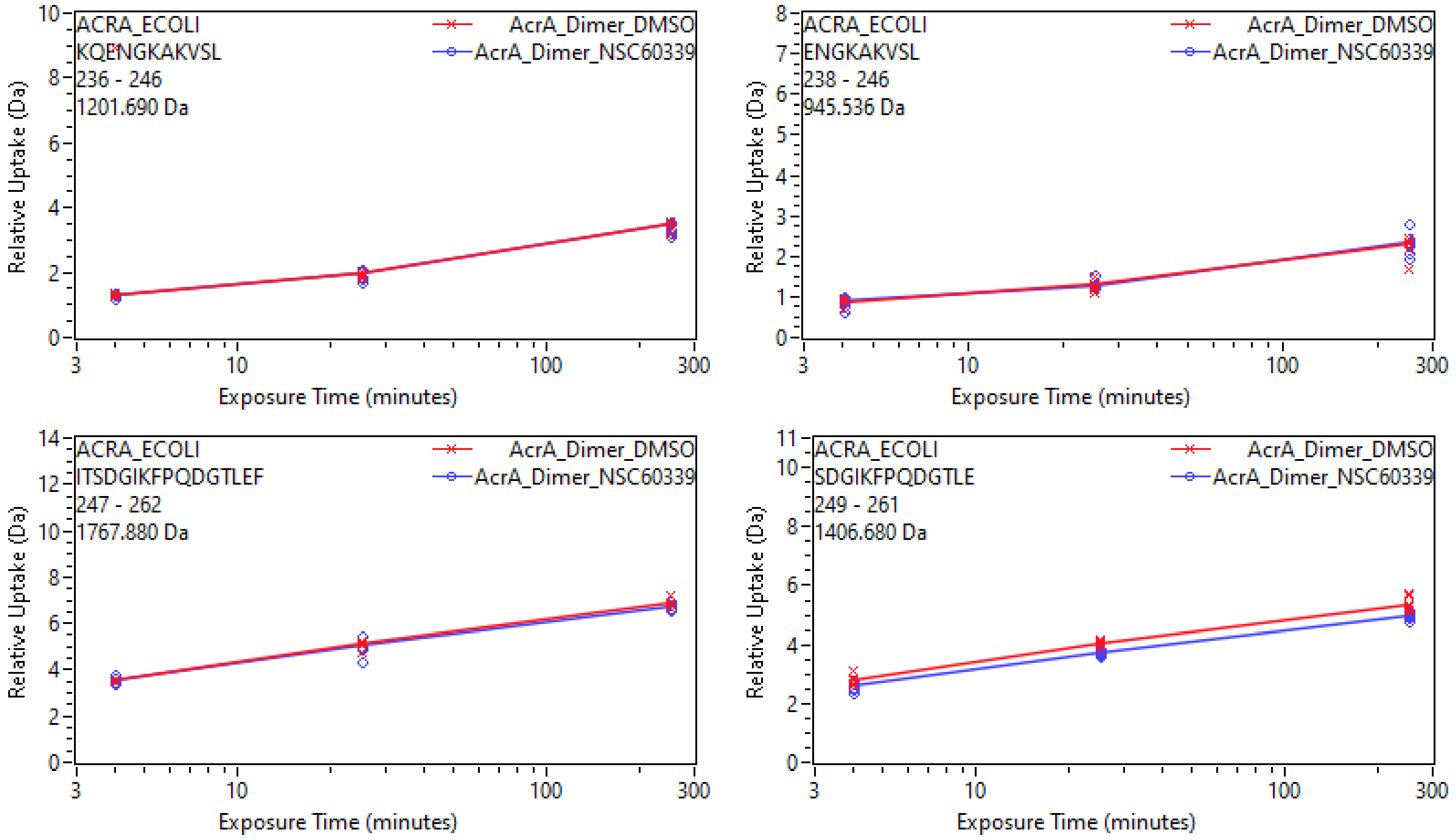
ACRA_ECOLI
KQENGKAKVSL
236 - 246
1201.690 Da
AcrA_Dimer_DMSO
AcrA_Dimer_NSC60339
Relative Uptake (Da)
Exposure Time (minutes)
ACRA_ECOLI
ENGKAKVSL
238 - 246
945.536 Da
ACRA_ECOLI
ITSDGIKFPQDGTLEF
247 - 262
1767.880 Da
ACRA_ECOLI
SDGIKFPQDGTLE
249 - 261
1406.680 Da

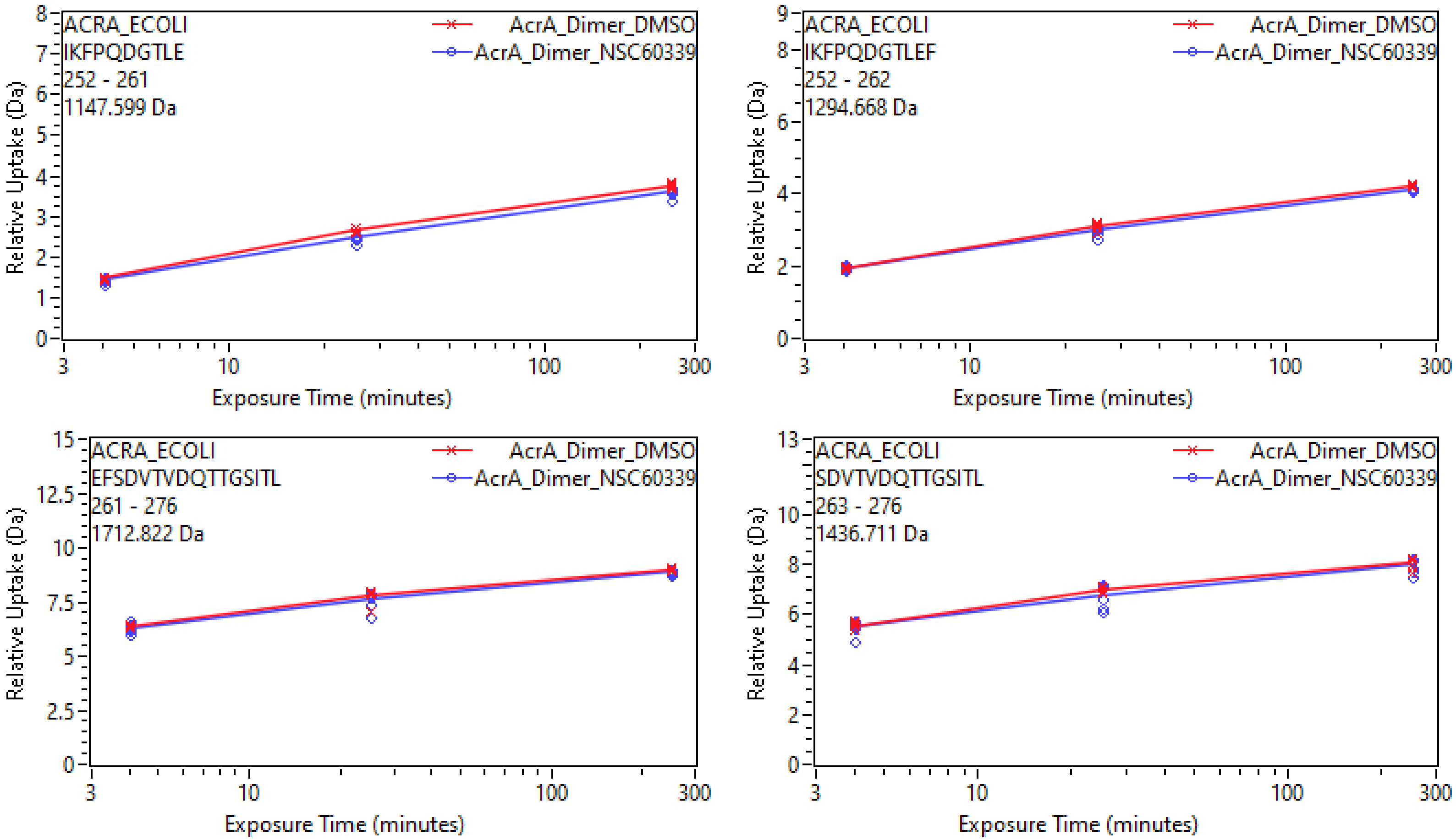
ACRA_ECOLI
IKFPQDGTLE
252 - 261
1147.599 Da
AcrA_Dimer_DMSO
AcrA_Dimer_NSC60339
Relative Uptake (Da)
Exposure Time (minutes)
ACRA_ECOLI
IKFPQDGTLEF
252 - 262
1294.668 Da
ACRA_ECOLI
EFSDVTVDQTTGSITL
261 - 276
1712.822 Da
ACRA_ECOLI
SDVTVDQTTGSITL
263 - 276
1436.711 Da

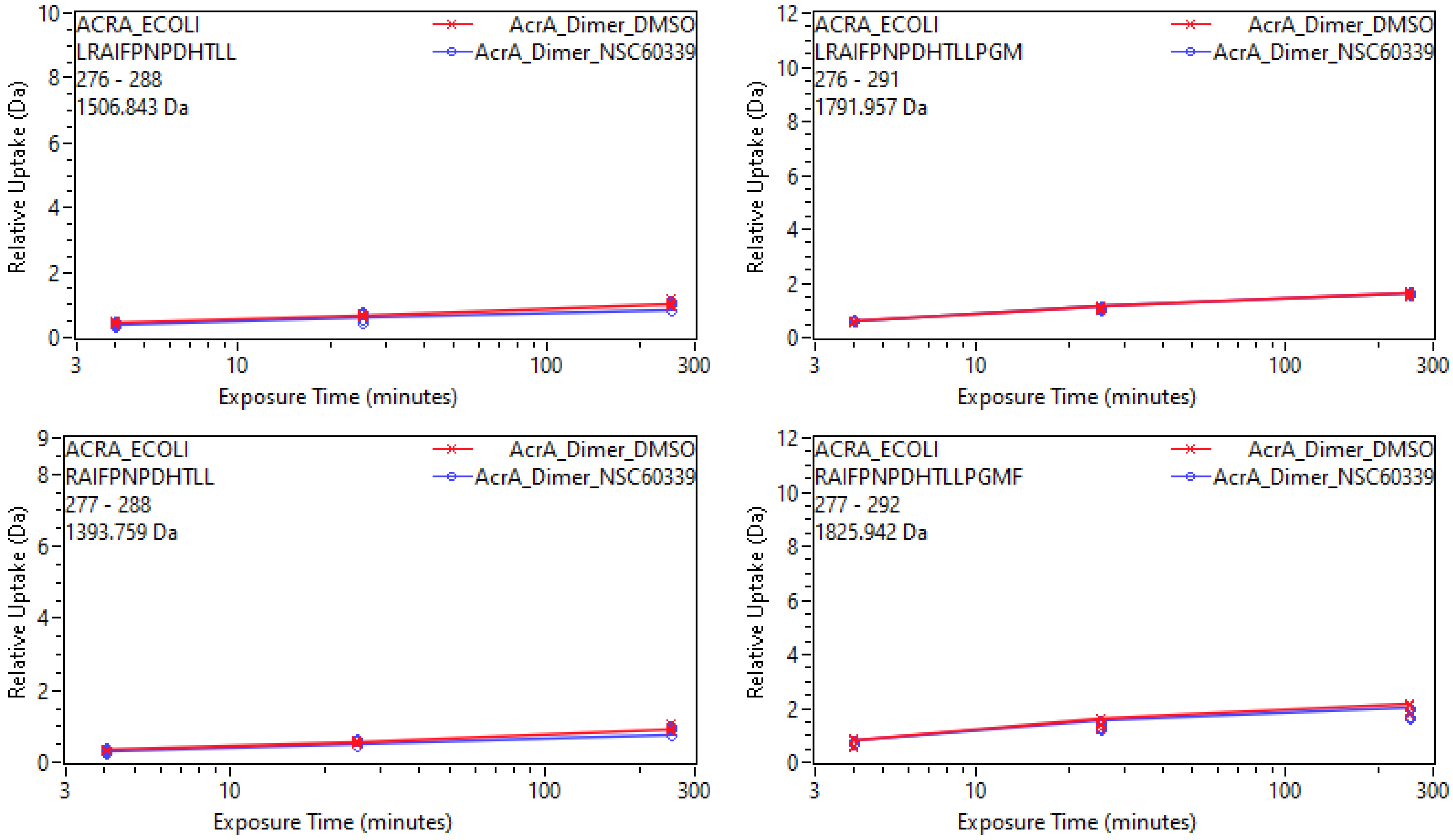

ACRA_ECOLI
LRAIFPNPDHTLL
276 - 288
1506.843 Da
AcrA_Dimer_DMSO
AcrA_Dimer_NSC60339
Relative Uptake (Da)
Exposure Time (minutes)
ACRA_ECOLI
LRAIFPNPDHTLLPGM
276 - 291
1791.957 Da
AcrA_Dimer_DMSO
AcrA_Dimer_NSC60339
Relative Uptake (Da)
Exposure Time (minutes)
ACRA_ECOLI
RAIFPNPDHTLL
277 - 288
1393.759 Da
AcrA_Dimer_DMSO
AcrA_Dimer_NSC60339
Relative Uptake (Da)
Exposure Time (minutes)
ACRA_ECOLI
RAIFPNPDHTLLPGMF
277 - 292
1825.942 Da
AcrA_Dimer_DMSO
AcrA_Dimer_NSC60339
Relative Uptake (Da)
Exposure Time (minutes)

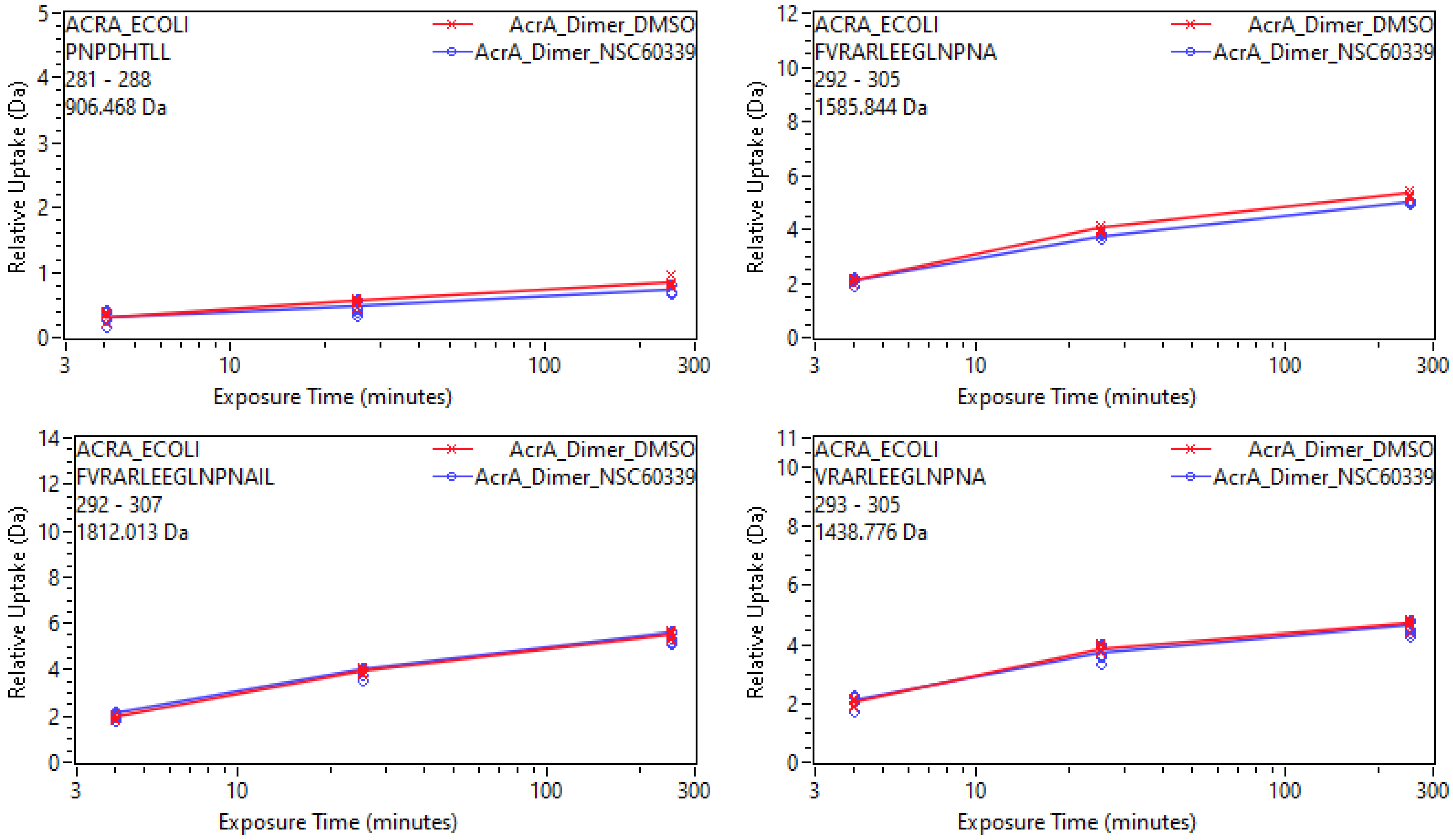
ACRA_ECOLI
PNPDHTLL
281 - 288
906.468 Da
AcrA_Dimer_DMSO
AcrA_Dimer_NSC60339
Relative Uptake (Da)
Exposure Time (minutes)
ACRA_ECOLI
FVRARLEEGLNPNA
292 - 305
1585.844 Da
ACRA_ECOLI
FVRARLEEGLNPNAIL
292 - 307
1812.013 Da
ACRA_ECOLI
VRARLEEGLNPNA
293 - 305
1438.776 Da

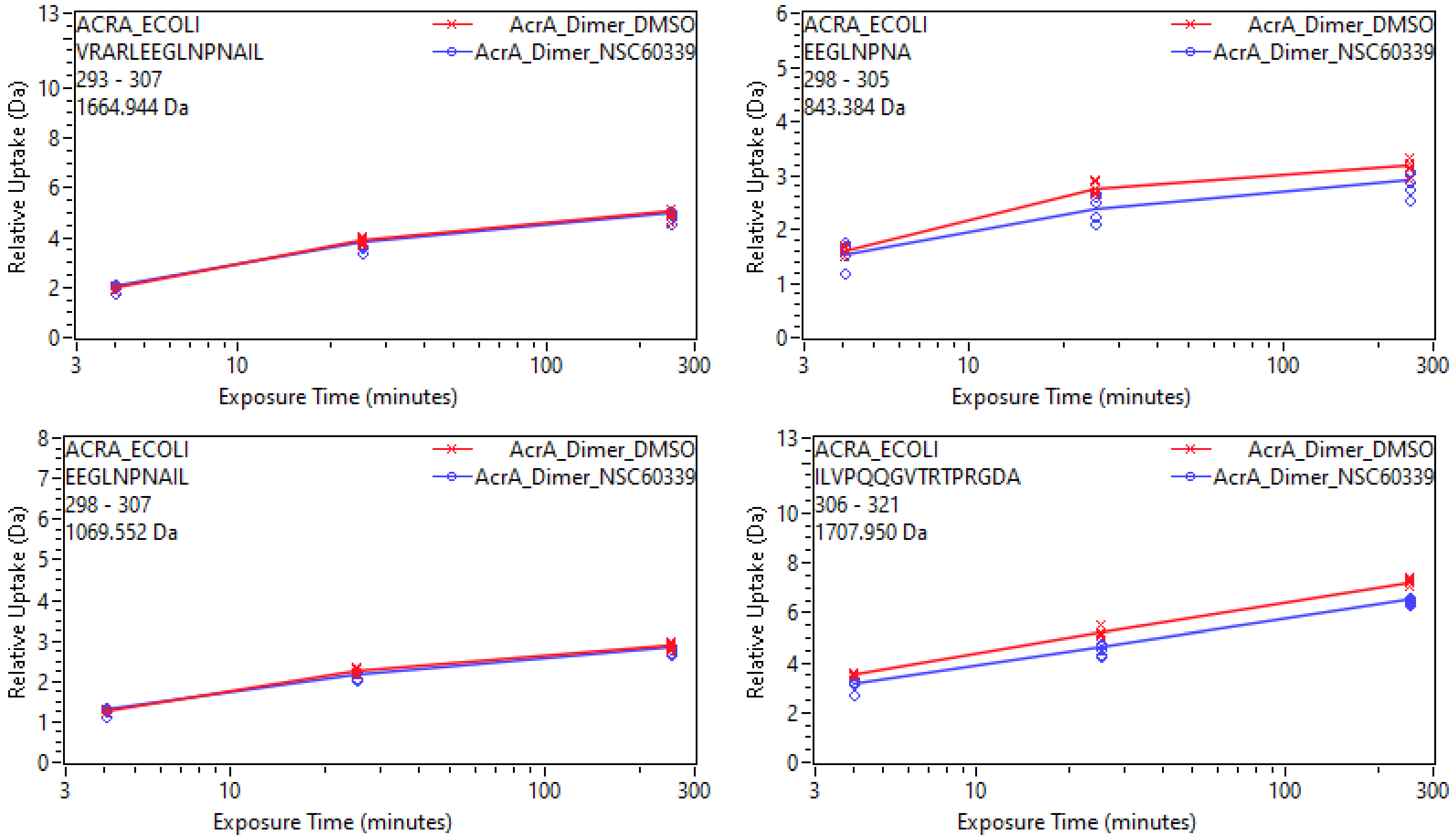
ACRA_ECOLI
VRARLEEGLNPNAIL
293 - 307
1664.944 Da
AcrA_Dimer_DMSO
AcrA_Dimer_NSC60339
Relative Uptake (Da)
Exposure Time (minutes)
ACRA_ECOLI
EEGLNPNA
298 - 305
843.384 Da
ACRA_ECOLI
EEGLNPNAIL
298 - 307
1069.552 Da
ACRA_ECOLI
ILVPQQGVTRTPRGDA
306 - 321
1707.950 Da

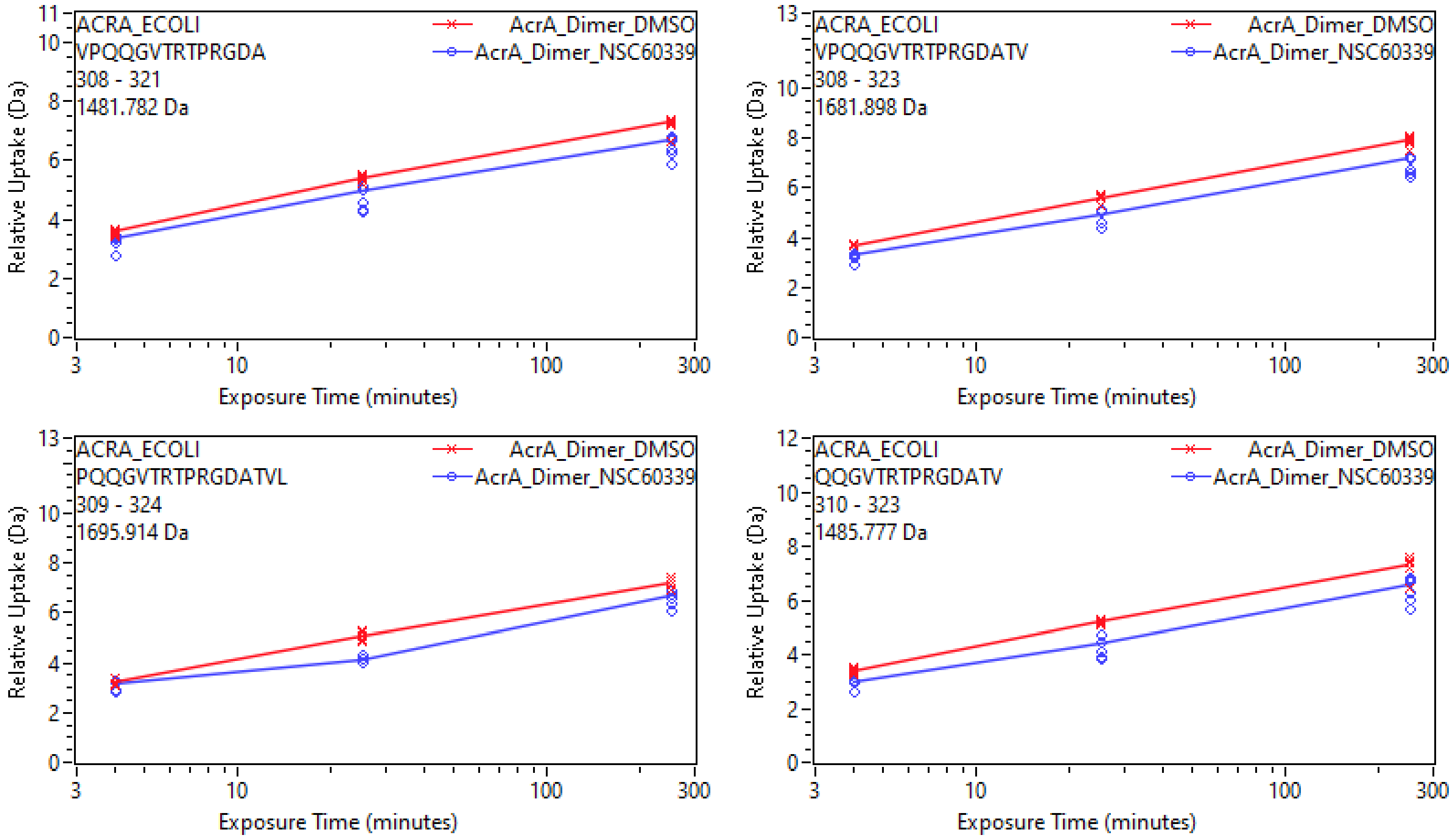
ACRA_ECOLI
VPQQGVTRTPRGDA
308 - 321
1481.782 Da
AcrA_Dimer_DMSO
AcrA_Dimer_NSC60339
Relative Uptake (Da)
Exposure Time (minutes)
ACRA_ECOLI
VPQQGVTRTPRGDATV
308 - 323
1681.898 Da
AcrA_Dimer_DMSO
AcrA_Dimer_NSC60339
Relative Uptake (Da)
Exposure Time (minutes)
ACRA_ECOLI
PQQGVTRTPRGDATVL
309 - 324
1695.914 Da
AcrA_Dimer_DMSO
AcrA_Dimer_NSC60339
Relative Uptake (Da)
Exposure Time (minutes)
ACRA_ECOLI
QQGVTRTPRGDATV
310 - 323
1485.777 Da
AcrA_Dimer_DMSO
AcrA_Dimer_NSC60339
Relative Uptake (Da)
Exposure Time (minutes)

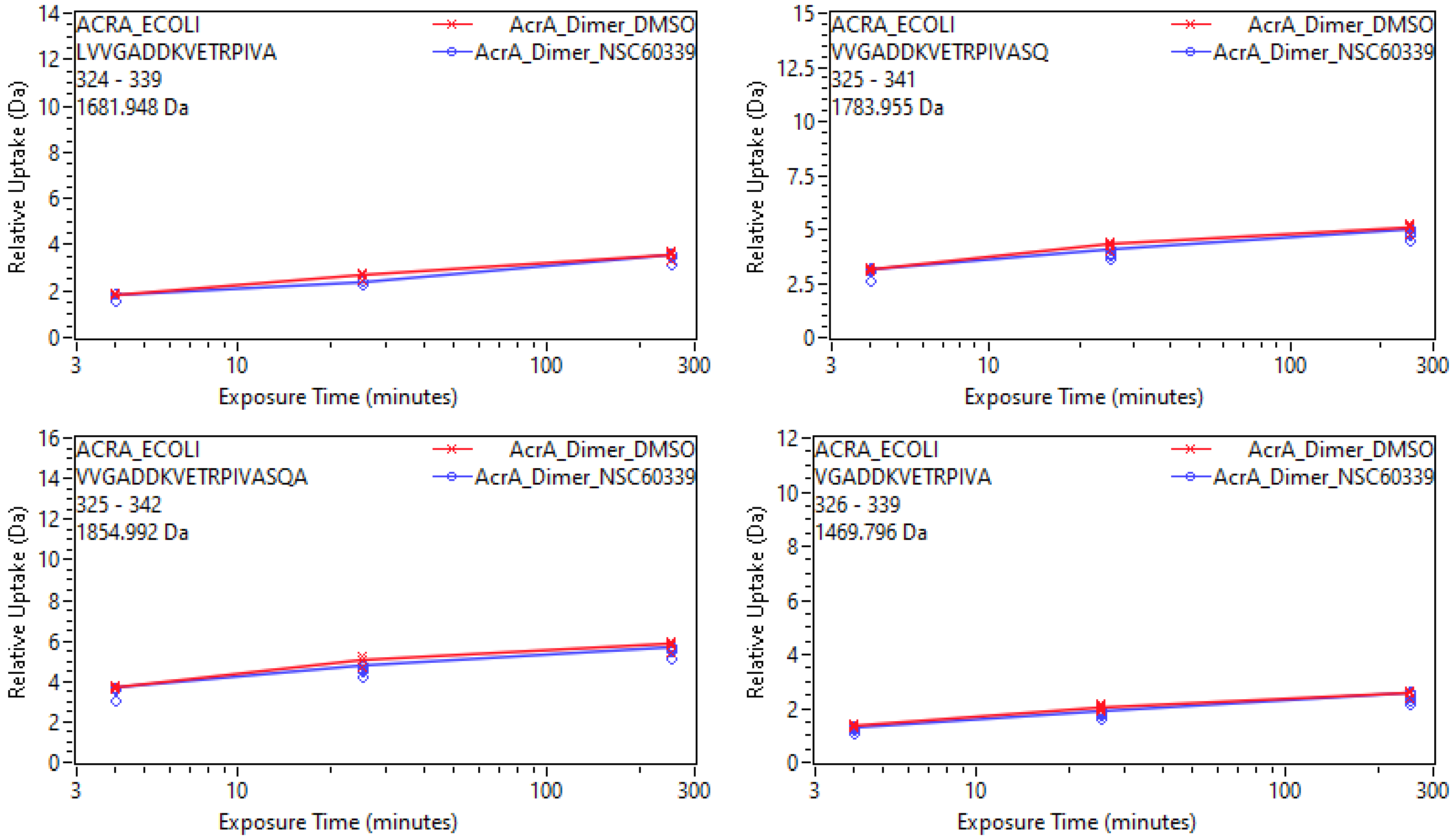
ACRA_ECOLI
LVVGADDKVETRPIVA
324 - 339
1681.948 Da
AcrA_Dimer_DMSO
AcrA_Dimer_NSC60339
Relative Uptake (Da)
Exposure Time (minutes)
ACRA_ECOLI
VVGADDKVETRPIVASQ
325 - 341
1783.955 Da
AcrA_Dimer_DMSO
AcrA_Dimer_NSC60339
Relative Uptake (Da)
Exposure Time (minutes)
ACRA_ECOLI
VVGADDKVETRPIVASQA
325 - 342
1854.992 Da
AcrA_Dimer_DMSO
AcrA_Dimer_NSC60339
Relative Uptake (Da)
Exposure Time (minutes)
ACRA_ECOLI
VGADDKVETRPIVA
326 - 339
1469.796 Da
AcrA_Dimer_DMSO
AcrA_Dimer_NSC60339
Relative Uptake (Da)
Exposure Time (minutes)

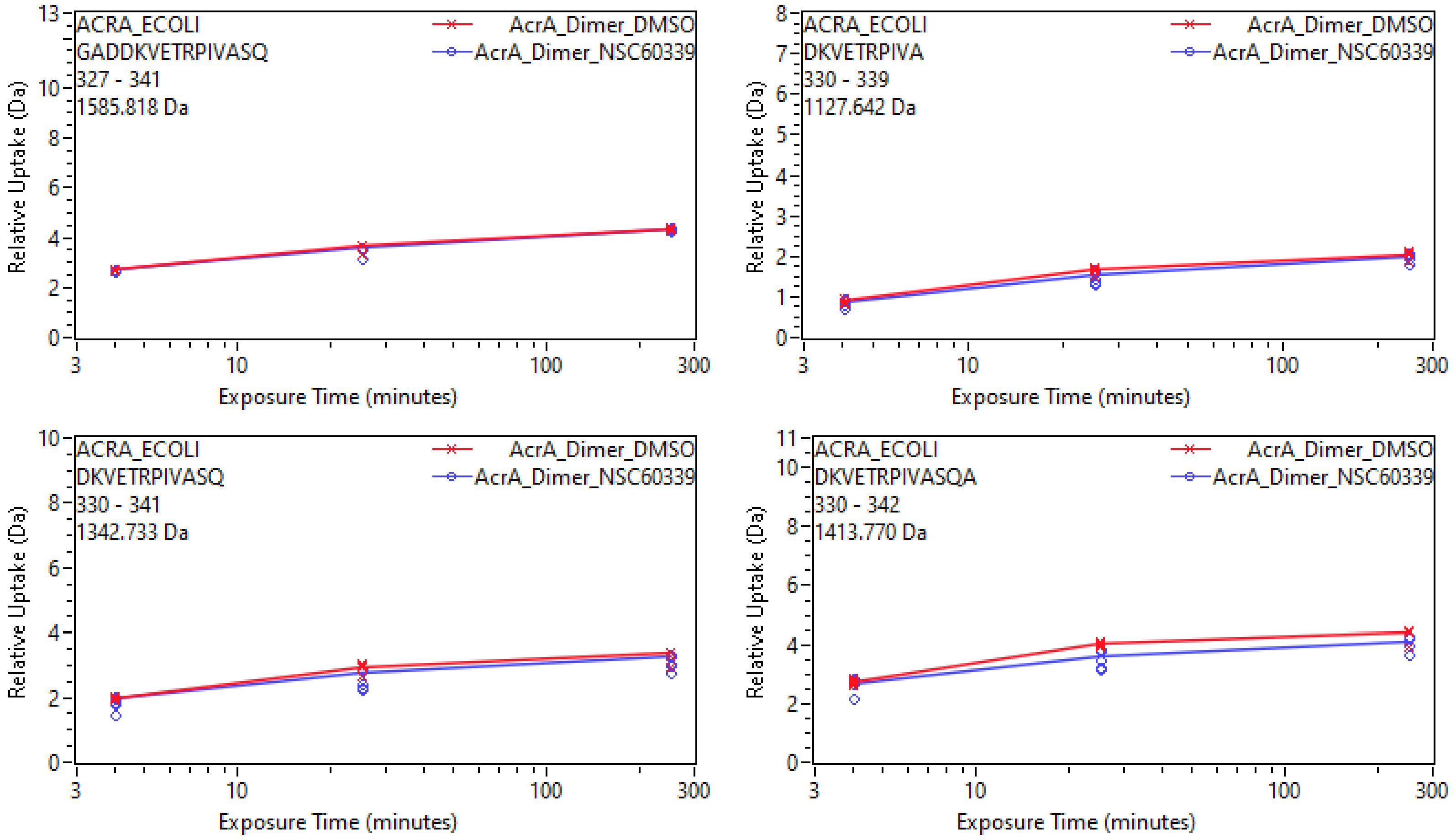
ACRA_ECOLI
GADDKVETRPIVASQ
327 - 341
1585.818 Da
AcrA_Dimer_DMSO
AcrA_Dimer_NSC60339
Relative Uptake (Da)
Exposure Time (minutes)
ACRA_ECOLI
DKVETRPIVA
330 - 339
1127.642 Da
AcrA_Dimer_DMSO
AcrA_Dimer_NSC60339
Relative Uptake (Da)
Exposure Time (minutes)
ACRA_ECOLI
DKVETRPIVASQ
330 - 341
1342.733 Da
AcrA_Dimer_DMSO
AcrA_Dimer_NSC60339
Relative Uptake (Da)
Exposure Time (minutes)
ACRA_ECOLI
DKVETRPIVASQA
330 - 342
1413.770 Da
AcrA_Dimer_DMSO
AcrA_Dimer_NSC60339
Relative Uptake (Da)
Exposure Time (minutes)

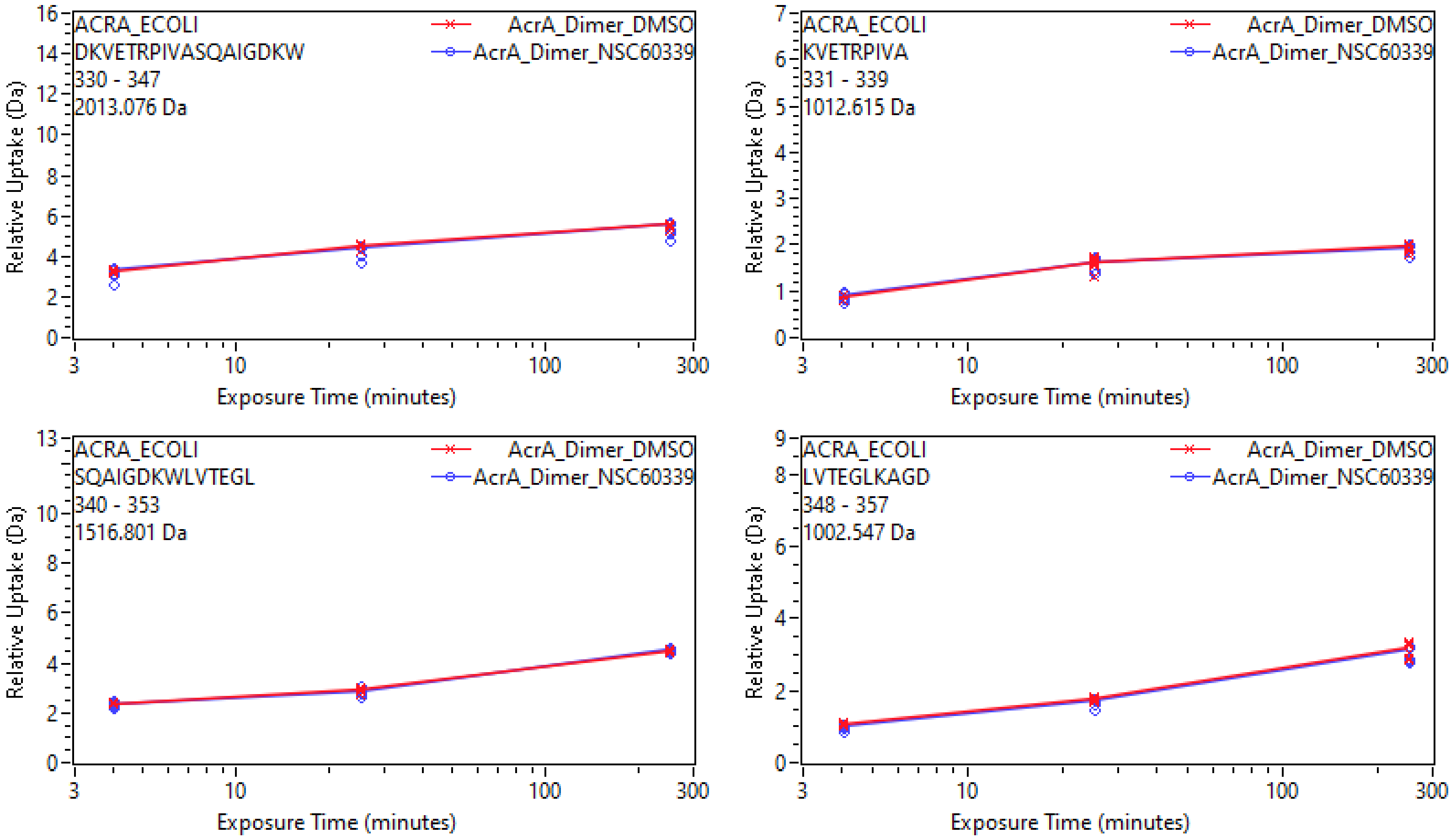
ACRA_ECOLI
DKVETRPIVASQAIGDKW
330 - 347
2013.076 Da
AcrA_Dimer_DMSO
AcrA_Dimer_NSC60339
Relative Uptake (Da)
Exposure Time (minutes)
ACRA_ECOLI
KVETRPIVA
331 - 339
1012.615 Da
AcrA_Dimer_DMSO
AcrA_Dimer_NSC60339
Relative Uptake (Da)
Exposure Time (minutes)
ACRA_ECOLI
SQAIGDKWLVTEGL
340 - 353
1516.801 Da
AcrA_Dimer_DMSO
AcrA_Dimer_NSC60339
Relative Uptake (Da)
Exposure Time (minutes)
ACRA_ECOLI
LVTEGLKAGD
348 - 357
1002.547 Da
AcrA_Dimer_DMSO
AcrA_Dimer_NSC60339
Relative Uptake (Da)
Exposure Time (minutes)

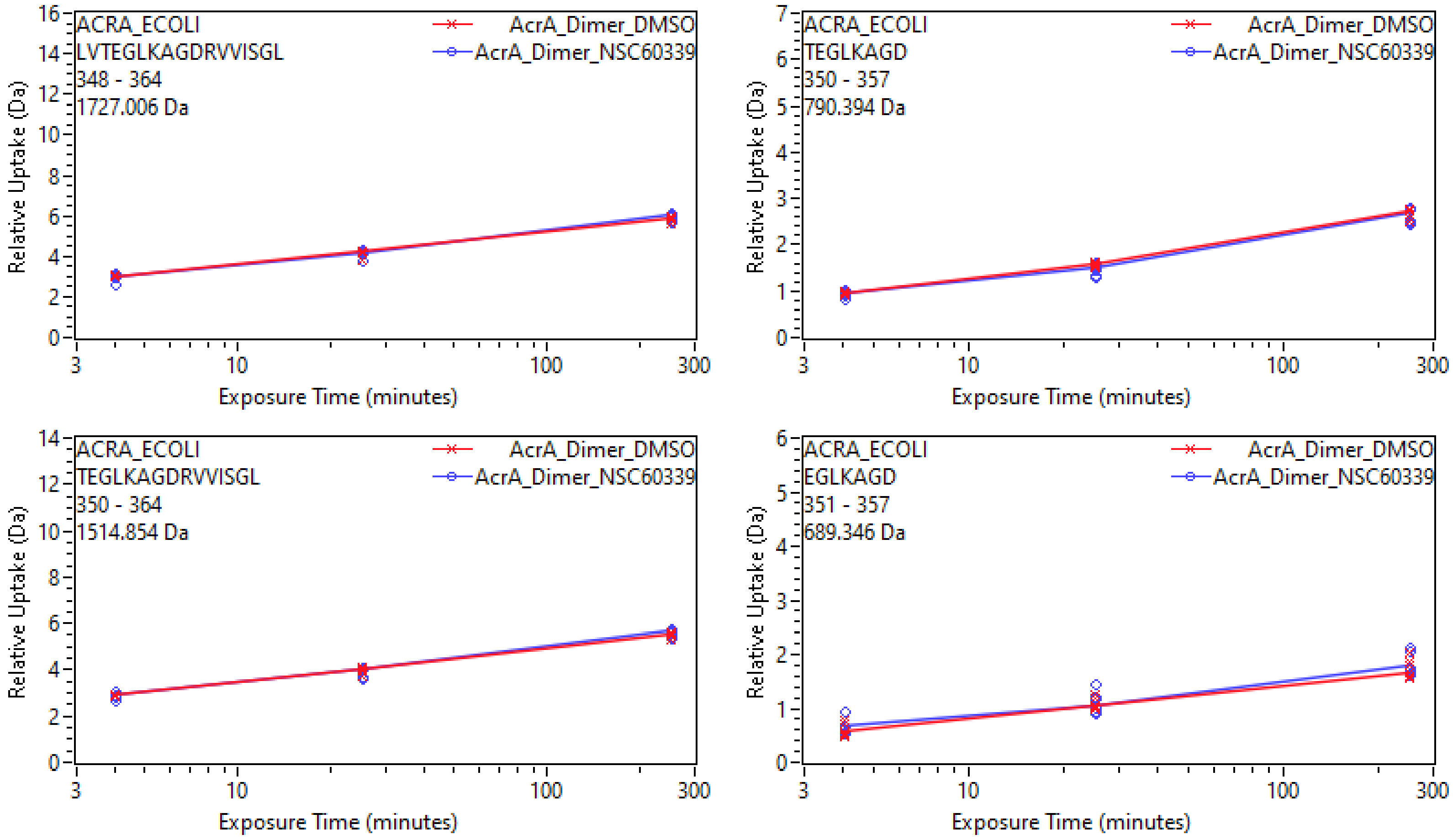
ACRA_ECOLI
LVTEGLKAGDRVVISGL
348 - 364
1727.006 Da
AcrA_Dimer_DMSO
AcrA_Dimer_NSC60339
Relative Uptake (Da)
Exposure Time (minutes)
ACRA_ECOLI
TEGLKAGD
350 - 357
790.394 Da
AcrA_Dimer_DMSO
AcrA_Dimer_NSC60339
Relative Uptake (Da)
Exposure Time (minutes)
ACRA_ECOLI
TEGLKAGDRVVISGL
350 - 364
1514.854 Da
AcrA_Dimer_DMSO
AcrA_Dimer_NSC60339
Relative Uptake (Da)
Exposure Time (minutes)
ACRA_ECOLI
EGLKAGD
351 - 357
689.346 Da
AcrA_Dimer_DMSO
AcrA_Dimer_NSC60339
Relative Uptake (Da)
Exposure Time (minutes)

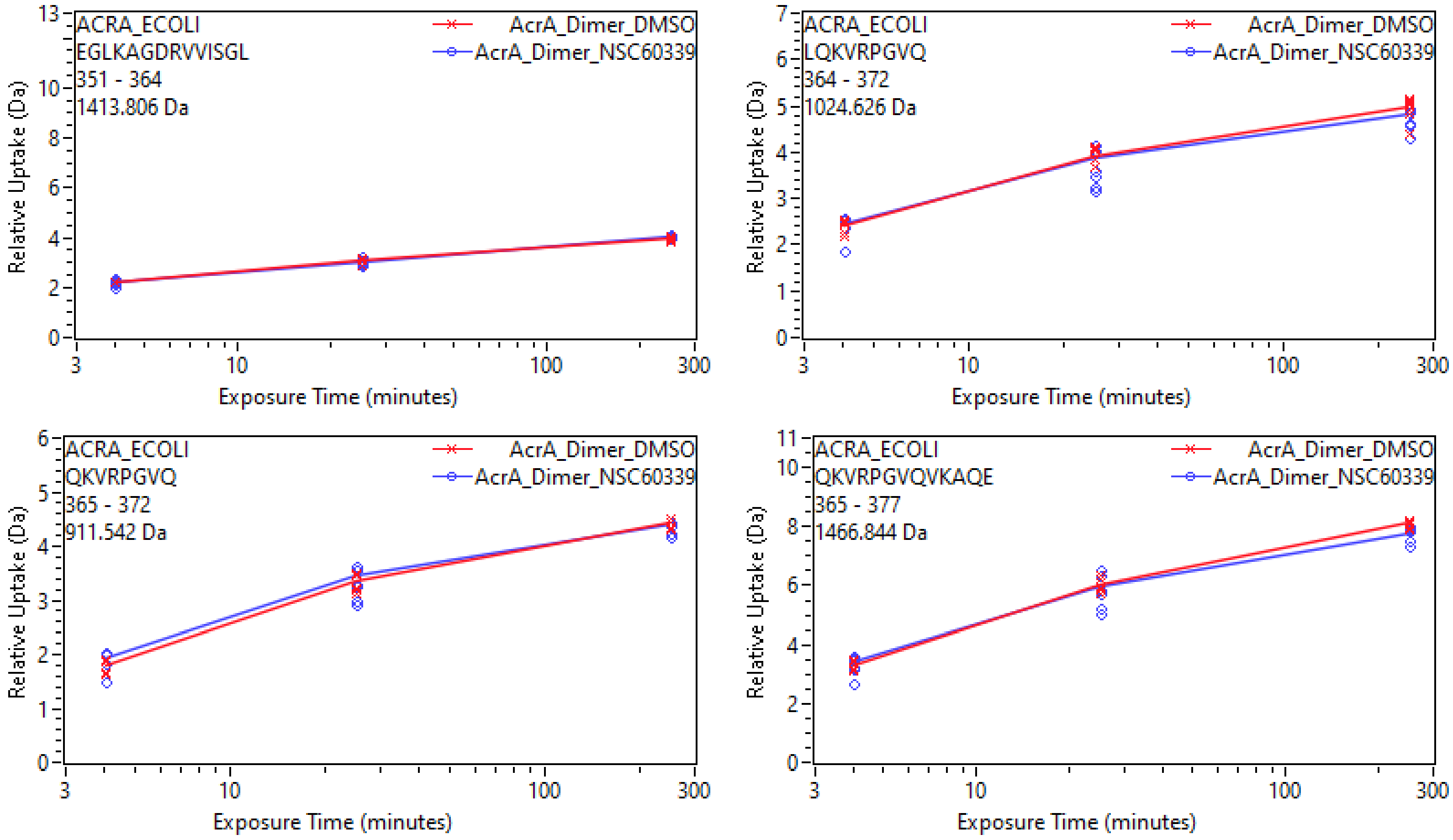
ACRA_ECOLI
EGLKAGDRVVISGL
351 - 364
1413.806 Da
AcrA_Dimer_DMSO
AcrA_Dimer_NSC60339
Relative Uptake (Da)
Exposure Time (minutes)
ACRA_ECOLI
LQKVRPGVQ
364 - 372
1024.626 Da
AcrA_Dimer_DMSO
AcrA_Dimer_NSC60339
Relative Uptake (Da)
Exposure Time (minutes)
ACRA_ECOLI
QKVRPGVQ
365 - 372
911.542 Da
AcrA_Dimer_DMSO
AcrA_Dimer_NSC60339
Relative Uptake (Da)
Exposure Time (minutes)
ACRA_ECOLI
QKVRPGVQVKAQE
365 - 377
1466.844 Da
AcrA_Dimer_DMSO
AcrA_Dimer_NSC60339
Relative Uptake (Da)
Exposure Time (minutes)

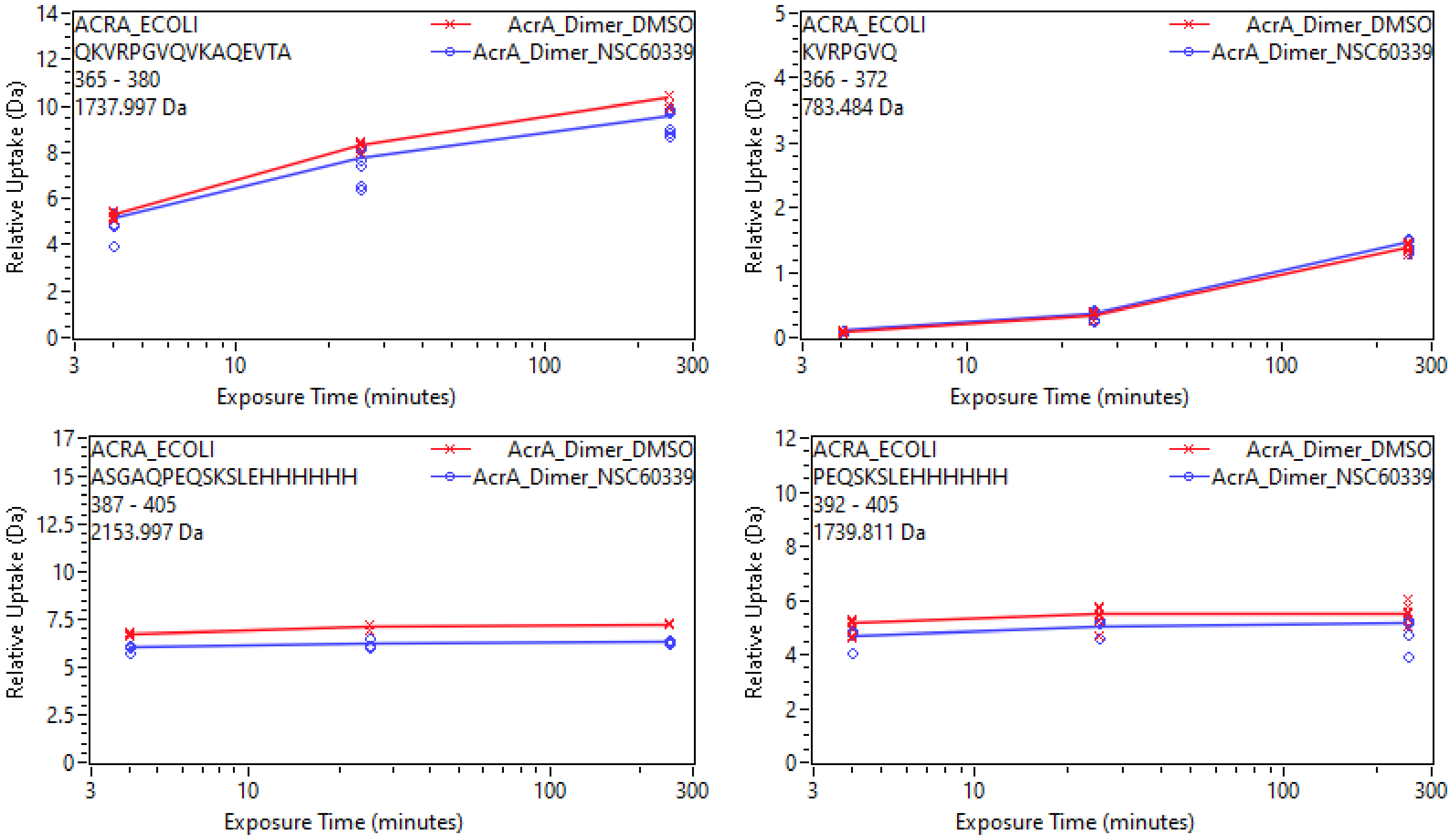
ACRA_ECOLI
QKVRPGVQVKAQEVTA
365 - 380
1737.997 Da
AcrA_Dimer_DMSO
AcrA_Dimer_NSC60339
Relative Uptake (Da)
Exposure Time (minutes)
ACRA_ECOLI
KVRPGVQ
366 - 372
783.484 Da
AcrA_Dimer_DMSO
AcrA_Dimer_NSC60339
Relative Uptake (Da)
Exposure Time (minutes)
ACRA_ECOLI
ASGAQPEQSKSLEHHHHHH
387 - 405
2153.997 Da
AcrA_Dimer_DMSO
AcrA_Dimer_NSC60339
Relative Uptake (Da)
Exposure Time (minutes)
ACRA_ECOLI
PEQSKSLEHHHHHH
392 - 405
1739.811 Da
AcrA_Dimer_DMSO
AcrA_Dimer_NSC60339
Relative Uptake (Da)
Exposure Time (minutes)
