## Supplementary material for "Conformational restriction shapes inhibition of a multidrug efflux adaptor protein": UptakePlots2

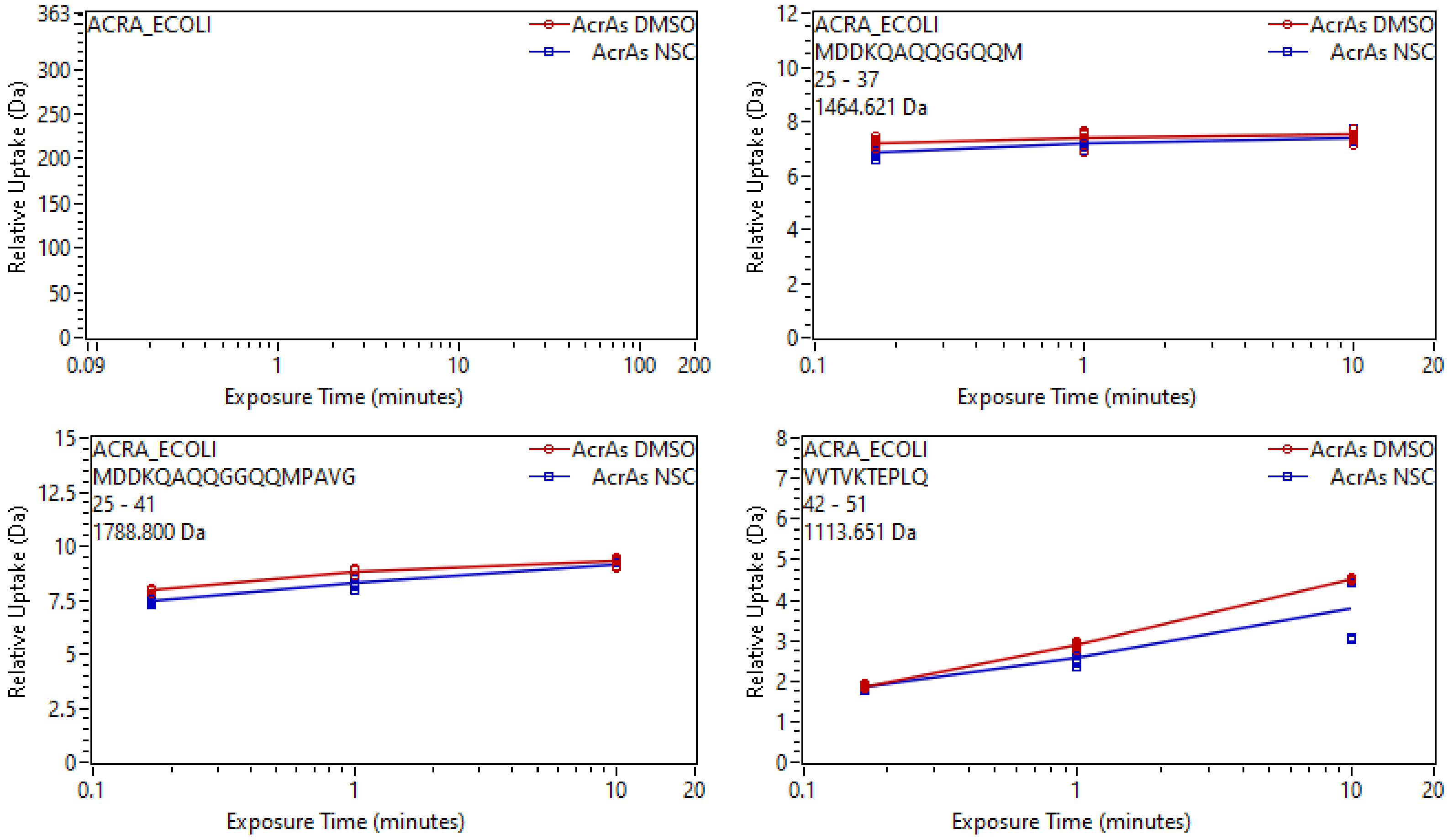

ACRA_ECOLI
AcrAs DMSO
AcrAs NSC
Relative Uptake (Da)
Exposure Time (minutes)
ACRA_ECOLI
MDDKQAQQGGQQM
25 - 37
1464.621 Da
AcrAs DMSO
AcrAs NSC
Relative Uptake (Da)
Exposure Time (minutes)
ACRA_ECOLI
MDDKQAQQGGQQMPAVG
25 - 41
1788.800 Da
AcrAs DMSO
AcrAs NSC
Relative Uptake (Da)
Exposure Time (minutes)
ACRA_ECOLI
VVTVKTEPLQ
42 - 51
1113.651 Da
AcrAs DMSO
AcrAs NSC
Relative Uptake (Da)
Exposure Time (minutes)

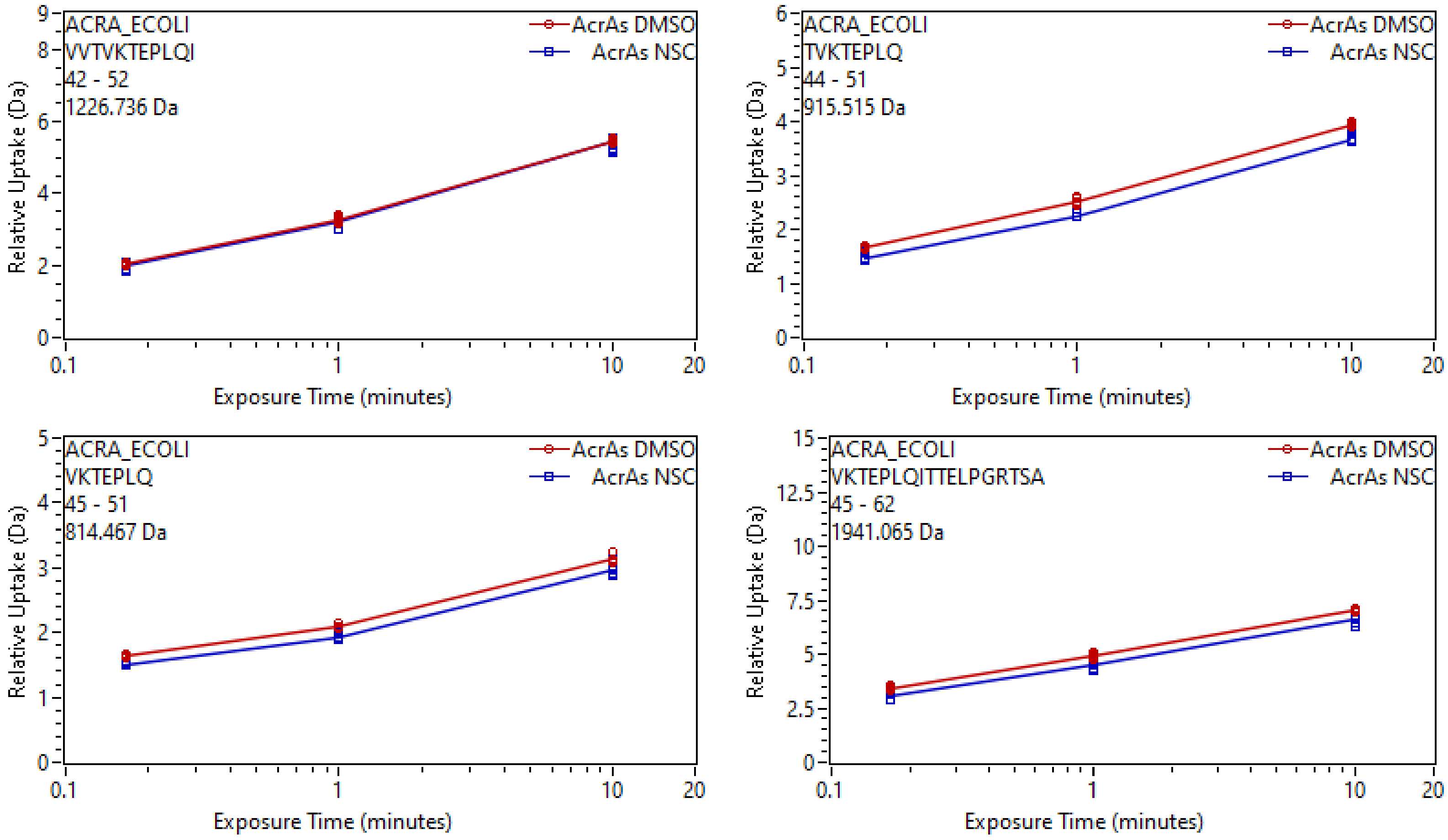
ACRA_ECOLI
VVTVKTEPLQI
42 - 52
1226.736 Da
AcrAs DMSO
AcrAs NSC
Relative Uptake (Da)
Exposure Time (minutes)
ACRA_ECOLI
TVKTEPLQ
44 - 51
915.515 Da
AcrAs DMSO
AcrAs NSC
Relative Uptake (Da)
Exposure Time (minutes)
ACRA_ECOLI
VKTEPLQ
45 - 51
814.467 Da
AcrAs DMSO
AcrAs NSC
Relative Uptake (Da)
Exposure Time (minutes)
ACRA_ECOLI
VKTEPLQITTELPGRTSA
45 - 62
1941.065 Da
AcrAs DMSO
AcrAs NSC
Relative Uptake (Da)
Exposure Time (minutes)

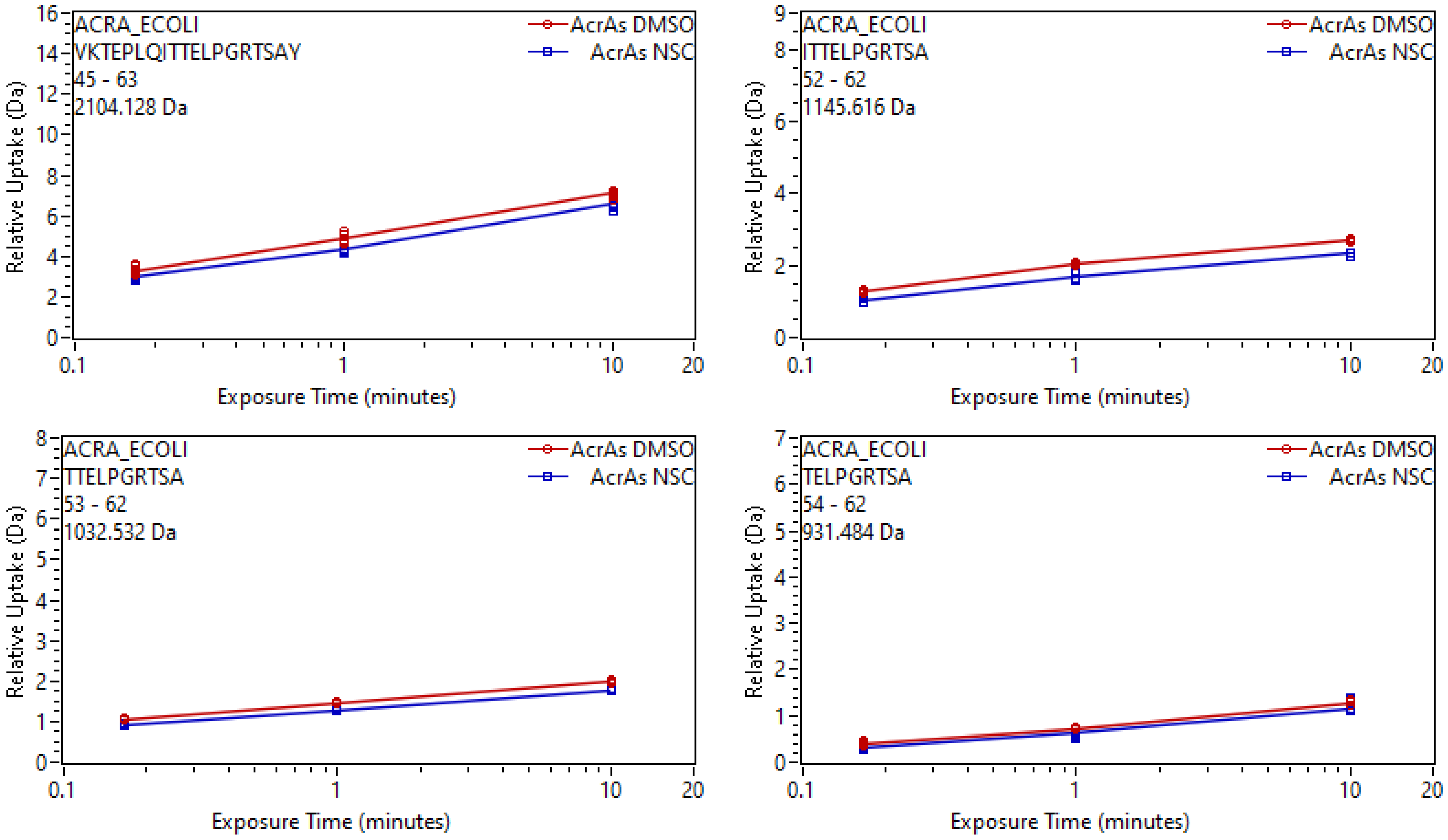
ACRA_ECOLI
VKTEPLQITTELPGRTSAY
45 - 63
2104.128 Da
AcrAs DMSO
AcrAs NSC
Relative Uptake (Da)
Exposure Time (minutes)
ACRA_ECOLI
ITTELPGRTSA
52 - 62
1145.616 Da
AcrAs DMSO
AcrAs NSC
Relative Uptake (Da)
Exposure Time (minutes)
ACRA_ECOLI
TTELPGRTSA
53 - 62
1032.532 Da
AcrAs DMSO
AcrAs NSC
Relative Uptake (Da)
Exposure Time (minutes)
ACRA_ECOLI
TELPGRTSA
54 - 62
931.484 Da
AcrAs DMSO
AcrAs NSC
Relative Uptake (Da)
Exposure Time (minutes)

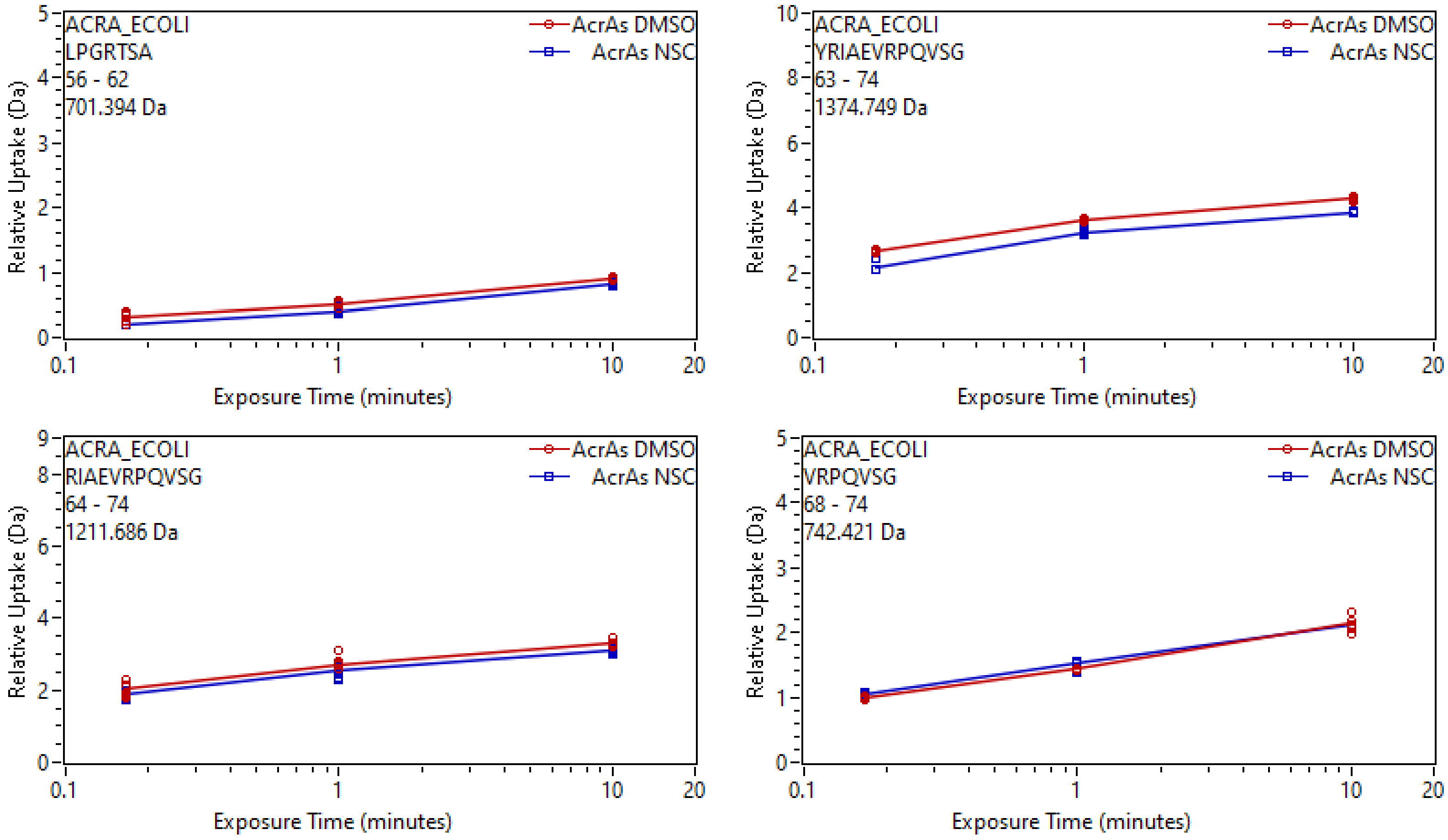
ACRA_ECOLI
LPGRTSA
56 - 62
701.394 Da
AcrAs DMSO
AcrAs NSC
Relative Uptake (Da)
Exposure Time (minutes)
ACRA_ECOLI
YRIAEVRPQVSG
63 - 74
1374.749 Da
ACRA_ECOLI
RIAEVRPQVSG
64 - 74
1211.686 Da
ACRA_ECOLI
VRPQVSG
68 - 74
742.421 Da

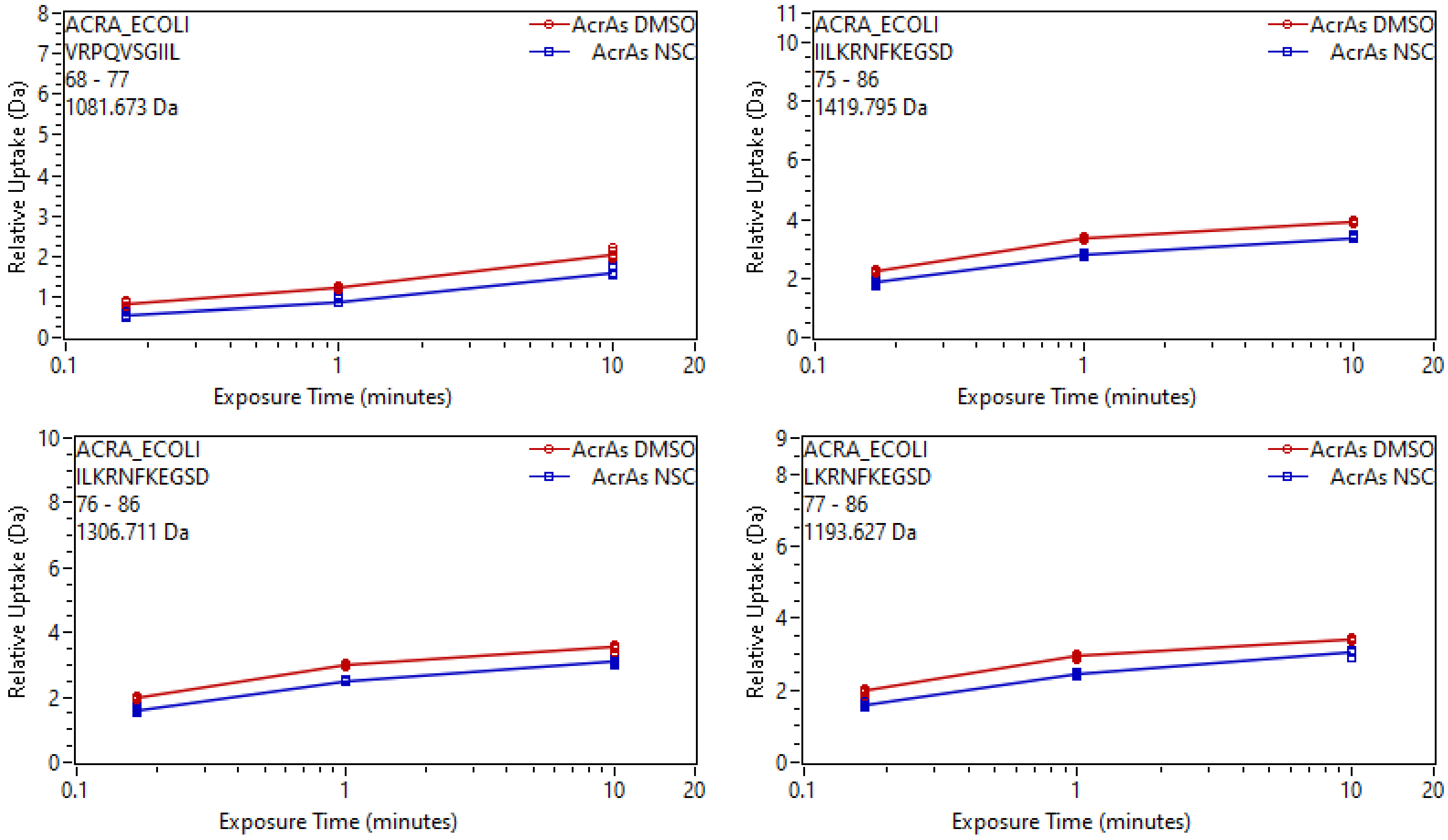
ACRA_ECOLI
VRPQVSGIIL
68 - 77
1081.673 Da
AcrAs DMSO
AcrAs NSC
Relative Uptake (Da)
Exposure Time (minutes)
ACRA_ECOLI
IILKRNFKEGSD
75 - 86
1419.795 Da
AcrAs DMSO
AcrAs NSC
Relative Uptake (Da)
Exposure Time (minutes)
ACRA_ECOLI
ILKRNFKEGSD
76 - 86
1306.711 Da
AcrAs DMSO
AcrAs NSC
Relative Uptake (Da)
Exposure Time (minutes)
ACRA_ECOLI
LKRNFKEGSD
77 - 86
1193.627 Da
AcrAs DMSO
AcrAs NSC
Relative Uptake (Da)
Exposure Time (minutes)

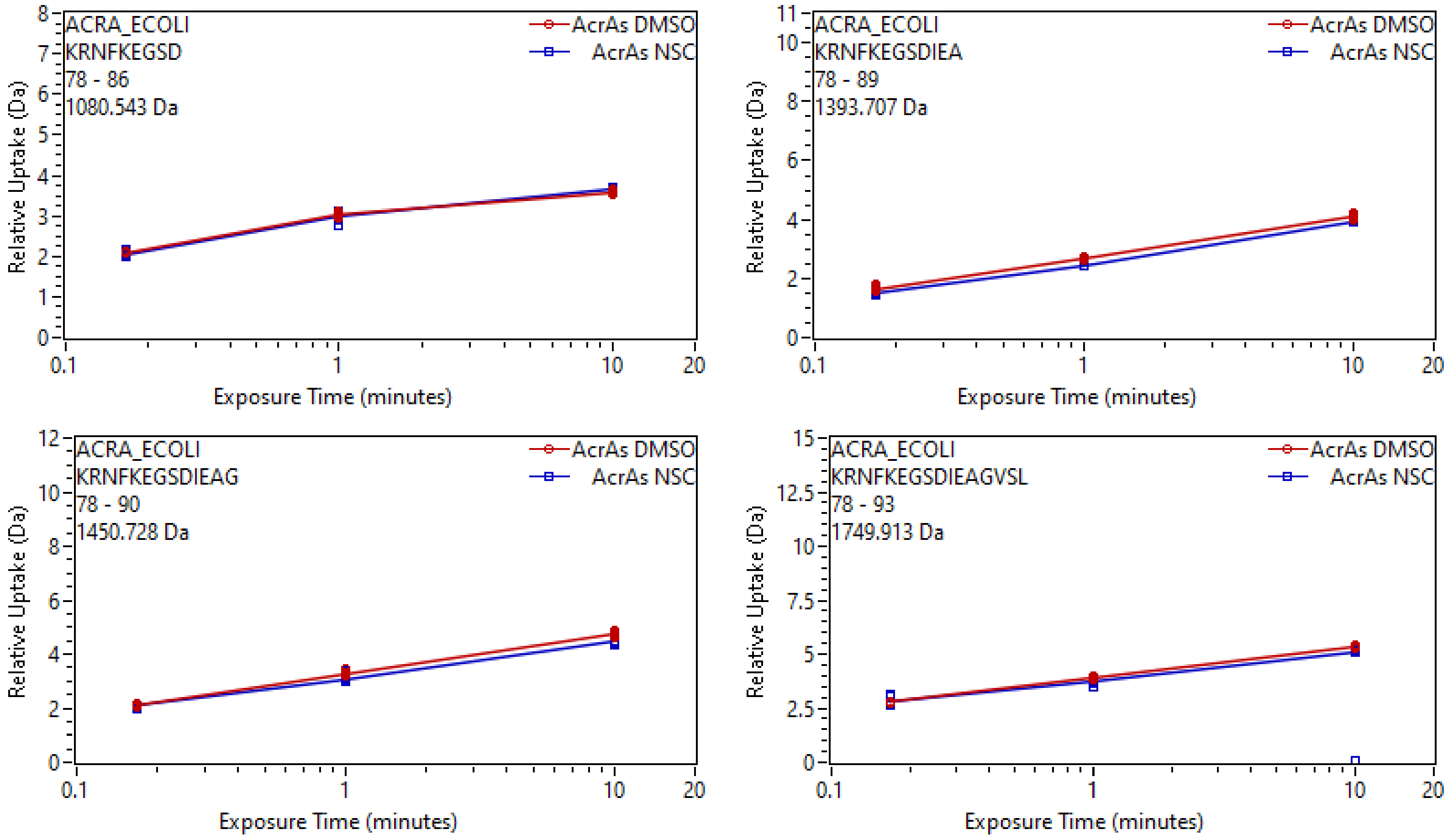

ACRA_ECOLI
KRNFKEGSD
78 - 86
1080.543 Da
AcrAs DMSO
AcrAs NSC
Relative Uptake (Da)
Exposure Time (minutes)
ACRA_ECOLI
KRNFKEGSDIEA
78 - 89
1393.707 Da
AcrAs DMSO
AcrAs NSC
Relative Uptake (Da)
Exposure Time (minutes)
ACRA_ECOLI
KRNFKEGSDIEAG
78 - 90
1450.728 Da
AcrAs DMSO
AcrAs NSC
Relative Uptake (Da)
Exposure Time (minutes)
ACRA_ECOLI
KRNFKEGSDIEAGVSL
78 - 93
1749.913 Da
AcrAs DMSO
AcrAs NSC
Relative Uptake (Da)
Exposure Time (minutes)

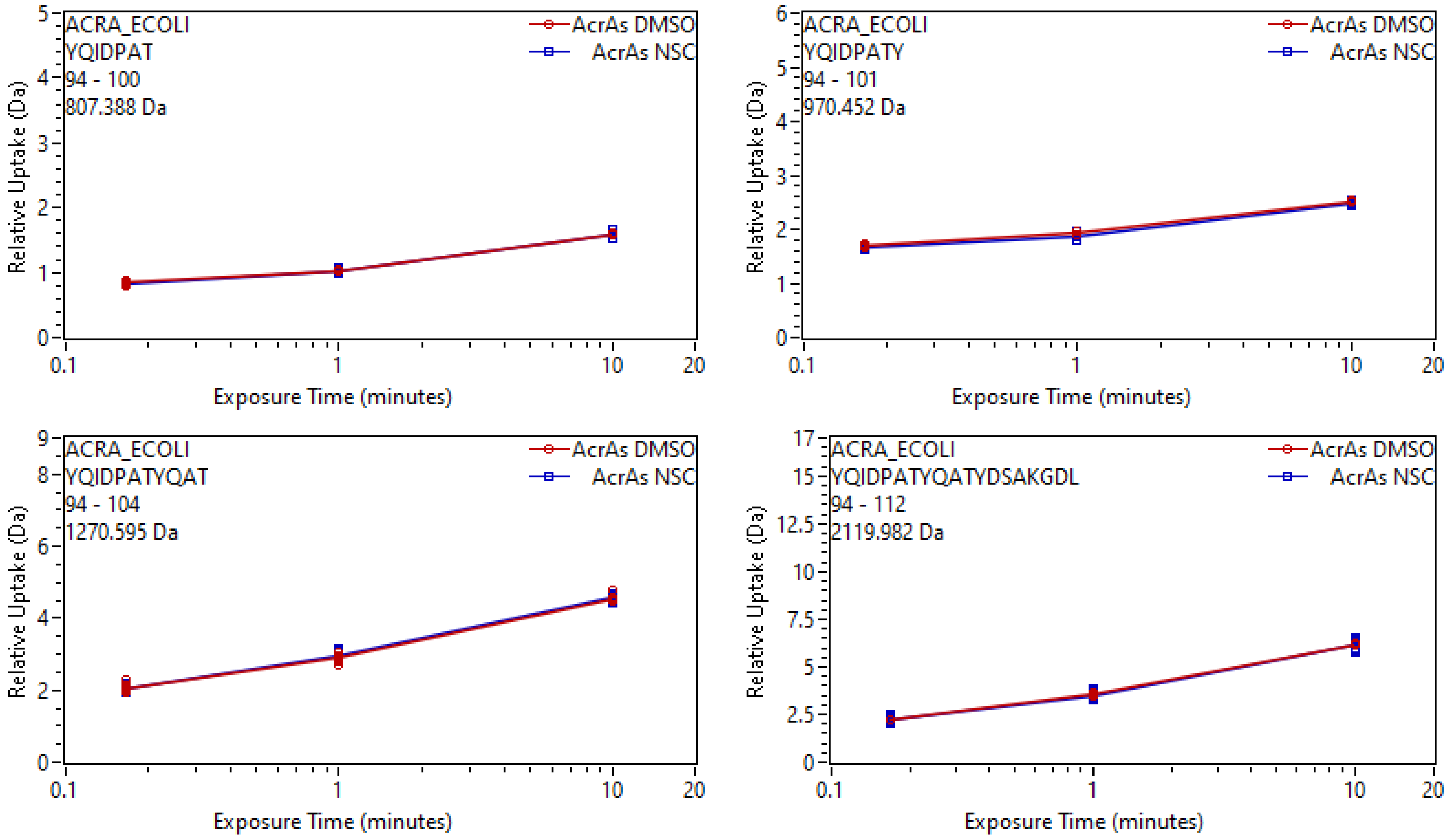
ACRA_ECOLI
YQIDPAT
94 - 100
807.388 Da
AcrAs DMSO
AcrAs NSC
Relative Uptake (Da)
Exposure Time (minutes)
ACRA_ECOLI
YQIDPATY
94 - 101
970.452 Da
AcrAs DMSO
AcrAs NSC
Relative Uptake (Da)
Exposure Time (minutes)
ACRA_ECOLI
YQIDPATYQAT
94 - 104
1270.595 Da
AcrAs DMSO
AcrAs NSC
Relative Uptake (Da)
Exposure Time (minutes)
ACRA_ECOLI
YQIDPATYQATYDSAKGDL
94 - 112
2119.982 Da
AcrAs DMSO
AcrAs NSC
Relative Uptake (Da)
Exposure Time (minutes)

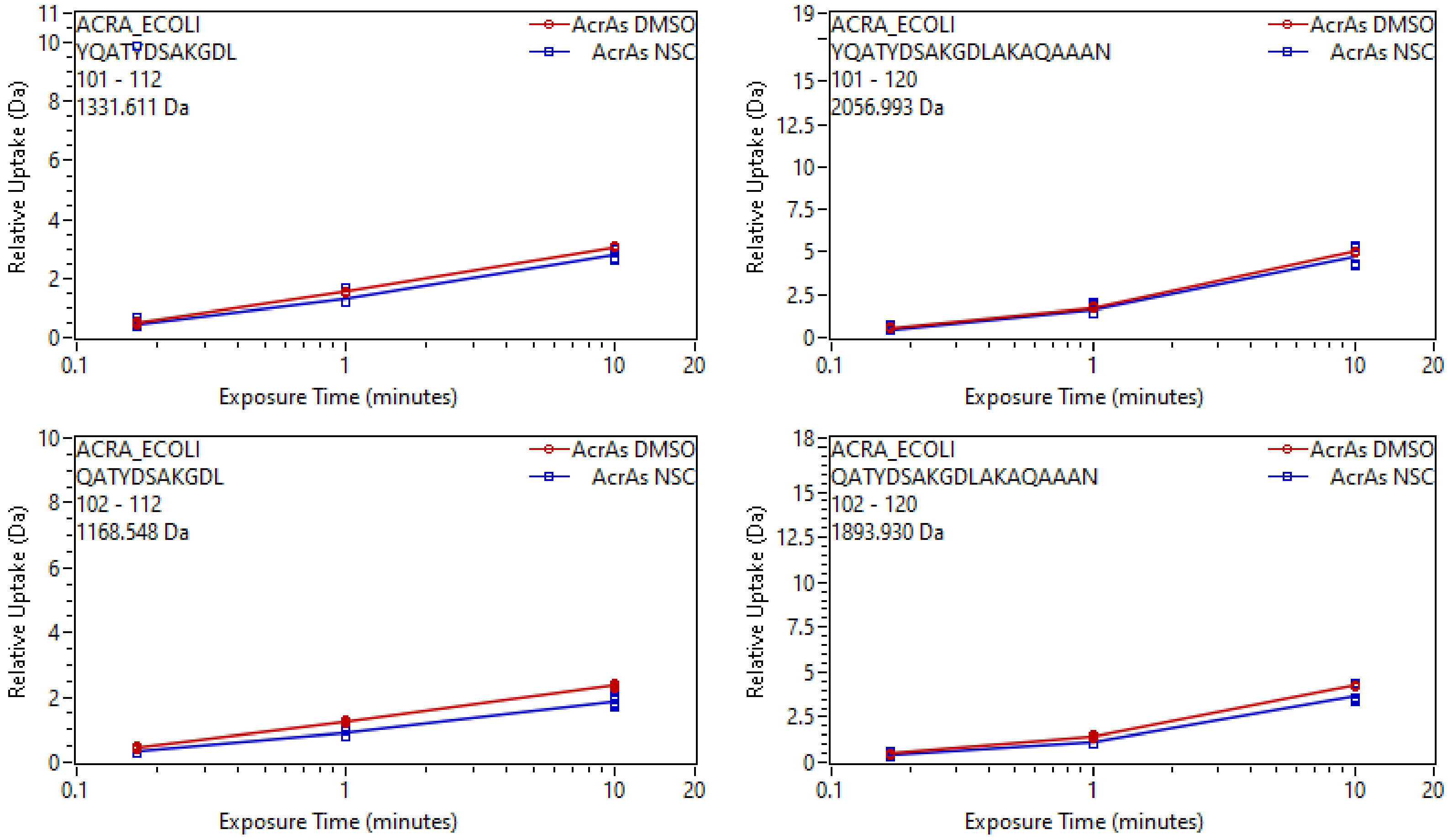
ACRA_ECOLI
YQATYDSAKGDL
101 - 112
1331.611 Da
AcrAs DMSO
AcrAs NSC
ACRA_ECOLI
YQATYDSAKGDLAKAQAAAN
101 - 120
2056.993 Da
AcrAs DMSO
AcrAs NSC
ACRA_ECOLI
QATYDSAKGDL
102 - 112
1168.548 Da
AcrAs DMSO
AcrAs NSC
ACRA_ECOLI
QATYDSAKGDLAKAQAAAN
102 - 120
1893.930 Da
AcrAs DMSO
AcrAs NSC
Relative Uptake (Da)
Exposure Time (minutes)

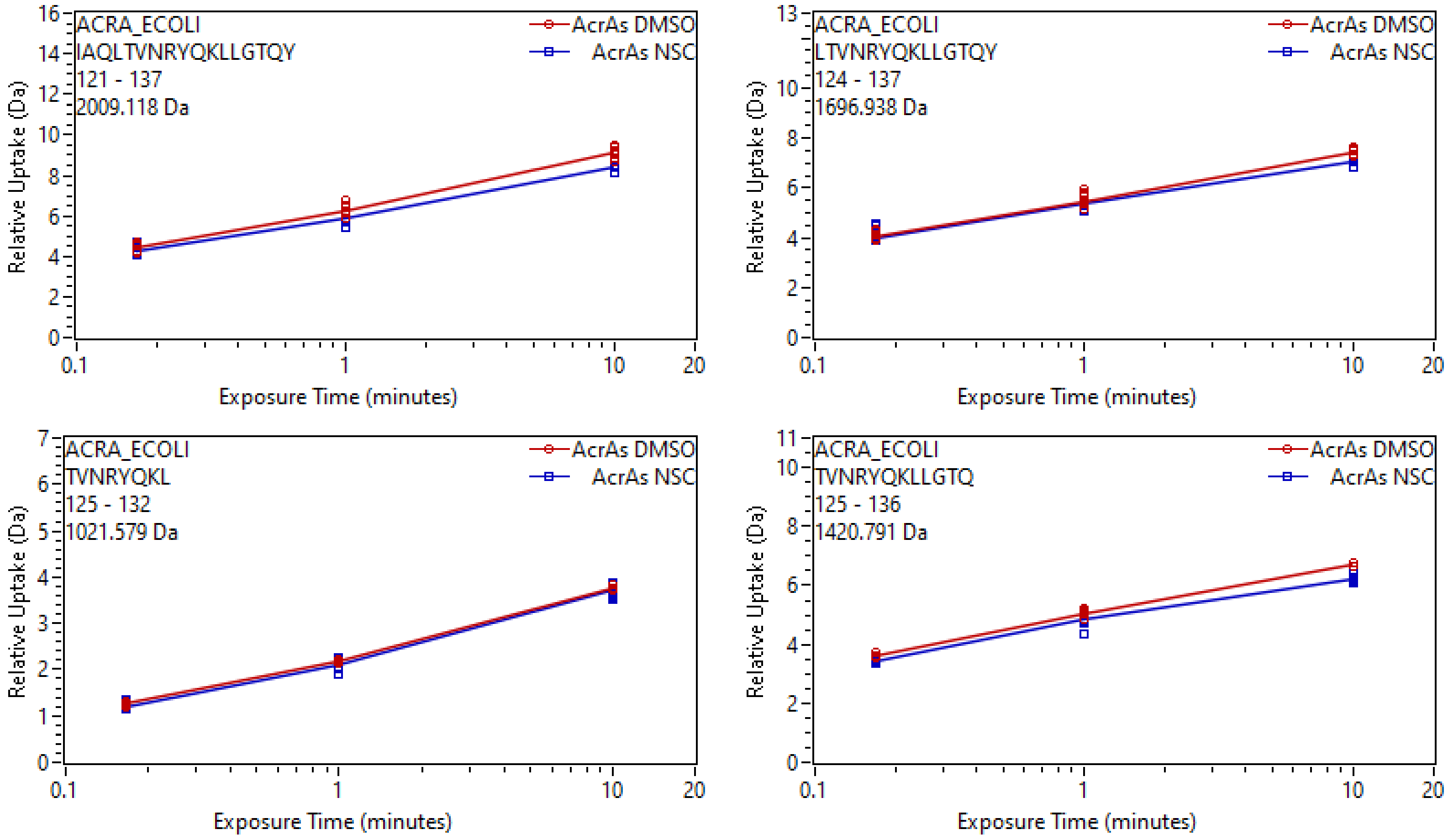
ACRA_ECOLI
IAQLTVNRYQKLLGTQY
121 - 137
2009.118 Da
AcrAs DMSO
AcrAs NSC
Relative Uptake (Da)
Exposure Time (minutes)
ACRA_ECOLI
LTVNRYQKLLGTQY
124 - 137
1696.938 Da
ACRA_ECOLI
TVNRYQKL
125 - 132
1021.579 Da
ACRA_ECOLI
TVNRYQKLLGTQ
125 - 136
1420.791 Da

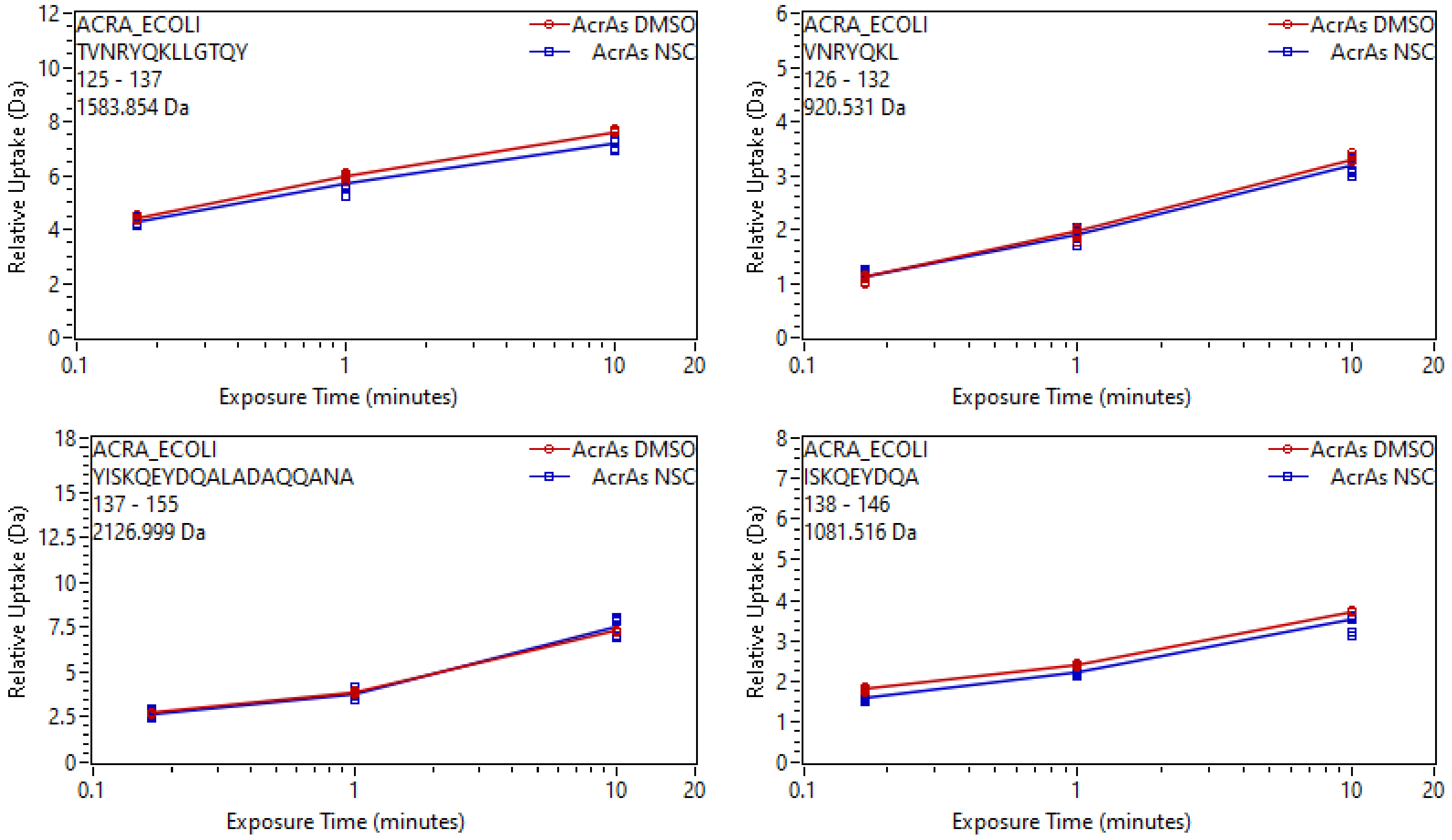
ACRA_ECOLI
TVNRYQKLLGTQY
125 - 137
1583.854 Da
AcrAs DMSO
AcrAs NSC
Relative Uptake (Da)
Exposure Time (minutes)
ACRA_ECOLI
VNRYQKL
126 - 132
920.531 Da
AcrAs DMSO
AcrAs NSC
Relative Uptake (Da)
Exposure Time (minutes)
ACRA_ECOLI
YISKQEYDQALADAQQANA
137 - 155
2126.999 Da
AcrAs DMSO
AcrAs NSC
Relative Uptake (Da)
Exposure Time (minutes)
ACRA_ECOLI
ISKQEYDQA
138 - 146
1081.516 Da
AcrAs DMSO
AcrAs NSC
Relative Uptake (Da)
Exposure Time (minutes)

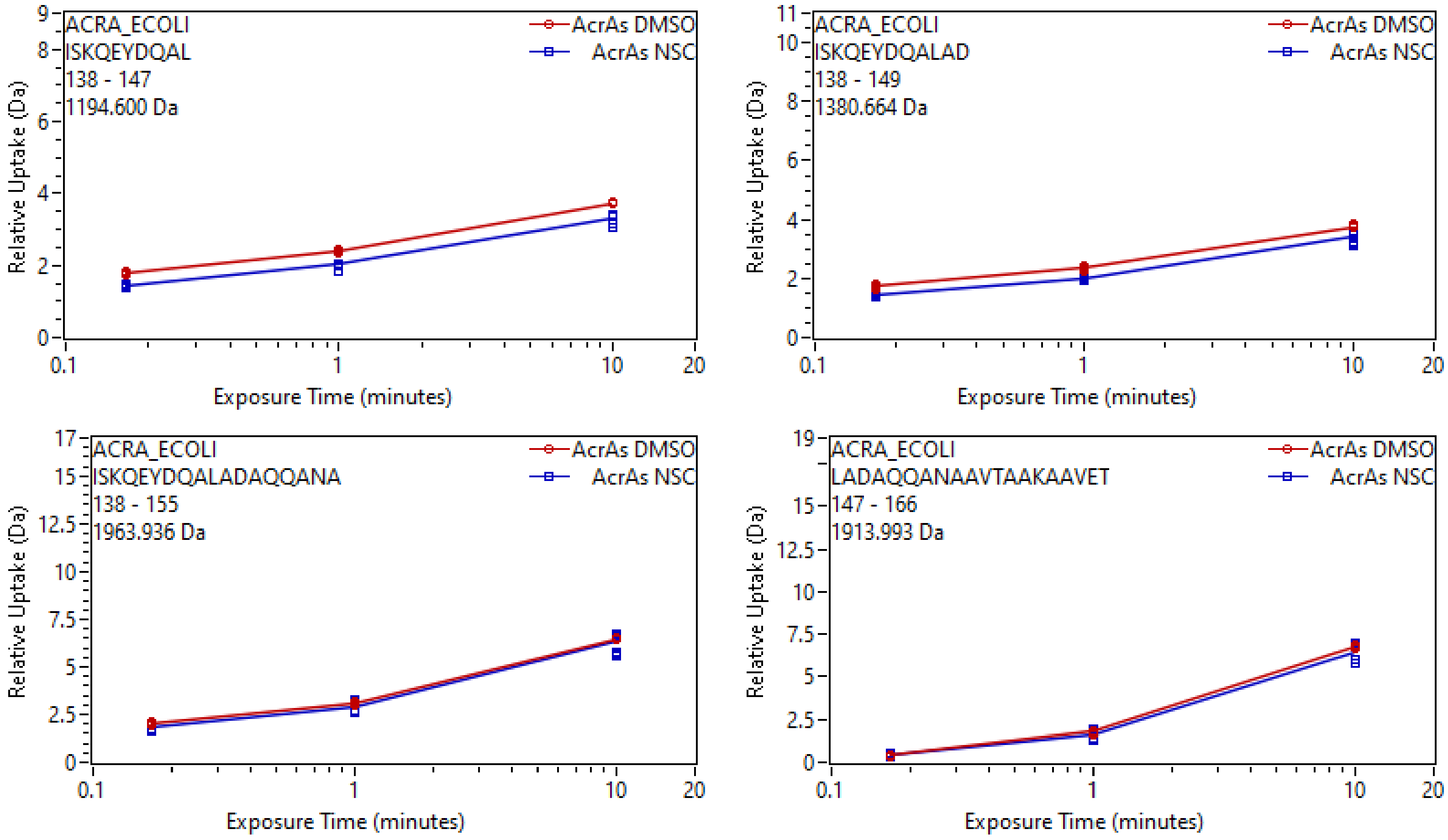
ACRA_ECOLI
ISKQEYDQAL
138 - 147
1194.600 Da
AcrAs DMSO
AcrAs NSC
Relative Uptake (Da)
Exposure Time (minutes)
ACRA_ECOLI
ISKQEYDQALAD
138 - 149
1380.664 Da
AcrAs DMSO
AcrAs NSC
Relative Uptake (Da)
Exposure Time (minutes)
ACRA_ECOLI
ISKQEYDQALADAQQANA
138 - 155
1963.936 Da
AcrAs DMSO
AcrAs NSC
Relative Uptake (Da)
Exposure Time (minutes)
ACRA_ECOLI
LADAQQANAAVTAAKAAVET
147 - 166
1913.993 Da
AcrAs DMSO
AcrAs NSC
Relative Uptake (Da)
Exposure Time (minutes)

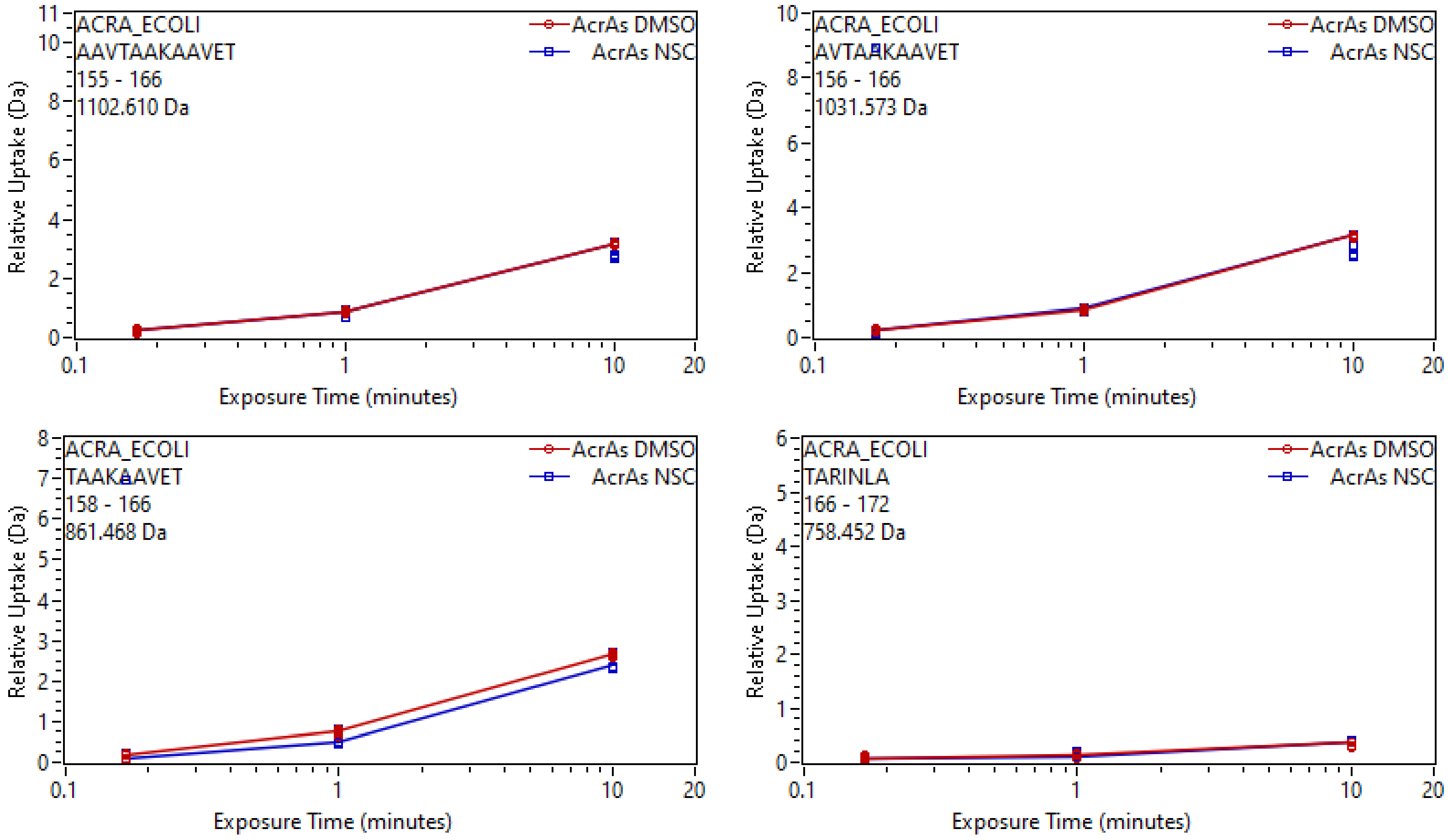
ACRA_ECOLI
AAVTAAKAAVET
155 - 166
1102.610 Da
AcrAs DMSO
AcrAs NSC
Relative Uptake (Da)
Exposure Time (minutes)
ACRA_ECOLI
AVTAAKAAVET
156 - 166
1031.573 Da
AcrAs DMSO
AcrAs NSC
Relative Uptake (Da)
Exposure Time (minutes)
ACRA_ECOLI
TAAKAAVET
158 - 166
861.468 Da
AcrAs DMSO
AcrAs NSC
Relative Uptake (Da)
Exposure Time (minutes)
ACRA_ECOLI
TARINLA
166 - 172
758.452 Da
AcrAs DMSO
AcrAs NSC
Relative Uptake (Da)
Exposure Time (minutes)

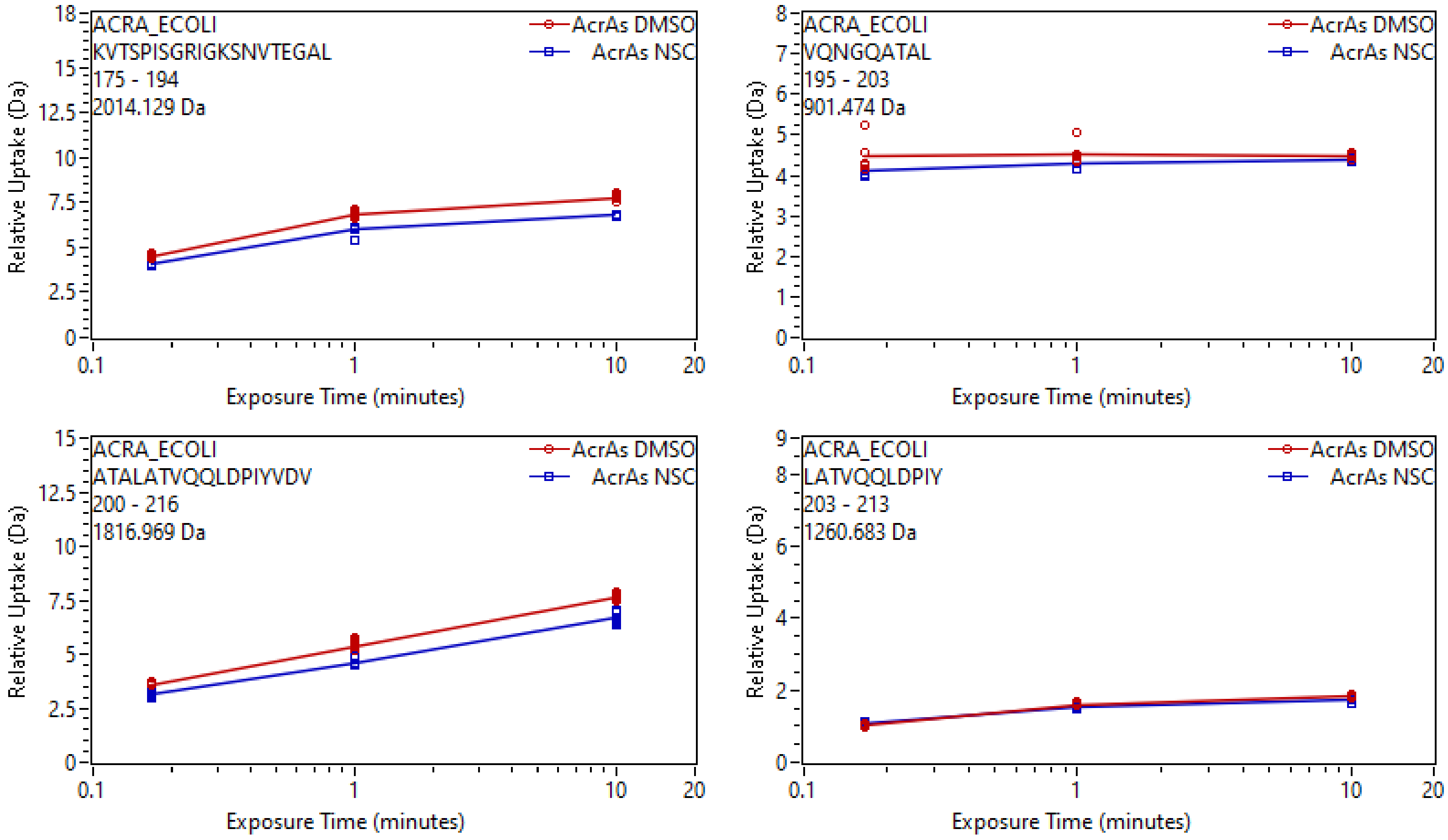
ACRA_ECOLI
KVTSPISGRIGKSNVTEGAL
175 - 194
2014.129 Da
AcrAs DMSO
AcrAs NSC
Relative Uptake (Da)
Exposure Time (minutes)
ACRA_ECOLI
VQNGQATAL
195 - 203
901.474 Da
ACRA_ECOLI
ATALATVQQLDPIYVDV
200 - 216
1816.969 Da
ACRA_ECOLI
LATVQQLDPIY
203 - 213
1260.683 Da

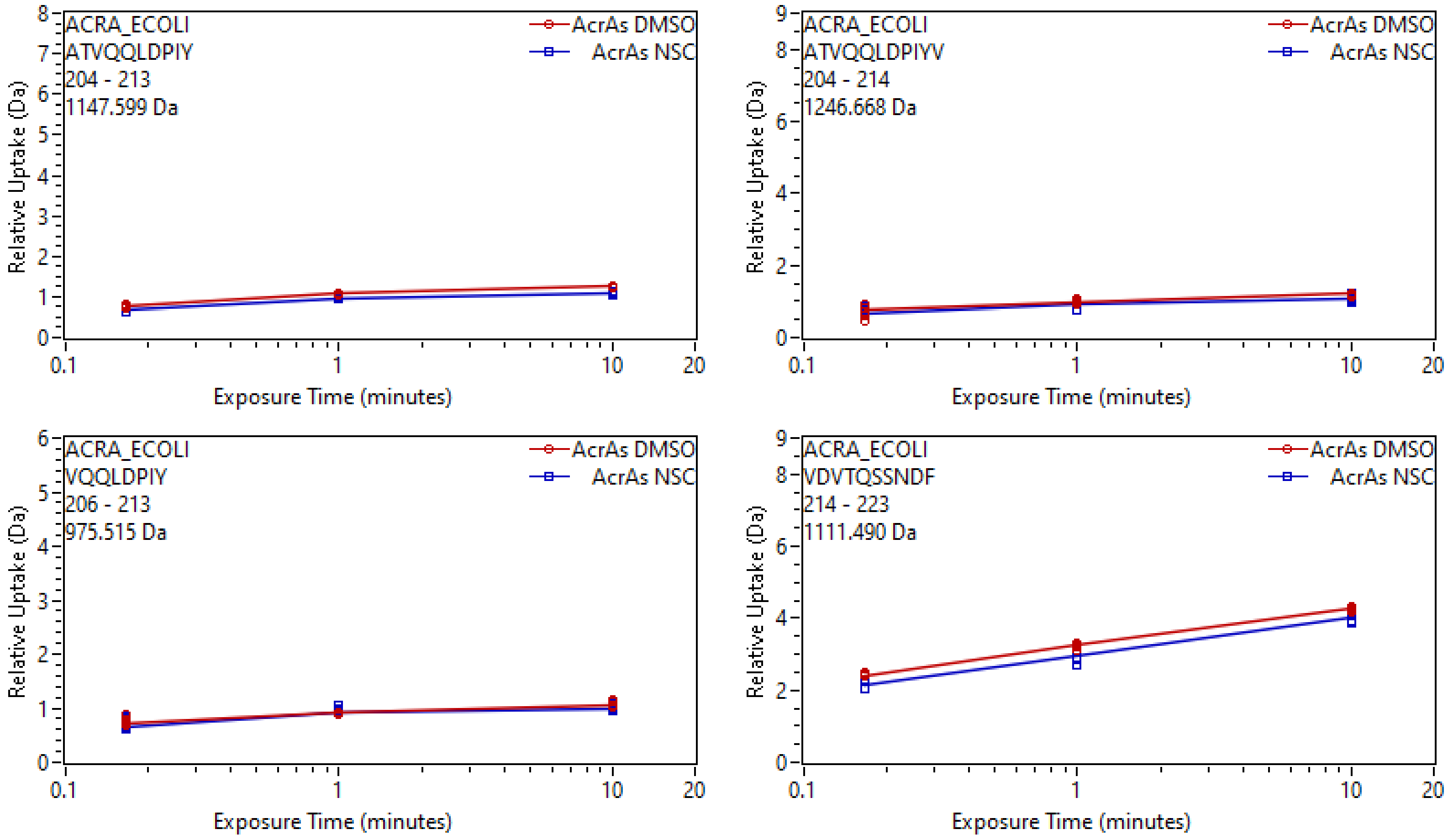
ACRA_ECOLI
ATVQQLDPIY
204 - 213
1147.599 Da
AcrAs DMSO
AcrAs NSC
Relative Uptake (Da)
Exposure Time (minutes)
ACRA_ECOLI
ATVQQLDPIYV
204 - 214
1246.668 Da
AcrAs DMSO
AcrAs NSC
Relative Uptake (Da)
Exposure Time (minutes)
ACRA_ECOLI
VQQLDPIY
206 - 213
975.515 Da
AcrAs DMSO
AcrAs NSC
Relative Uptake (Da)
Exposure Time (minutes)
ACRA_ECOLI
VDVTQSSNDF
214 - 223
1111.490 Da
AcrAs DMSO
AcrAs NSC
Relative Uptake (Da)
Exposure Time (minutes)

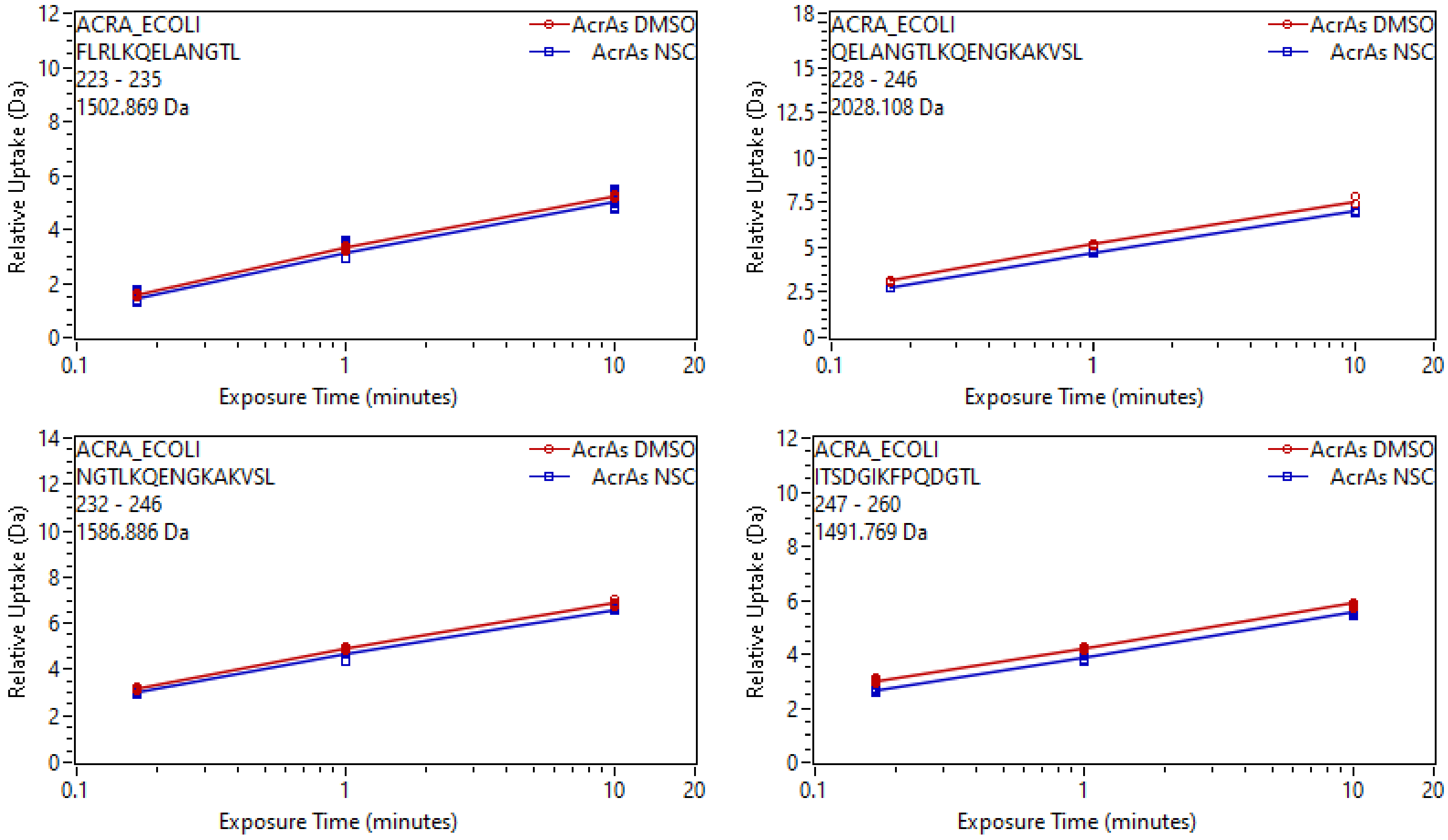
ACRA_ECOLI
FLRLKQELANGTL
223 - 235
1502.869 Da
AcrAs DMSO
AcrAs NSC
Relative Uptake (Da)
Exposure Time (minutes)
ACRA_ECOLI
QELANGTLKQENGKAKVSL
228 - 246
2028.108 Da
AcrAs DMSO
AcrAs NSC
Relative Uptake (Da)
Exposure Time (minutes)
ACRA_ECOLI
NGTLKQENGKAKVSL
232 - 246
1586.886 Da
AcrAs DMSO
AcrAs NSC
Relative Uptake (Da)
Exposure Time (minutes)
ACRA_ECOLI
ITSDGIKFPQDGTL
247 - 260
1491.769 Da
AcrAs DMSO
AcrAs NSC
Relative Uptake (Da)
Exposure Time (minutes)

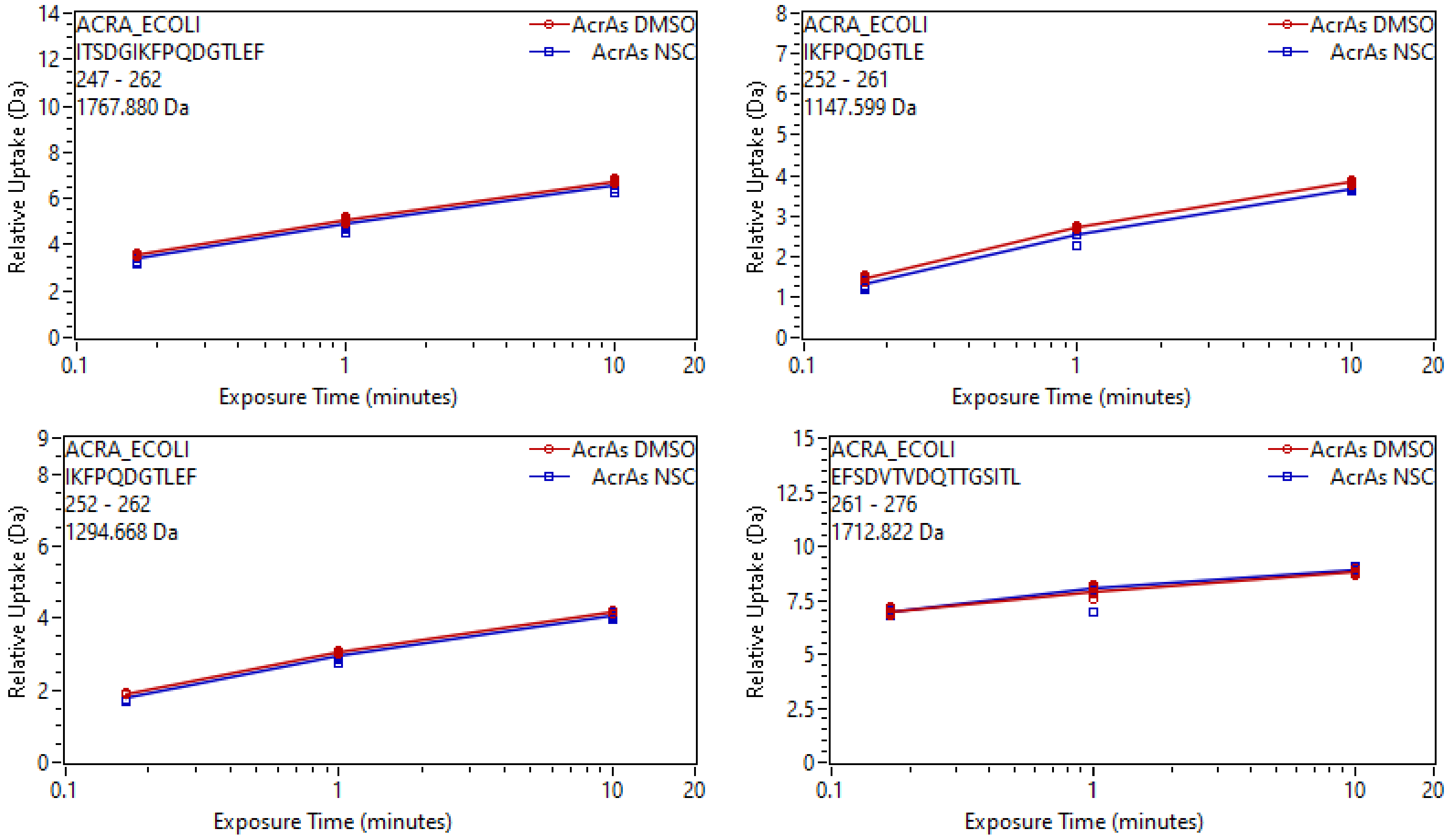
ACRA_ECOLI
ITSDGIKFPQDGTLEF
247 - 262
1767.880 Da
AcrAs DMSO
AcrAs NSC
Relative Uptake (Da)
Exposure Time (minutes)
ACRA_ECOLI
IKFPQDGTLE
252 - 261
1147.599 Da
AcrAs DMSO
AcrAs NSC
Relative Uptake (Da)
Exposure Time (minutes)
ACRA_ECOLI
IKFPQDGTLEF
252 - 262
1294.668 Da
AcrAs DMSO
AcrAs NSC
Relative Uptake (Da)
Exposure Time (minutes)
ACRA_ECOLI
EFSDVTVDQTTGSITL
261 - 276
1712.822 Da
AcrAs DMSO
AcrAs NSC
Relative Uptake (Da)
Exposure Time (minutes)

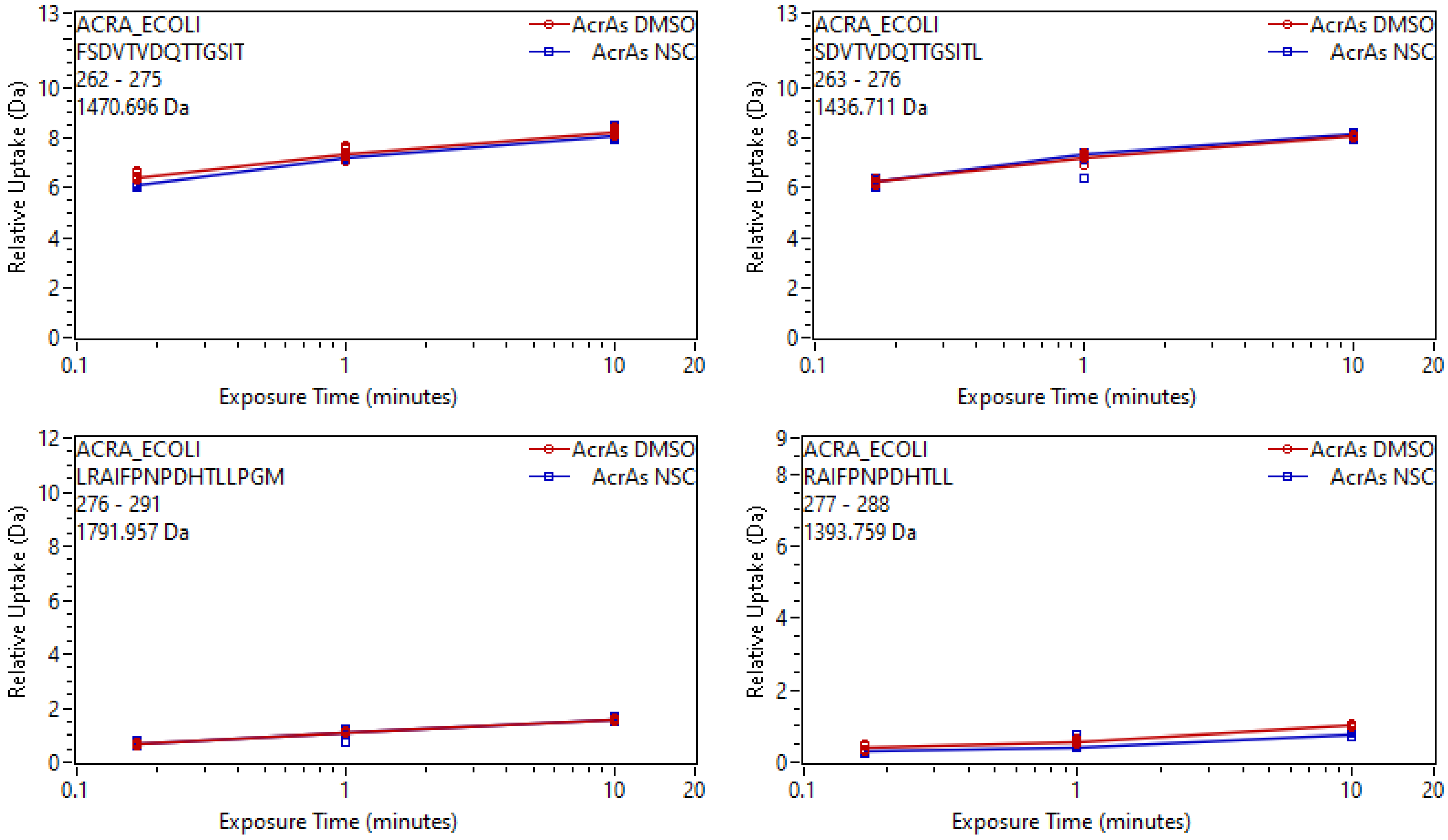
ACRA_ECOLI
FSDVTVDQTTGSIT
262 - 275
1470.696 Da
AcrAs DMSO
AcrAs NSC
Relative Uptake (Da)
Exposure Time (minutes)
ACRA_ECOLI
SDVTVDQTTGSITL
263 - 276
1436.711 Da
AcrAs DMSO
AcrAs NSC
Relative Uptake (Da)
Exposure Time (minutes)
ACRA_ECOLI
LRAIFPNPDHTLLPGM
276 - 291
1791.957 Da
AcrAs DMSO
AcrAs NSC
Relative Uptake (Da)
Exposure Time (minutes)
ACRA_ECOLI
RAIFPNPDHTLL
277 - 288
1393.759 Da
AcrAs DMSO
AcrAs NSC
Relative Uptake (Da)
Exposure Time (minutes)

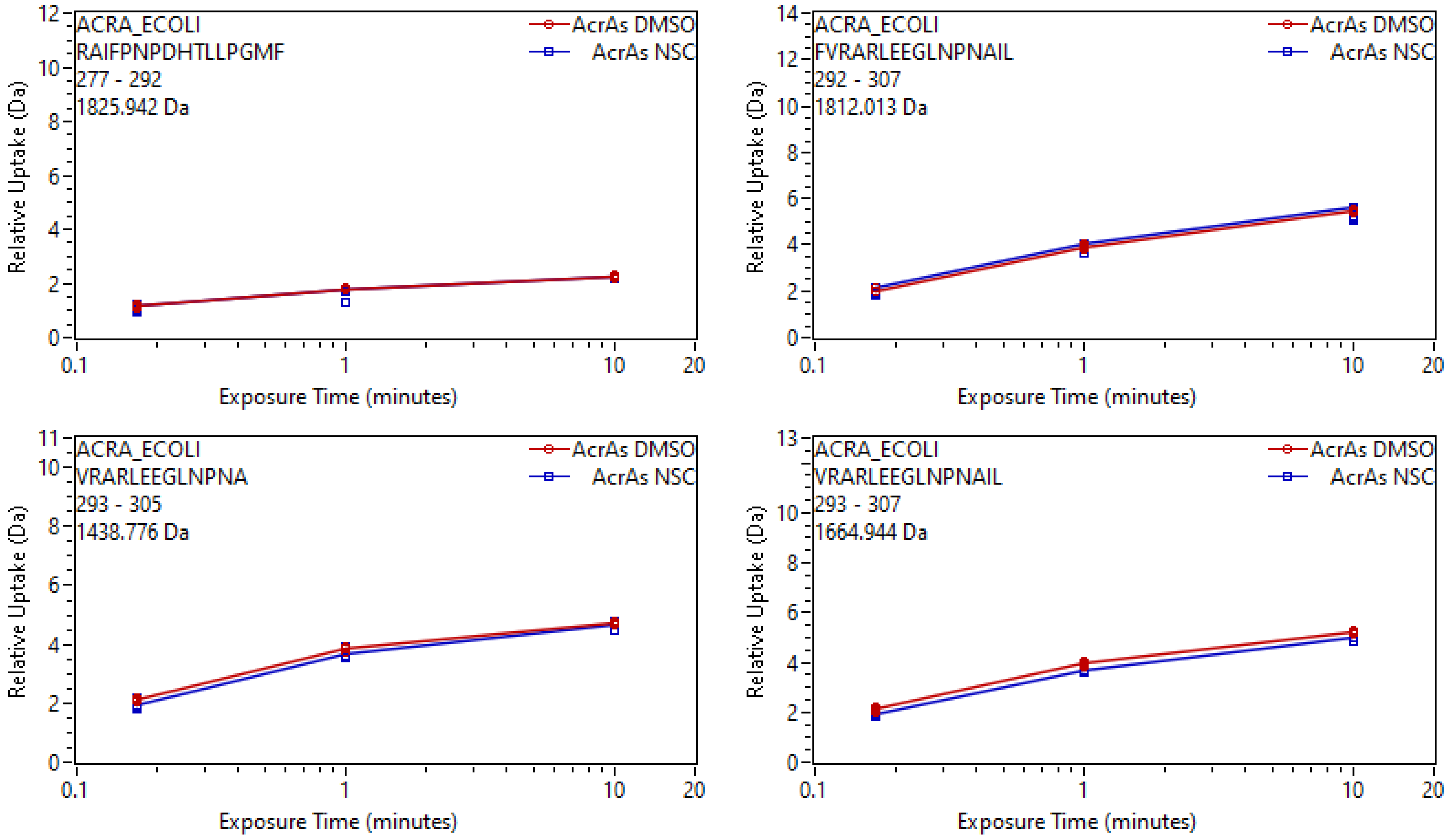
ACRA_ECOLI
RAIFPNPDHTLLPGMF
277 - 292
1825.942 Da
AcrAs DMSO
AcrAs NSC
Relative Uptake (Da)
Exposure Time (minutes)
ACRA_ECOLI
FVRARLEEGLNPNAIL
292 - 307
1812.013 Da
ACRA_ECOLI
VRARLEEGLNPNA
293 - 305
1438.776 Da
ACRA_ECOLI
VRARLEEGLNPNAIL
293 - 307
1664.944 Da

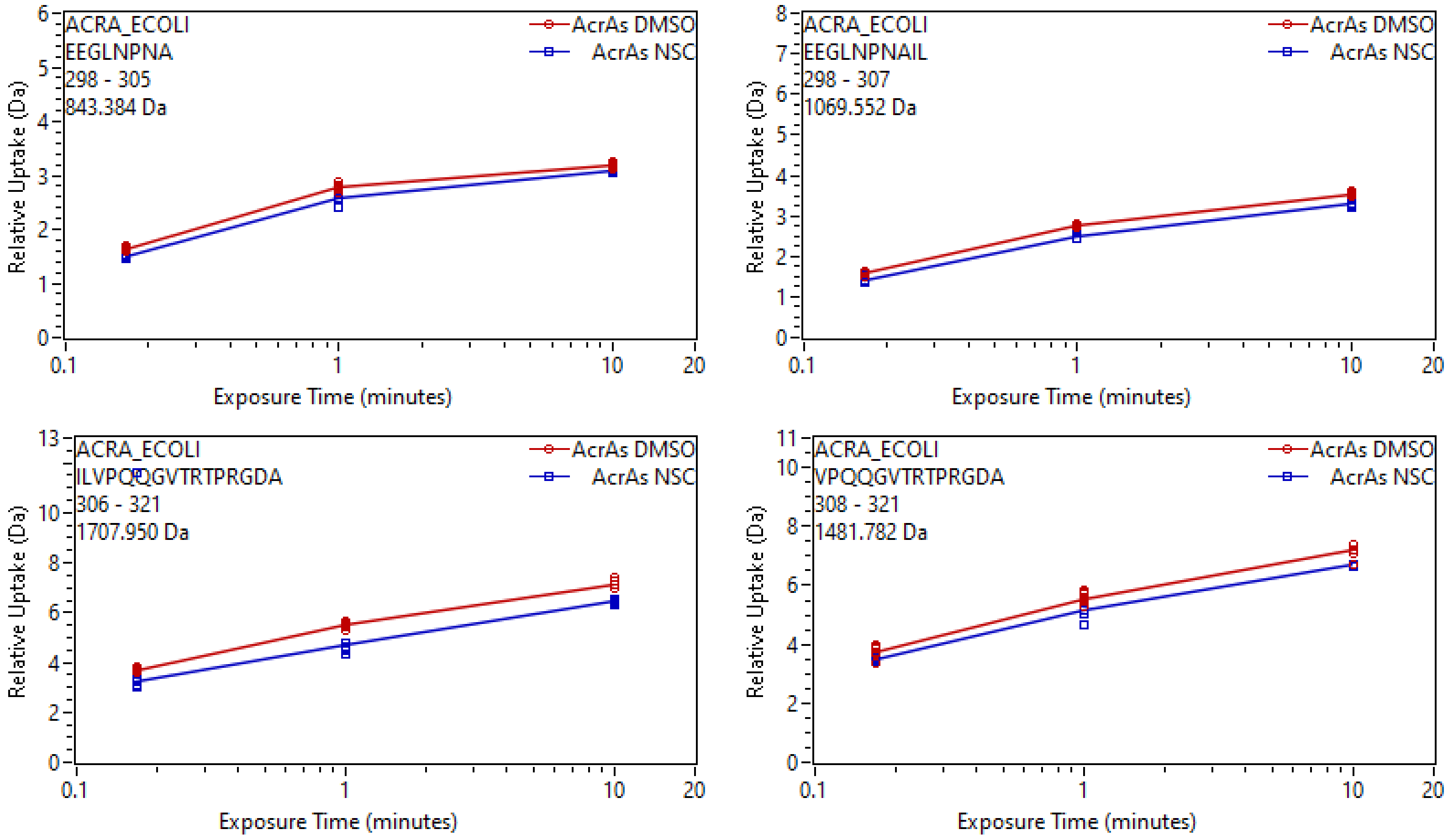

ACRA_ECOLI
EEGLNPNA
298 - 305
843.384 Da
AcrAs DMSO
AcrAs NSC
ACRA_ECOLI
EEGLNPNAIL
298 - 307
1069.552 Da
ACRA_ECOLI
ILVPQQGVTRTPRGDA
306 - 321
1707.950 Da
ACRA_ECOLI
VPQQGVTRTPRGDA
308 - 321
1481.782 Da
Relative Uptake (Da)
Exposure Time (minutes)

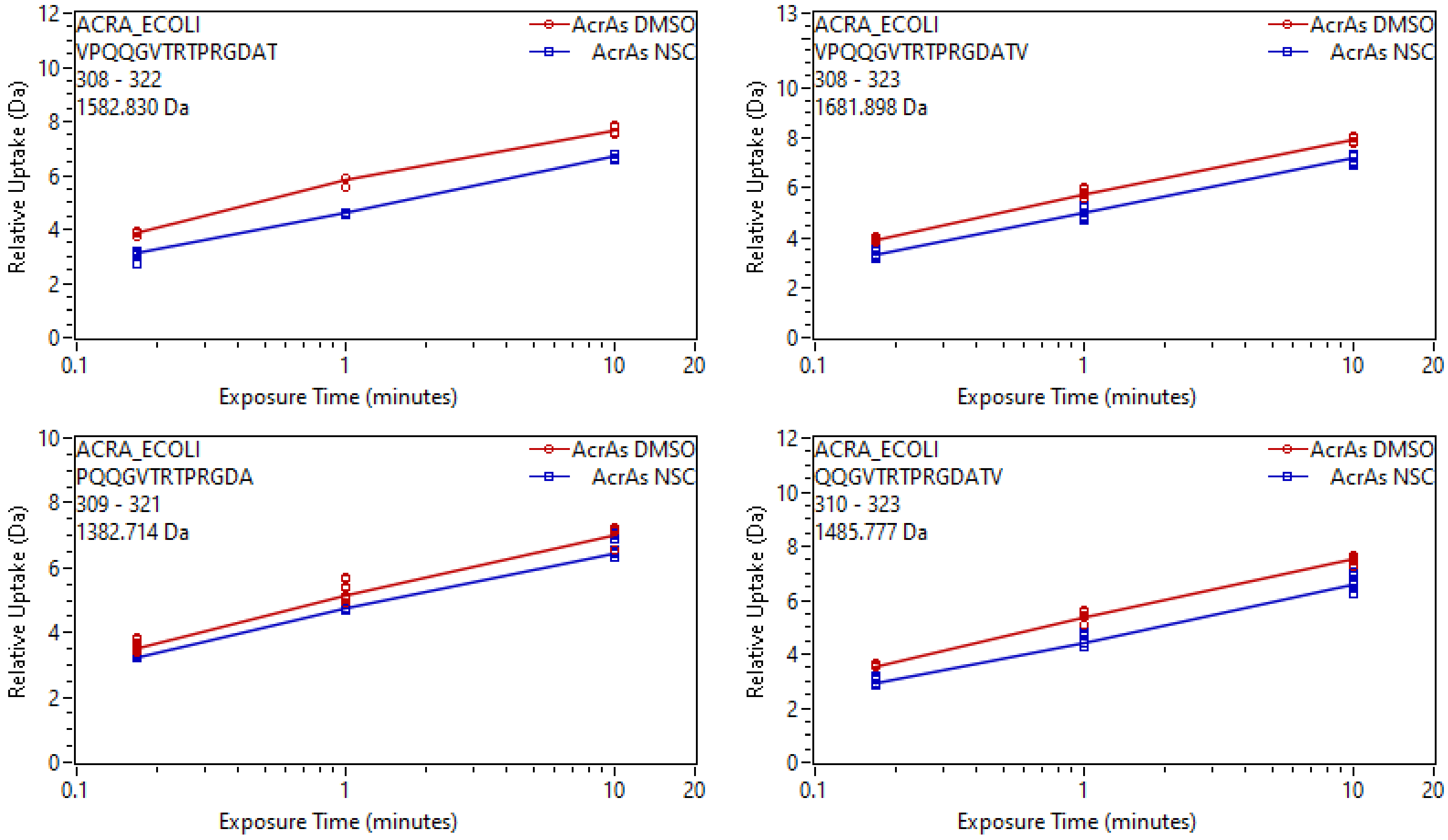

ACRA_ECOLI
VPQQGVTRTPRGDAT
308 - 322
1582.830 Da
AcrAs DMSO
AcrAs NSC
Relative Uptake (Da)
Exposure Time (minutes)
ACRA_ECOLI
VPQQGVTRTPRGDATV
308 - 323
1681.898 Da
ACRA_ECOLI
PQQGVTRTPRGDA
309 - 321
1382.714 Da
ACRA_ECOLI
QQGVTRTPRGDATV
310 - 323
1485.777 Da

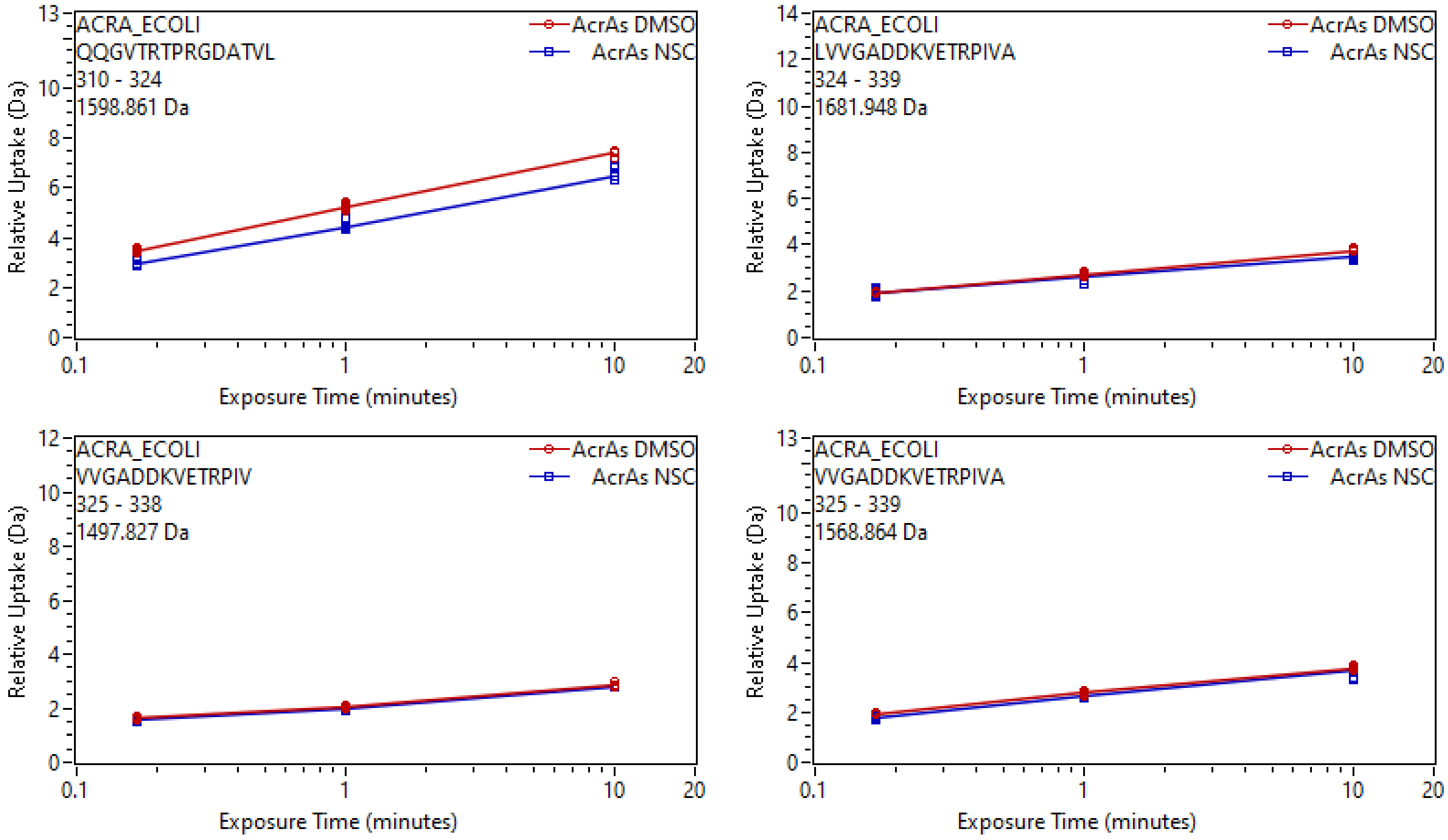
ACRA_ECOLI
QQGVTRTPRGDATVL
310 - 324
1598.861 Da
AcrAs DMSO
AcrAs NSC
Relative Uptake (Da)
Exposure Time (minutes)
ACRA_ECOLI
LVVGADDKVETRPIVA
324 - 339
1681.948 Da
ACRA_ECOLI
VVGADDKVETRPIV
325 - 338
1497.827 Da
ACRA_ECOLI
VVGADDKVETRPIVA
325 - 339
1568.864 Da

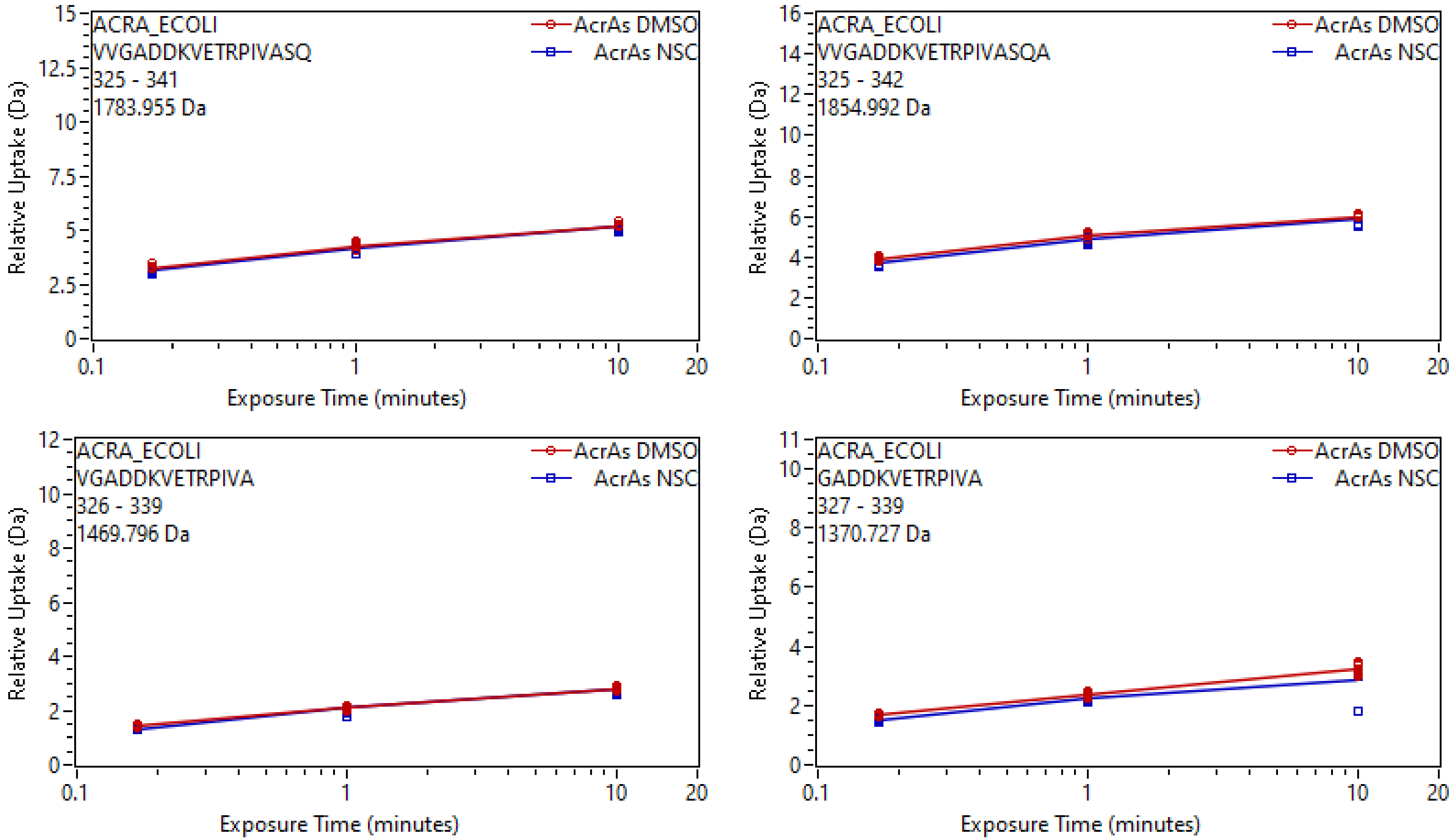
ACRA_ECOLI
VVGADDKVETRPIVASQ
325 - 341
1783.955 Da
AcrAs DMSO
AcrAs NSC
Relative Uptake (Da)
Exposure Time (minutes)
ACRA_ECOLI
VVGADDKVETRPIVASQA
325 - 342
1854.992 Da
AcrAs DMSO
AcrAs NSC
Relative Uptake (Da)
Exposure Time (minutes)
ACRA_ECOLI
VGADDKVETRPIVA
326 - 339
1469.796 Da
AcrAs DMSO
AcrAs NSC
Relative Uptake (Da)
Exposure Time (minutes)
ACRA_ECOLI
GADDKVETRPIVA
327 - 339
1370.727 Da
AcrAs DMSO
AcrAs NSC
Relative Uptake (Da)
Exposure Time (minutes)

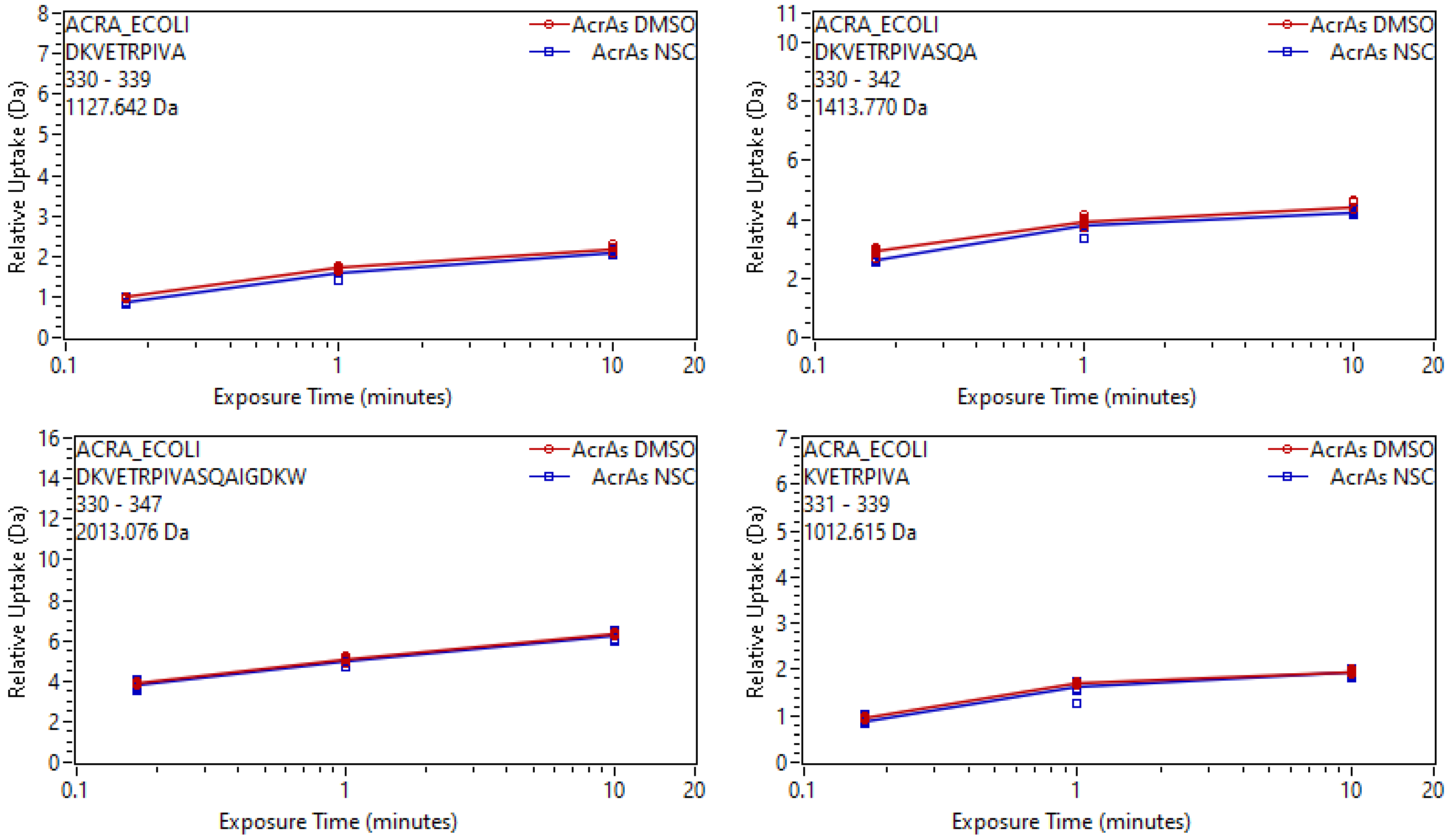
ACRA_ECOLI
DKVETRPIVA
330 - 339
1127.642 Da
AcrAs DMSO
AcrAs NSC
Relative Uptake (Da)
Exposure Time (minutes)
ACRA_ECOLI
DKVETRPIVASQA
330 - 342
1413.770 Da
AcrAs DMSO
AcrAs NSC
Relative Uptake (Da)
Exposure Time (minutes)
ACRA_ECOLI
DKVETRPIVASQAIGDKW
330 - 347
2013.076 Da
AcrAs DMSO
AcrAs NSC
Relative Uptake (Da)
Exposure Time (minutes)
ACRA_ECOLI
KVETRPIVA
331 - 339
1012.615 Da
AcrAs DMSO
AcrAs NSC
Relative Uptake (Da)
Exposure Time (minutes)

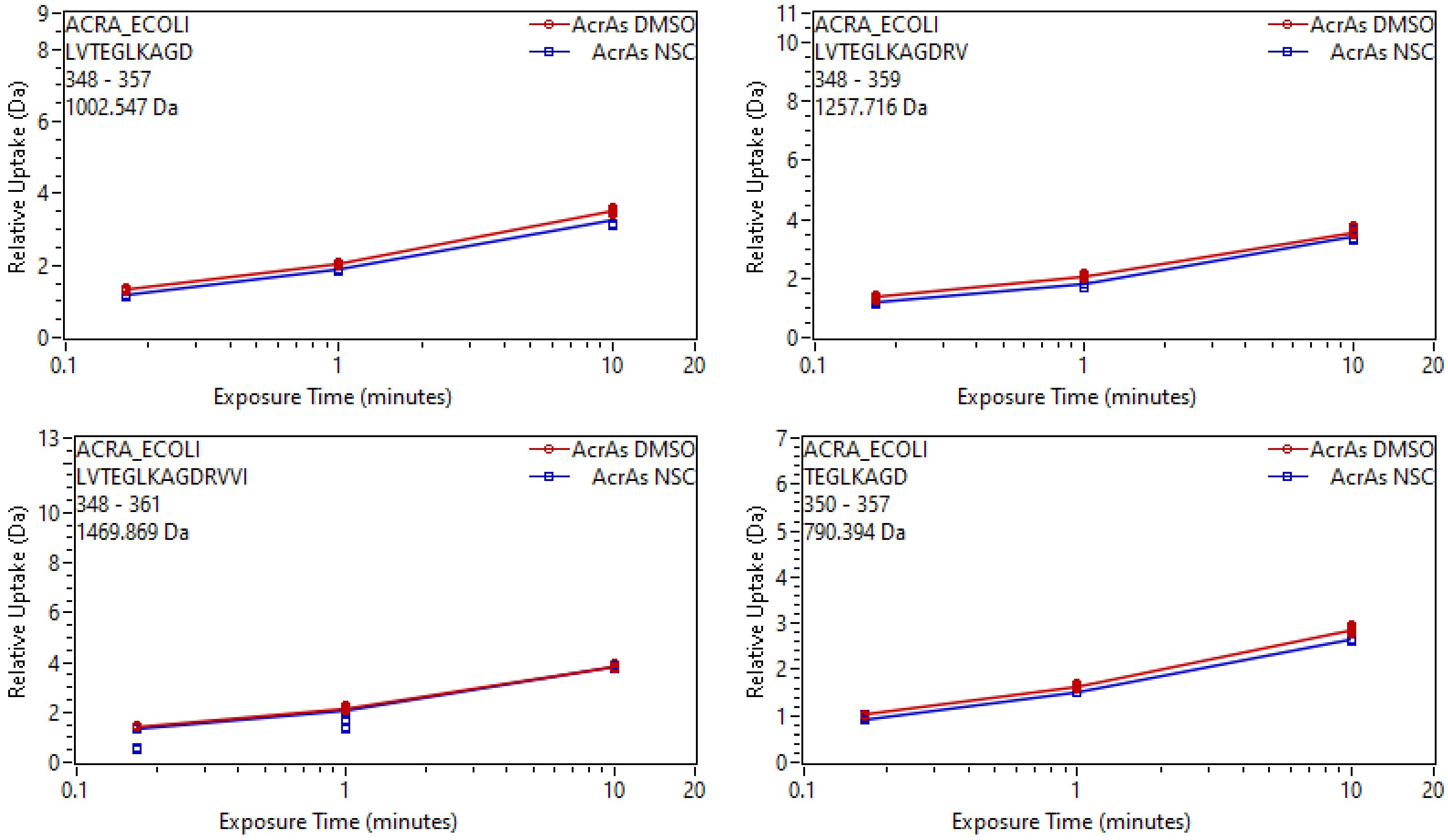
ACRA_ECOLI
LVTEGLKAGD
348 - 357
1002.547 Da
AcrAs DMSO
AcrAs NSC
Relative Uptake (Da)
Exposure Time (minutes)
ACRA_ECOLI
LVTEGLKAGDRV
348 - 359
1257.716 Da
AcrAs DMSO
AcrAs NSC
Relative Uptake (Da)
Exposure Time (minutes)
ACRA_ECOLI
LVTEGLKAGDRVVI
348 - 361
1469.869 Da
AcrAs DMSO
AcrAs NSC
Relative Uptake (Da)
Exposure Time (minutes)
ACRA_ECOLI
TEGLKAGD
350 - 357
790.394 Da
AcrAs DMSO
AcrAs NSC
Relative Uptake (Da)
Exposure Time (minutes)

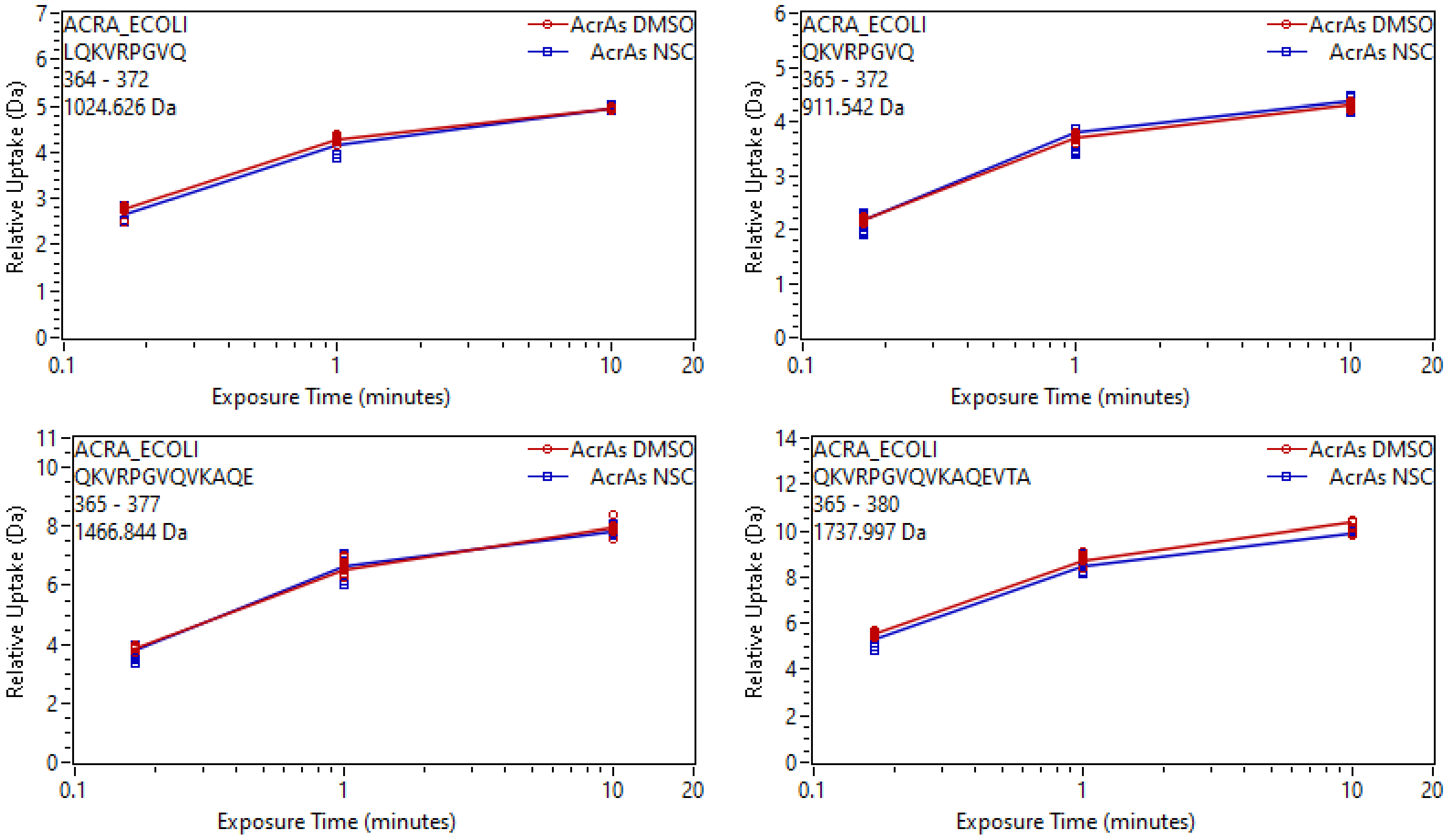
ACRA_ECOLI
LQKVRPGVQ
364 - 372
1024.626 Da
AcrAs DMSO
AcrAs NSC
Relative Uptake (Da)
Exposure Time (minutes)
ACRA_ECOLI
QKVRPGVQ
365 - 372
911.542 Da
AcrAs DMSO
AcrAs NSC
Relative Uptake (Da)
Exposure Time (minutes)
ACRA_ECOLI
QKVRPGVQVKAQE
365 - 377
1466.844 Da
AcrAs DMSO
AcrAs NSC
Relative Uptake (Da)
Exposure Time (minutes)
ACRA_ECOLI
QKVRPGVQVKAQEVTA
365 - 380
1737.997 Da
AcrAs DMSO
AcrAs NSC
Relative Uptake (Da)
Exposure Time (minutes)

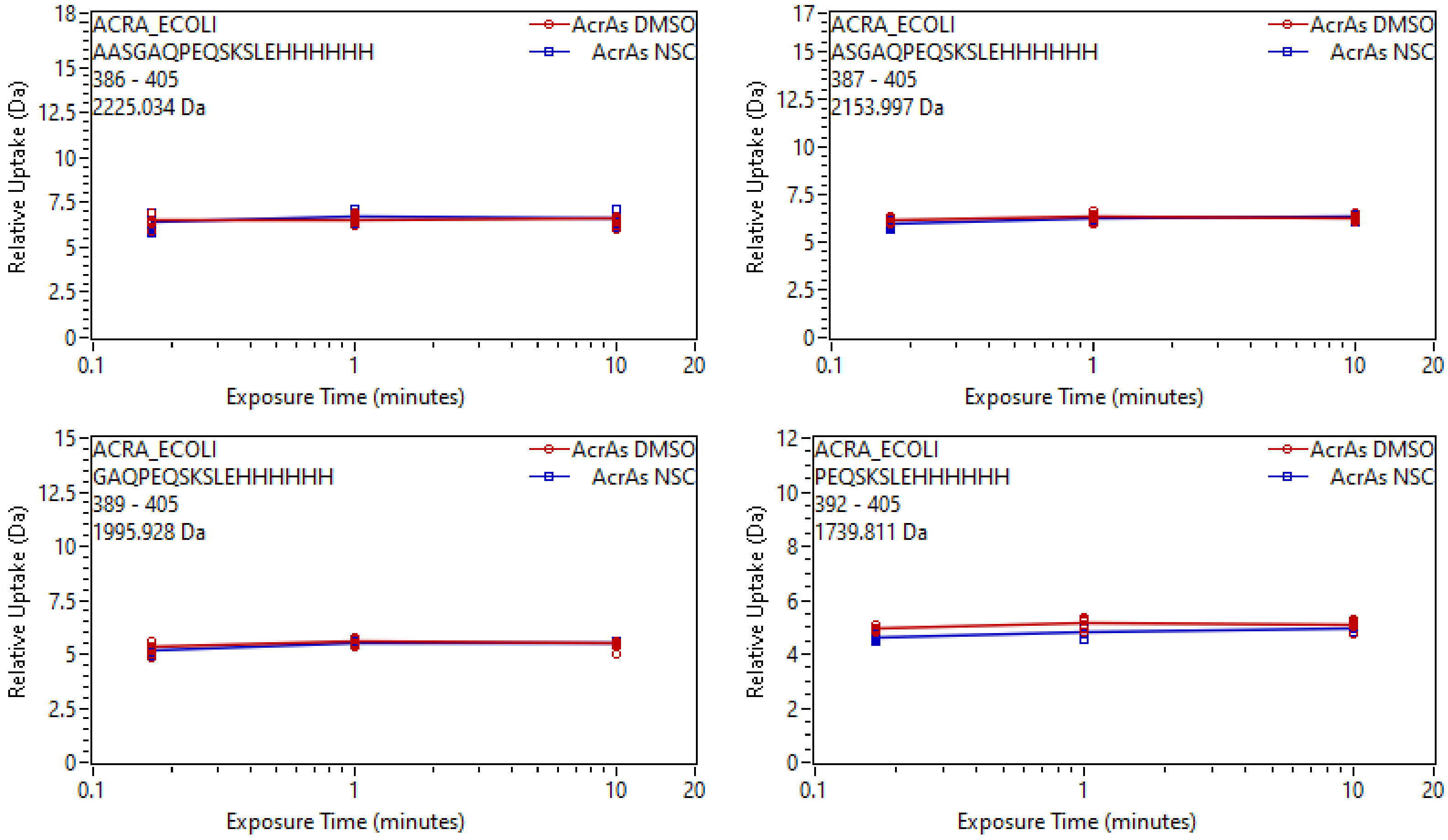
ACRA_ECOLI
AASGAQPEQSKSLEHHHHHH
386 - 405
2225.034 Da
AcrAs DMSO
AcrAs NSC
Relative Uptake (Da)
Exposure Time (minutes)
ACRA_ECOLI
ASGAQPEQSKSLEHHHHHH
387 - 405
2153.997 Da
ACRA_ECOLI
GAQPEQSKSLEHHHHHH
389 - 405
1995.928 Da
ACRA_ECOLI
PEQSKSLEHHHHHH
392 - 405
1739.811 Da
