## Supplementary material for "Conformational restriction shapes inhibition of a multidrug efflux adaptor protein": UptakePlots3

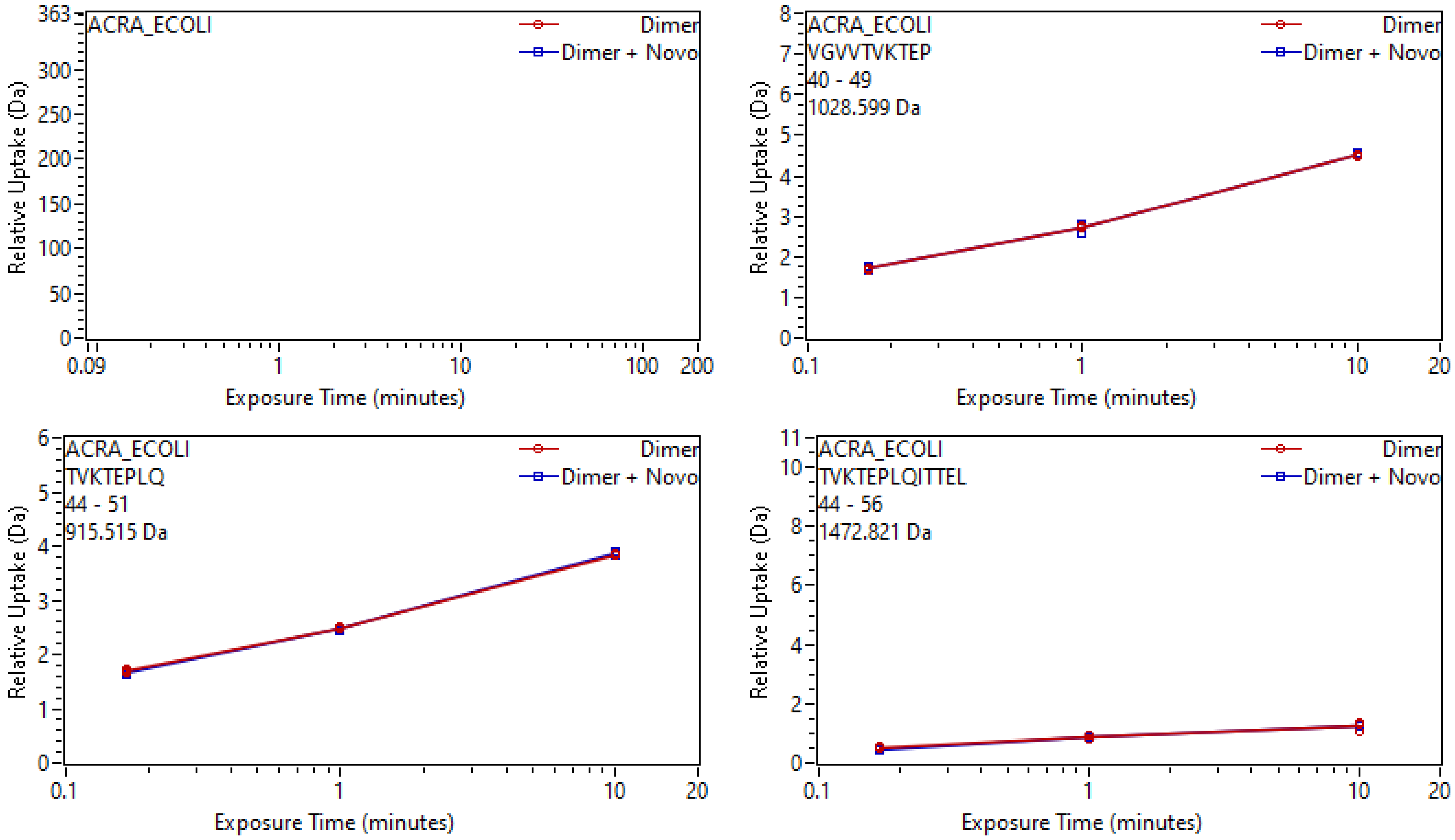
ACRA_ECOLI
Dimer
Dimer + Novo
Relative Uptake (Da)
Exposure Time (minutes)
363
300
250
200
150
100
50
0
0.09
1
10
100
200
ACRA_ECOLI
VGVVTVKTEP
40 - 49
1028.599 Da
Dimer
Dimer + Novo
Relative Uptake (Da)
Exposure Time (minutes)
ACRA_ECOLI
TVKTEPLQ
44 - 51
915.515 Da
Dimer
Dimer + Novo
Relative Uptake (Da)
Exposure Time (minutes)
ACRA_ECOLI
TVKTEPLQITTEL
44 - 56
1472.821 Da
Dimer
Dimer + Novo
Relative Uptake (Da)
Exposure Time (minutes)

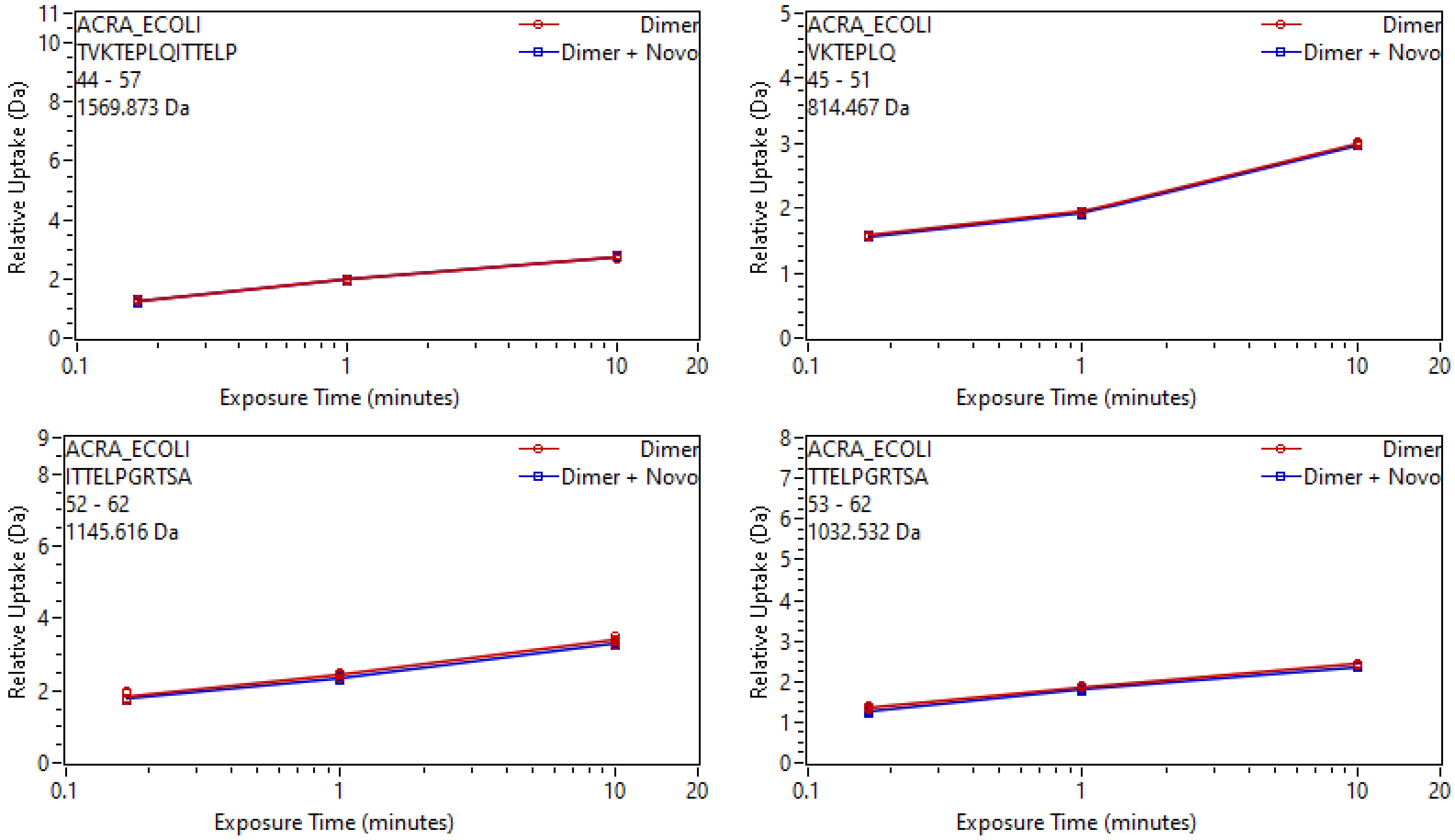

ACRA_ECOLI
TVKTEPLQITTELP
44 - 57
1569.873 Da
Dimer
Dimer + Novo
Relative Uptake (Da)
Exposure Time (minutes)
ACRA_ECOLI
VKTEPLQ
45 - 51
814.467 Da
Dimer
Dimer + Novo
Relative Uptake (Da)
Exposure Time (minutes)
ACRA_ECOLI
ITTELPGRTSA
52 - 62
1145.616 Da
Dimer
Dimer + Novo
Relative Uptake (Da)
Exposure Time (minutes)
ACRA_ECOLI
TTELPGRTSA
53 - 62
1032.532 Da
Dimer
Dimer + Novo
Relative Uptake (Da)
Exposure Time (minutes)

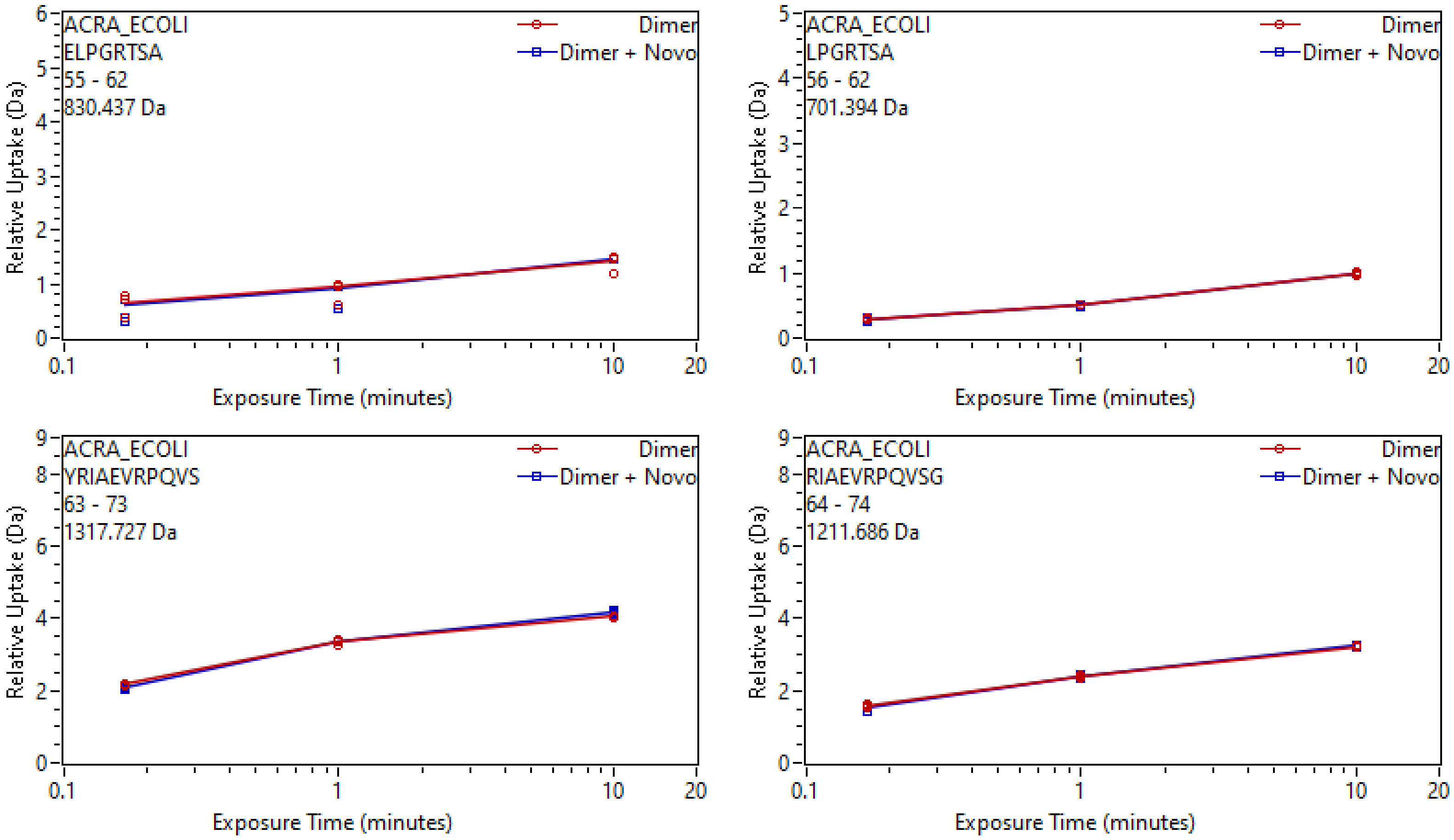
ACRA_ECOLI
ELPGRTSA
55 - 62
830.437 Da
Dimer
Dimer + Novo
Relative Uptake (Da)
Exposure Time (minutes)
ACRA_ECOLI
LPGRTSA
56 - 62
701.394 Da
Dimer
Dimer + Novo
Relative Uptake (Da)
Exposure Time (minutes)
ACRA_ECOLI
YRIAEVRPQVS
63 - 73
1317.727 Da
Dimer
Dimer + Novo
Relative Uptake (Da)
Exposure Time (minutes)
ACRA_ECOLI
RIAEVRPQVSG
64 - 74
1211.686 Da
Dimer
Dimer + Novo
Relative Uptake (Da)
Exposure Time (minutes)

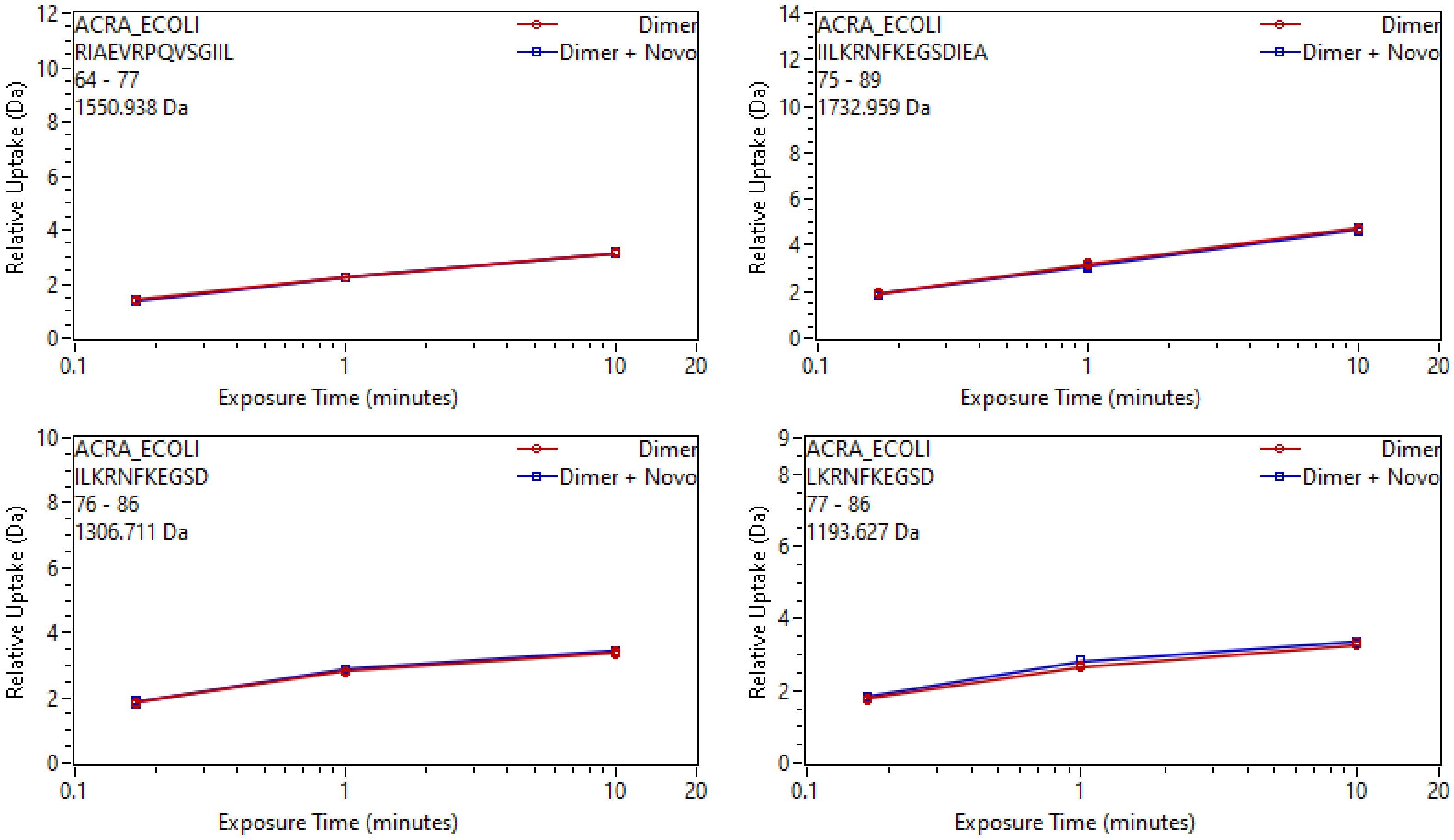
ACRA_ECOLI
RIAEVRPQVSGIIL
64 - 77
1550.938 Da
Dimer
Dimer + Novo
Relative Uptake (Da)
Exposure Time (minutes)
ACRA_ECOLI
IILKRNFKEGSDIEA
75 - 89
1732.959 Da
Dimer
Dimer + Novo
Relative Uptake (Da)
Exposure Time (minutes)
ACRA_ECOLI
ILKRNFKEGSD
76 - 86
1306.711 Da
Dimer
Dimer + Novo
Relative Uptake (Da)
Exposure Time (minutes)
ACRA_ECOLI
LKRNFKEGSD
77 - 86
1193.627 Da
Dimer
Dimer + Novo
Relative Uptake (Da)
Exposure Time (minutes)

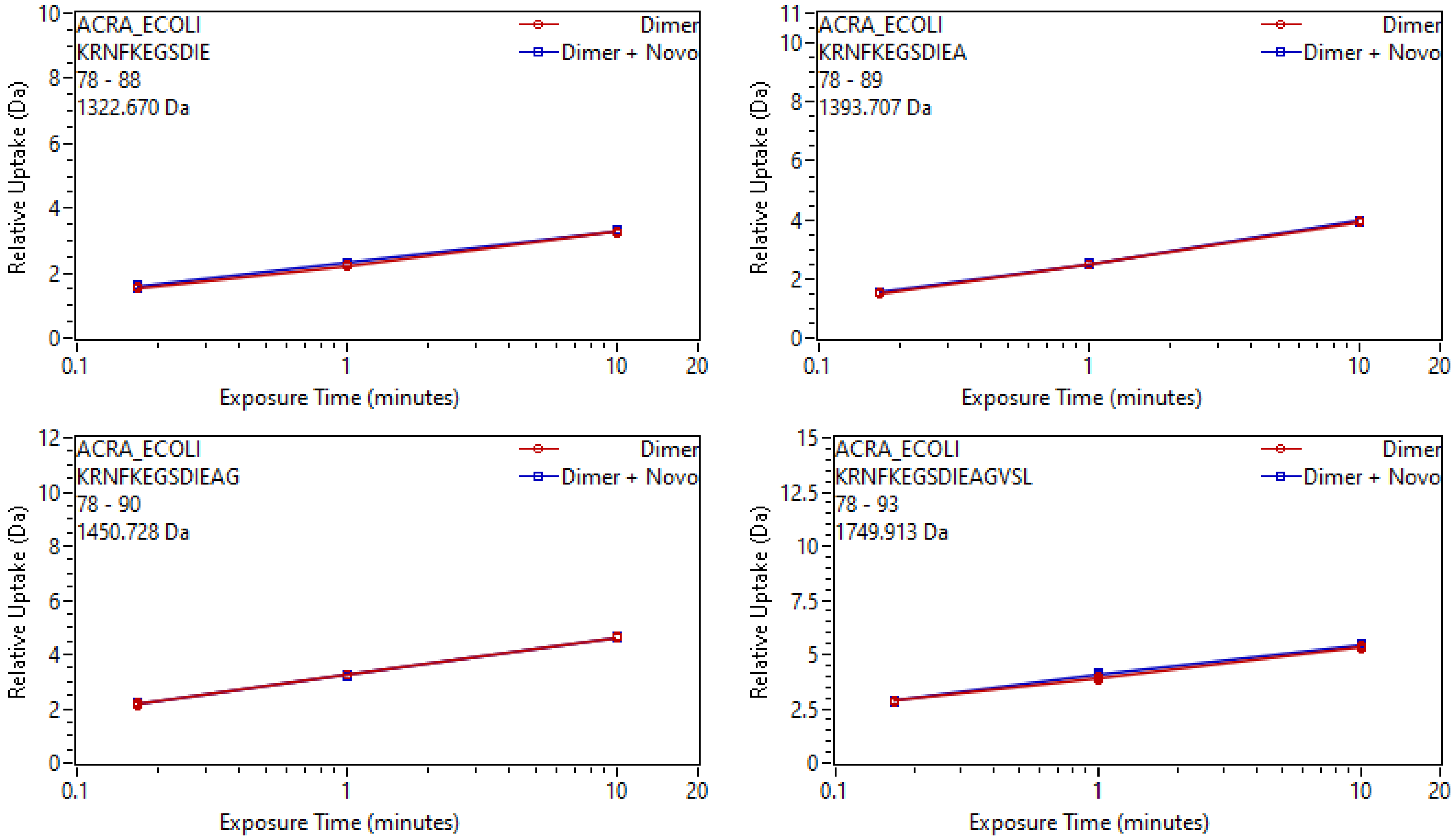
ACRA_ECOLI
KRNFKEGSDIE
78 - 88
1322.670 Da
Dimer
Dimer + Novo
Relative Uptake (Da)
Exposure Time (minutes)
ACRA_ECOLI
KRNFKEGSDIEA
78 - 89
1393.707 Da
Dimer
Dimer + Novo
Relative Uptake (Da)
Exposure Time (minutes)
ACRA_ECOLI
KRNFKEGSDIEAG
78 - 90
1450.728 Da
Dimer
Dimer + Novo
Relative Uptake (Da)
Exposure Time (minutes)
ACRA_ECOLI
KRNFKEGSDIEAGVSL
78 - 93
1749.913 Da
Dimer
Dimer + Novo
Relative Uptake (Da)
Exposure Time (minutes)

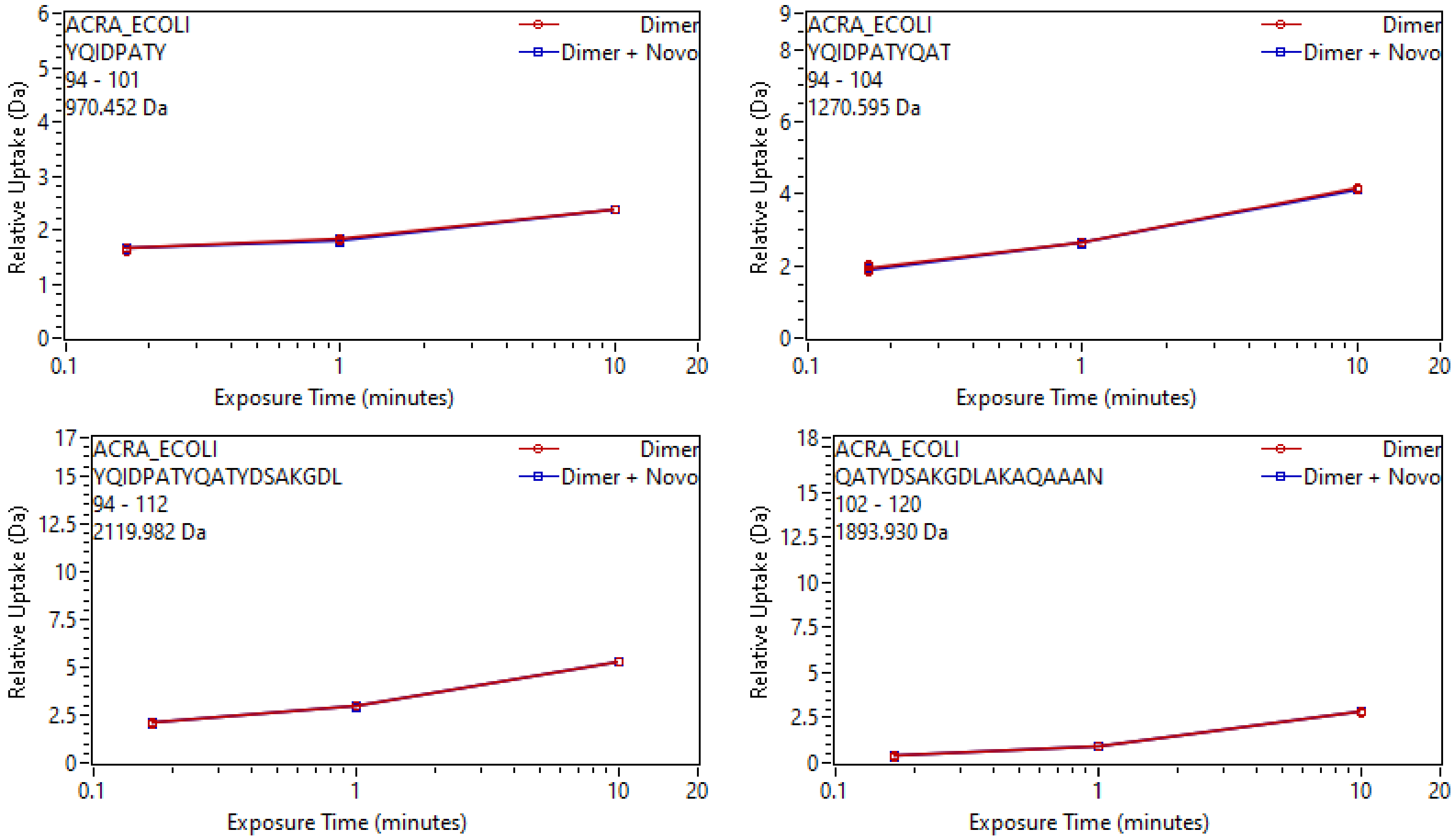
ACRA_ECOLI
YQIDPATY
94 - 101
970.452 Da
Dimer
Dimer + Novo
Relative Uptake (Da)
Exposure Time (minutes)
ACRA_ECOLI
YQIDPATYQAT
94 - 104
1270.595 Da
Dimer
Dimer + Novo
Relative Uptake (Da)
Exposure Time (minutes)
ACRA_ECOLI
YQIDPATYQATYDSAKGDL
94 - 112
2119.982 Da
Dimer
Dimer + Novo
Relative Uptake (Da)
Exposure Time (minutes)
ACRA_ECOLI
QATYDSAKGDLAKAQAAAN
102 - 120
1893.930 Da
Dimer
Dimer + Novo
Relative Uptake (Da)
Exposure Time (minutes)

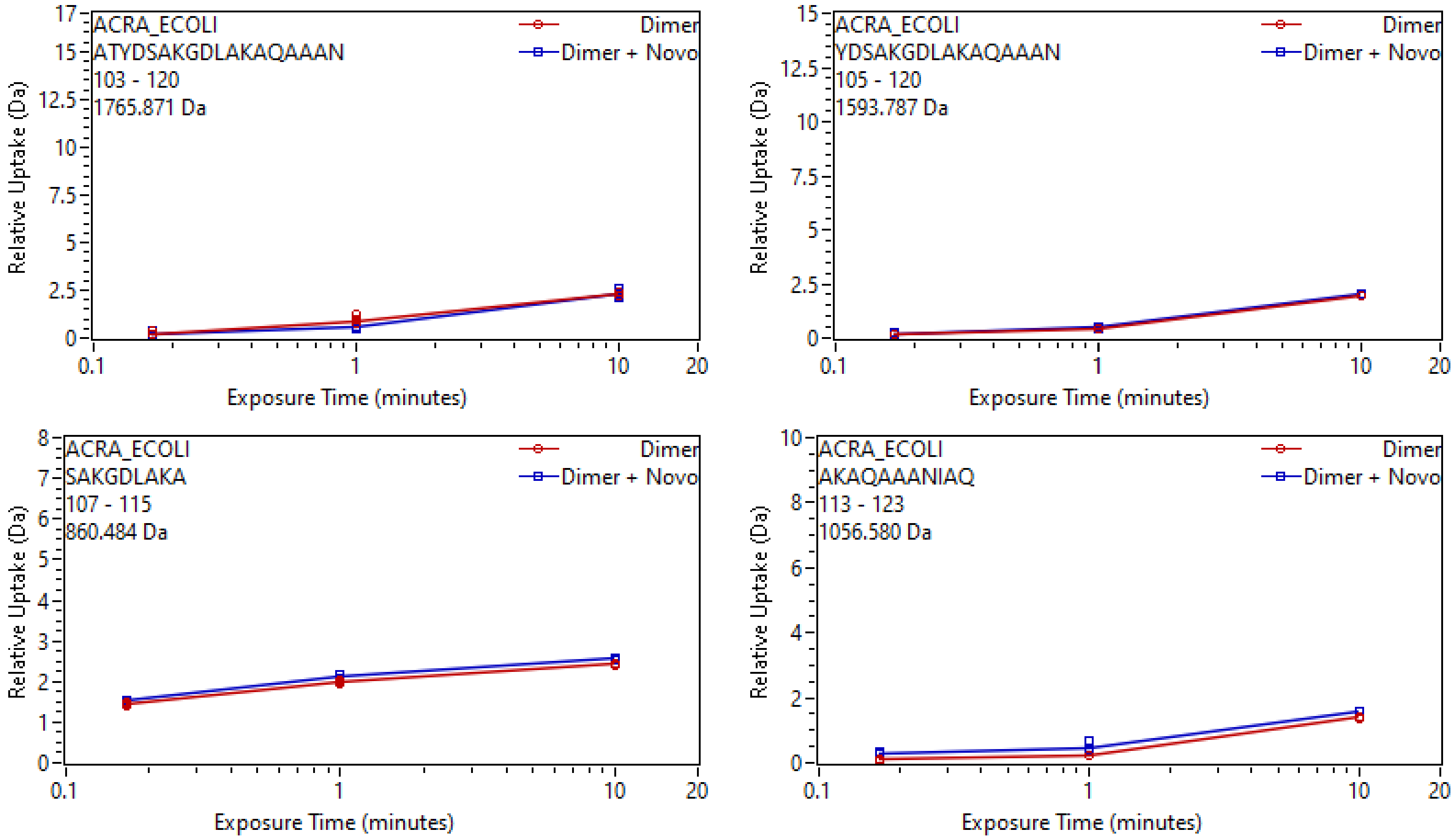
ACRA_ECOLI
ATYDSAKGDLAKAQAAAN
103 - 120
1765.871 Da
Dimer
Dimer + Novo
Relative Uptake (Da)
Exposure Time (minutes)
ACRA_ECOLI
YDSAKGDLAKAQAAAN
105 - 120
1593.787 Da
Dimer
Dimer + Novo
Relative Uptake (Da)
Exposure Time (minutes)
ACRA_ECOLI
SAKGDLAKA
107 - 115
860.484 Da
Dimer
Dimer + Novo
Relative Uptake (Da)
Exposure Time (minutes)
ACRA_ECOLI
AKAQAAANIAQ
113 - 123
1056.580 Da
Dimer
Dimer + Novo
Relative Uptake (Da)
Exposure Time (minutes)

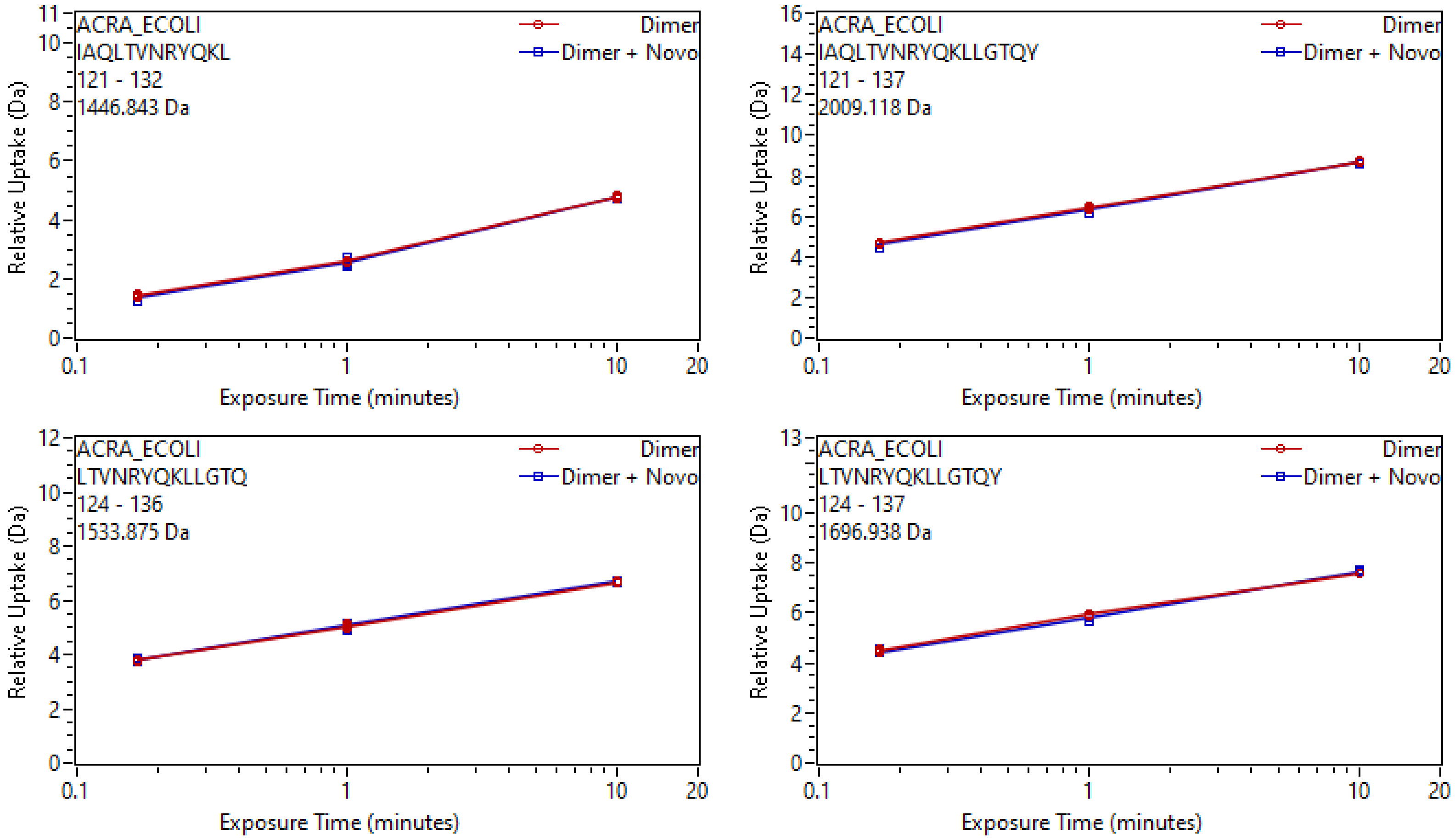
ACRA_ECOLI
IAQLTVNRYQKL
121 - 132
1446.843 Da
Dimer
Dimer + Novo
Relative Uptake (Da)
Exposure Time (minutes)
ACRA_ECOLI
IAQLTVNRYQKLLGTQY
121 - 137
2009.118 Da
Dimer
Dimer + Novo
Relative Uptake (Da)
Exposure Time (minutes)
ACRA_ECOLI
LTVNRYQKLLGTQ
124 - 136
1533.875 Da
Dimer
Dimer + Novo
Relative Uptake (Da)
Exposure Time (minutes)
ACRA_ECOLI
LTVNRYQKLLGTQY
124 - 137
1696.938 Da
Dimer
Dimer + Novo
Relative Uptake (Da)
Exposure Time (minutes)

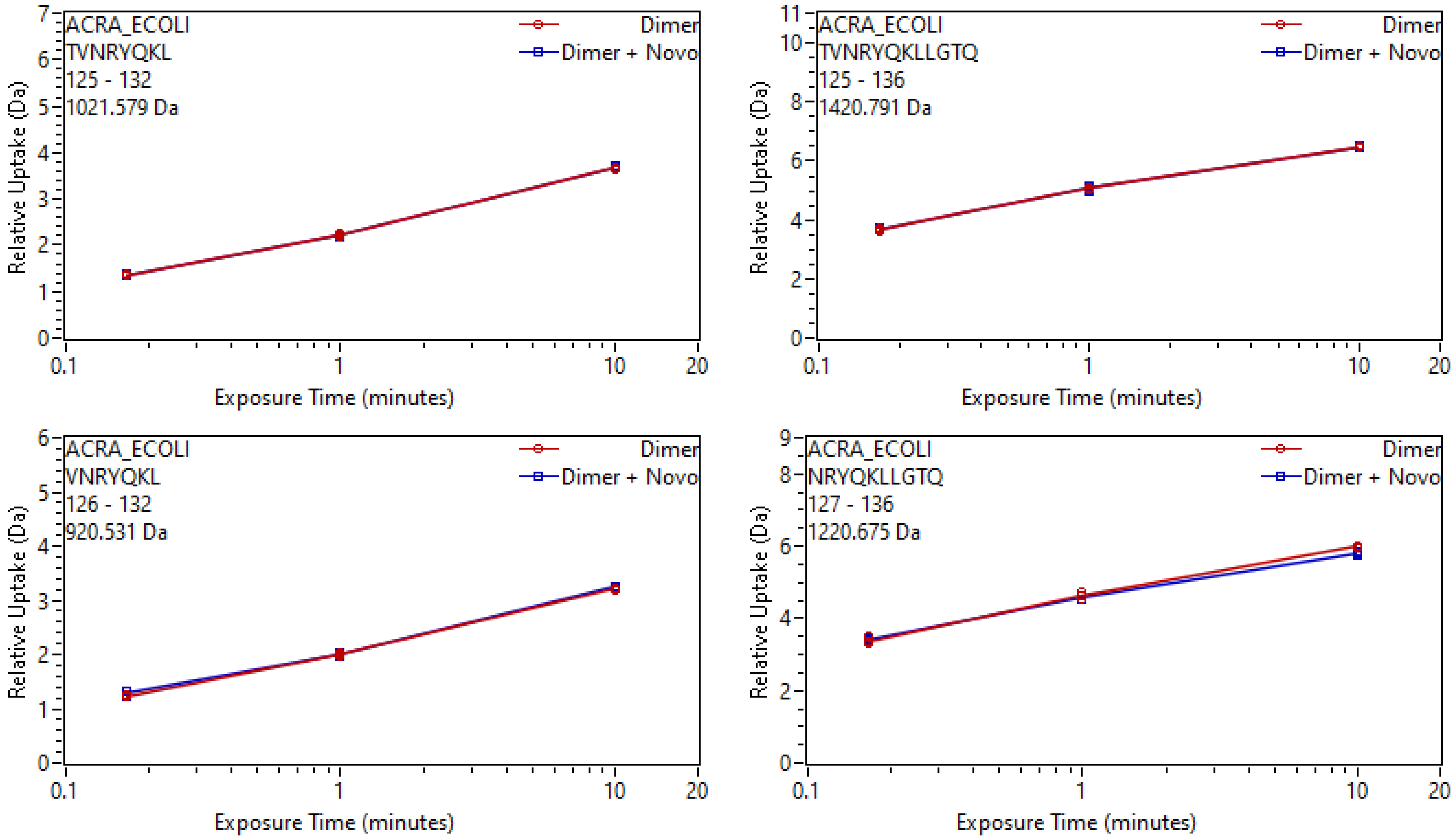

ACRA_ECOLI
TVNRYQKL
125 - 132
1021.579 Da
Dimer
Dimer + Novo
Relative Uptake (Da)
Exposure Time (minutes)
ACRA_ECOLI
TVNRYQKLLGTQ
125 - 136
1420.791 Da
Dimer
Dimer + Novo
Relative Uptake (Da)
Exposure Time (minutes)
ACRA_ECOLI
VNRYQKL
126 - 132
920.531 Da
Dimer
Dimer + Novo
Relative Uptake (Da)
Exposure Time (minutes)
ACRA_ECOLI
NRYQKLLGTQ
127 - 136
1220.675 Da
Dimer
Dimer + Novo
Relative Uptake (Da)
Exposure Time (minutes)

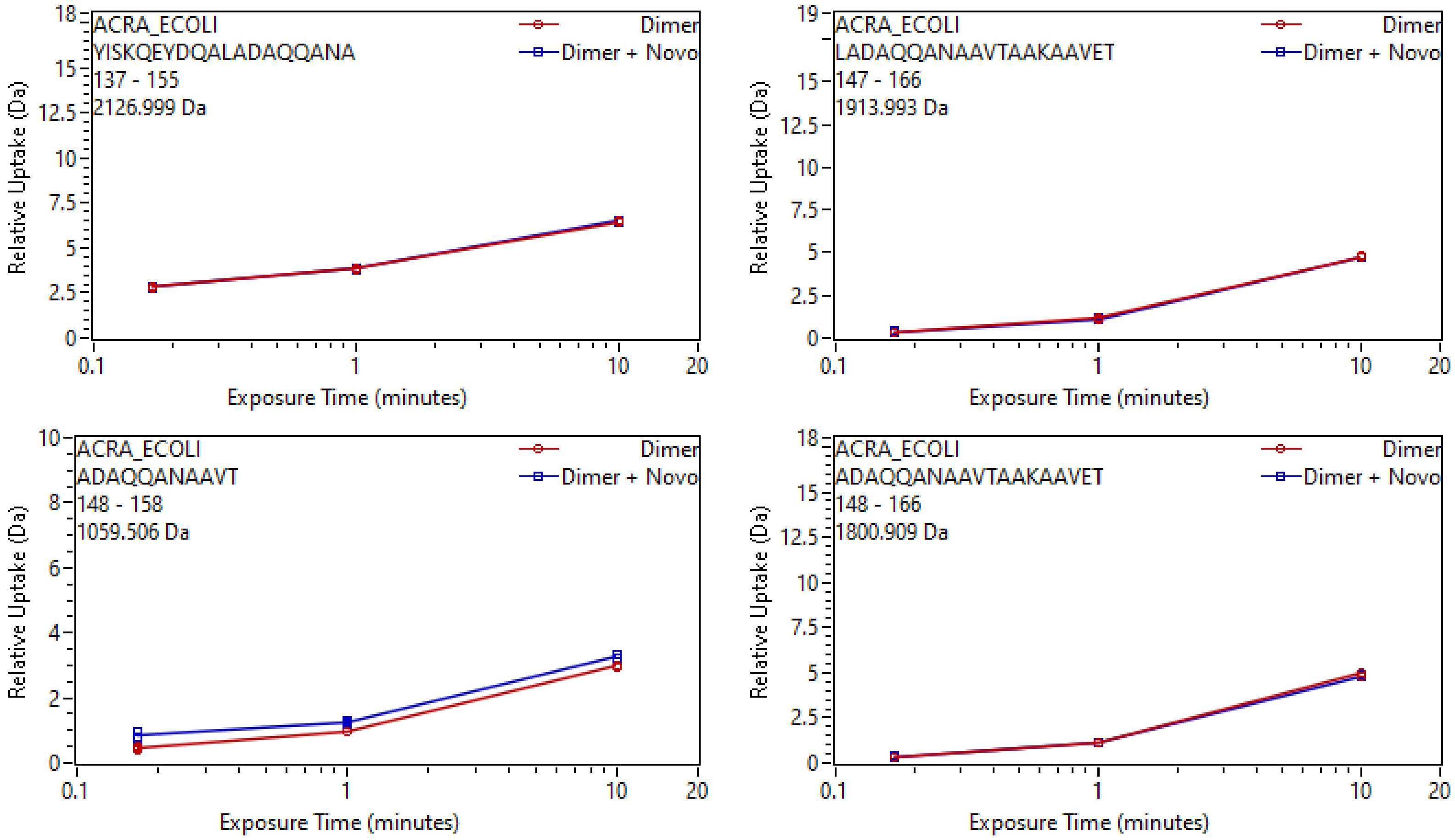
ACRA_ECOLI
YISKQEYDQALADAQQANA
137 - 155
2126.999 Da
Dimer
Dimer + Novo
Relative Uptake (Da)
Exposure Time (minutes)
ACRA_ECOLI
LADAQQANAAVTAAKAAVET
147 - 166
1913.993 Da
Dimer
Dimer + Novo
Relative Uptake (Da)
Exposure Time (minutes)
ACRA_ECOLI
ADAQQANAAVT
148 - 158
1059.506 Da
Dimer
Dimer + Novo
Relative Uptake (Da)
Exposure Time (minutes)
ACRA_ECOLI
ADAQQANAAVTAAKAAVET
148 - 166
1800.909 Da
Dimer
Dimer + Novo
Relative Uptake (Da)
Exposure Time (minutes)

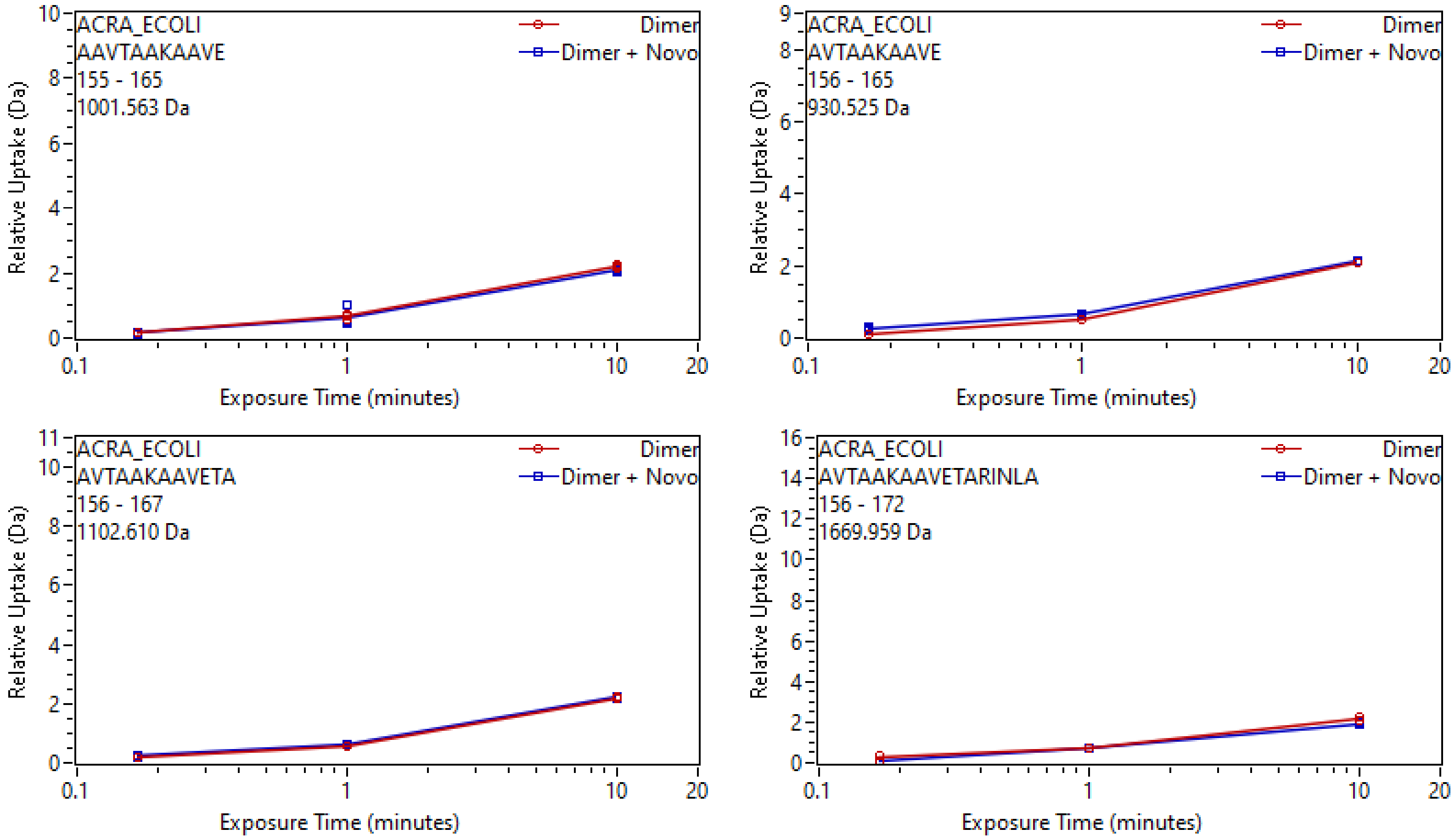

ACRA_ECOLI
AAVTAAKAAVE
155 - 165
1001.563 Da
Dimer
Dimer + Novo
Relative Uptake (Da)
Exposure Time (minutes)
ACRA_ECOLI
AVTAAKAAVE
156 - 165
930.525 Da
Dimer
Dimer + Novo
Relative Uptake (Da)
Exposure Time (minutes)
ACRA_ECOLI
AVTAAKAAVETA
156 - 167
1102.610 Da
Dimer
Dimer + Novo
Relative Uptake (Da)
Exposure Time (minutes)
ACRA_ECOLI
AVTAAKAAVETARINLA
156 - 172
1669.959 Da
Dimer
Dimer + Novo
Relative Uptake (Da)
Exposure Time (minutes)

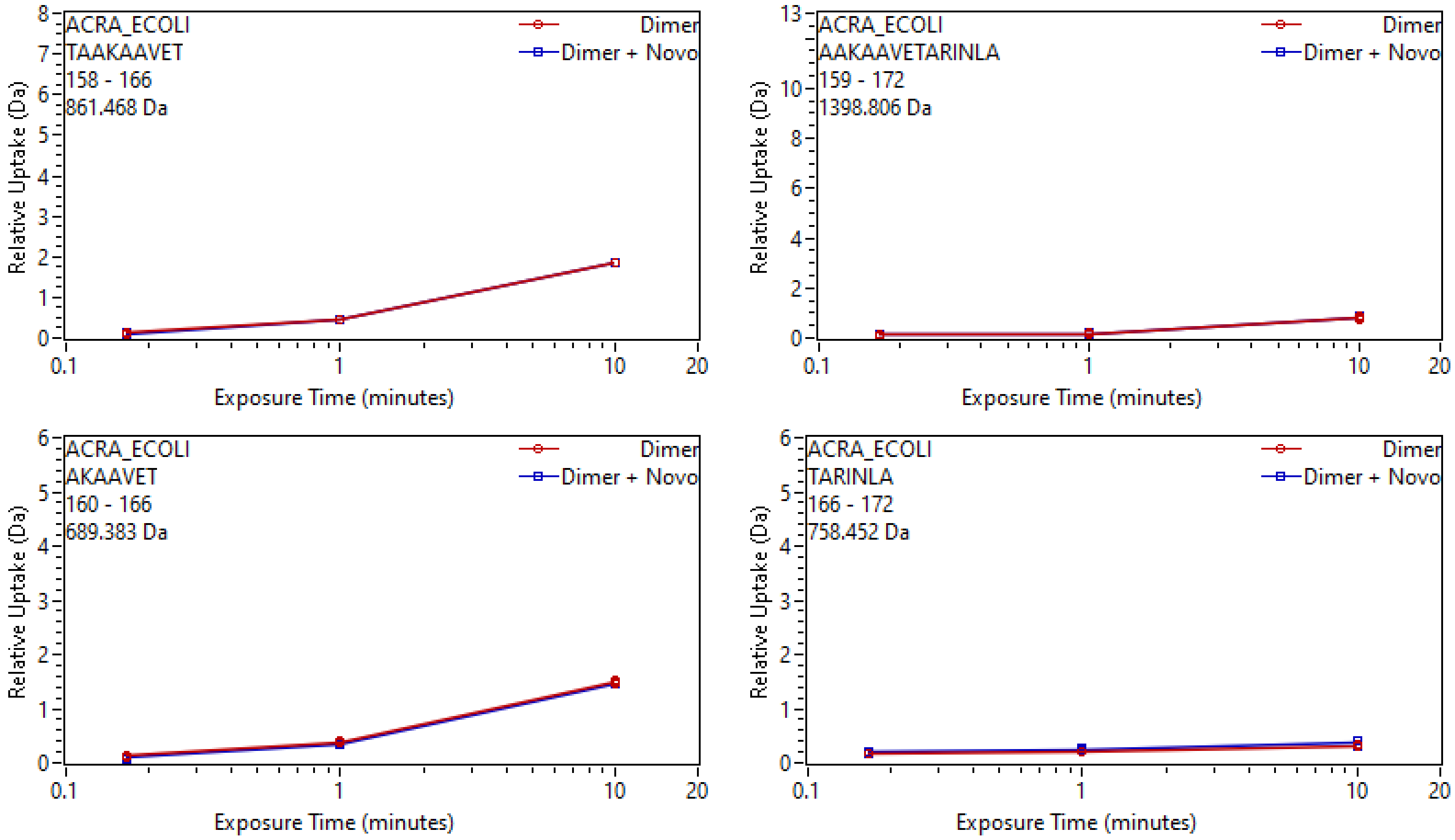
ACRA_ECOLI
TAAKAAVET
158 - 166
861.468 Da
Dimer
Dimer + Novo
Relative Uptake (Da)
Exposure Time (minutes)
ACRA_ECOLI
AAKAAVETARINLA
159 - 172
1398.806 Da
Dimer
Dimer + Novo
Relative Uptake (Da)
Exposure Time (minutes)
ACRA_ECOLI
AKAAVET
160 - 166
689.383 Da
Dimer
Dimer + Novo
Relative Uptake (Da)
Exposure Time (minutes)
ACRA_ECOLI
TARINLA
166 - 172
758.452 Da
Dimer
Dimer + Novo
Relative Uptake (Da)
Exposure Time (minutes)

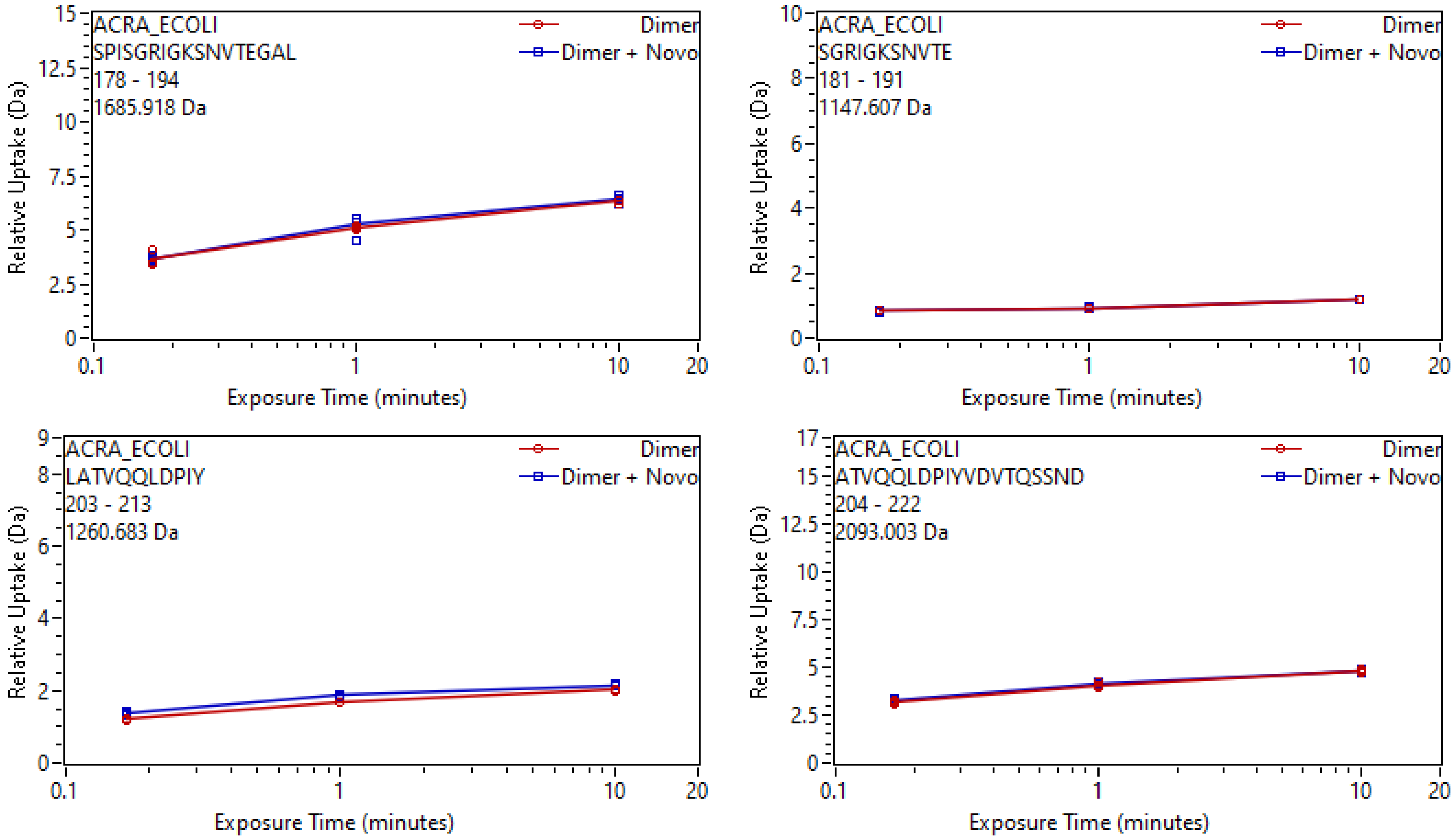
ACRA_ECOLI
SPISGRIGKSNVTEGAL
178 - 194
1685.918 Da
Dimer
Dimer + Novo
Relative Uptake (Da)
Exposure Time (minutes)
ACRA_ECOLI
SGRIGKSNVTE
181 - 191
1147.607 Da
Dimer
Dimer + Novo
Relative Uptake (Da)
Exposure Time (minutes)
ACRA_ECOLI
LATVQQLDPIY
203 - 213
1260.683 Da
Dimer
Dimer + Novo
Relative Uptake (Da)
Exposure Time (minutes)
ACRA_ECOLI
ATVQQLDPIYVDVTQSSND
204 - 222
2093.003 Da
Dimer
Dimer + Novo
Relative Uptake (Da)
Exposure Time (minutes)

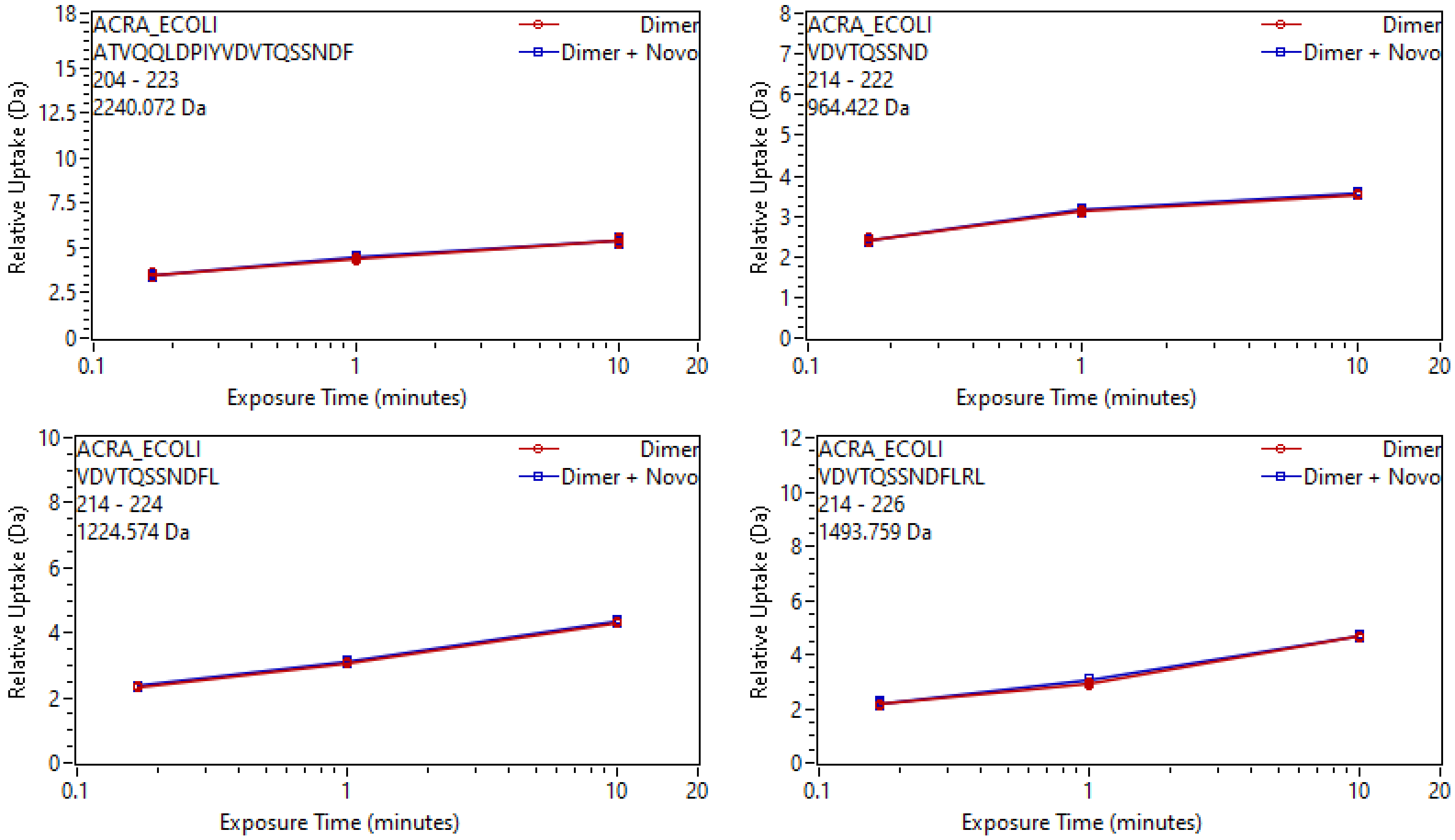

ACRA_ECOLI
ATVQQLDPIYVDVTQSSNDF
204 - 223
2240.072 Da
Dimer
Dimer + Novo
Relative Uptake (Da)
Exposure Time (minutes)
ACRA_ECOLI
VDVTQSSND
214 - 222
964.422 Da
Dimer
Dimer + Novo
Relative Uptake (Da)
Exposure Time (minutes)
ACRA_ECOLI
VDVTQSSNDFL
214 - 224
1224.574 Da
Dimer
Dimer + Novo
Relative Uptake (Da)
Exposure Time (minutes)
ACRA_ECOLI
VDVTQSSNDFLRL
214 - 226
1493.759 Da
Dimer
Dimer + Novo
Relative Uptake (Da)
Exposure Time (minutes)

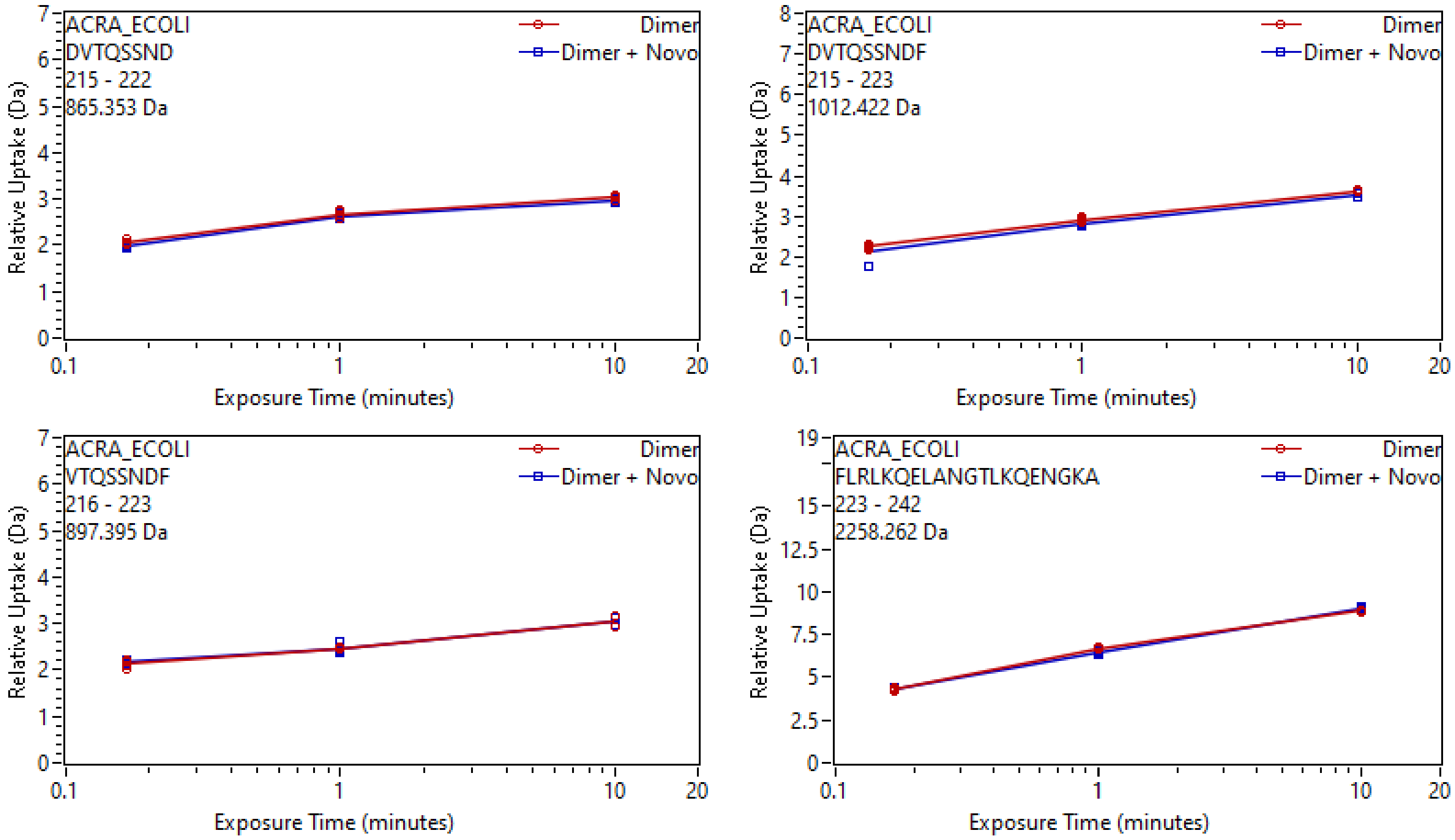

ACRA_ECOLI
DVTQSSND
215 - 222
865.353 Da
Dimer
Dimer + Novo
Relative Uptake (Da)
Exposure Time (minutes)
ACRA_ECOLI
DVTQSSNDF
215 - 223
1012.422 Da
Dimer
Dimer + Novo
Relative Uptake (Da)
Exposure Time (minutes)
ACRA_ECOLI
VTQSSNDF
216 - 223
897.395 Da
Dimer
Dimer + Novo
Relative Uptake (Da)
Exposure Time (minutes)
ACRA_ECOLI
FLRLKQELANGTLKQENGKA
223 - 242
2258.262 Da
Dimer
Dimer + Novo
Relative Uptake (Da)
Exposure Time (minutes)

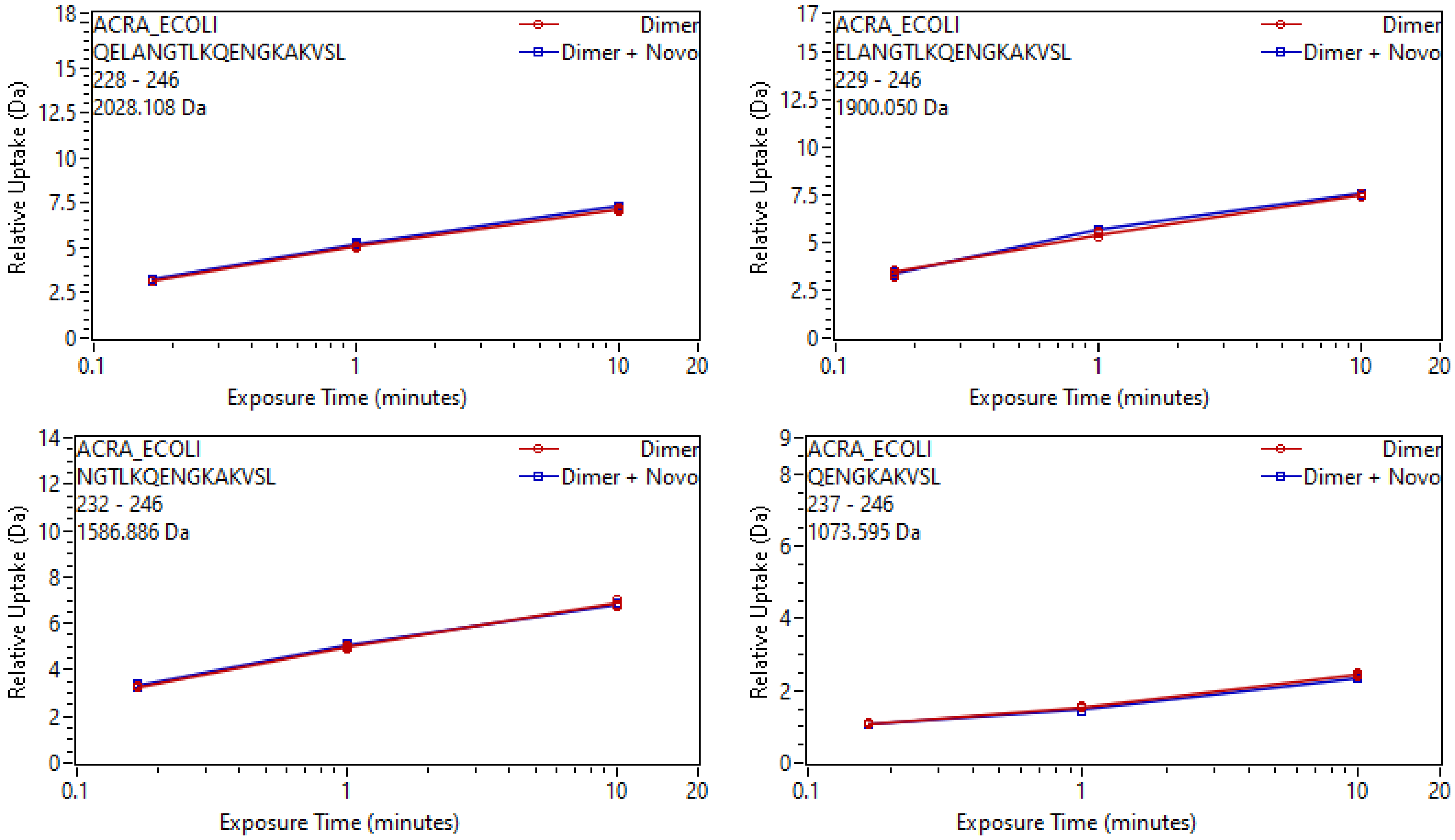
ACRA_ECOLI
QELANGTLKQENGKAKVSL
228 - 246
2028.108 Da
Dimer
Dimer + Novo
Relative Uptake (Da)
Exposure Time (minutes)
ACRA_ECOLI
ELANGTLKQENGKAKVSL
229 - 246
1900.050 Da
Dimer
Dimer + Novo
Relative Uptake (Da)
Exposure Time (minutes)
ACRA_ECOLI
NGTLKQENGKAKVSL
232 - 246
1586.886 Da
Dimer
Dimer + Novo
Relative Uptake (Da)
Exposure Time (minutes)
ACRA_ECOLI
QENGKAKVSL
237 - 246
1073.595 Da
Dimer
Dimer + Novo
Relative Uptake (Da)
Exposure Time (minutes)

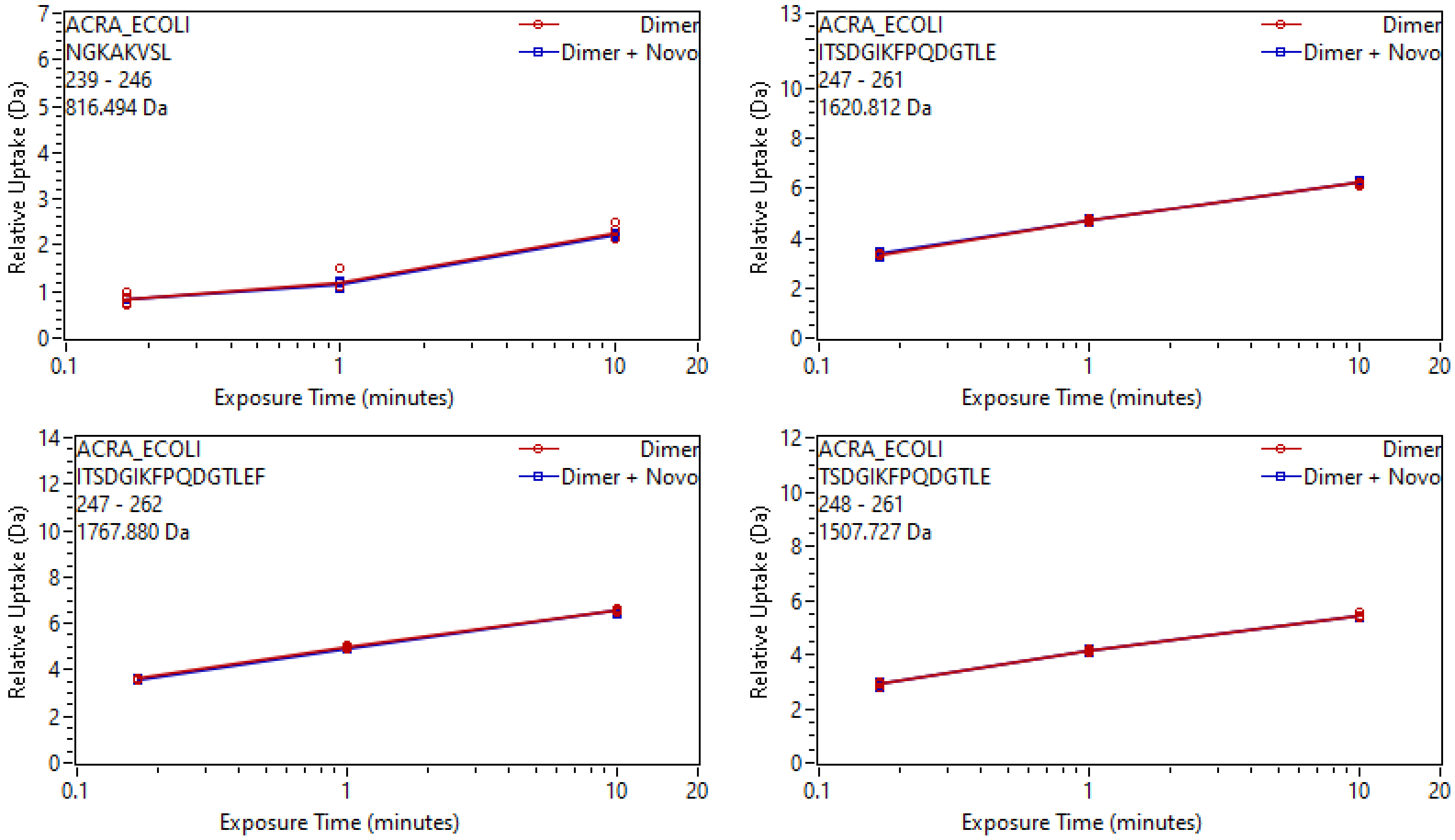
ACRA_ECOLI
NGKAKVSL
239 - 246
816.494 Da
Dimer
Dimer + Novo
Relative Uptake (Da)
Exposure Time (minutes)
ACRA_ECOLI
ITSDGIKFPQDGTLE
247 - 261
1620.812 Da
ACRA_ECOLI
ITSDGIKFPQDGTLEF
247 - 262
1767.880 Da
ACRA_ECOLI
TSDGIKFPQDGTLE
248 - 261
1507.727 Da

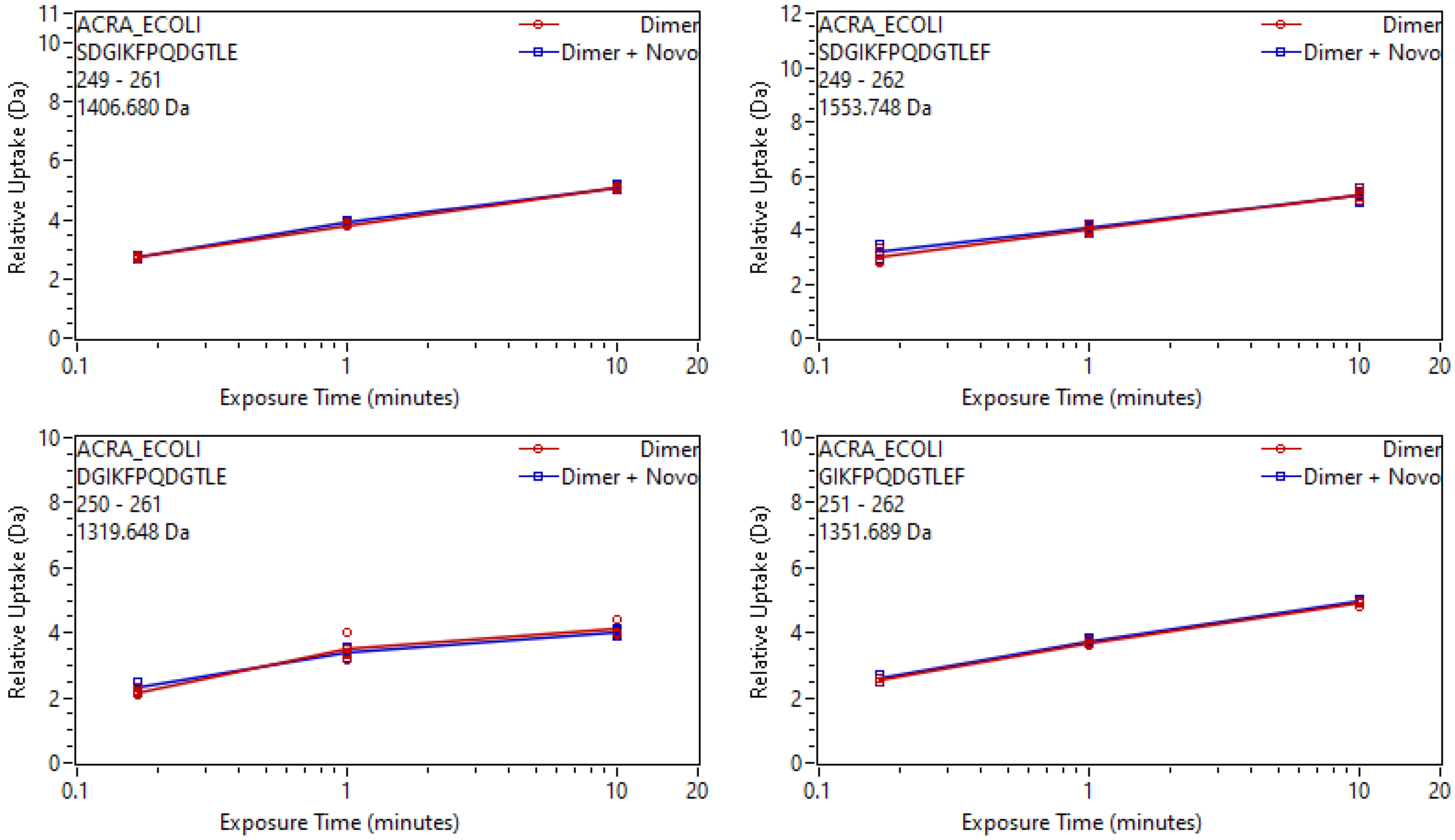
ACRA_ECOLI
SDGIKFPQDGTLE
249 - 261
1406.680 Da
Dimer
Dimer + Novo
Relative Uptake (Da)
Exposure Time (minutes)
ACRA_ECOLI
SDGIKFPQDGTLEF
249 - 262
1553.748 Da
Dimer
Dimer + Novo
Relative Uptake (Da)
Exposure Time (minutes)
ACRA_ECOLI
DGIKFPQDGTLE
250 - 261
1319.648 Da
Dimer
Dimer + Novo
Relative Uptake (Da)
Exposure Time (minutes)
ACRA_ECOLI
GIKFPQDGTLEF
251 - 262
1351.689 Da
Dimer
Dimer + Novo
Relative Uptake (Da)
Exposure Time (minutes)

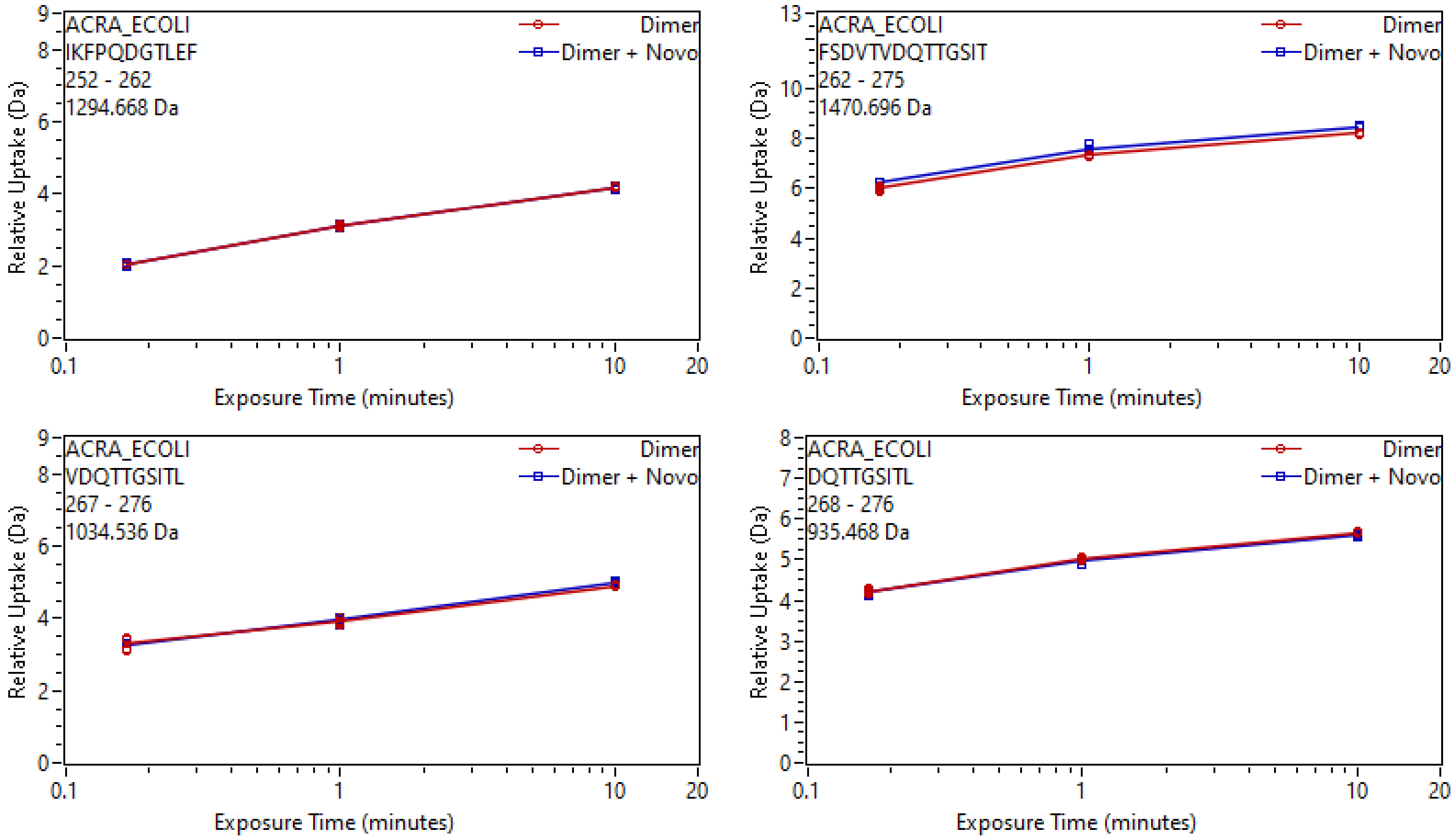

ACRA_ECOLI
IKFPQDGTLEF
252 - 262
1294.668 Da
Dimer
Dimer + Novo
Relative Uptake (Da)
Exposure Time (minutes)
ACRA_ECOLI
FSDVTVDQTTGSIT
262 - 275
1470.696 Da
Dimer
Dimer + Novo
Relative Uptake (Da)
Exposure Time (minutes)
ACRA_ECOLI
VDQTTGSITL
267 - 276
1034.536 Da
Dimer
Dimer + Novo
Relative Uptake (Da)
Exposure Time (minutes)
ACRA_ECOLI
DQTTGSITL
268 - 276
935.468 Da
Dimer
Dimer + Novo
Relative Uptake (Da)
Exposure Time (minutes)

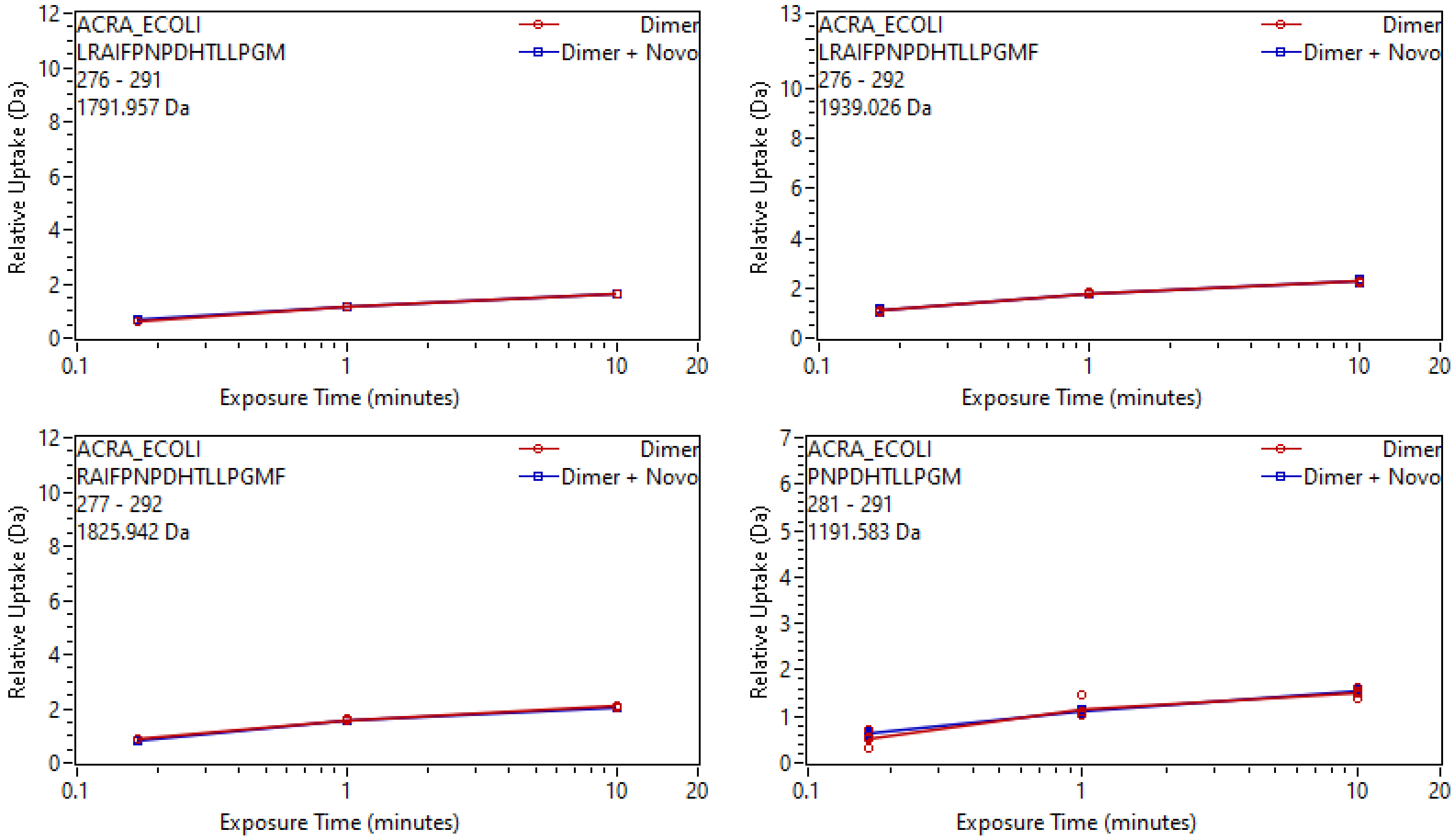
ACRA_ECOLI
LRAIFPNPDHTLLPGM
276 - 291
1791.957 Da
Dimer
Dimer + Novo
Relative Uptake (Da)
Exposure Time (minutes)
ACRA_ECOLI
LRAIFPNPDHTLLPGMF
276 - 292
1939.026 Da
Dimer
Dimer + Novo
Relative Uptake (Da)
Exposure Time (minutes)
ACRA_ECOLI
RAIFPNPDHTLLPGMF
277 - 292
1825.942 Da
Dimer
Dimer + Novo
Relative Uptake (Da)
Exposure Time (minutes)
ACRA_ECOLI
PNPDHTLLPGM
281 - 291
1191.583 Da
Dimer
Dimer + Novo
Relative Uptake (Da)
Exposure Time (minutes)

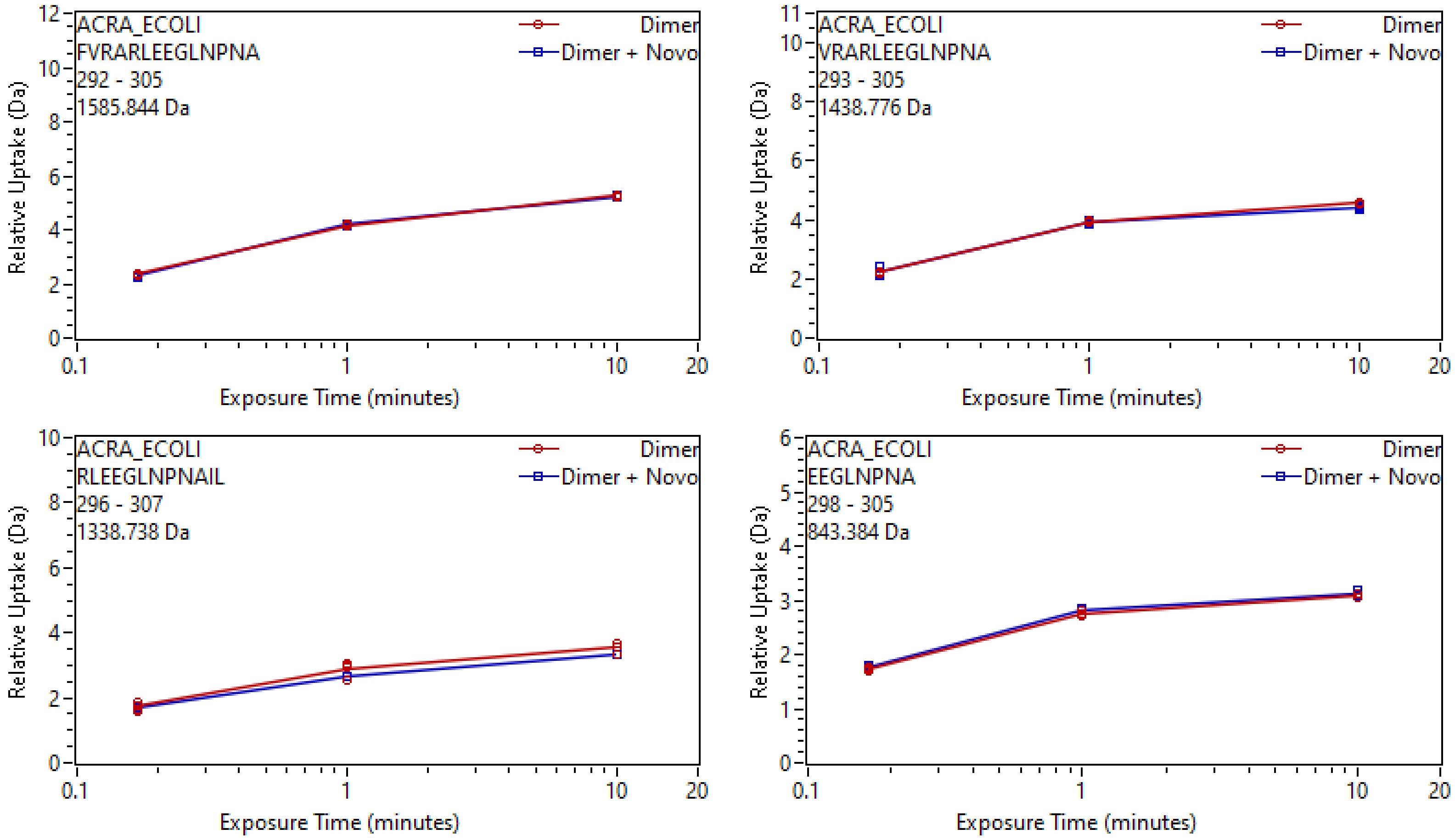

ACRA_ECOLI
FVRARLEEGLNPNA
292 - 305
1585.844 Da
Dimer
Dimer + Novo
Relative Uptake (Da)
Exposure Time (minutes)
ACRA_ECOLI
VRARLEEGLNPNA
293 - 305
1438.776 Da
Dimer
Dimer + Novo
Relative Uptake (Da)
Exposure Time (minutes)
ACRA_ECOLI
RLEEGLNPNAIL
296 - 307
1338.738 Da
Dimer
Dimer + Novo
Relative Uptake (Da)
Exposure Time (minutes)
ACRA_ECOLI
EEGLNPNA
298 - 305
843.384 Da
Dimer
Dimer + Novo
Relative Uptake (Da)
Exposure Time (minutes)

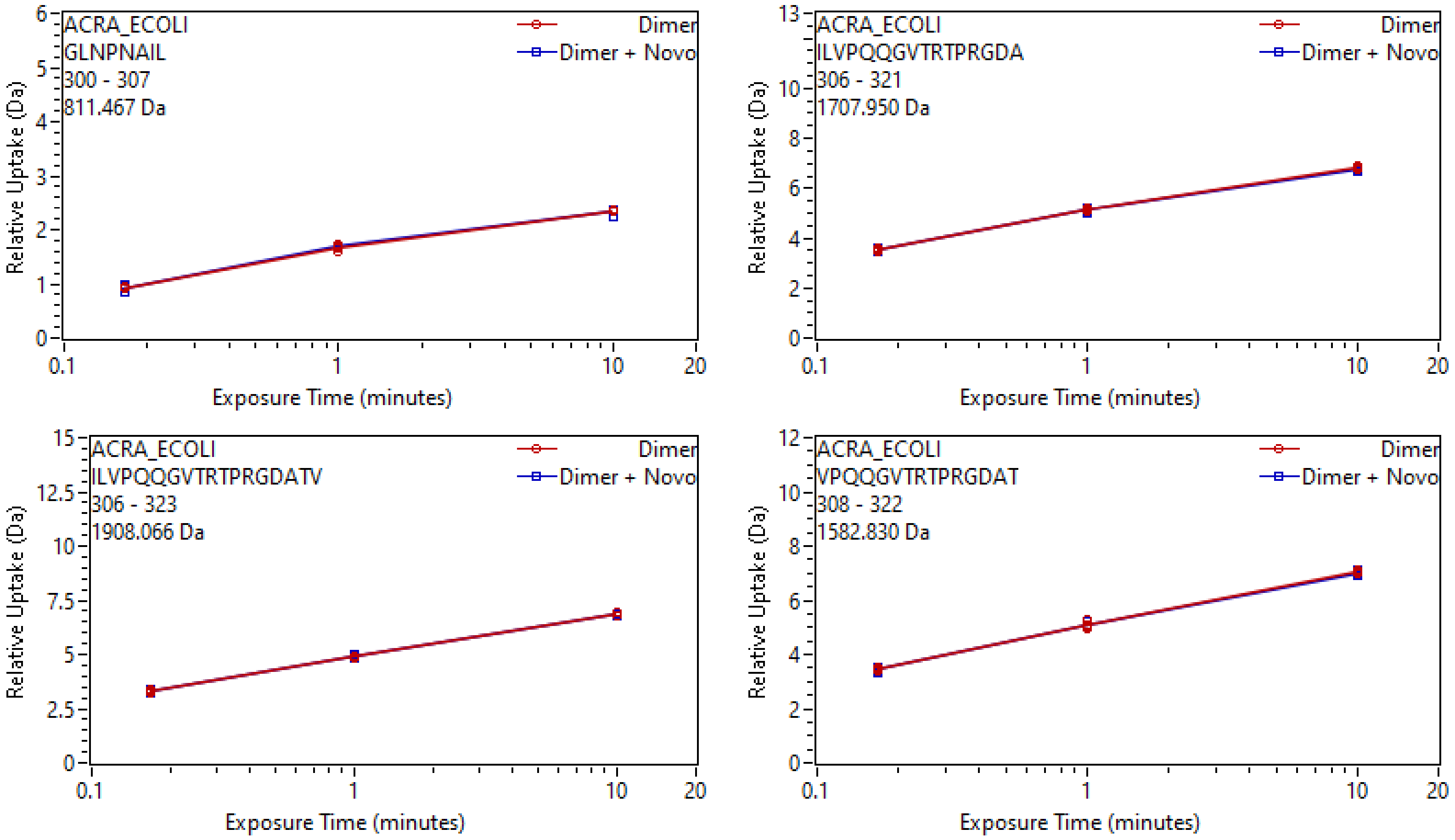
ACRA_ECOLI
GLNPNAIL
300 - 307
811.467 Da
Dimer
Dimer + Novo
Relative Uptake (Da)
Exposure Time (minutes)
ACRA_ECOLI
ILVPQQGVTRTPRGDA
306 - 321
1707.950 Da
Dimer
Dimer + Novo
Relative Uptake (Da)
Exposure Time (minutes)
ACRA_ECOLI
ILVPQQGVTRTPRGDATV
306 - 323
1908.066 Da
Dimer
Dimer + Novo
Relative Uptake (Da)
Exposure Time (minutes)
ACRA_ECOLI
VPQQGVTRTPRGDAT
308 - 322
1582.830 Da
Dimer
Dimer + Novo
Relative Uptake (Da)
Exposure Time (minutes)

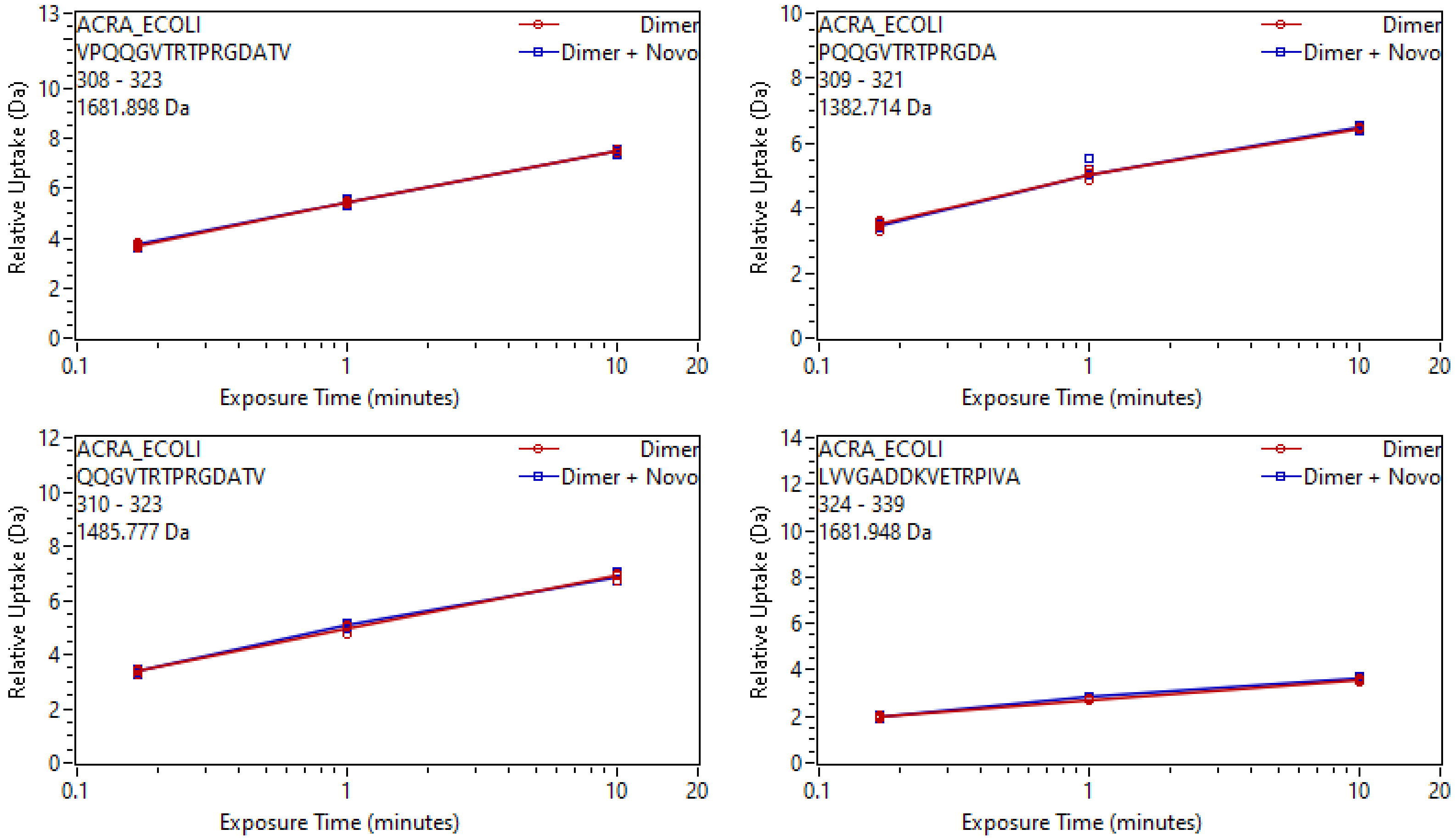
ACRA_ECOLI
VPQQGVTRTPRGDATV
308 - 323
1681.898 Da
Dimer
Dimer + Novo
Relative Uptake (Da)
Exposure Time (minutes)
ACRA_ECOLI
PQQGVTRTPRGDA
309 - 321
1382.714 Da
Dimer
Dimer + Novo
Relative Uptake (Da)
Exposure Time (minutes)
ACRA_ECOLI
QQGVTRTPRGDATV
310 - 323
1485.777 Da
Dimer
Dimer + Novo
Relative Uptake (Da)
Exposure Time (minutes)
ACRA_ECOLI
LVVGADDKVETRPIVA
324 - 339
1681.948 Da
Dimer
Dimer + Novo
Relative Uptake (Da)
Exposure Time (minutes)

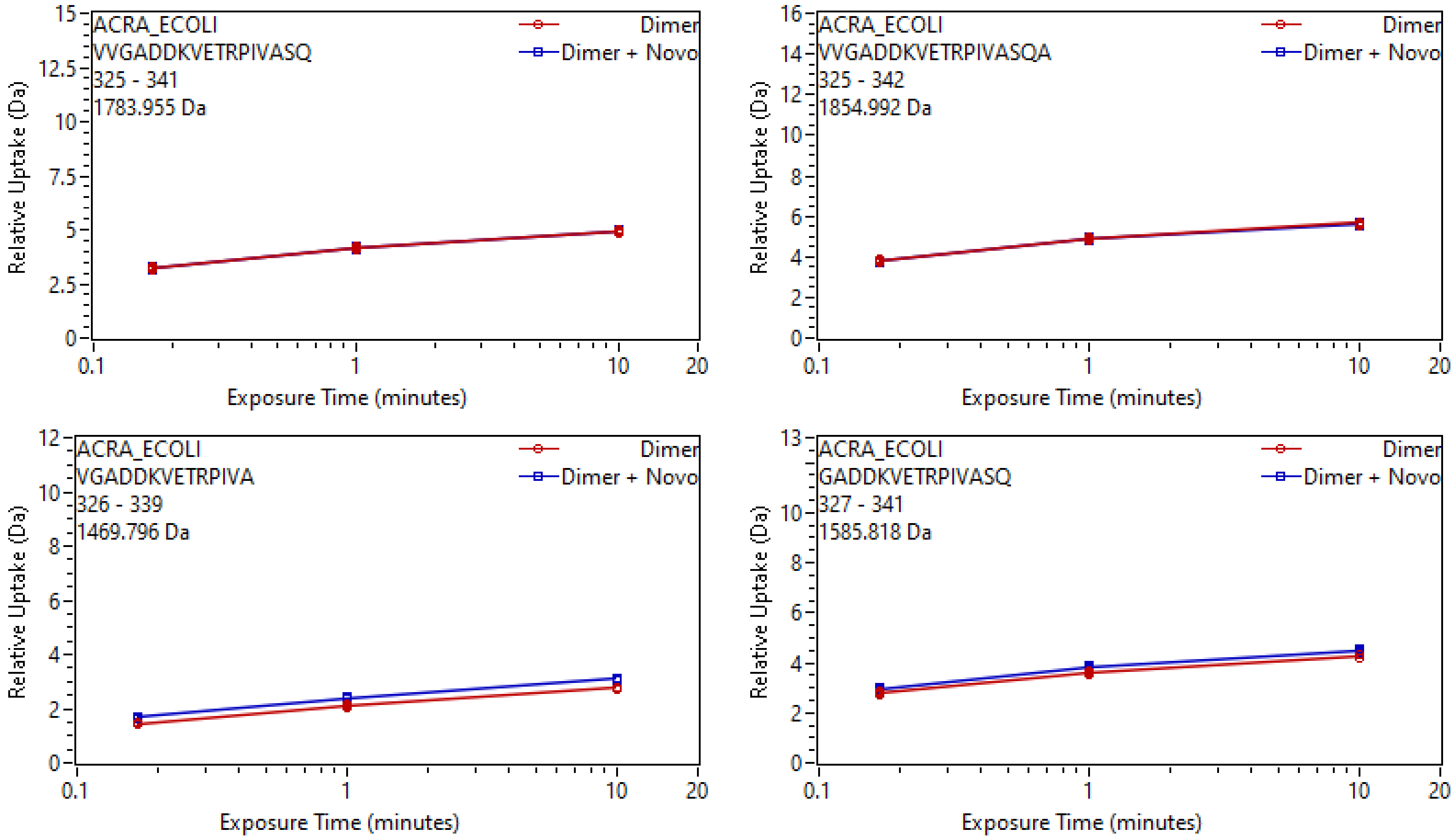
ACRA_ECOLI
VVGADDKVETRPIVASQ
325 - 341
1783.955 Da
Dimer
Dimer + Novo
Relative Uptake (Da)
Exposure Time (minutes)
ACRA_ECOLI
VVGADDKVETRPIVASQA
325 - 342
1854.992 Da
Dimer
Dimer + Novo
Relative Uptake (Da)
Exposure Time (minutes)
ACRA_ECOLI
VGADDKVETRPIVA
326 - 339
1469.796 Da
Dimer
Dimer + Novo
Relative Uptake (Da)
Exposure Time (minutes)
ACRA_ECOLI
GADDKVETRPIVASQ
327 - 341
1585.818 Da
Dimer
Dimer + Novo
Relative Uptake (Da)
Exposure Time (minutes)

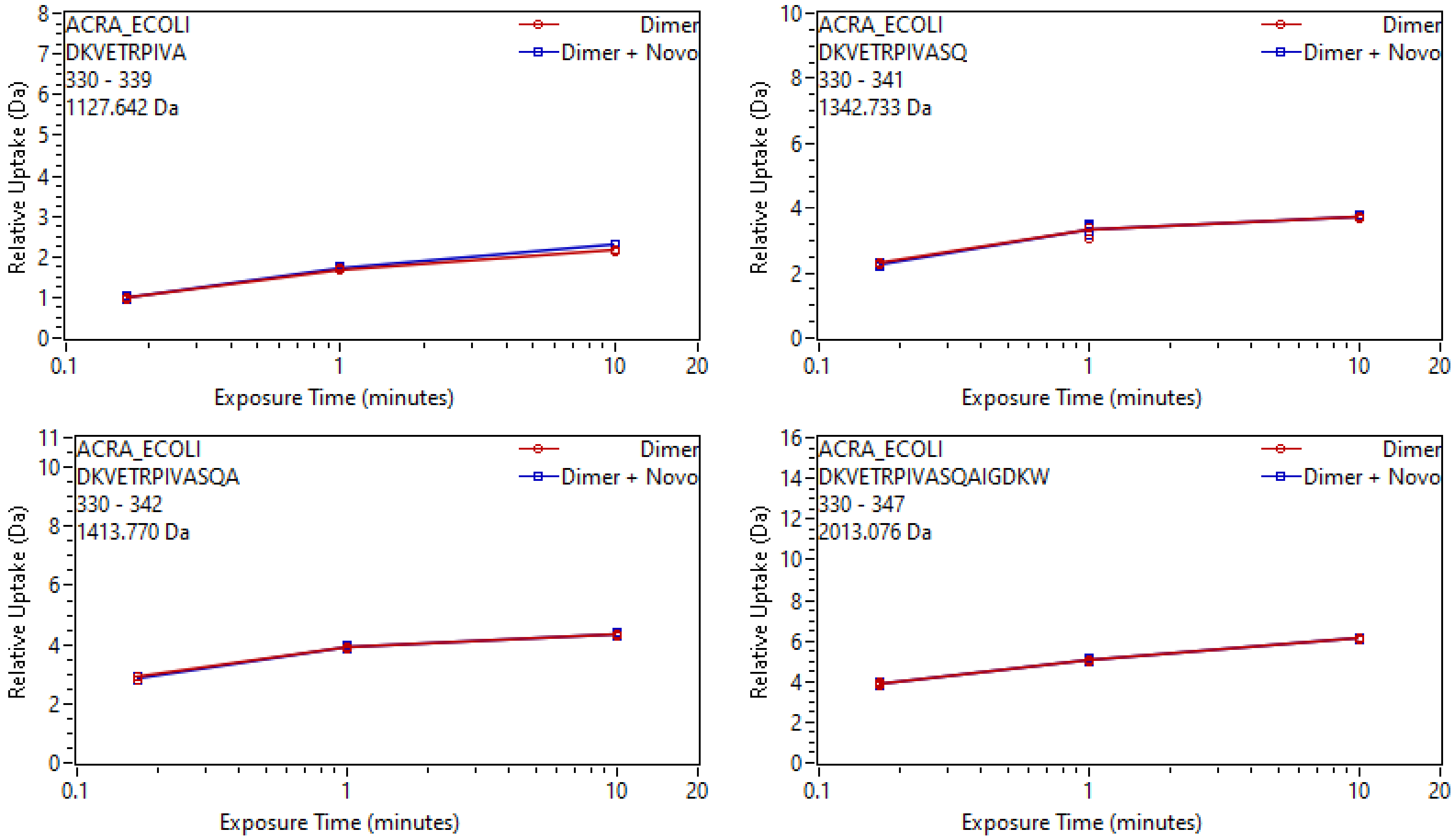
ACRA_ECOLI
DKVETRPIVA
330 - 339
1127.642 Da
Dimer
Dimer + Novo
Relative Uptake (Da)
Exposure Time (minutes)
ACRA_ECOLI
DKVETRPIVASQ
330 - 341
1342.733 Da
Dimer
Dimer + Novo
Relative Uptake (Da)
Exposure Time (minutes)
ACRA_ECOLI
DKVETRPIVASQA
330 - 342
1413.770 Da
Dimer
Dimer + Novo
Relative Uptake (Da)
Exposure Time (minutes)
ACRA_ECOLI
DKVETRPIVASQAIGDKW
330 - 347
2013.076 Da
Dimer
Dimer + Novo
Relative Uptake (Da)
Exposure Time (minutes)

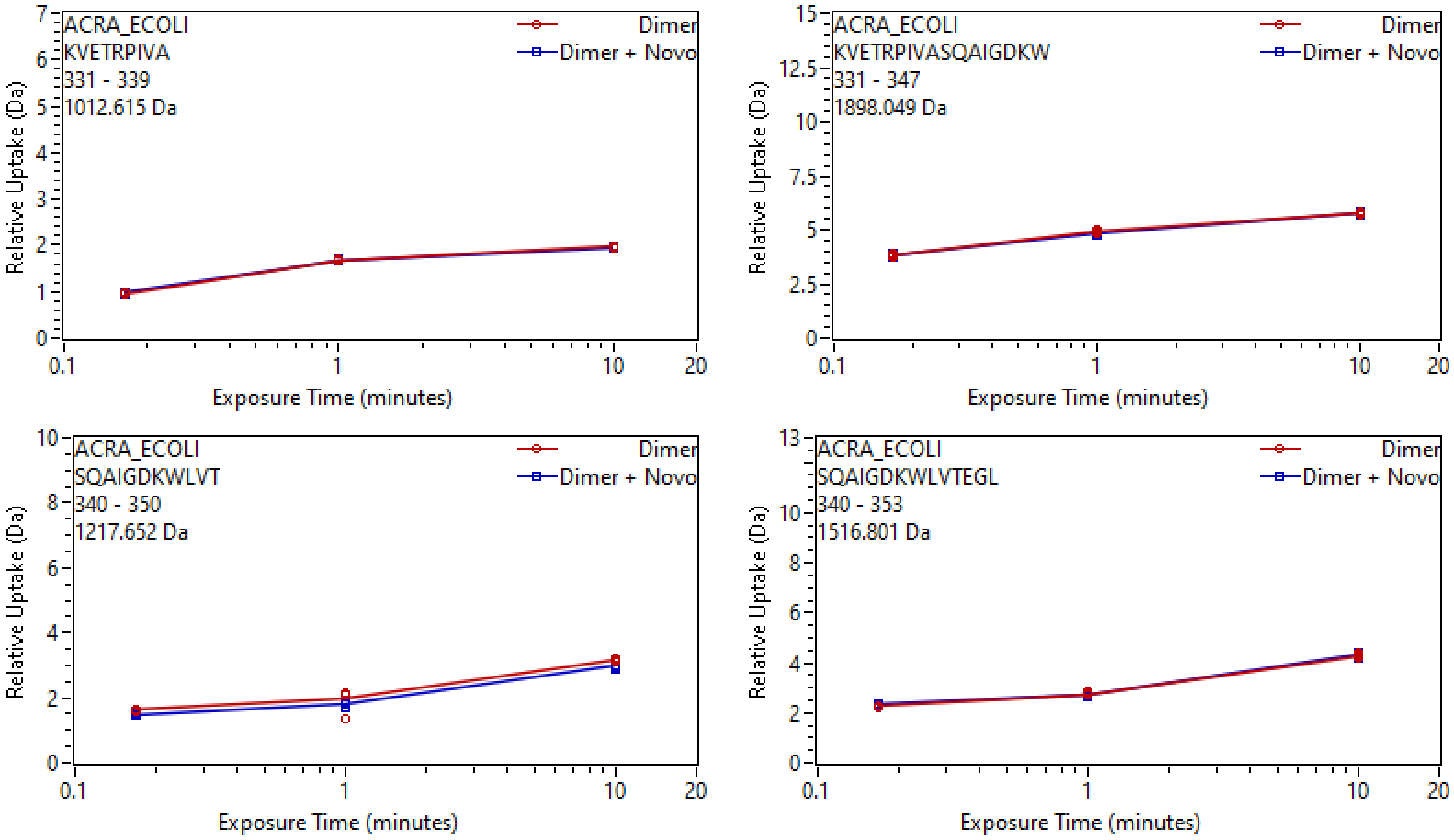
ACRA_ECOLI
KVETRPIVA
331 - 339
1012.615 Da
Dimer
Dimer + Novo
Relative Uptake (Da)
Exposure Time (minutes)
ACRA_ECOLI
KVETRPIVASQAIGDKW
331 - 347
1898.049 Da
Dimer
Dimer + Novo
Relative Uptake (Da)
Exposure Time (minutes)
ACRA_ECOLI
SQAIGDKWLVT
340 - 350
1217.652 Da
Dimer
Dimer + Novo
Relative Uptake (Da)
Exposure Time (minutes)
ACRA_ECOLI
SQAIGDKWLVTEGL
340 - 353
1516.801 Da
Dimer
Dimer + Novo
Relative Uptake (Da)
Exposure Time (minutes)

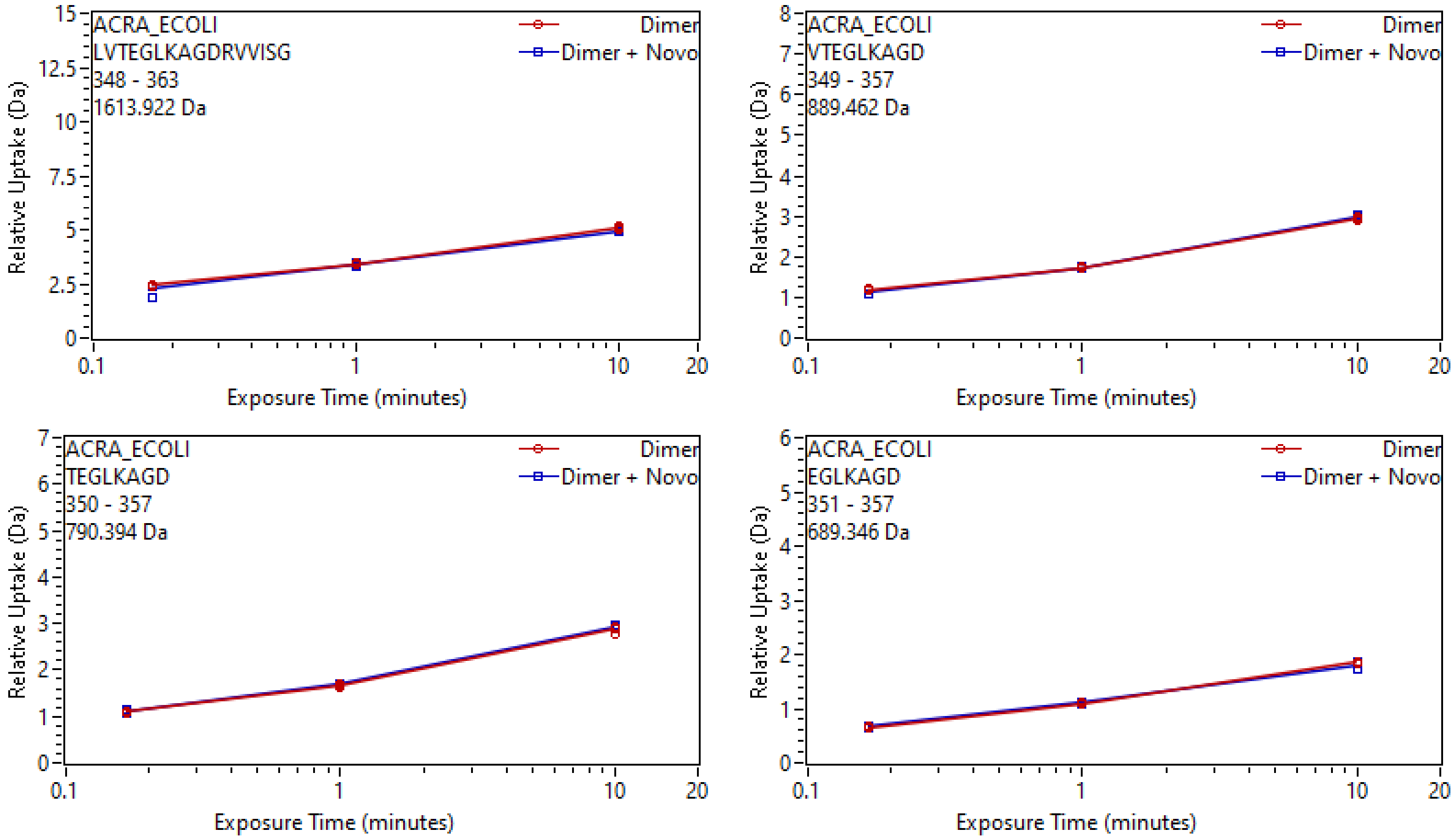
ACRA_ECOLI
LVTEGLKAGDRVVISG
348 - 363
1613.922 Da
Dimer
Dimer + Novo
Relative Uptake (Da)
Exposure Time (minutes)
ACRA_ECOLI
VTEGLKAGD
349 - 357
889.462 Da
Dimer
Dimer + Novo
Relative Uptake (Da)
Exposure Time (minutes)
ACRA_ECOLI
TEGLKAGD
350 - 357
790.394 Da
Dimer
Dimer + Novo
Relative Uptake (Da)
Exposure Time (minutes)
ACRA_ECOLI
EGLKAGD
351 - 357
689.346 Da
Dimer
Dimer + Novo
Relative Uptake (Da)
Exposure Time (minutes)

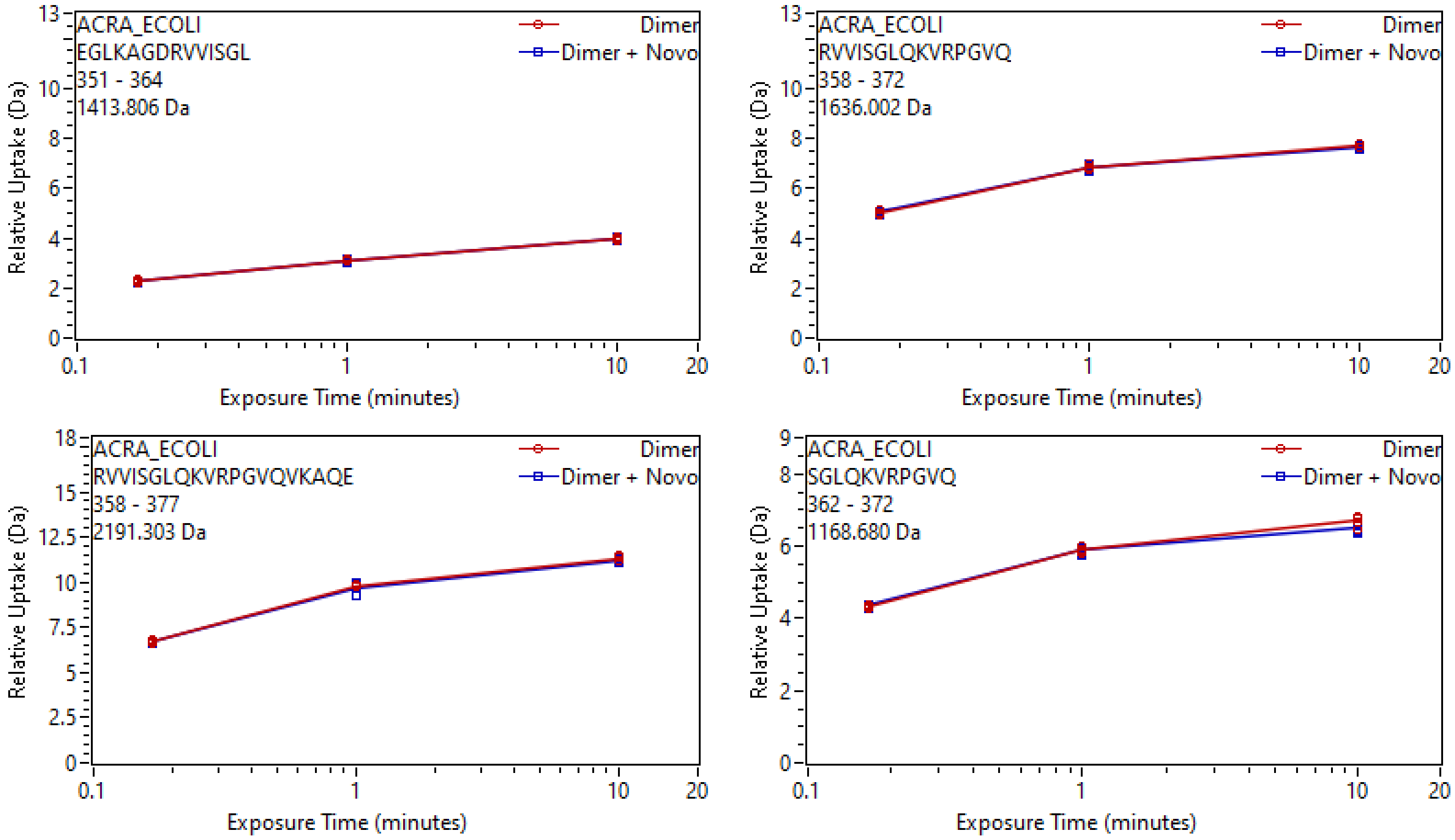
ACRA_ECOLI
EGLKAGDRVVISGL
351 - 364
1413.806 Da
Dimer
Dimer + Novo
Relative Uptake (Da)
Exposure Time (minutes)
ACRA_ECOLI
RVVISGLQKVRPGVQ
358 - 372
1636.002 Da
Dimer
Dimer + Novo
Relative Uptake (Da)
Exposure Time (minutes)
ACRA_ECOLI
RVVISGLQKVRPGVQVKAQE
358 - 377
2191.303 Da
Dimer
Dimer + Novo
Relative Uptake (Da)
Exposure Time (minutes)
ACRA_ECOLI
SGLQKVRPGVQ
362 - 372
1168.680 Da
Dimer
Dimer + Novo
Relative Uptake (Da)
Exposure Time (minutes)

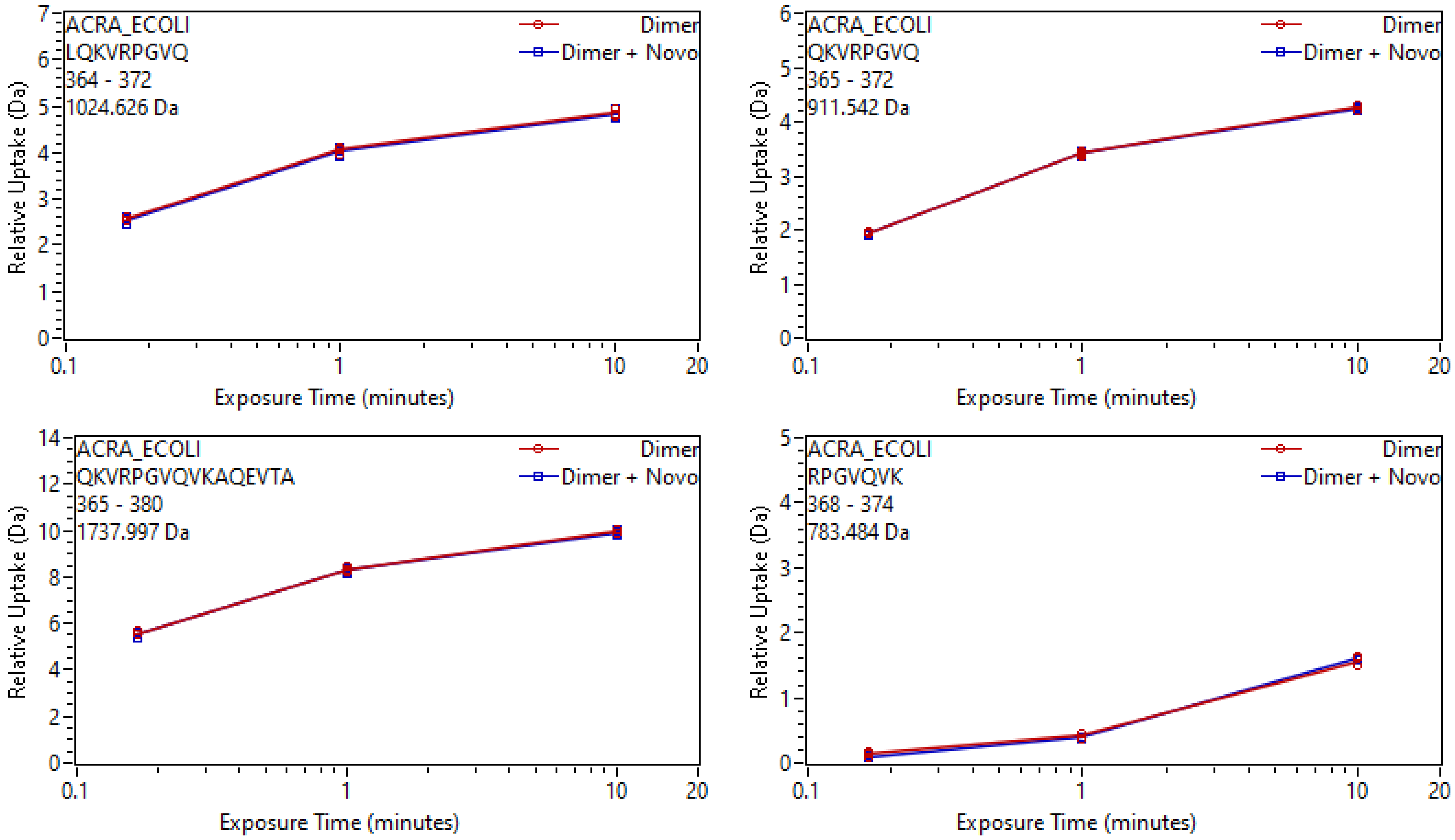

ACRA_ECOLI
LQKVRPGVQ
364 - 372
1024.626 Da
Dimer
Dimer + Novo
Relative Uptake (Da)
Exposure Time (minutes)
ACRA_ECOLI
QKVRPGVQ
365 - 372
911.542 Da
Dimer
Dimer + Novo
Relative Uptake (Da)
Exposure Time (minutes)
ACRA_ECOLI
QKVRPGVQVKAQEVTA
365 - 380
1737.997 Da
Dimer
Dimer + Novo
Relative Uptake (Da)
Exposure Time (minutes)
ACRA_ECOLI
RPGVQVK
368 - 374
783.484 Da
Dimer
Dimer + Novo
Relative Uptake (Da)
Exposure Time (minutes)

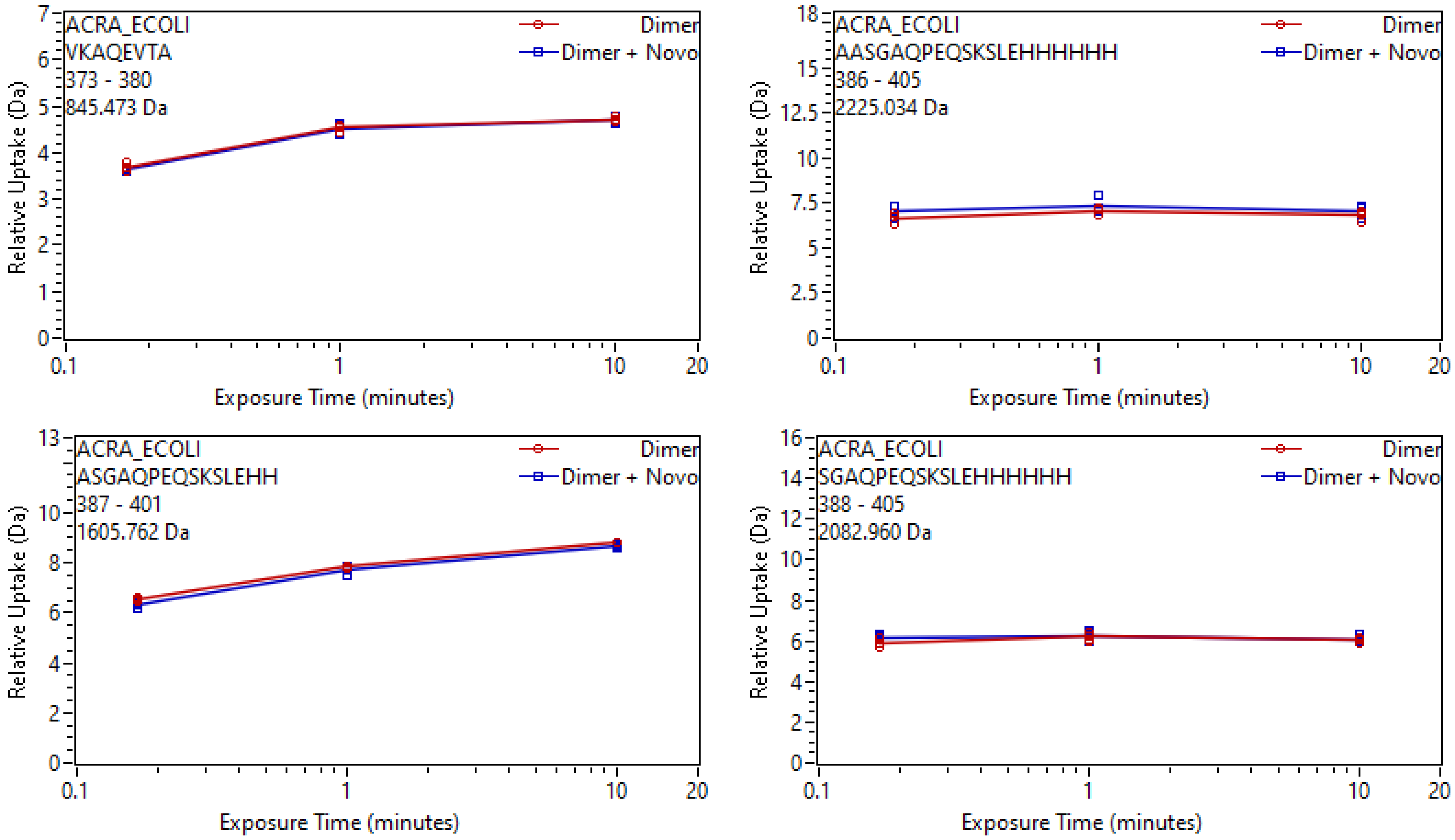
ACRA_ECOLI
VKAQEVTA
373 - 380
845.473 Da
Dimer
Dimer + Novo
Relative Uptake (Da)
Exposure Time (minutes)
ACRA_ECOLI
AASGAQPEQSKSLEHHHHHH
386 - 405
2225.034 Da
Dimer
Dimer + Novo
Relative Uptake (Da)
Exposure Time (minutes)
ACRA_ECOLI
ASGAQPEQSKSLEHH
387 - 401
1605.762 Da
Dimer
Dimer + Novo
Relative Uptake (Da)
Exposure Time (minutes)
ACRA_ECOLI
SGAQPEQSKSLEHHHHHH
388 - 405
2082.960 Da
Dimer
Dimer + Novo
Relative Uptake (Da)
Exposure Time (minutes)

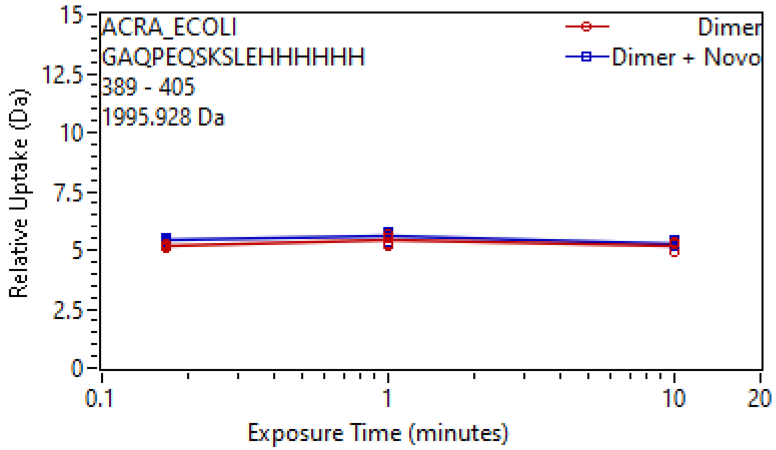
ACRA_ECOLI
GAQPEQSKSLEHHHHHH
389 - 405
1995.928 Da
Dimer
Dimer + Novo
Relative Uptake (Da)
Exposure Time (minutes)
0
2.5
5
7.5
10
12.5
15
0.1
1
10
20

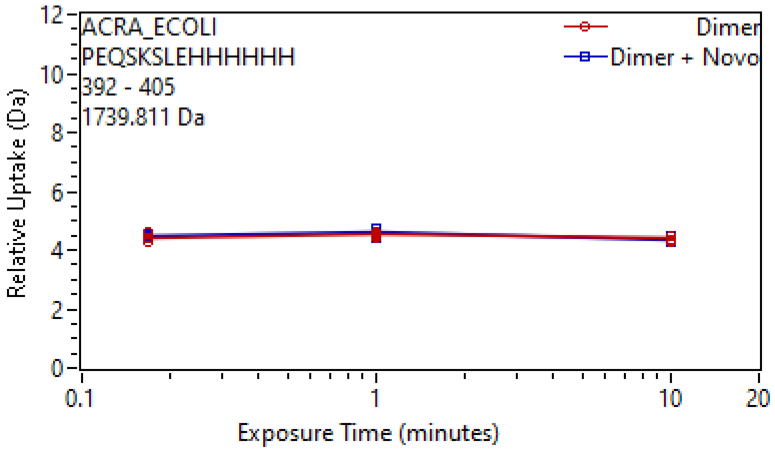
ACRA_ECOLI
PEQSKSLEHHHHHH
392 - 405
1739.811 Da
Dimer
Dimer + Novo
Relative Uptake (Da)
Exposure Time (minutes)
0
2
4
6
8
10
12
0.1
1
10
20
