## Supplementary material for "Conformational restriction shapes inhibition of a multidrug efflux adaptor protein": UptakePlots5

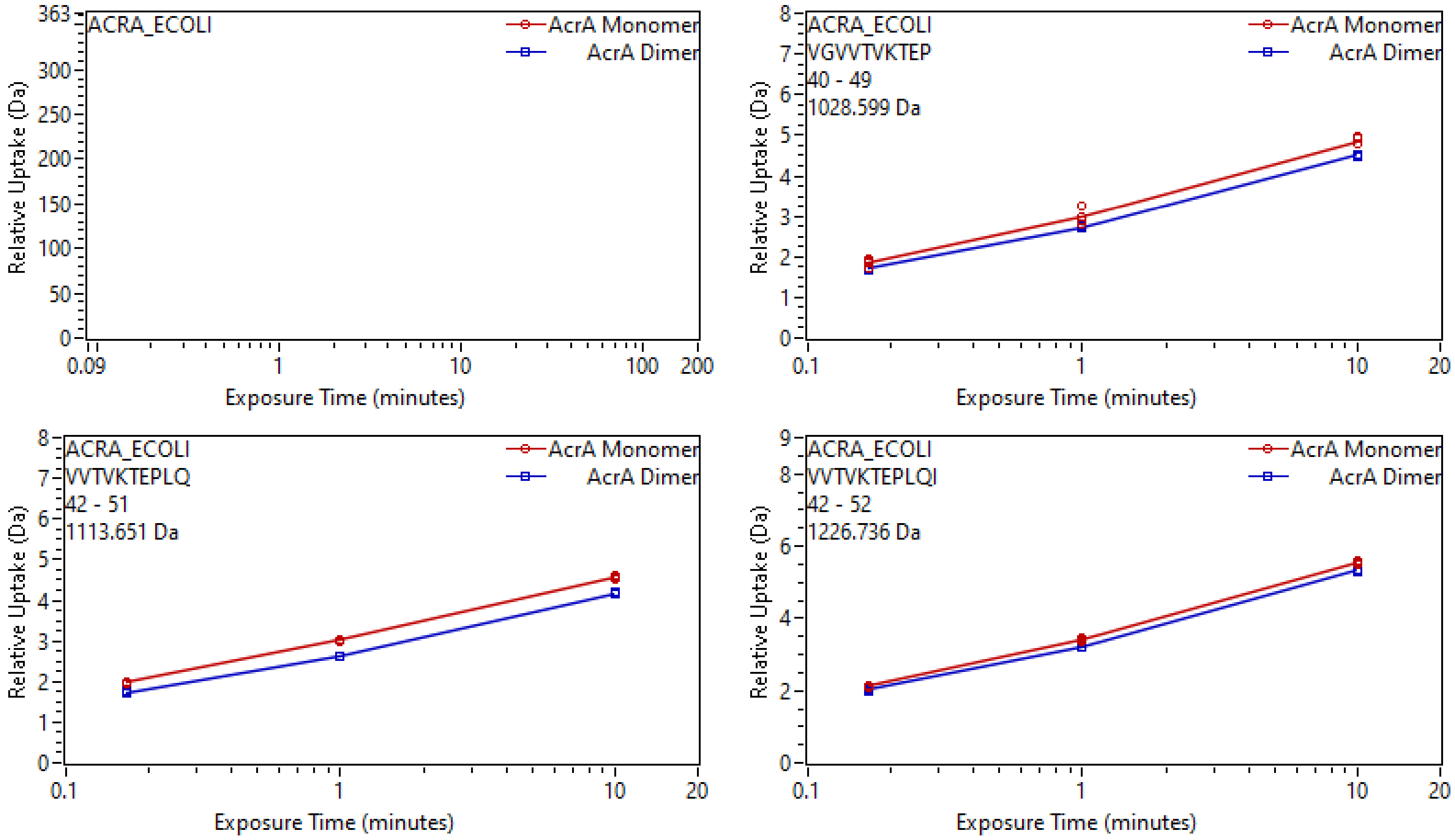
ACRA_ECOLI
AcrA Monomer
AcrA Dimer
Relative Uptake (Da)
Exposure Time (minutes)
ACRA_ECOLI
VGVVTVKTEP
40 - 49
1028.599 Da
AcrA Monomer
AcrA Dimer
Relative Uptake (Da)
Exposure Time (minutes)
ACRA_ECOLI
VVTVKTEPLQ
42 - 51
1113.651 Da
AcrA Monomer
AcrA Dimer
Relative Uptake (Da)
Exposure Time (minutes)
ACRA_ECOLI
VVTVKTEPLQI
42 - 52
1226.736 Da
AcrA Monomer
AcrA Dimer
Relative Uptake (Da)
Exposure Time (minutes)

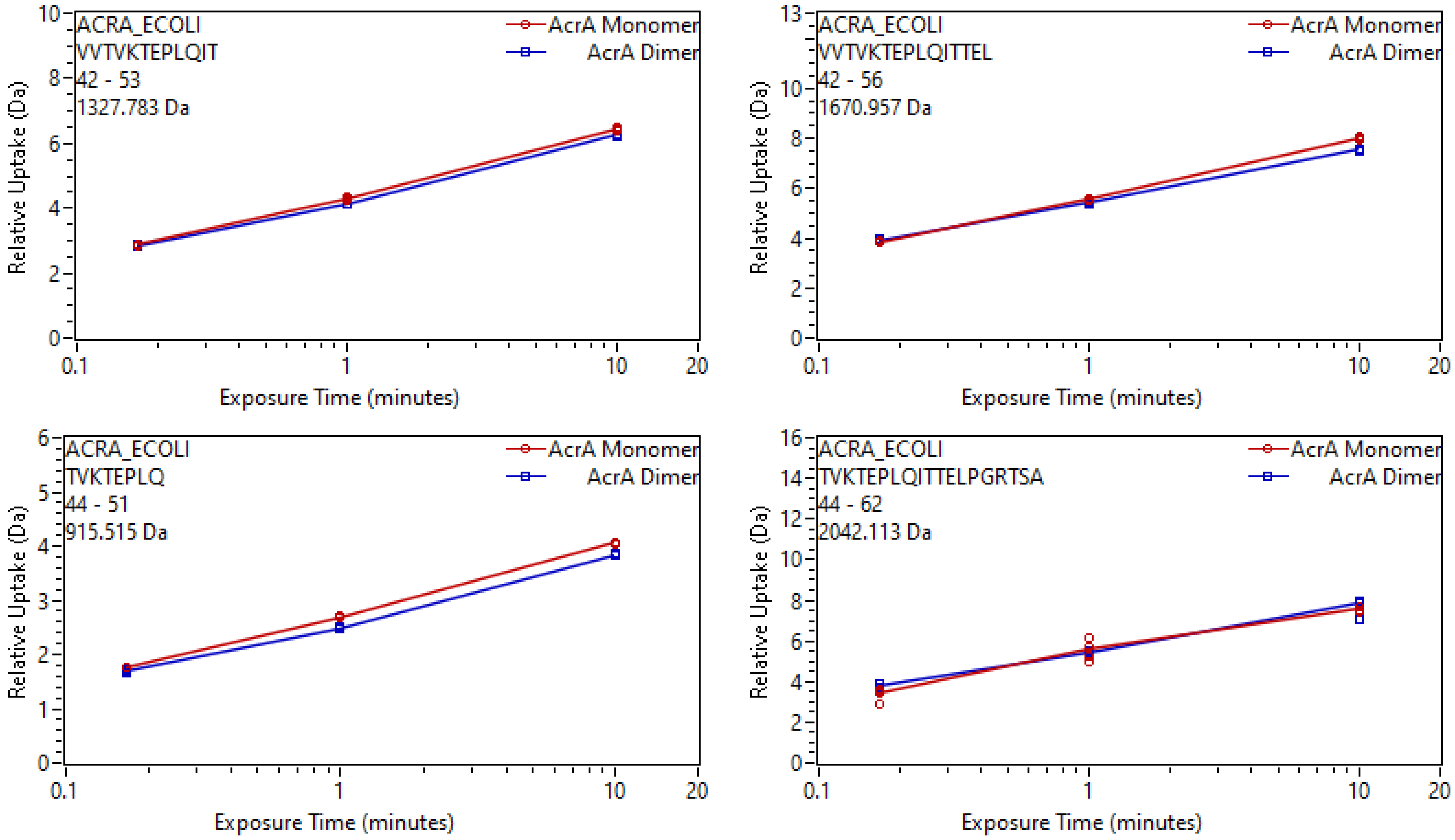
ACRA_ECOLI
VVTVKTEPLQIT
42 - 53
1327.783 Da
AcrA Monomer
AcrA Dimer
Relative Uptake (Da)
Exposure Time (minutes)
ACRA_ECOLI
VVTVKTEPLQITTEL
42 - 56
1670.957 Da
AcrA Monomer
AcrA Dimer
Relative Uptake (Da)
Exposure Time (minutes)
ACRA_ECOLI
TVKTEPLQ
44 - 51
915.515 Da
AcrA Monomer
AcrA Dimer
Relative Uptake (Da)
Exposure Time (minutes)
ACRA_ECOLI
TVKTEPLQITTELPGRTSA
44 - 62
2042.113 Da
AcrA Monomer
AcrA Dimer
Relative Uptake (Da)
Exposure Time (minutes)

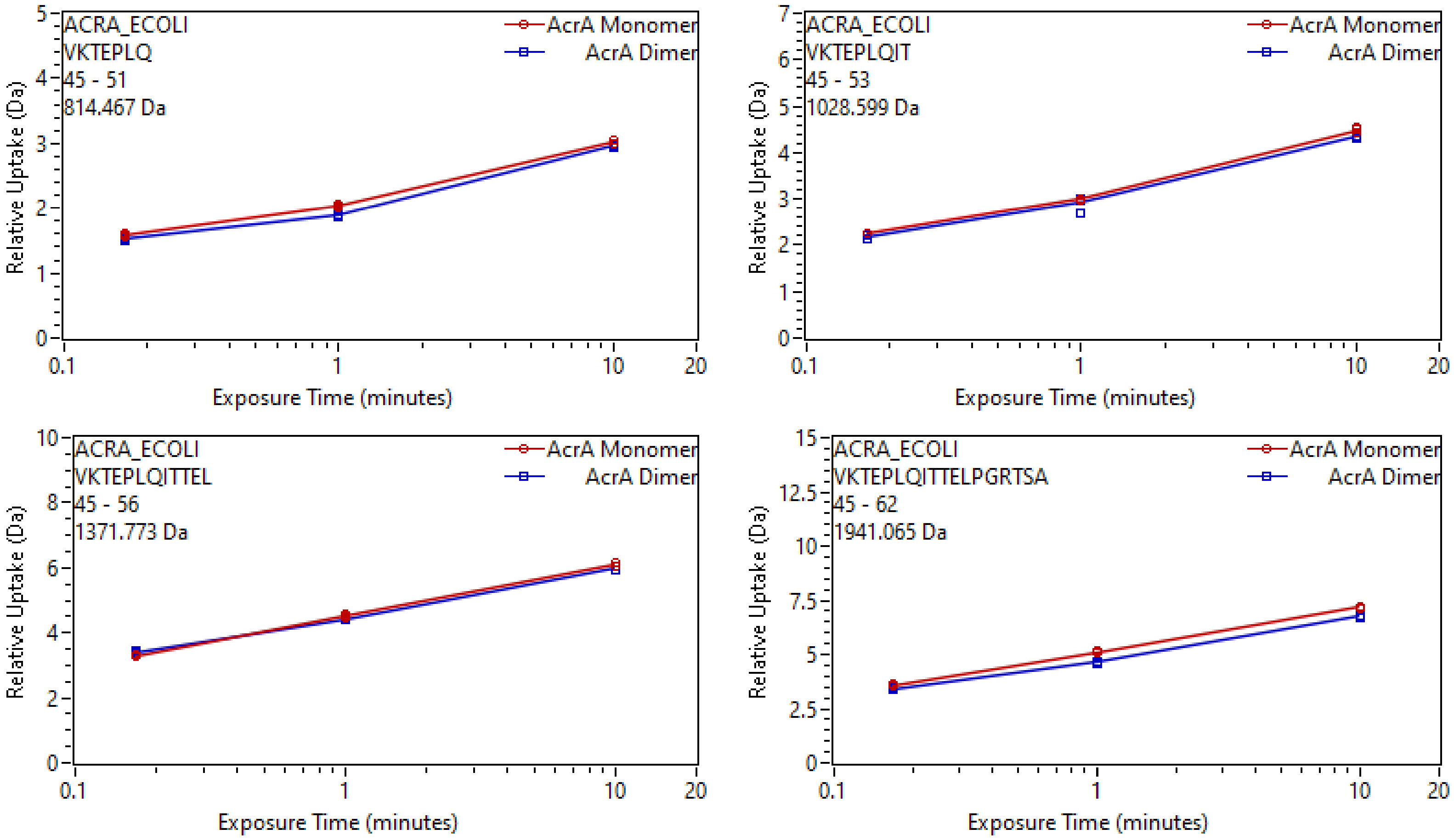
ACRA_ECOLI
VKTEPLQ
45 - 51
814.467 Da
AcrA Monomer
AcrA Dimer
Relative Uptake (Da)
Exposure Time (minutes)
0 1 2 3 4 5
0.1 1 10 20
ACRA_ECOLI
VKTEPLQIT
45 - 53
1028.599 Da
AcrA Monomer
AcrA Dimer
Relative Uptake (Da)
Exposure Time (minutes)
0 1 2 3 4 5 6 7
0.1 1 10 20
ACRA_ECOLI
VKTEPLQITTEL
45 - 56
1371.773 Da
AcrA Monomer
AcrA Dimer
Relative Uptake (Da)
Exposure Time (minutes)
0 2 4 6 8 10
0.1 1 10 20
ACRA_ECOLI
VKTEPLQITTELPGRTSA
45 - 62
1941.065 Da
AcrA Monomer
AcrA Dimer
Relative Uptake (Da)
Exposure Time (minutes)
0 2.5 5 7.5 10 12.5 15
0.1 1 10 20

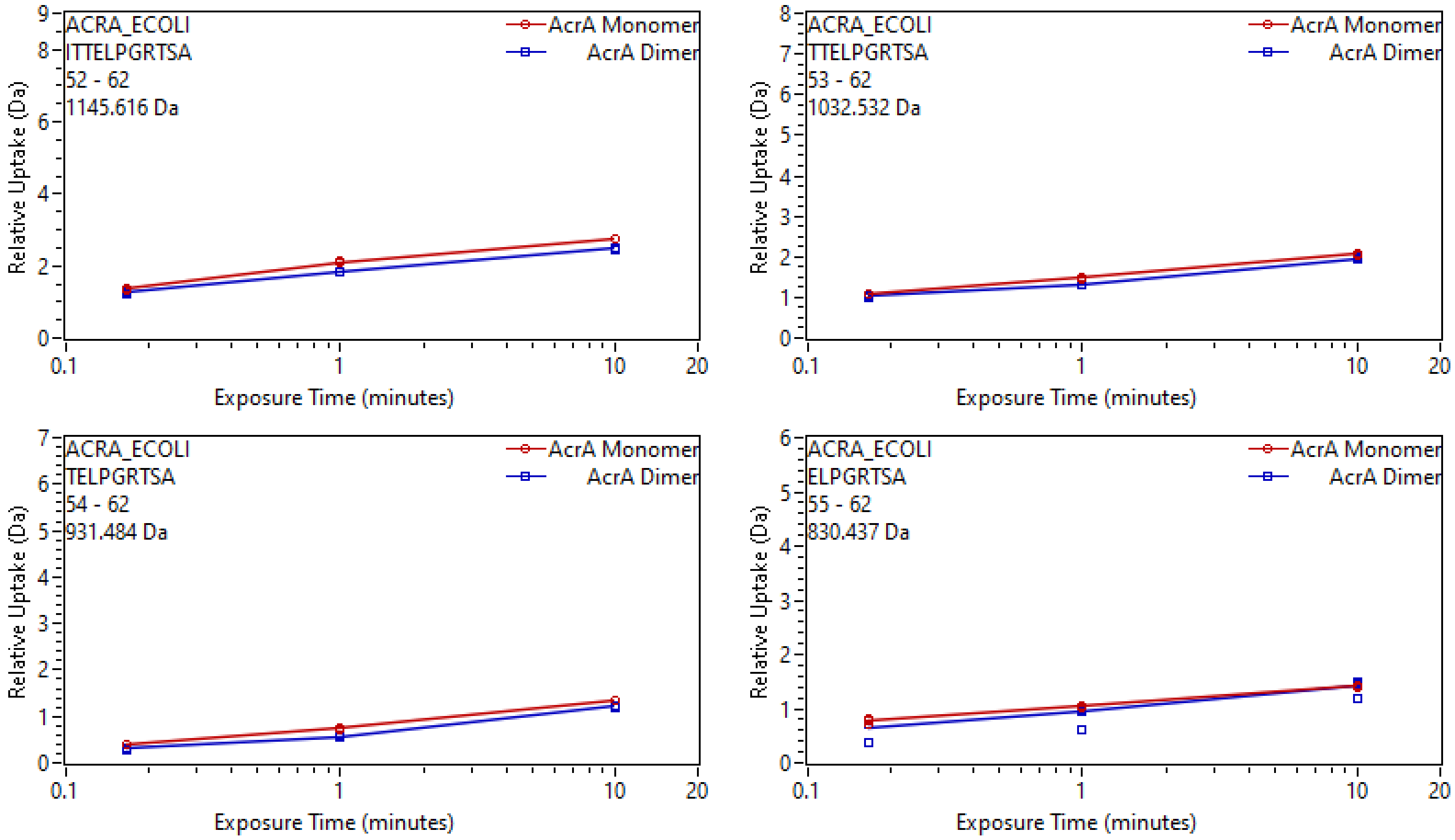

ACRA_ECOLI
ITTELPGRTSA
52 - 62
1145.616 Da
AcrA Monomer
AcrA Dimer
Relative Uptake (Da)
Exposure Time (minutes)
ACRA_ECOLI
TTELPGRTSA
53 - 62
1032.532 Da
AcrA Monomer
AcrA Dimer
Relative Uptake (Da)
Exposure Time (minutes)
ACRA_ECOLI
TELPGRTSA
54 - 62
931.484 Da
AcrA Monomer
AcrA Dimer
Relative Uptake (Da)
Exposure Time (minutes)
ACRA_ECOLI
ELPGRTSA
55 - 62
830.437 Da
AcrA Monomer
AcrA Dimer
Relative Uptake (Da)
Exposure Time (minutes)

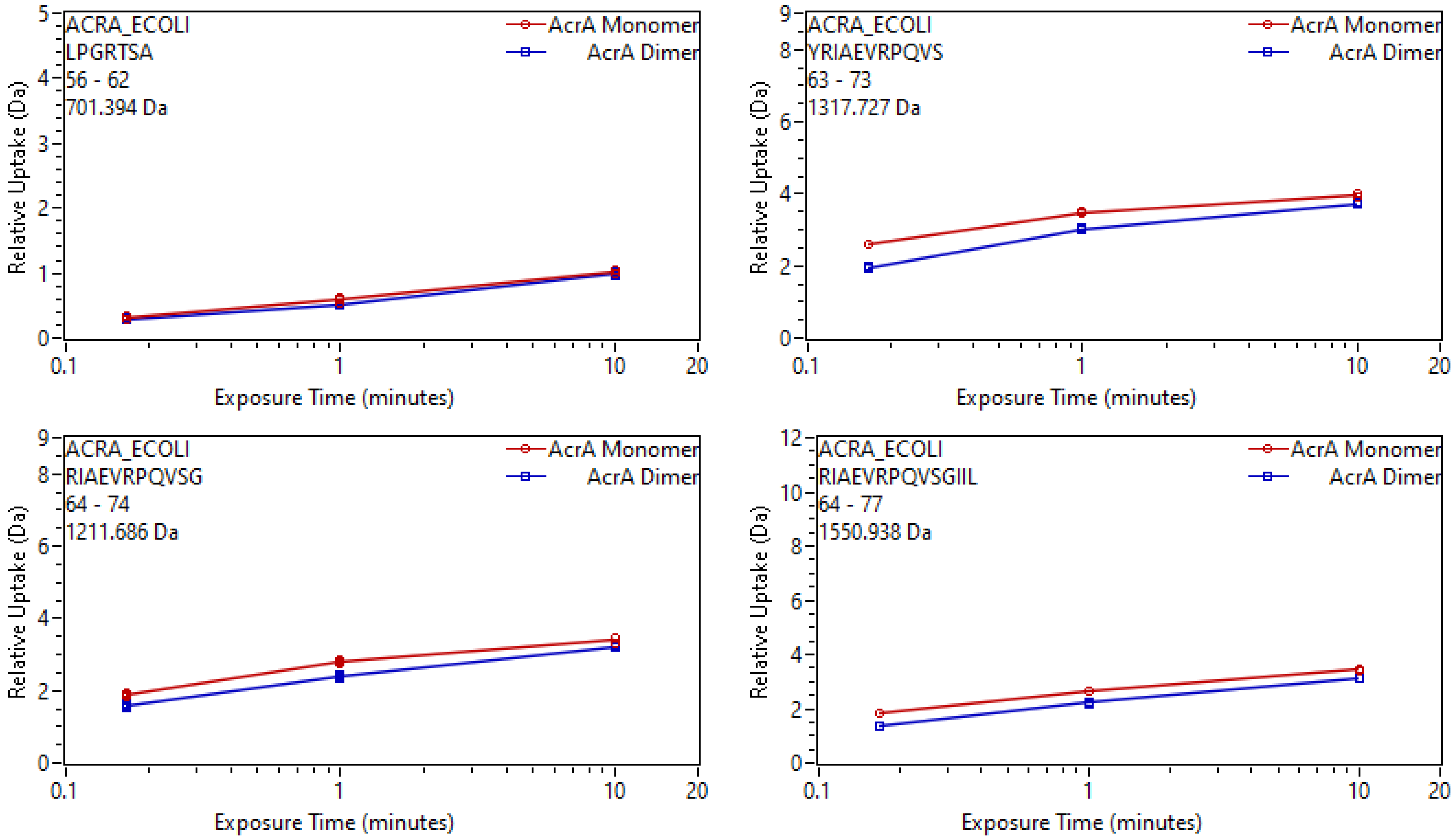
ACRA_ECOLI
LPGRTSA
56 - 62
701.394 Da
AcrA Monomer
AcrA Dimer
Relative Uptake (Da)
Exposure Time (minutes)
ACRA_ECOLI
YRIAEVRPQVS
63 - 73
1317.727 Da
AcrA Monomer
AcrA Dimer
Relative Uptake (Da)
Exposure Time (minutes)
ACRA_ECOLI
RIAEVRPQVSG
64 - 74
1211.686 Da
AcrA Monomer
AcrA Dimer
Relative Uptake (Da)
Exposure Time (minutes)
ACRA_ECOLI
RIAEVRPQVSGIIL
64 - 77
1550.938 Da
AcrA Monomer
AcrA Dimer
Relative Uptake (Da)
Exposure Time (minutes)

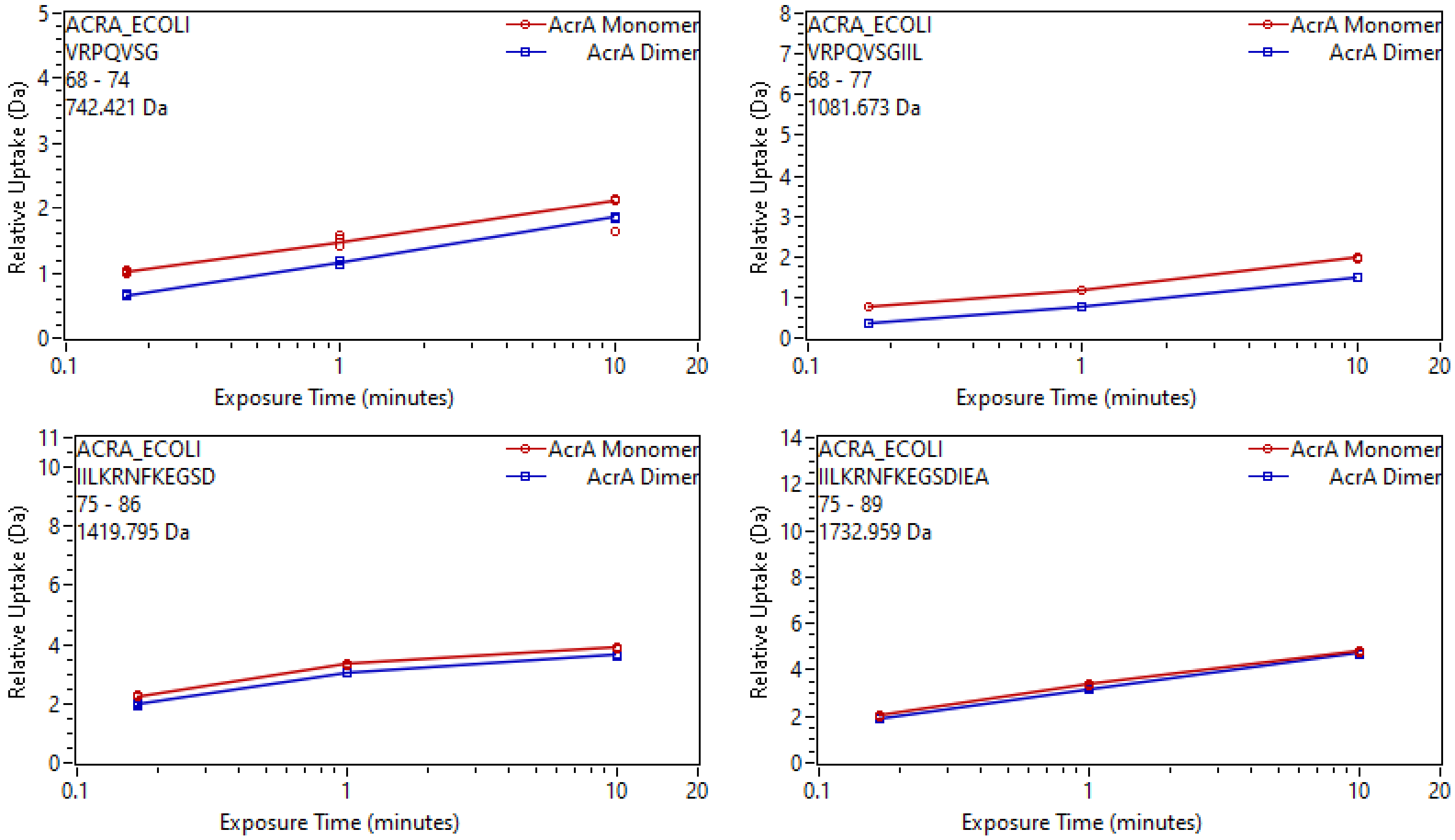
ACRA_ECOLI
VRPQVSG
68 - 74
742.421 Da
AcrA Monomer
AcrA Dimer
Relative Uptake (Da)
Exposure Time (minutes)
ACRA_ECOLI
VRPQVSGIIL
68 - 77
1081.673 Da
AcrA Monomer
AcrA Dimer
Relative Uptake (Da)
Exposure Time (minutes)
ACRA_ECOLI
IILKRNFKEGSD
75 - 86
1419.795 Da
AcrA Monomer
AcrA Dimer
Relative Uptake (Da)
Exposure Time (minutes)
ACRA_ECOLI
IILKRNFKEGSDIEA
75 - 89
1732.959 Da
AcrA Monomer
AcrA Dimer
Relative Uptake (Da)
Exposure Time (minutes)

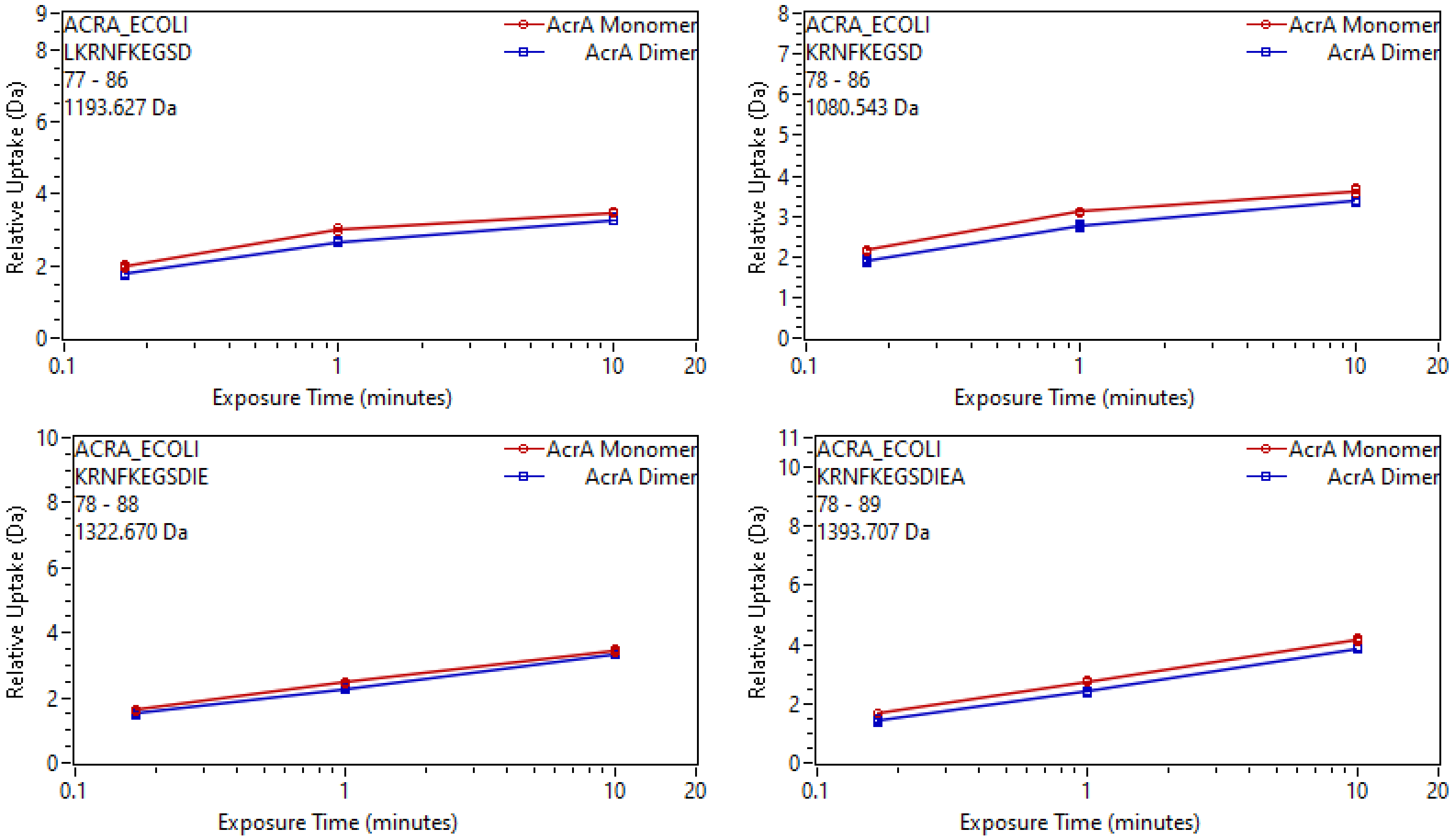
ACRA_ECOLI
LKRNFKEGSD
77 - 86
1193.627 Da
AcrA Monomer
AcrA Dimer
Relative Uptake (Da)
Exposure Time (minutes)
ACRA_ECOLI
KRNFKEGSD
78 - 86
1080.543 Da
AcrA Monomer
AcrA Dimer
Relative Uptake (Da)
Exposure Time (minutes)
ACRA_ECOLI
KRNFKEGSDIE
78 - 88
1322.670 Da
AcrA Monomer
AcrA Dimer
Relative Uptake (Da)
Exposure Time (minutes)
ACRA_ECOLI
KRNFKEGSDIEA
78 - 89
1393.707 Da
AcrA Monomer
AcrA Dimer
Relative Uptake (Da)
Exposure Time (minutes)

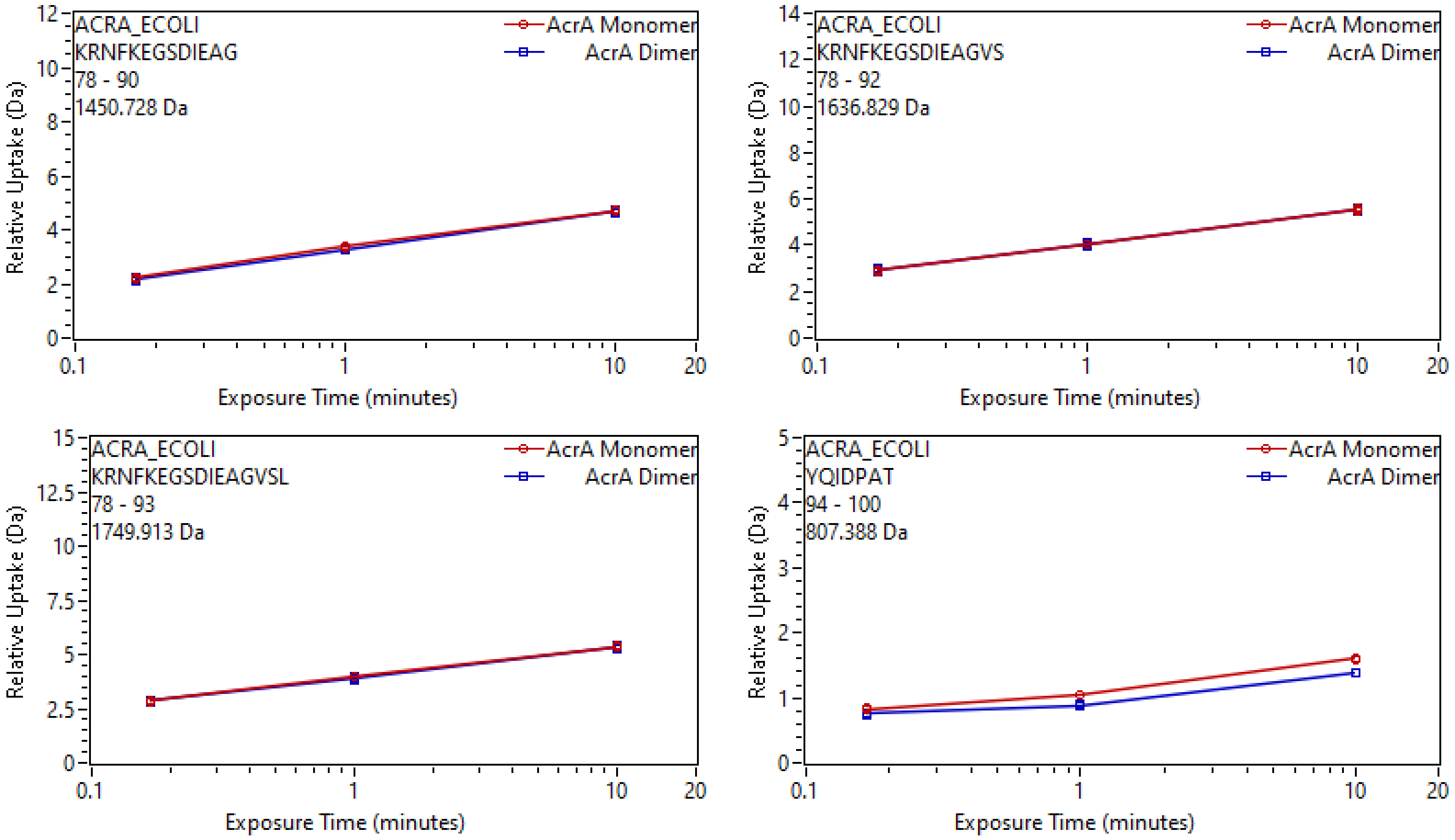
ACRA_ECOLI
KRNFKEGSDIEAG
78 - 90
1450.728 Da
AcrA Monomer
AcrA Dimer
Relative Uptake (Da)
Exposure Time (minutes)
ACRA_ECOLI
KRNFKEGSDIEAGVS
78 - 92
1636.829 Da
AcrA Monomer
AcrA Dimer
Relative Uptake (Da)
Exposure Time (minutes)
ACRA_ECOLI
KRNFKEGSDIEAGVSL
78 - 93
1749.913 Da
AcrA Monomer
AcrA Dimer
Relative Uptake (Da)
Exposure Time (minutes)
ACRA_ECOLI
YQIDPAT
94 - 100
807.388 Da
AcrA Monomer
AcrA Dimer
Relative Uptake (Da)
Exposure Time (minutes)

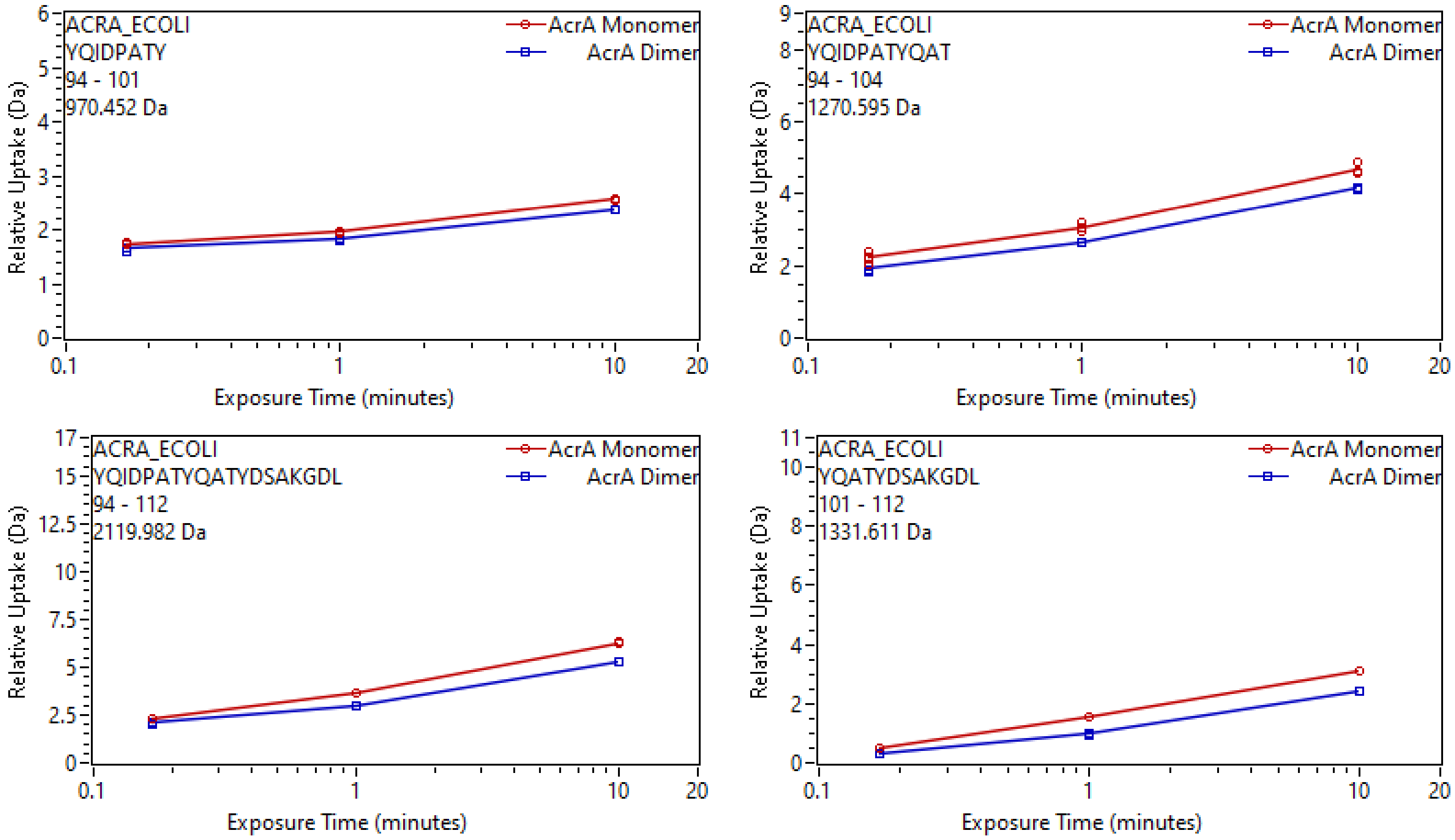
ACRA_ECOLI
YQIDPATY
94 - 101
970.452 Da
AcrA Monomer
AcrA Dimer
Relative Uptake (Da)
Exposure Time (minutes)
ACRA_ECOLI
YQIDPATYQAT
94 - 104
1270.595 Da
AcrA Monomer
AcrA Dimer
Relative Uptake (Da)
Exposure Time (minutes)
ACRA_ECOLI
YQIDPATYQATYDSAKGDL
94 - 112
2119.982 Da
AcrA Monomer
AcrA Dimer
Relative Uptake (Da)
Exposure Time (minutes)
ACRA_ECOLI
YQATYDSAKGDL
101 - 112
1331.611 Da
AcrA Monomer
AcrA Dimer
Relative Uptake (Da)
Exposure Time (minutes)

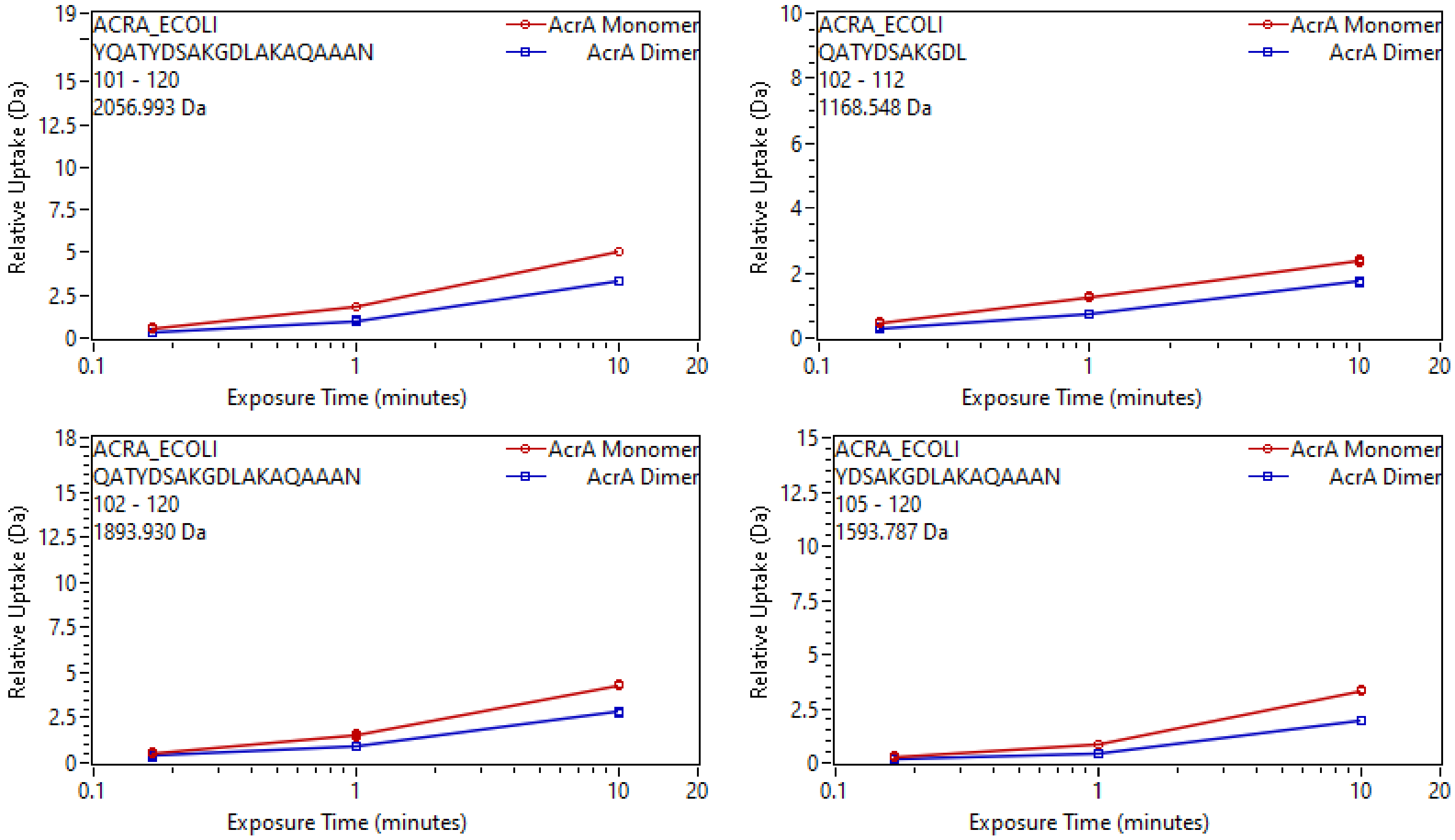
ACRA_ECOLI
YQATYDSAKGDLAKAQAAAN
101 - 120
2056.993 Da
AcrA Monomer
AcrA Dimer
Relative Uptake (Da)
Exposure Time (minutes)
ACRA_ECOLI
QATYDSAKGDL
102 - 112
1168.548 Da
AcrA Monomer
AcrA Dimer
Relative Uptake (Da)
Exposure Time (minutes)
ACRA_ECOLI
QATYDSAKGDLAKAQAAAN
102 - 120
1893.930 Da
AcrA Monomer
AcrA Dimer
Relative Uptake (Da)
Exposure Time (minutes)
ACRA_ECOLI
YDSAKGDLAKAQAAAN
105 - 120
1593.787 Da
AcrA Monomer
AcrA Dimer
Relative Uptake (Da)
Exposure Time (minutes)

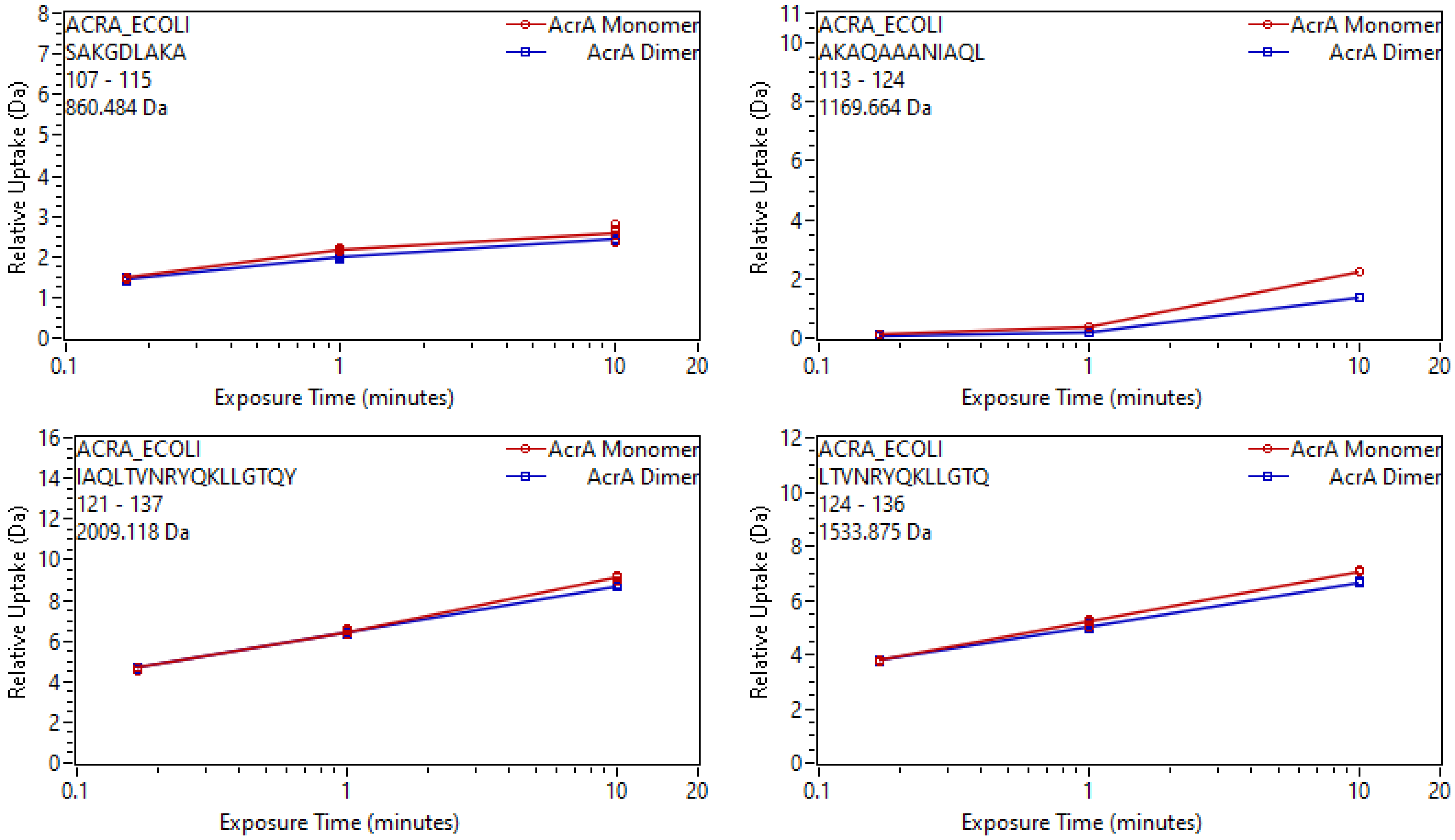
ACRA_ECOLI
SAKGDLAKA
107 - 115
860.484 Da
AcrA Monomer
AcrA Dimer
Relative Uptake (Da)
Exposure Time (minutes)
ACRA_ECOLI
AKAQAAANIAQL
113 - 124
1169.664 Da
AcrA Monomer
AcrA Dimer
Relative Uptake (Da)
Exposure Time (minutes)
ACRA_ECOLI
IAQLTVNRYQKLLGTQY
121 - 137
2009.118 Da
AcrA Monomer
AcrA Dimer
Relative Uptake (Da)
Exposure Time (minutes)
ACRA_ECOLI
LTVNRYQKLLGTQ
124 - 136
1533.875 Da
AcrA Monomer
AcrA Dimer
Relative Uptake (Da)
Exposure Time (minutes)

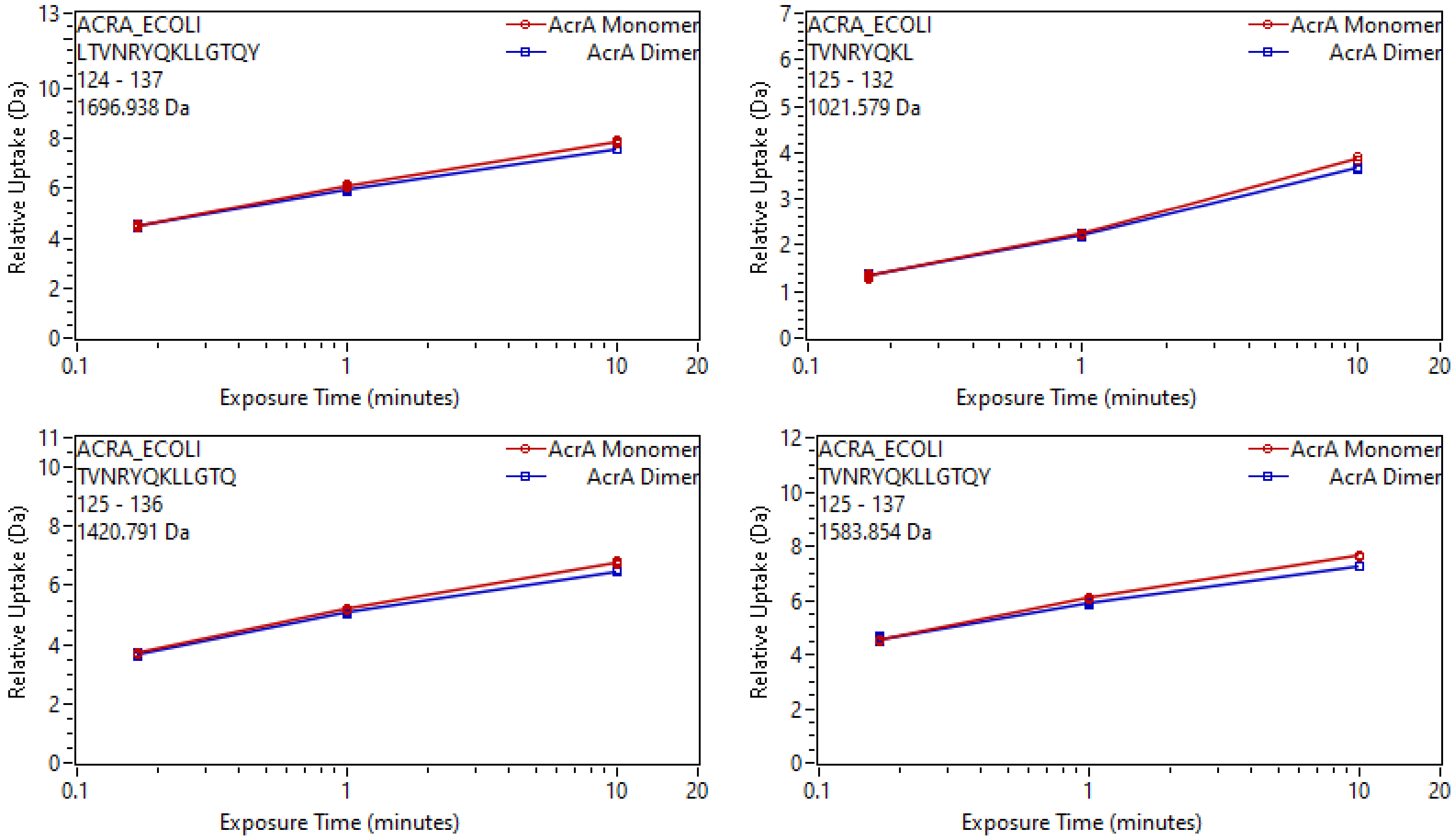
ACRA_ECOLI
LTVNRYQKLLGTQY
124 - 137
1696.938 Da
AcrA Monomer
AcrA Dimer
Relative Uptake (Da)
Exposure Time (minutes)
ACRA_ECOLI
TVNRYQKL
125 - 132
1021.579 Da
AcrA Monomer
AcrA Dimer
Relative Uptake (Da)
Exposure Time (minutes)
ACRA_ECOLI
TVNRYQKLLGTQ
125 - 136
1420.791 Da
AcrA Monomer
AcrA Dimer
Relative Uptake (Da)
Exposure Time (minutes)
ACRA_ECOLI
TVNRYQKLLGTQY
125 - 137
1583.854 Da
AcrA Monomer
AcrA Dimer
Relative Uptake (Da)
Exposure Time (minutes)

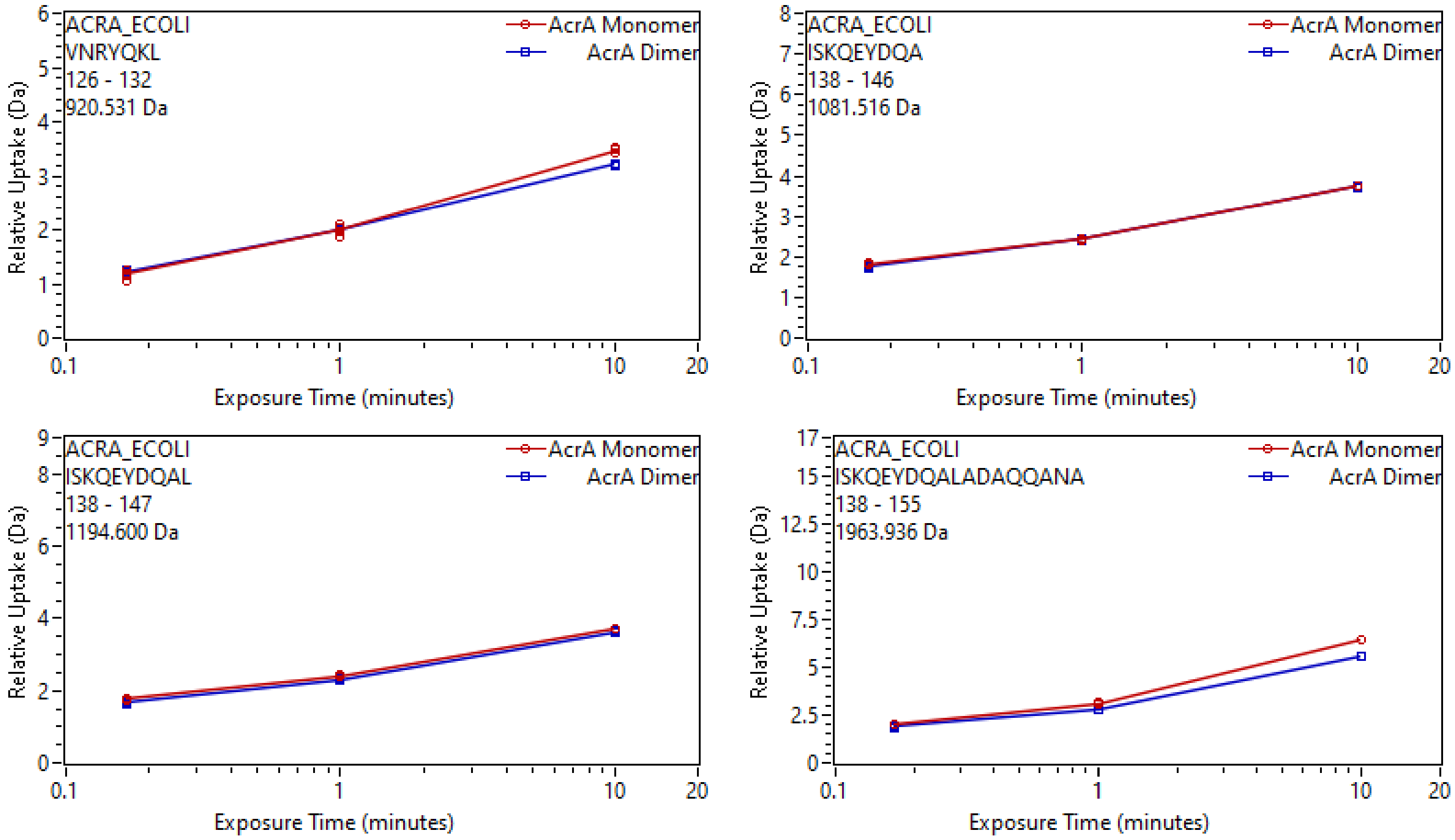
ACRA_ECOLI
VNRYQKL
126 - 132
920.531 Da
AcrA Monomer
AcrA Dimer
Relative Uptake (Da)
Exposure Time (minutes)
ACRA_ECOLI
ISKQEYDQA
138 - 146
1081.516 Da
AcrA Monomer
AcrA Dimer
Relative Uptake (Da)
Exposure Time (minutes)
ACRA_ECOLI
ISKQEYDQAL
138 - 147
1194.600 Da
AcrA Monomer
AcrA Dimer
Relative Uptake (Da)
Exposure Time (minutes)
ACRA_ECOLI
ISKQEYDQALADAQQANA
138 - 155
1963.936 Da
AcrA Monomer
AcrA Dimer
Relative Uptake (Da)
Exposure Time (minutes)

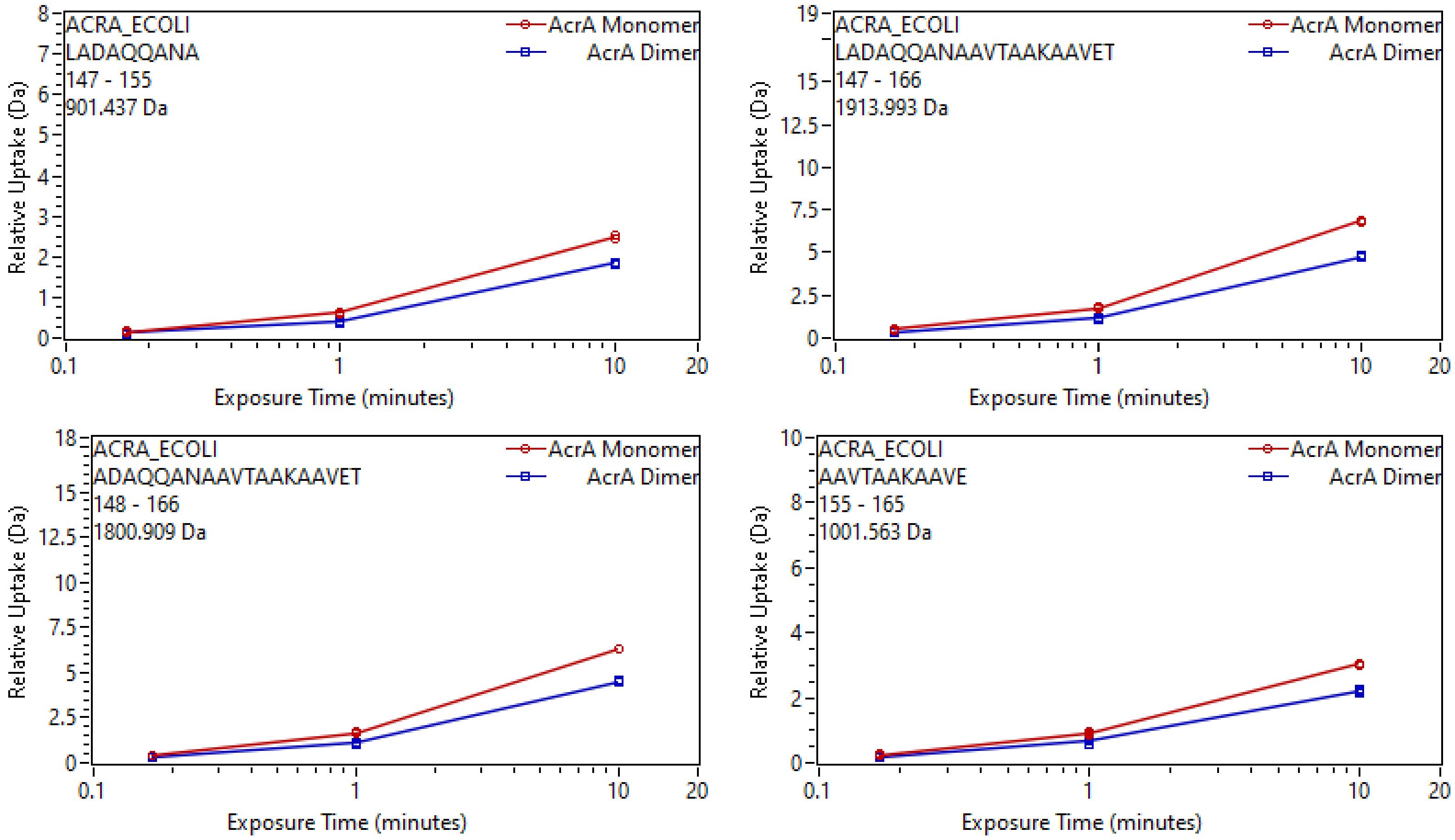
ACRA_ECOLI
LADAQQANA
147 - 155
901.437 Da
AcrA Monomer
AcrA Dimer
Relative Uptake (Da)
Exposure Time (minutes)
ACRA_ECOLI
LADAQQANAAVTAAKAAVET
147 - 166
1913.993 Da
AcrA Monomer
AcrA Dimer
Relative Uptake (Da)
Exposure Time (minutes)
ACRA_ECOLI
ADAQQANAAVTAAKAAVET
148 - 166
1800.909 Da
AcrA Monomer
AcrA Dimer
Relative Uptake (Da)
Exposure Time (minutes)
ACRA_ECOLI
AAVTAAKAAVE
155 - 165
1001.563 Da
AcrA Monomer
AcrA Dimer
Relative Uptake (Da)
Exposure Time (minutes)

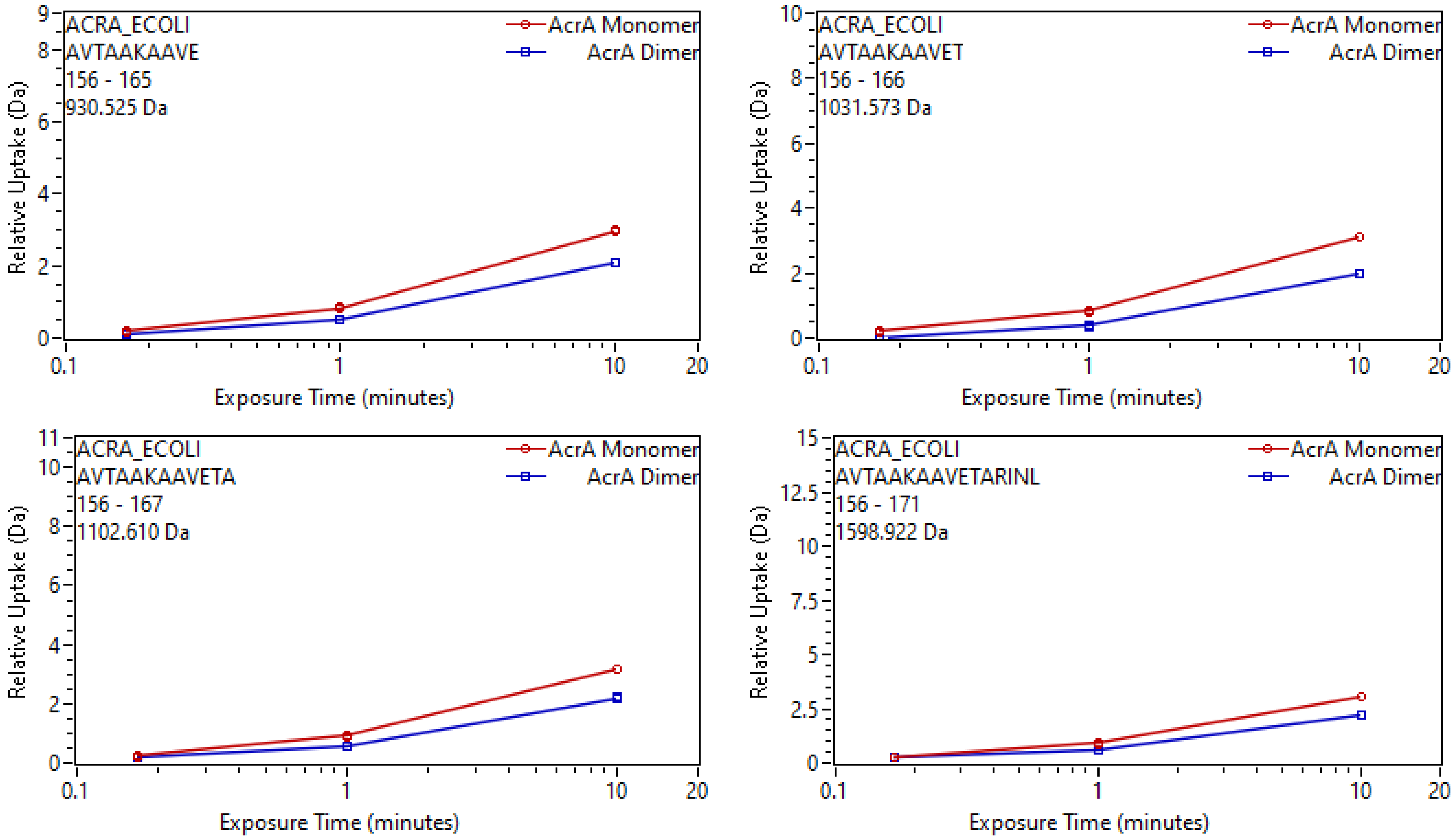
ACRA_ECOLI
AVTAAKAAVE
156 - 165
930.525 Da
AcrA Monomer
AcrA Dimer
Relative Uptake (Da)
Exposure Time (minutes)
ACRA_ECOLI
AVTAAKAAVET
156 - 166
1031.573 Da
AcrA Monomer
AcrA Dimer
Relative Uptake (Da)
Exposure Time (minutes)
ACRA_ECOLI
AVTAAKAAVETA
156 - 167
1102.610 Da
AcrA Monomer
AcrA Dimer
Relative Uptake (Da)
Exposure Time (minutes)
ACRA_ECOLI
AVTAAKAAVETARINL
156 - 171
1598.922 Da
AcrA Monomer
AcrA Dimer
Relative Uptake (Da)
Exposure Time (minutes)

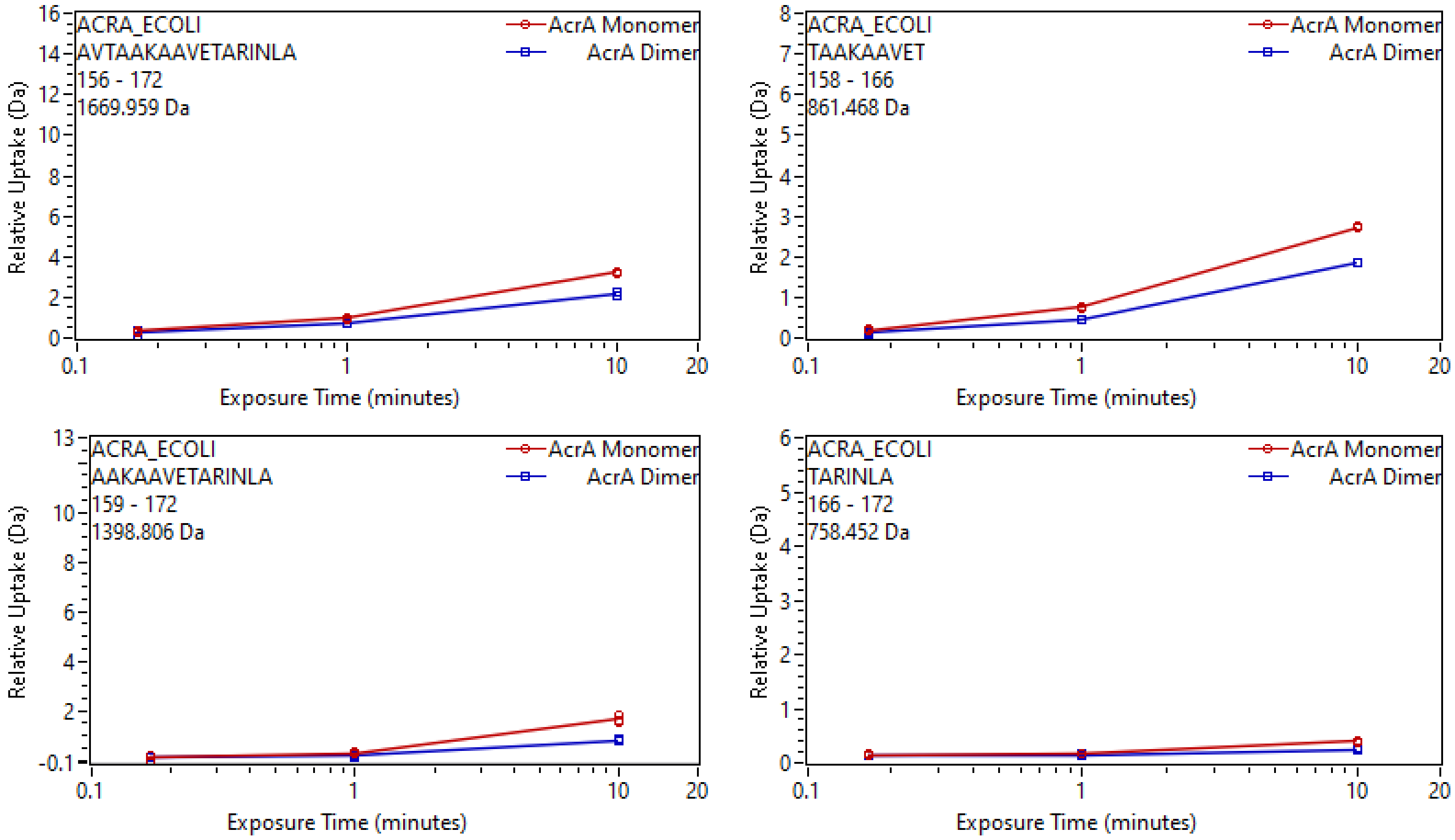
ACRA_ECOLI
AVTAAKAAVETARINLA
156 - 172
1669.959 Da
AcrA Monomer
AcrA Dimer
Relative Uptake (Da)
Exposure Time (minutes)
ACRA_ECOLI
TAAKAAVET
158 - 166
861.468 Da
AcrA Monomer
AcrA Dimer
Relative Uptake (Da)
Exposure Time (minutes)
ACRA_ECOLI
AAKAAVETARINLA
159 - 172
1398.806 Da
AcrA Monomer
AcrA Dimer
Relative Uptake (Da)
Exposure Time (minutes)
ACRA_ECOLI
TARINLA
166 - 172
758.452 Da
AcrA Monomer
AcrA Dimer
Relative Uptake (Da)
Exposure Time (minutes)

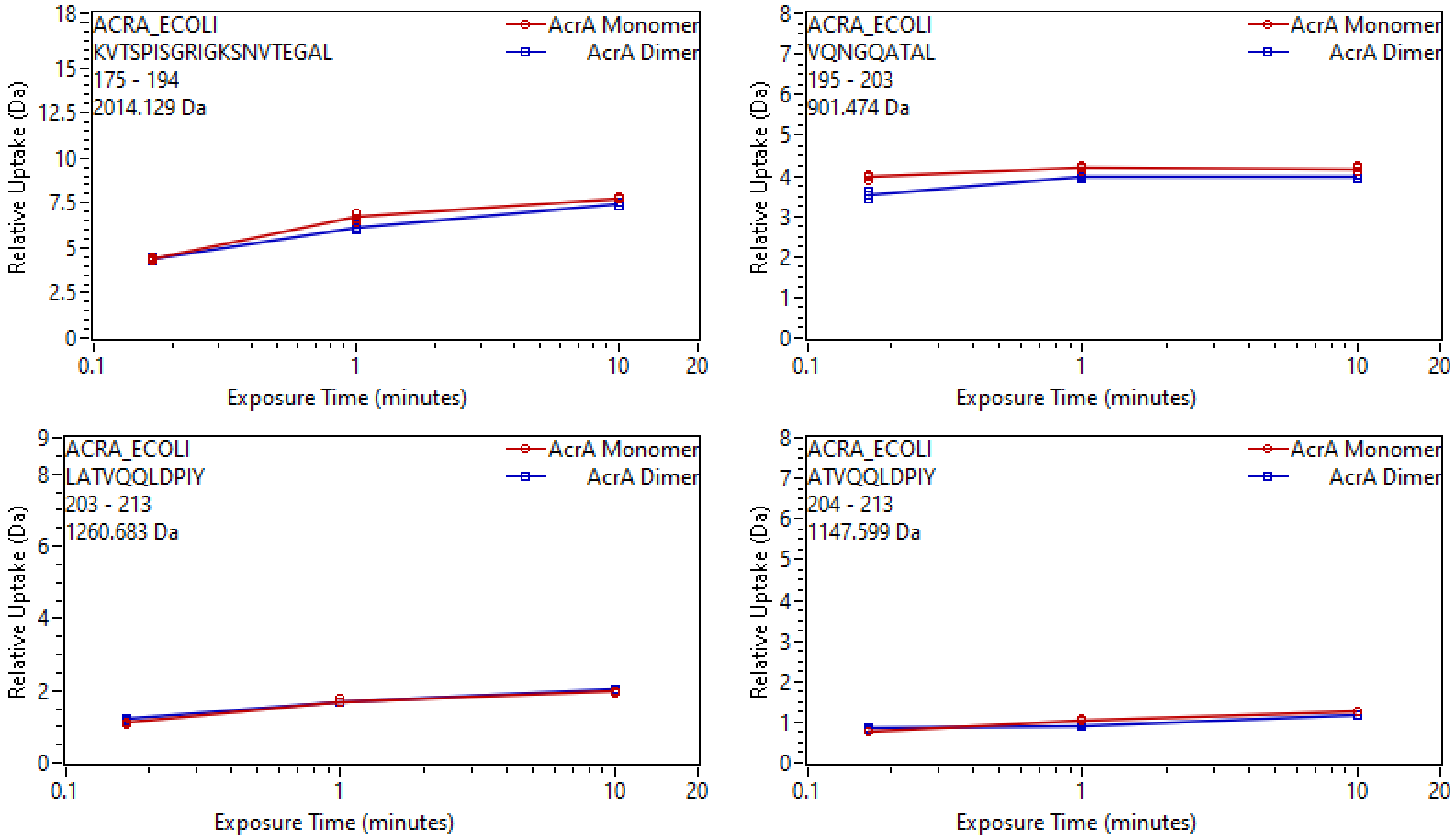
ACRA_ECOLI
KVTSPISGRIGKSNVTEGAL
175 - 194
2014.129 Da
AcrA Monomer
AcrA Dimer
Relative Uptake (Da)
Exposure Time (minutes)
ACRA_ECOLI
VQNGQATAL
195 - 203
901.474 Da
ACRA_ECOLI
LATVQQLDPIY
203 - 213
1260.683 Da
ACRA_ECOLI
ATVQQLDPIY
204 - 213
1147.599 Da

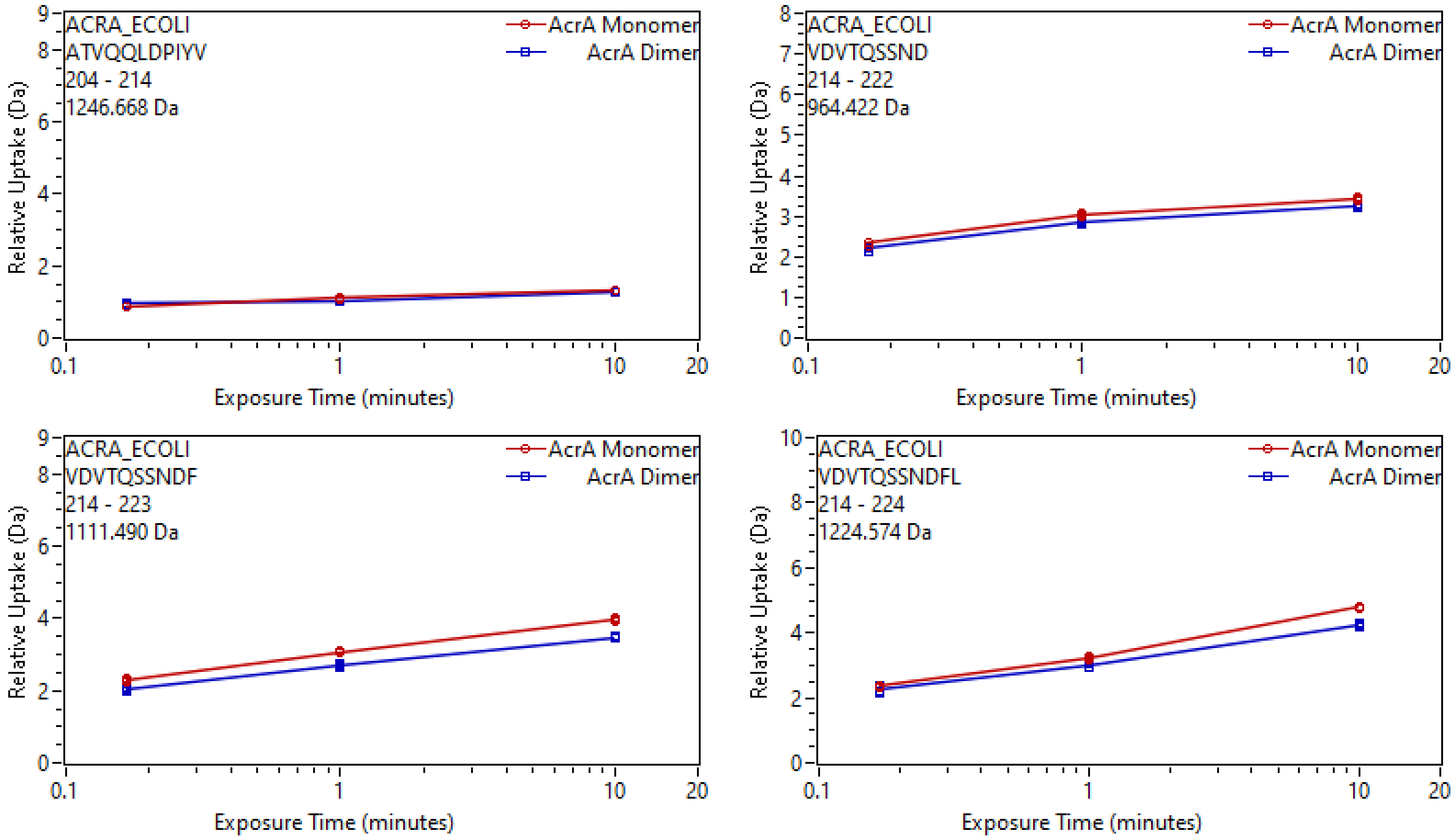
ACRA_ECOLI
ATVQQLDPIYV
204 - 214
1246.668 Da
AcrA Monomer
AcrA Dimer
Relative Uptake (Da)
Exposure Time (minutes)
ACRA_ECOLI
VDVTQSSND
214 - 222
964.422 Da
AcrA Monomer
AcrA Dimer
Relative Uptake (Da)
Exposure Time (minutes)
ACRA_ECOLI
VDVTQSSNDF
214 - 223
1111.490 Da
AcrA Monomer
AcrA Dimer
Relative Uptake (Da)
Exposure Time (minutes)
ACRA_ECOLI
VDVTQSSNDFL
214 - 224
1224.574 Da
AcrA Monomer
AcrA Dimer
Relative Uptake (Da)
Exposure Time (minutes)

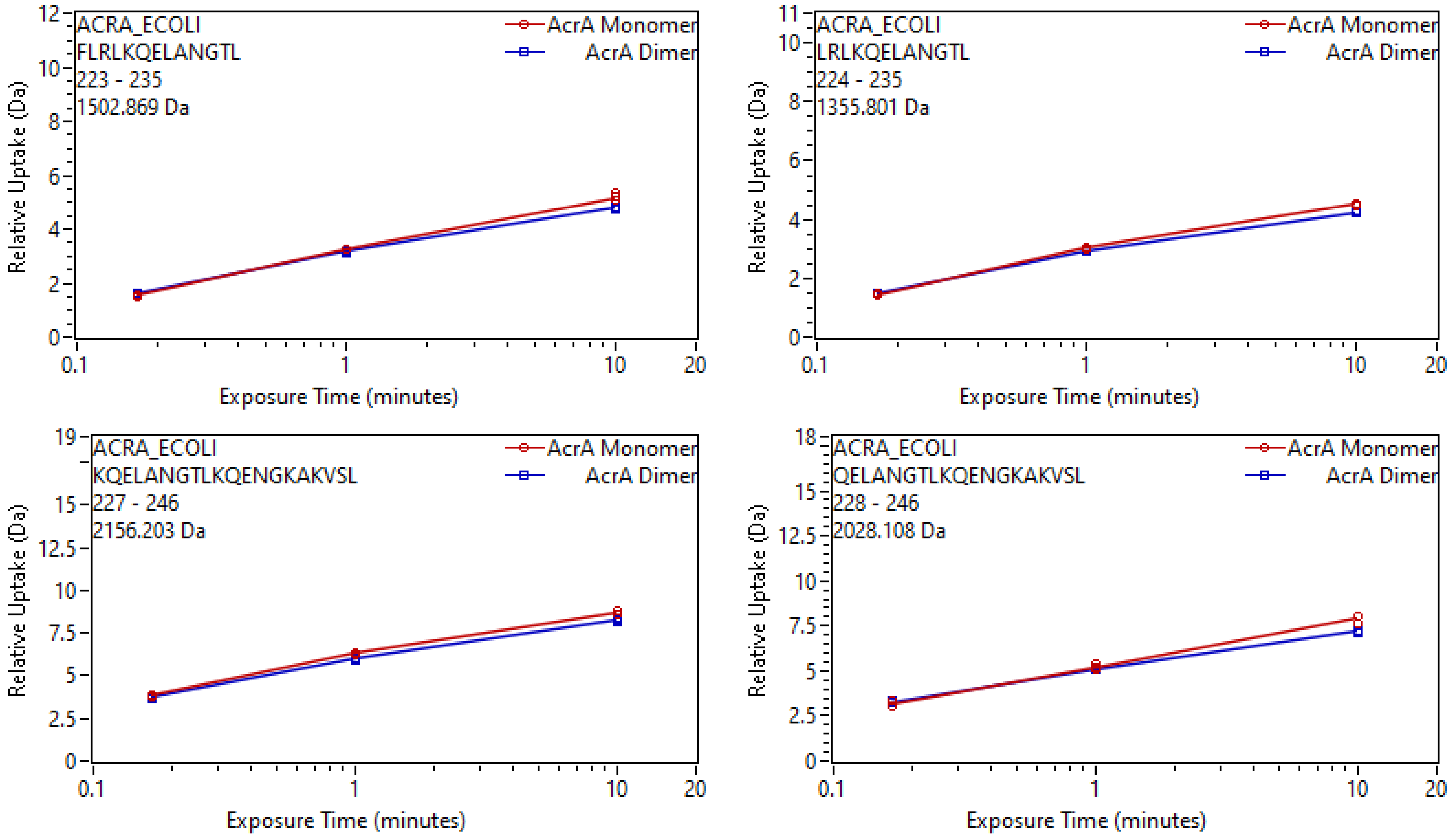
ACRA_ECOLI
FLRLKQELANGTL
223 - 235
1502.869 Da
AcrA Monomer
AcrA Dimer
Relative Uptake (Da)
Exposure Time (minutes)
0
2
4
6
8
10
12
0.1
1
10
20
ACRA_ECOLI
LRLKQELANGTL
224 - 235
1355.801 Da
AcrA Monomer
AcrA Dimer
Relative Uptake (Da)
Exposure Time (minutes)
0
2
4
6
8
10
11
0.1
1
10
20
ACRA_ECOLI
KQELANGTLKQENGKAKVSL
227 - 246
2156.203 Da
AcrA Monomer
AcrA Dimer
Relative Uptake (Da)
Exposure Time (minutes)
0
2.5
5
7.5
10
12.5
15
19
0.1
1
10
20
ACRA_ECOLI
QELANGTLKQENGKAKVSL
228 - 246
2028.108 Da
AcrA Monomer
AcrA Dimer
Relative Uptake (Da)
Exposure Time (minutes)
0
2.5
5
7.5
10
12.5
15
18
0.1
1
10
20

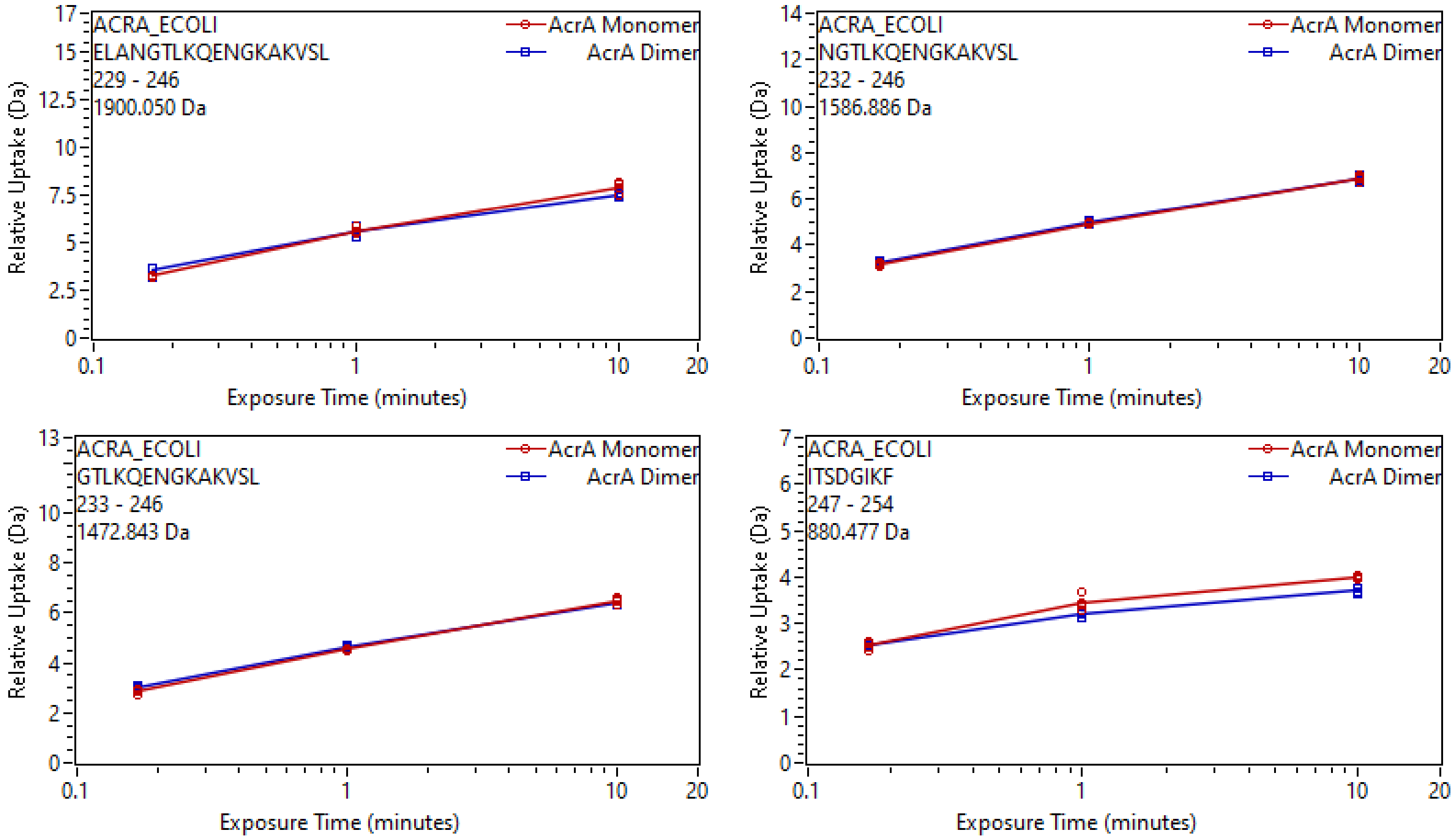
ACRA_ECOLI
ELANGTLKQENGKAKVSL
229 - 246
1900.050 Da
AcrA Monomer
AcrA Dimer
Relative Uptake (Da)
Exposure Time (minutes)
ACRA_ECOLI
NGTLKQENGKAKVSL
232 - 246
1586.886 Da
ACRA_ECOLI
GTLKQENGKAKVSL
233 - 246
1472.843 Da
ACRA_ECOLI
ITSDGIKF
247 - 254
880.477 Da

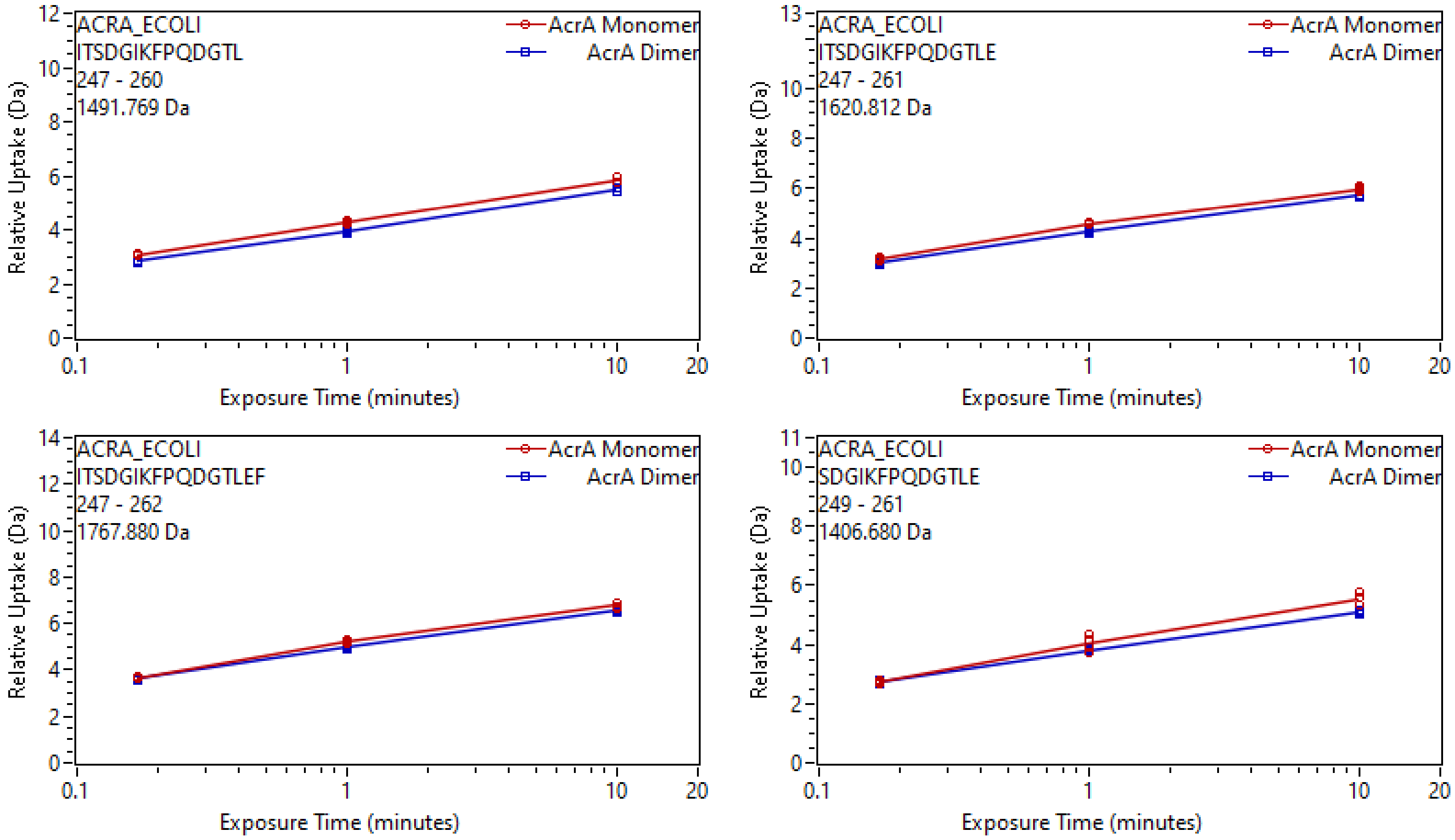
ACRA_ECOLI
ITSDGIKFPQDGTL
247 - 260
1491.769 Da
AcrA Monomer
AcrA Dimer
Relative Uptake (Da)
Exposure Time (minutes)
ACRA_ECOLI
ITSDGIKFPQDGTLE
247 - 261
1620.812 Da
AcrA Monomer
AcrA Dimer
Relative Uptake (Da)
Exposure Time (minutes)
ACRA_ECOLI
ITSDGIKFPQDGTLEF
247 - 262
1767.880 Da
AcrA Monomer
AcrA Dimer
Relative Uptake (Da)
Exposure Time (minutes)
ACRA_ECOLI
SDGIKFPQDGTLE
249 - 261
1406.680 Da
AcrA Monomer
AcrA Dimer
Relative Uptake (Da)
Exposure Time (minutes)

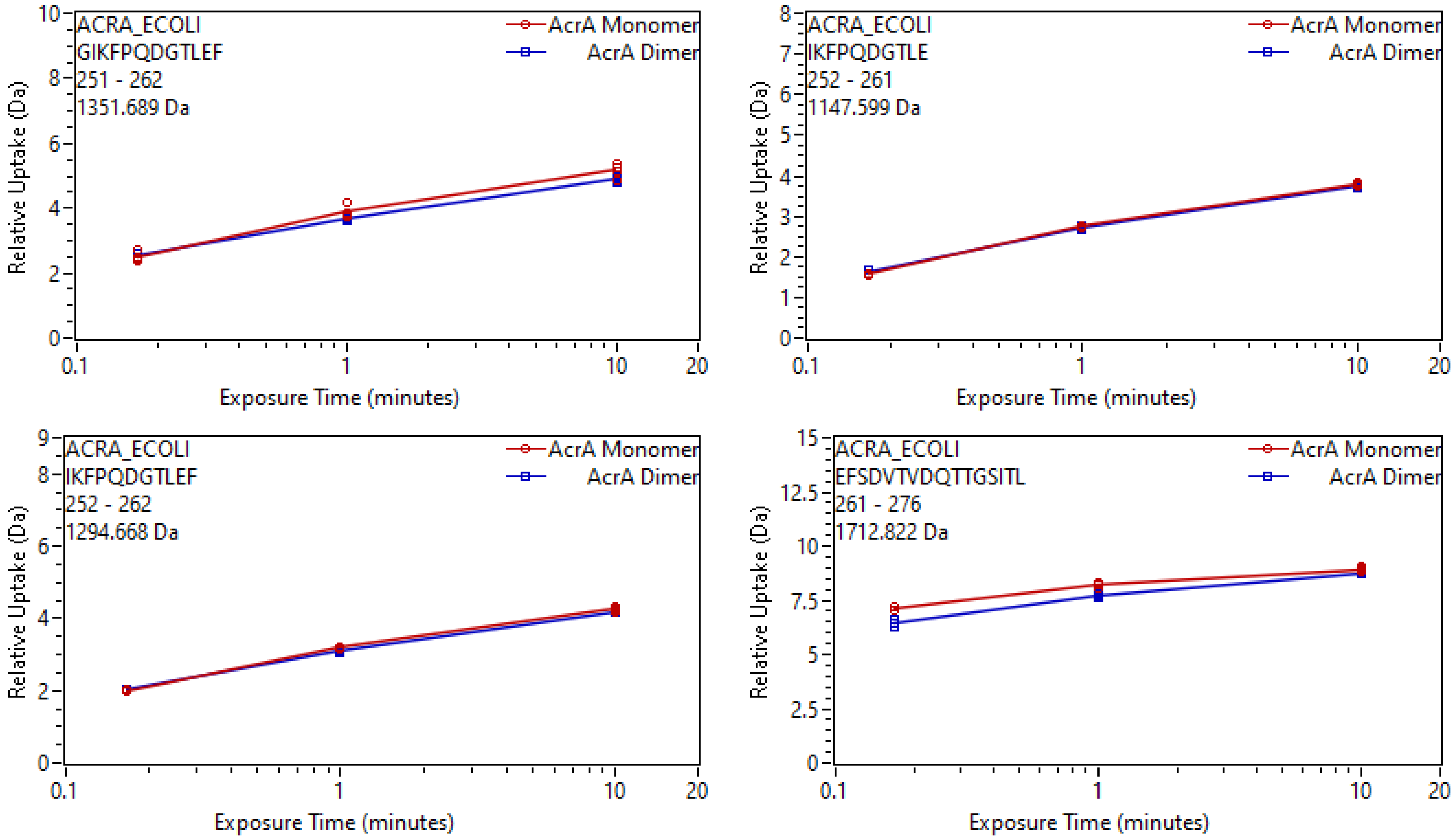
ACRA_ECOLI
GIKFPQDGTLEF
251 - 262
1351.689 Da
AcrA Monomer
AcrA Dimer
Relative Uptake (Da)
Exposure Time (minutes)
ACRA_ECOLI
IKFPQDGTLE
252 - 261
1147.599 Da
AcrA Monomer
AcrA Dimer
Relative Uptake (Da)
Exposure Time (minutes)
ACRA_ECOLI
IKFPQDGTLEF
252 - 262
1294.668 Da
AcrA Monomer
AcrA Dimer
Relative Uptake (Da)
Exposure Time (minutes)
ACRA_ECOLI
EFSDVTVDQTTGSITL
261 - 276
1712.822 Da
AcrA Monomer
AcrA Dimer
Relative Uptake (Da)
Exposure Time (minutes)

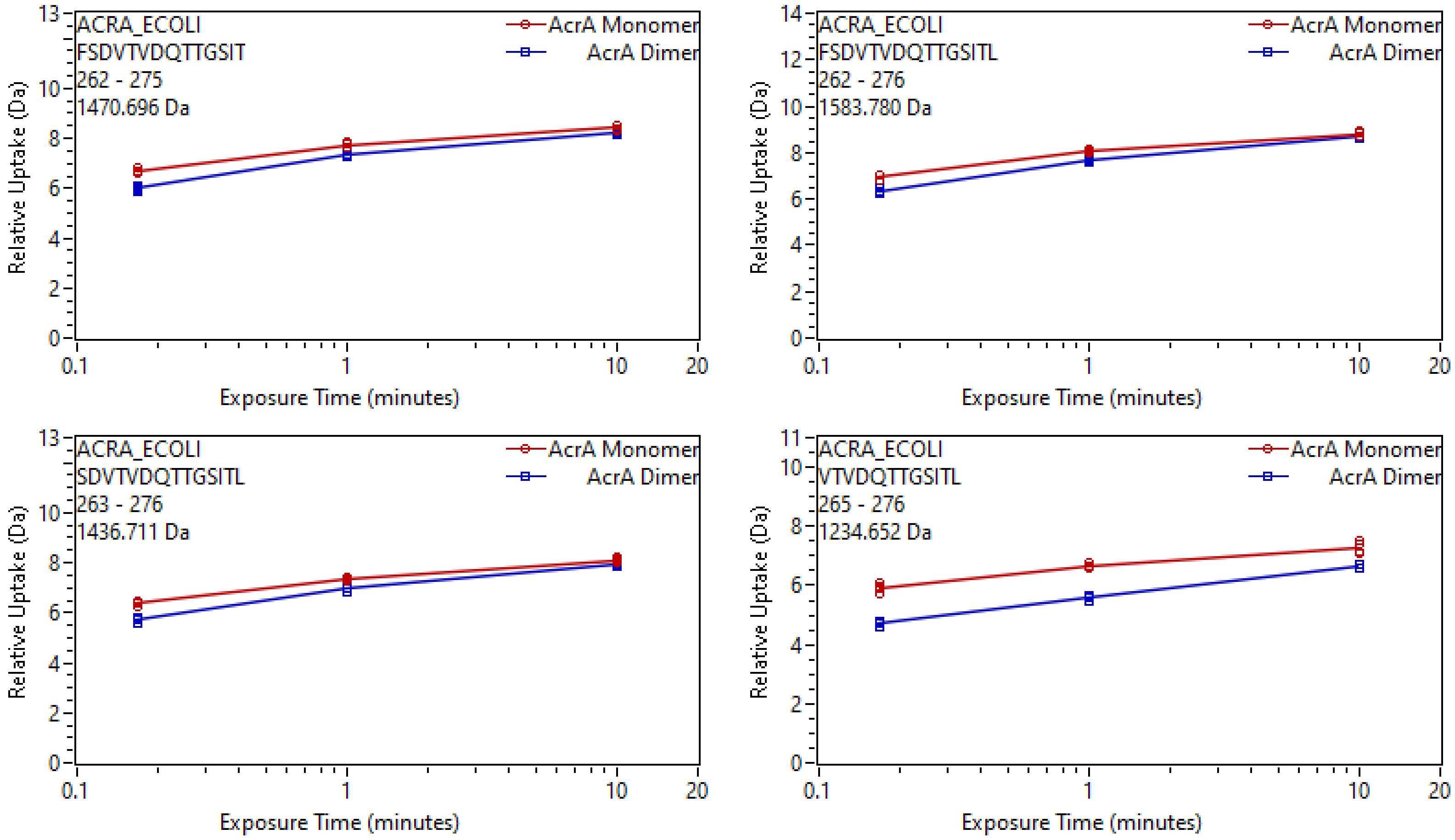
ACRA_ECOLI
FSDVTVDQTTGSIT
262 - 275
1470.696 Da
AcrA Monomer
AcrA Dimer
ACRA_ECOLI
FSDVTVDQTTGSITL
262 - 276
1583.780 Da
ACRA_ECOLI
SDVTVDQTTGSITL
263 - 276
1436.711 Da
ACRA_ECOLI
VTVDQTTGSITL
265 - 276
1234.652 Da
Relative Uptake (Da)
Exposure Time (minutes)
0.1
1
10
20

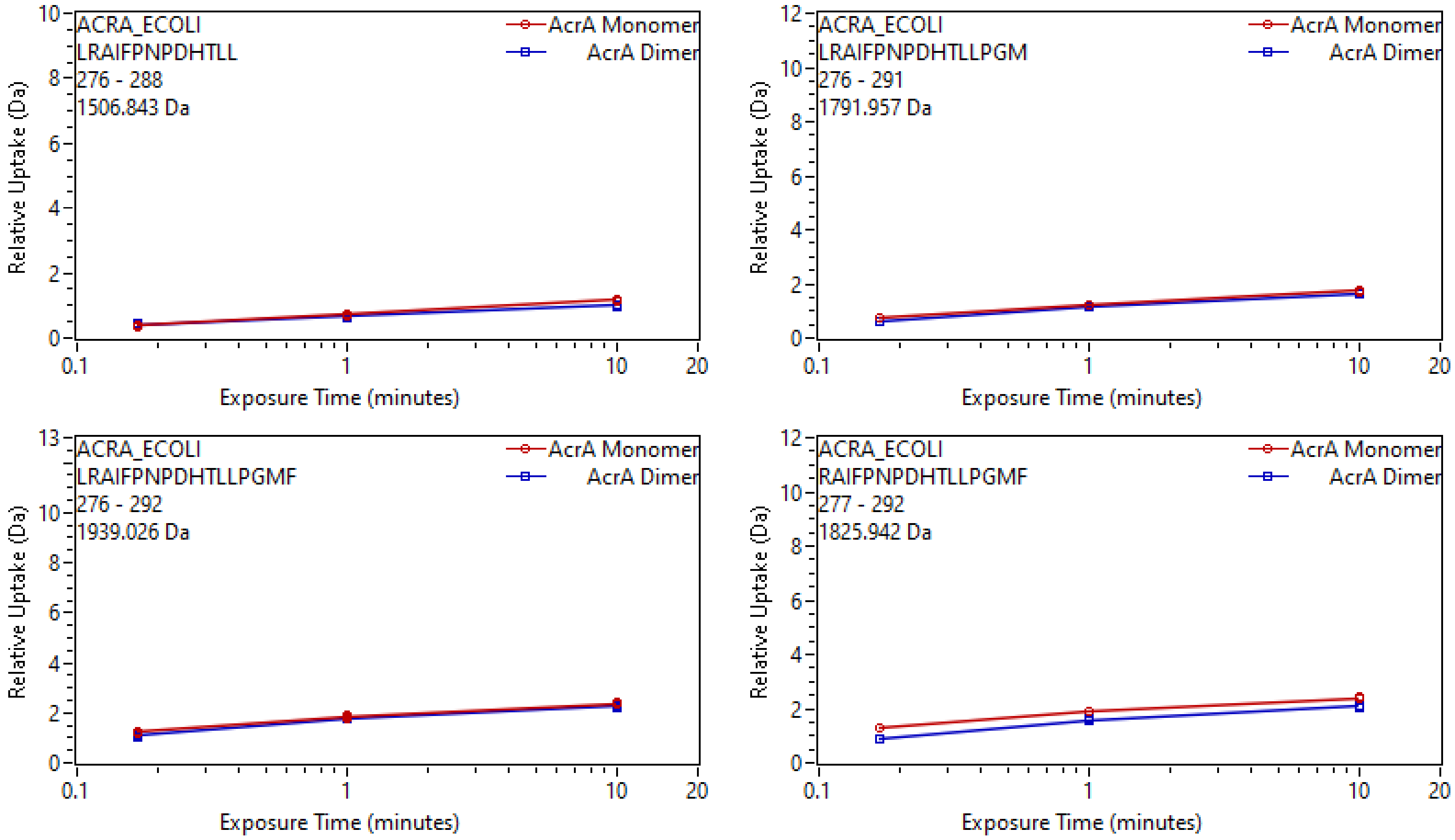
ACRA_ECOLI
LRAIFPNPDHTLL
276 - 288
1506.843 Da
AcrA Monomer
AcrA Dimer
Relative Uptake (Da)
Exposure Time (minutes)
ACRA_ECOLI
LRAIFPNPDHTLLPGM
276 - 291
1791.957 Da
AcrA Monomer
AcrA Dimer
Relative Uptake (Da)
Exposure Time (minutes)
ACRA_ECOLI
LRAIFPNPDHTLLPGMF
276 - 292
1939.026 Da
AcrA Monomer
AcrA Dimer
Relative Uptake (Da)
Exposure Time (minutes)
ACRA_ECOLI
RAIFPNPDHTLLPGMF
277 - 292
1825.942 Da
AcrA Monomer
AcrA Dimer
Relative Uptake (Da)
Exposure Time (minutes)

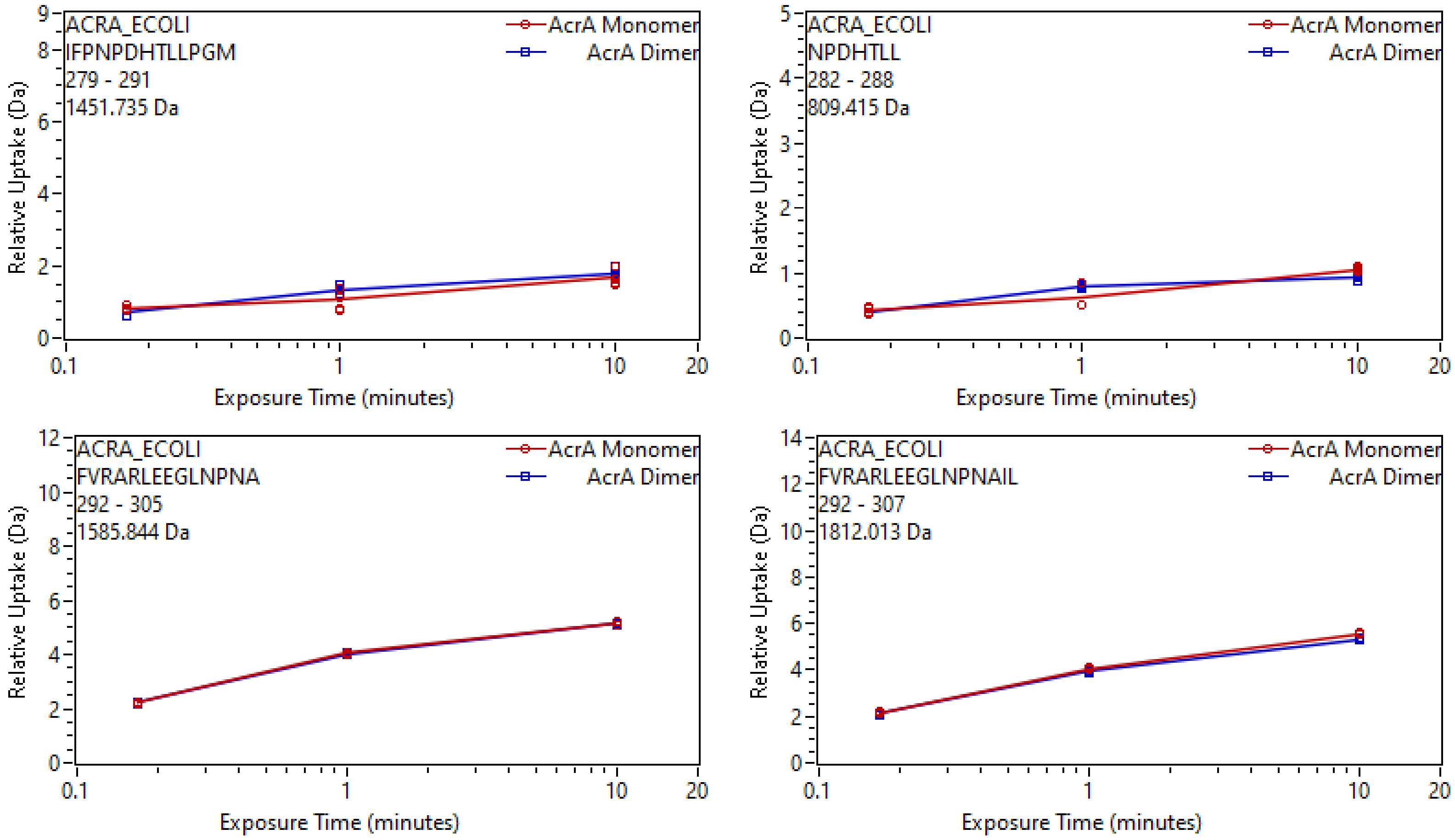
ACRA_ECOLI
IFPNPDHTLLPGM
279 - 291
1451.735 Da
AcrA Monomer
AcrA Dimer
Relative Uptake (Da)
Exposure Time (minutes)
ACRA_ECOLI
NPDHTLL
282 - 288
809.415 Da
ACRA_ECOLI
FVRARLEEGLNPNA
292 - 305
1585.844 Da
ACRA_ECOLI
FVRARLEEGLNPNAIL
292 - 307
1812.013 Da

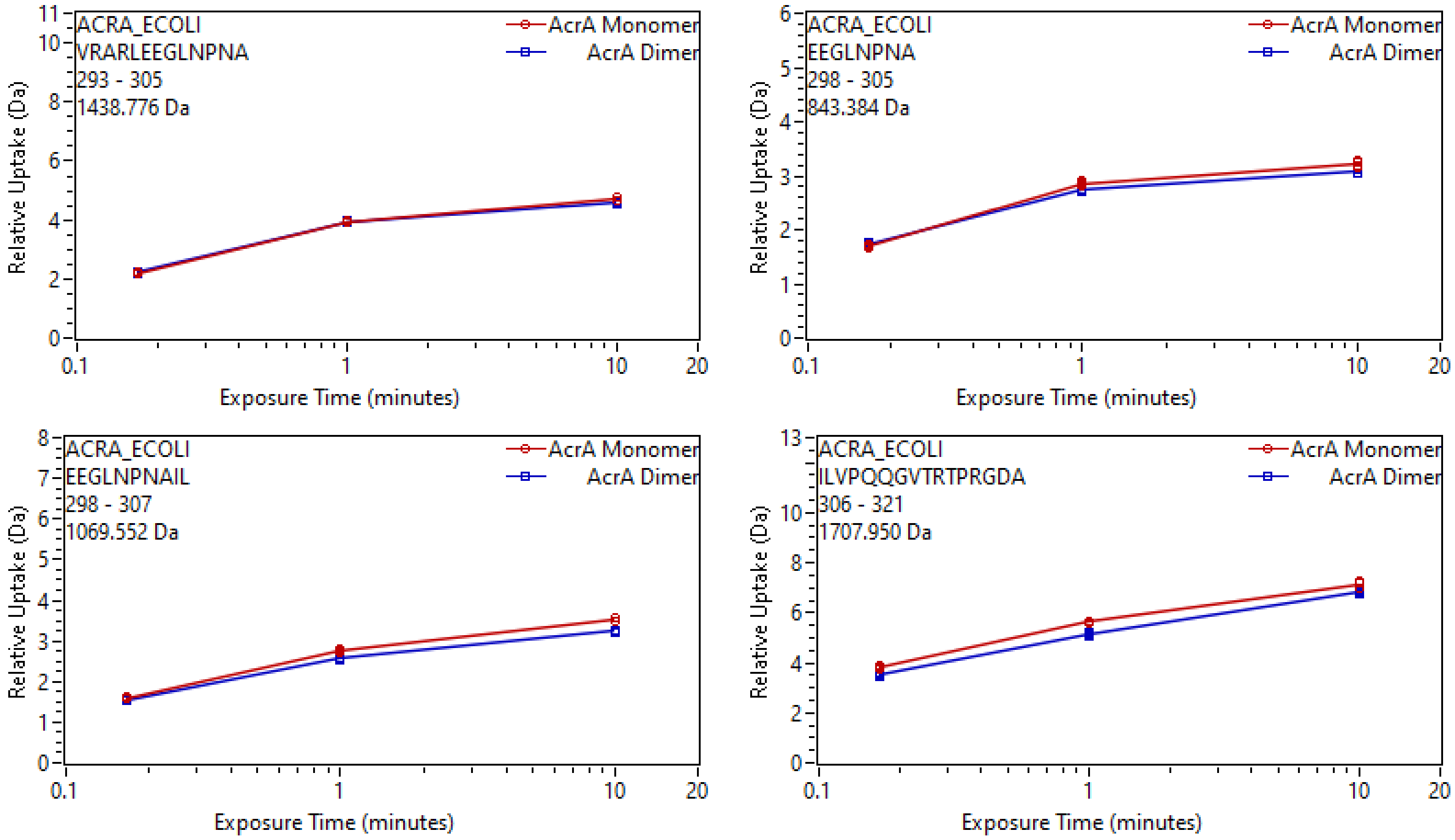
ACRA_ECOLI
VRARLEEGLNPNA
293 - 305
1438.776 Da
AcrA Monomer
AcrA Dimer
Relative Uptake (Da)
Exposure Time (minutes)
ACRA_ECOLI
EEGLNPNA
298 - 305
843.384 Da
ACRA_ECOLI
EEGLNPNAIL
298 - 307
1069.552 Da
ACRA_ECOLI
ILVPQQGVTRTPRGDA
306 - 321
1707.950 Da

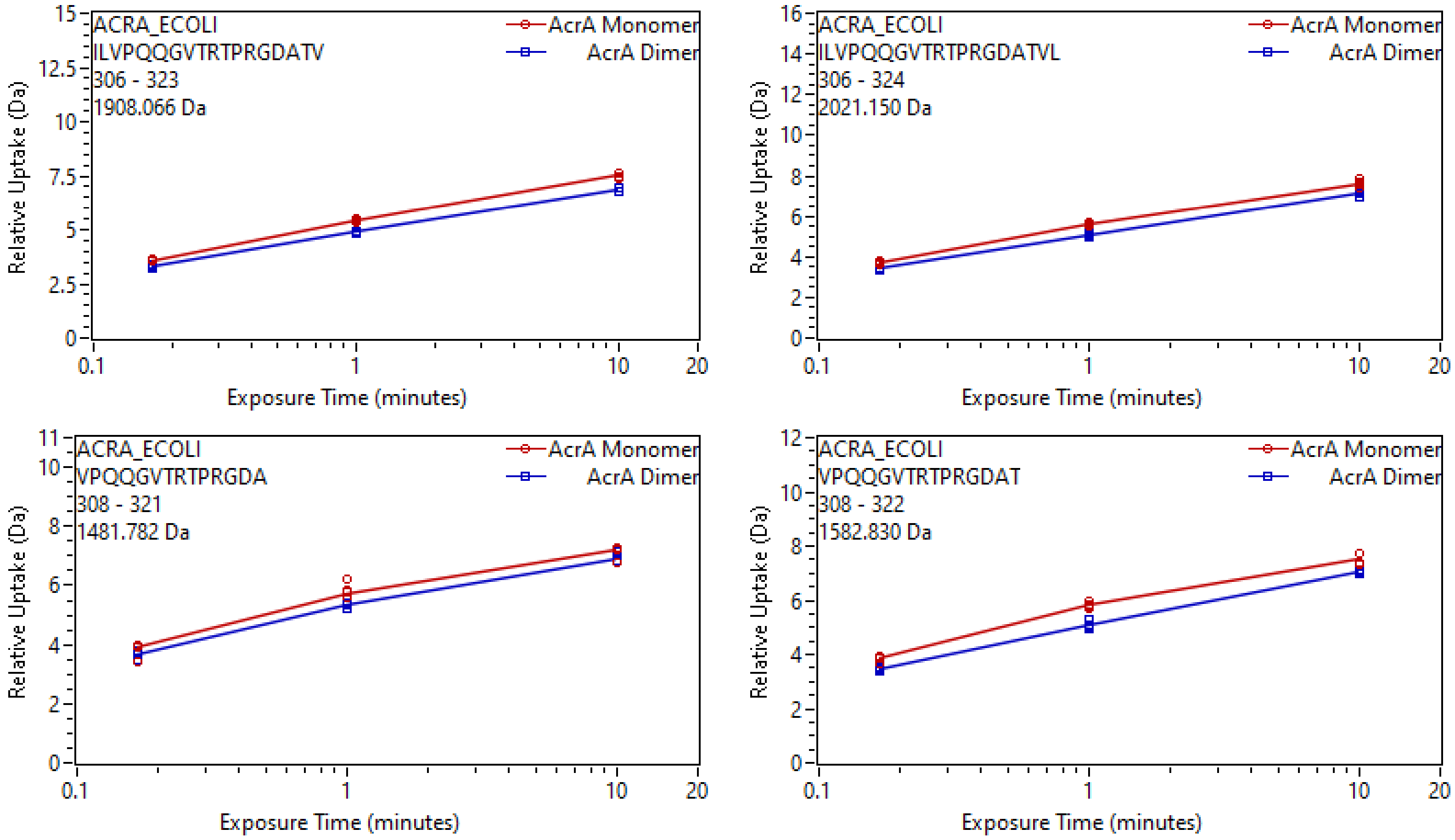
ACRA_ECOLI
ILVPQQGVTRTPRGDATV
306 - 323
1908.066 Da
AcrA Monomer
AcrA Dimer
Relative Uptake (Da)
Exposure Time (minutes)
ACRA_ECOLI
ILVPQQGVTRTPRGDATVL
306 - 324
2021.150 Da
AcrA Monomer
AcrA Dimer
Relative Uptake (Da)
Exposure Time (minutes)
ACRA_ECOLI
VPQQGVTRTPRGDA
308 - 321
1481.782 Da
AcrA Monomer
AcrA Dimer
Relative Uptake (Da)
Exposure Time (minutes)
ACRA_ECOLI
VPQQGVTRTPRGDAT
308 - 322
1582.830 Da
AcrA Monomer
AcrA Dimer
Relative Uptake (Da)
Exposure Time (minutes)

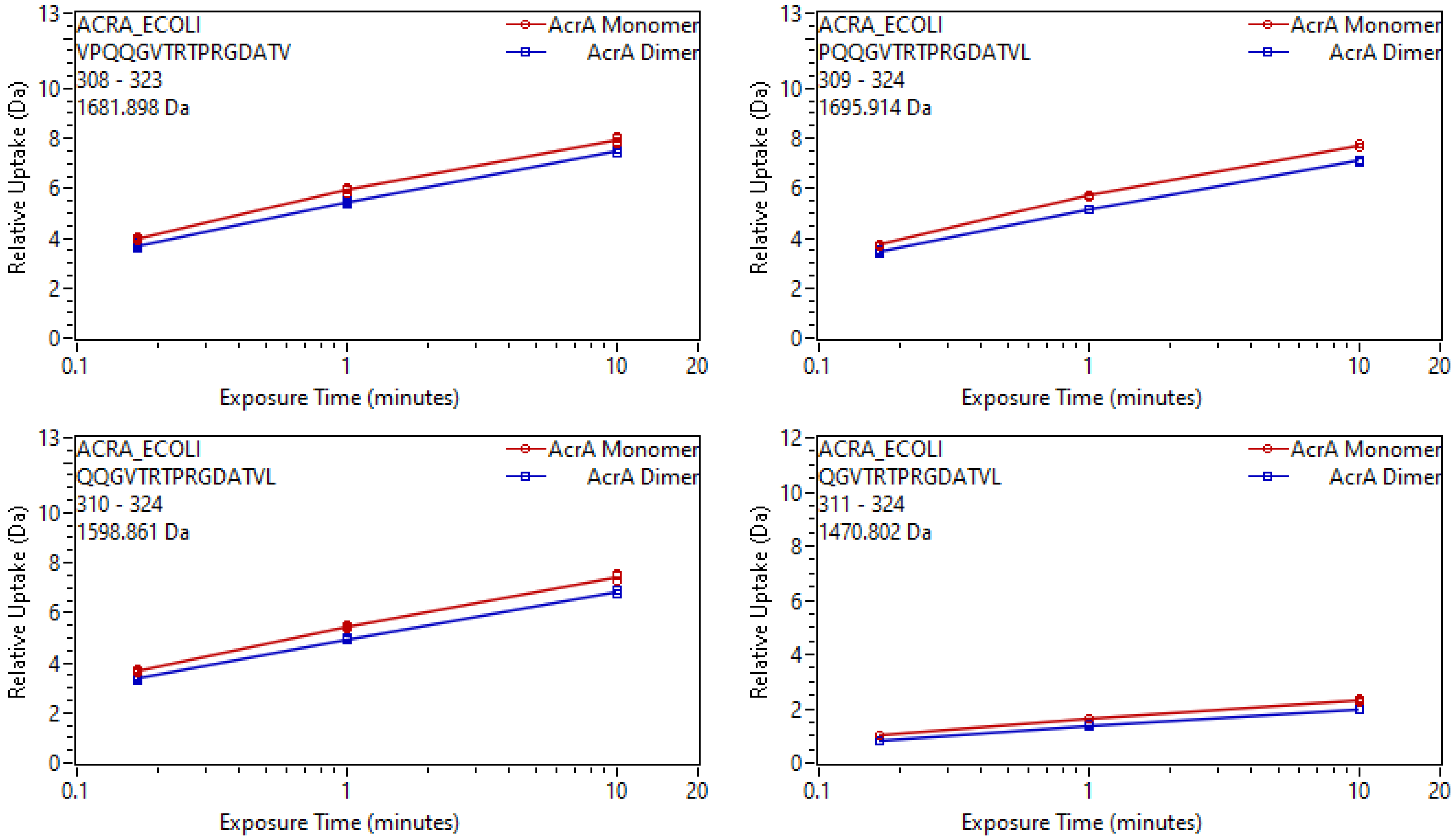
ACRA_ECOLI
VPQQGVTRTPRGDATV
308 - 323
1681.898 Da
AcrA Monomer
AcrA Dimer
Relative Uptake (Da)
Exposure Time (minutes)
ACRA_ECOLI
PQQGVTRTPRGDATVL
309 - 324
1695.914 Da
AcrA Monomer
AcrA Dimer
Relative Uptake (Da)
Exposure Time (minutes)
ACRA_ECOLI
QQGVTRTPRGDATVL
310 - 324
1598.861 Da
AcrA Monomer
AcrA Dimer
Relative Uptake (Da)
Exposure Time (minutes)
ACRA_ECOLI
QGVTRTPRGDATVL
311 - 324
1470.802 Da
AcrA Monomer
AcrA Dimer
Relative Uptake (Da)
Exposure Time (minutes)

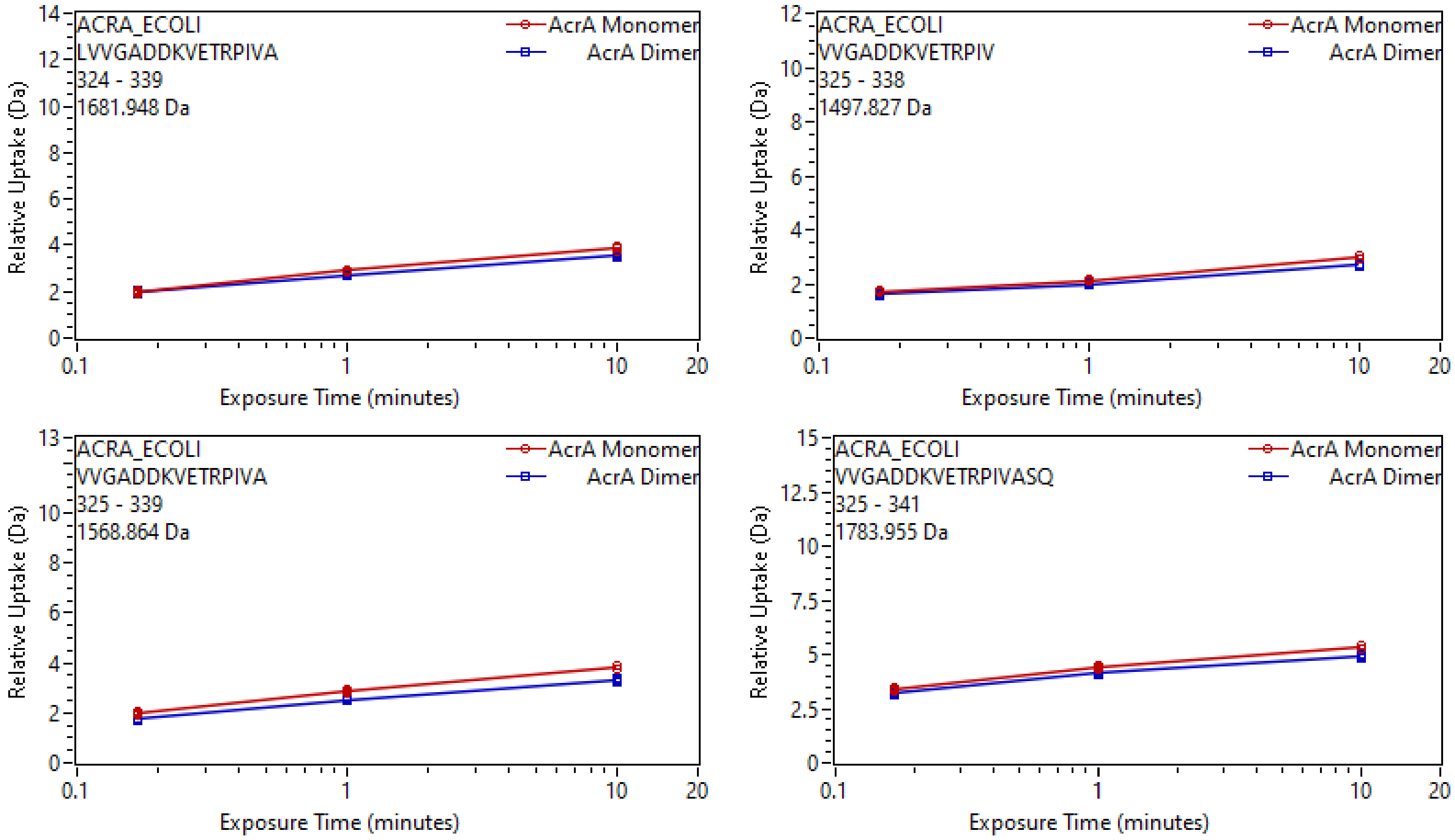
ACRA_ECOLI
LVVGADDKVETRPIVA
324 - 339
1681.948 Da
AcrA Monomer
AcrA Dimer
Relative Uptake (Da)
Exposure Time (minutes)
ACRA_ECOLI
VVGADDKVETRPIV
325 - 338
1497.827 Da
AcrA Monomer
AcrA Dimer
Relative Uptake (Da)
Exposure Time (minutes)
ACRA_ECOLI
VVGADDKVETRPIVA
325 - 339
1568.864 Da
AcrA Monomer
AcrA Dimer
Relative Uptake (Da)
Exposure Time (minutes)
ACRA_ECOLI
VVGADDKVETRPIVASQ
325 - 341
1783.955 Da
AcrA Monomer
AcrA Dimer
Relative Uptake (Da)
Exposure Time (minutes)

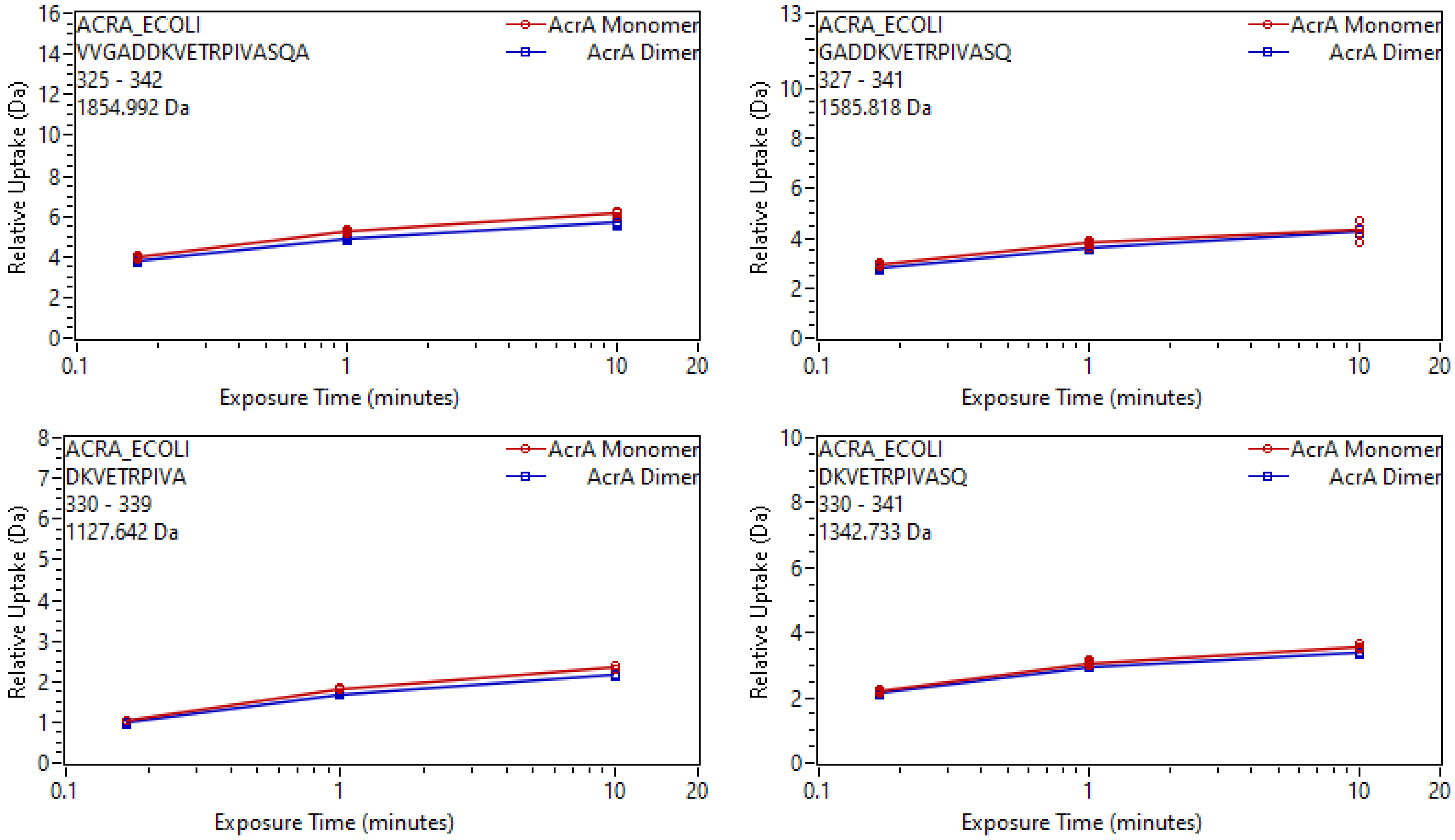
ACRA_ECOLI
VVGADDKVETRPIVASQA
325 - 342
1854.992 Da
AcrA Monomer
AcrA Dimer
Relative Uptake (Da)
Exposure Time (minutes)
ACRA_ECOLI
GADDKVETRPIVASQ
327 - 341
1585.818 Da
AcrA Monomer
AcrA Dimer
Relative Uptake (Da)
Exposure Time (minutes)
ACRA_ECOLI
DKVETRPIVA
330 - 339
1127.642 Da
AcrA Monomer
AcrA Dimer
Relative Uptake (Da)
Exposure Time (minutes)
ACRA_ECOLI
DKVETRPIVASQ
330 - 341
1342.733 Da
AcrA Monomer
AcrA Dimer
Relative Uptake (Da)
Exposure Time (minutes)

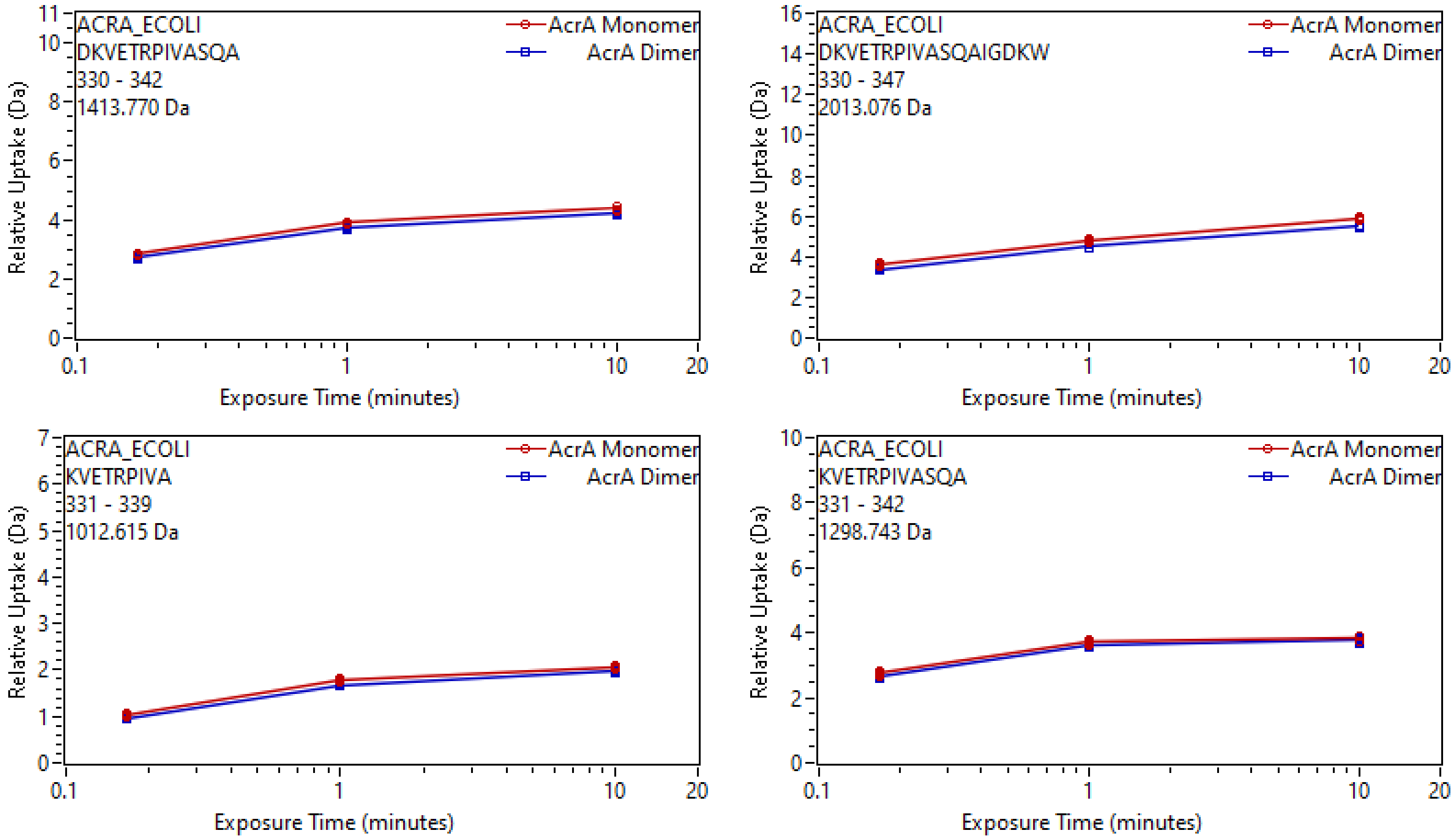
ACRA_ECOLI
DKVETRPIVASQA
330 - 342
1413.770 Da
AcrA Monomer
AcrA Dimer
Relative Uptake (Da)
Exposure Time (minutes)
ACRA_ECOLI
DKVETRPIVASQAIGDKW
330 - 347
2013.076 Da
AcrA Monomer
AcrA Dimer
Relative Uptake (Da)
Exposure Time (minutes)
ACRA_ECOLI
KVETRPIVA
331 - 339
1012.615 Da
AcrA Monomer
AcrA Dimer
Relative Uptake (Da)
Exposure Time (minutes)
ACRA_ECOLI
KVETRPIVASQA
331 - 342
1298.743 Da
AcrA Monomer
AcrA Dimer
Relative Uptake (Da)
Exposure Time (minutes)

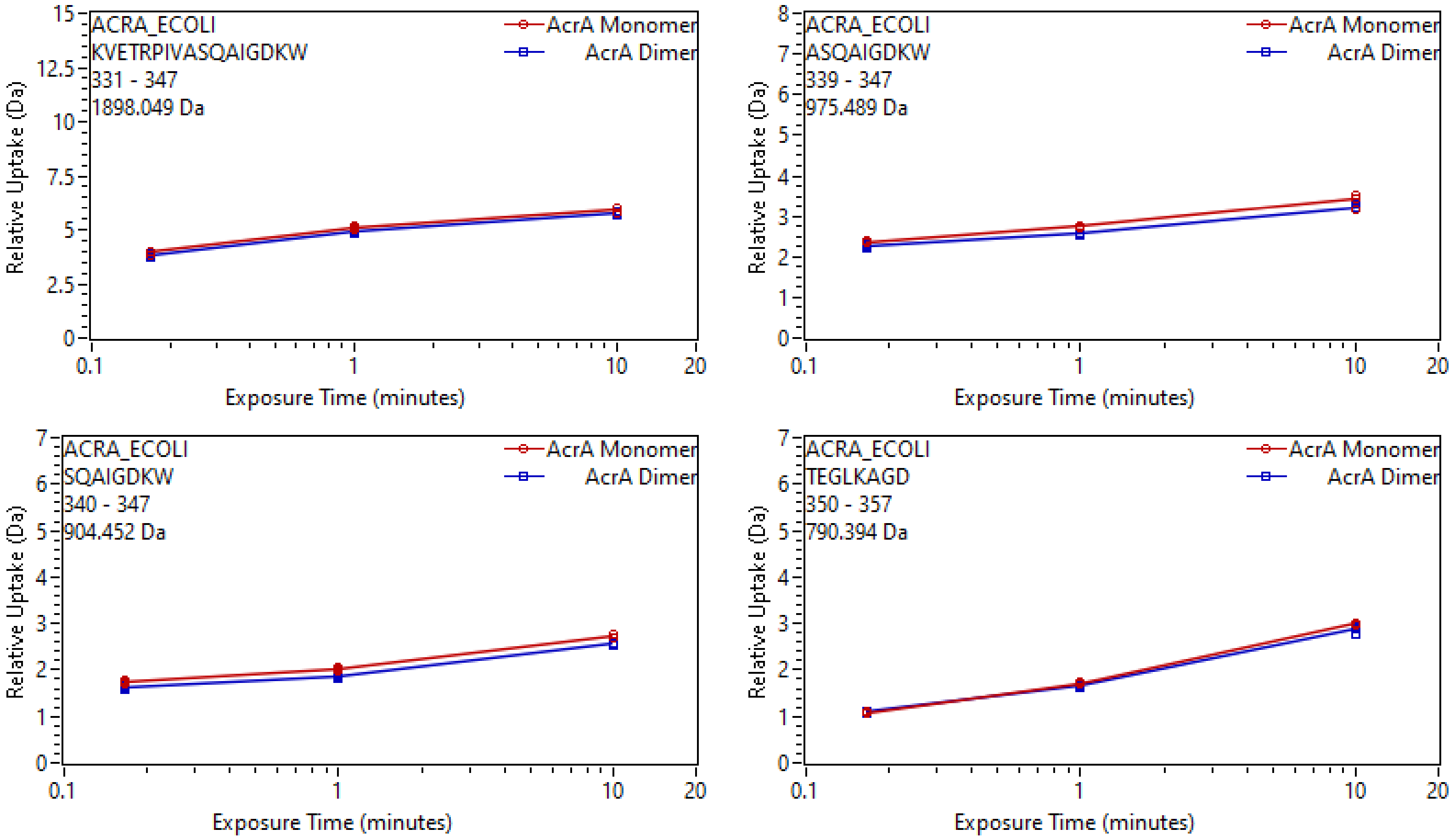
ACRA_ECOLI
KVETRPIVASQAIGDKW
331 - 347
1898.049 Da
AcrA Monomer
AcrA Dimer
Relative Uptake (Da)
Exposure Time (minutes)
ACRA_ECOLI
ASQAIGDKW
339 - 347
975.489 Da
AcrA Monomer
AcrA Dimer
Relative Uptake (Da)
Exposure Time (minutes)
ACRA_ECOLI
SQAIGDKW
340 - 347
904.452 Da
AcrA Monomer
AcrA Dimer
Relative Uptake (Da)
Exposure Time (minutes)
ACRA_ECOLI
TEGLKAGD
350 - 357
790.394 Da
AcrA Monomer
AcrA Dimer
Relative Uptake (Da)
Exposure Time (minutes)

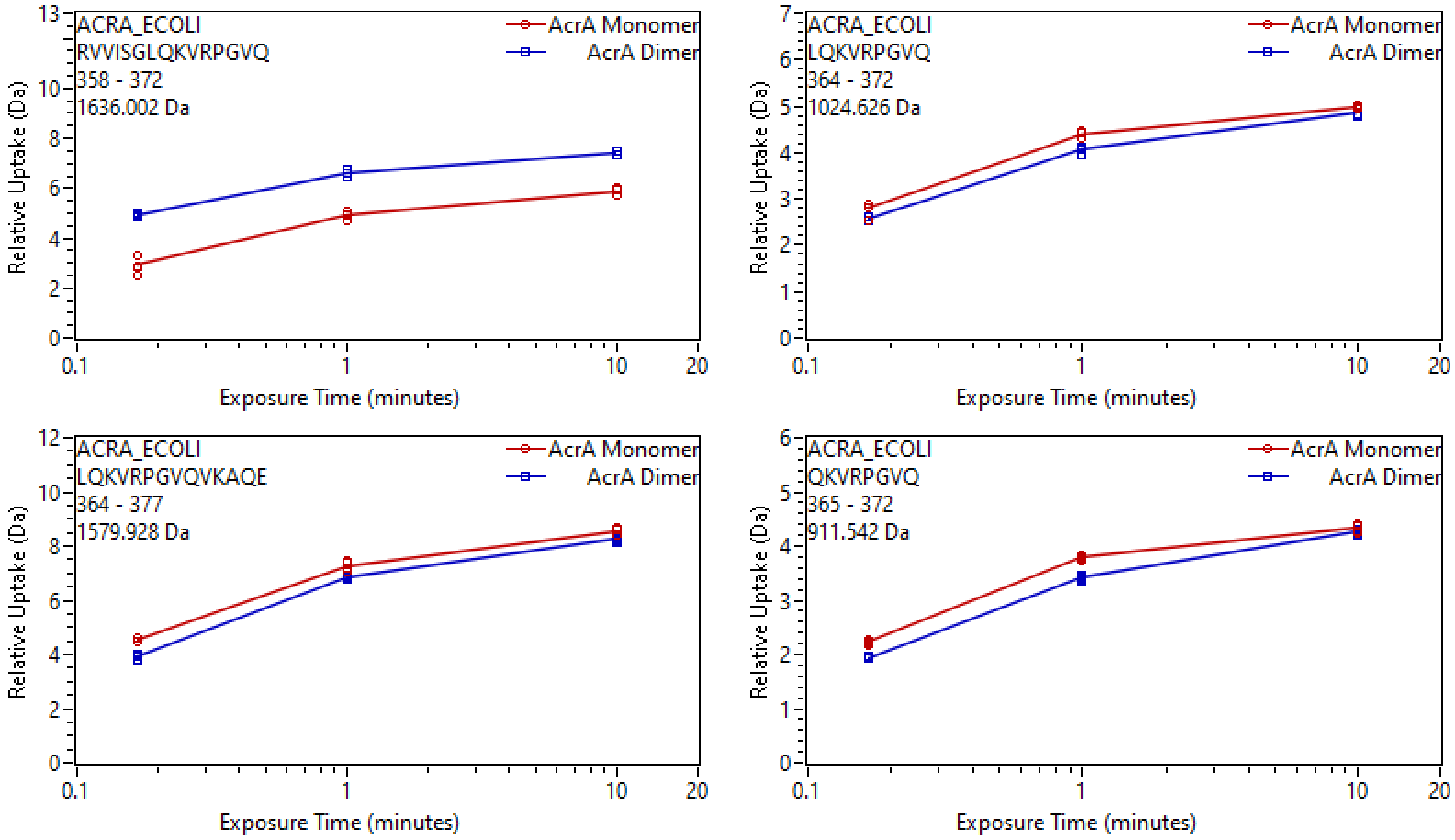
ACRA_ECOLI
RVVISGLQKVRPGVQ
358 - 372
1636.002 Da
AcrA Monomer
AcrA Dimer
Relative Uptake (Da)
Exposure Time (minutes)
ACRA_ECOLI
LQKVRPGVQ
364 - 372
1024.626 Da
AcrA Monomer
AcrA Dimer
Relative Uptake (Da)
Exposure Time (minutes)
ACRA_ECOLI
LQKVRPGVQVKAQE
364 - 377
1579.928 Da
AcrA Monomer
AcrA Dimer
Relative Uptake (Da)
Exposure Time (minutes)
ACRA_ECOLI
QKVRPGVQ
365 - 372
911.542 Da
AcrA Monomer
AcrA Dimer
Relative Uptake (Da)
Exposure Time (minutes)

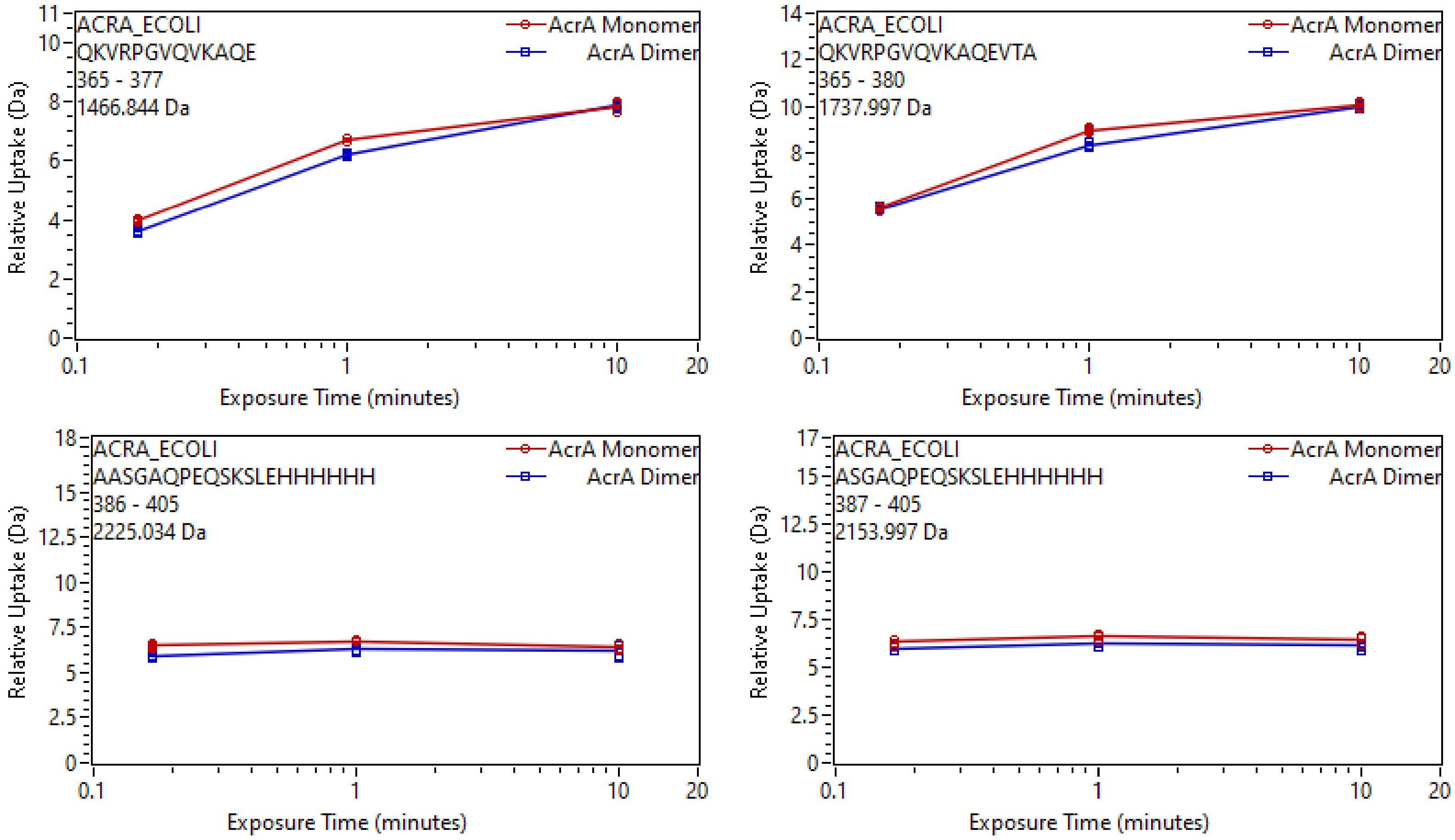

ACRA_ECOLI
QKVRPGVQVKAQE
365 - 377
1466.844 Da
AcrA Monomer
AcrA Dimer
Relative Uptake (Da)
Exposure Time (minutes)
ACRA_ECOLI
QKVRPGVQVKAQEVTA
365 - 380
1737.997 Da
AcrA Monomer
AcrA Dimer
Relative Uptake (Da)
Exposure Time (minutes)
ACRA_ECOLI
AASGAQPEQSKSLEHHHHHH
386 - 405
2225.034 Da
AcrA Monomer
AcrA Dimer
Relative Uptake (Da)
Exposure Time (minutes)
ACRA_ECOLI
ASGAQPEQSKSLEHHHHHH
387 - 405
2153.997 Da
AcrA Monomer
AcrA Dimer
Relative Uptake (Da)
Exposure Time (minutes)

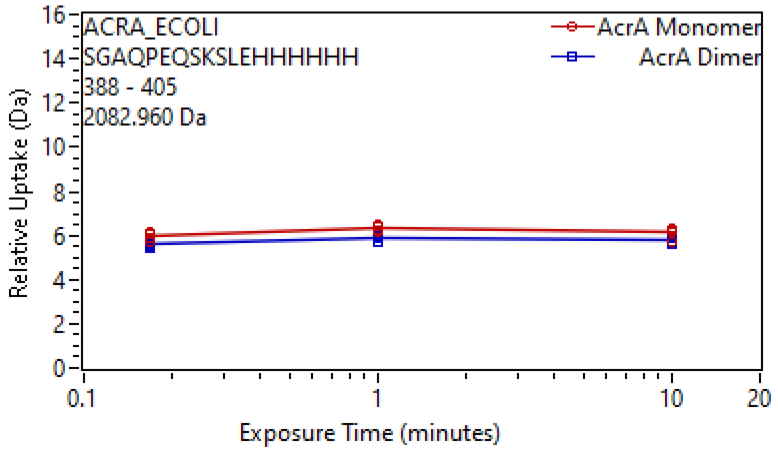
ACRA_ECOLI
SGAQPEQSKSLEHHHHHH
388 - 405
2082.960 Da
AcrA Monomer
AcrA Dimer
Relative Uptake (Da)
Exposure Time (minutes)

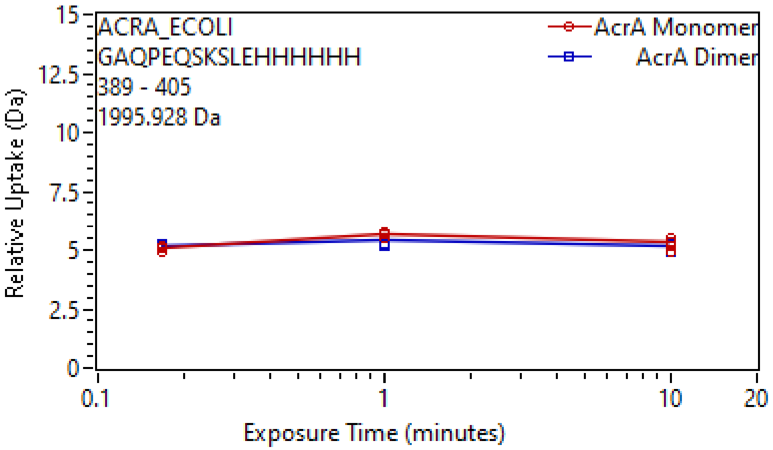
ACRA_ECOLI
GAQPEQSKSLEHHHHHH
389 - 405
1995.928 Da
AcrA Monomer
AcrA Dimer
Relative Uptake (Da)
Exposure Time (minutes)

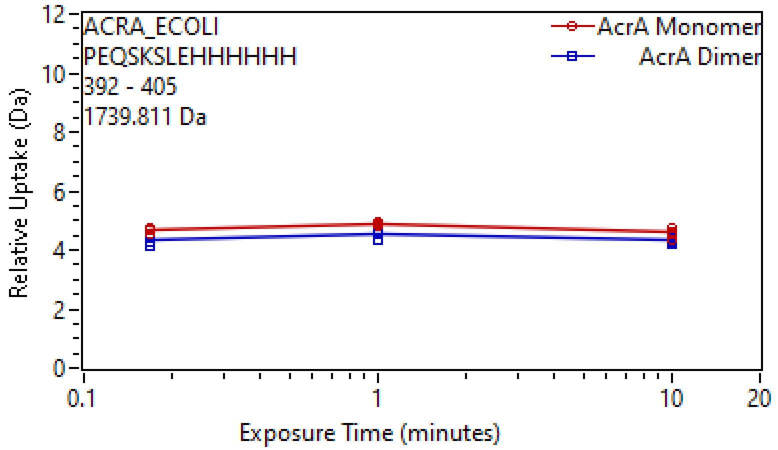
ACRA_ECOLI
PEQSKSLEHHHHHH
392 - 405
1739.811 Da
AcrA Monomer
AcrA Dimer
Relative Uptake (Da)
Exposure Time (minutes)
